# Organellomics: AI-driven deep organellar phenotyping reveals novel ALS mechanisms in human neurons

**DOI:** 10.1101/2024.01.31.572110

**Authors:** Sagy Krispin, Welmoed van Zuiden, Yehuda M. Danino, Noam Rudberg, Lena Molitor, Bar Abarbanel, Gal Aviram, Gili Wolf, Chen Bar, Aviad Siani, Thea Meimoun, Alyssa N. Coyne, Marianita Santiana, Cory A. Weller, Mark R Cookson, Michael E Ward, Fergal M Waldron, Jenna M Gregory, Tal Fisher, Aharon Nachshon, Noam Stern-Ginossar, Nancy Yacovzada, Eran Hornstein

## Abstract

Systematic assessment of organelle architectures, termed the *organellome*, offers valuable insights into cellular states and pathomechanisms, but remains largely uncharted. Here, we present a deep phenotypic learning based on vision transformers, resulting in the Neuronal Organellomics Vision Atlas (NOVA) model that studies confocal images of more than 30 markers of distinct membrane-bound and membraneless organelles in 11.5 million images of human neurons. Organellomics analysis quantifies perturbation-induced changes in organelle localization and morphology using a rigorous mixed-effects meta-analytic framework that accounts for sampling variance and experimental heterogeneity. Applying this approach, we delineate phenotypic alterations in neurons carrying ALS-associated mutations and uncover a physical and functional crosstalk between cytoplasmic mislocalized TDP-43, a hallmark of ALS, and processing bodies (P-bodies), membraneless organelles regulating mRNA stability. These findings are validated in patient-derived neurons and human neuropathology. NOVA establishes a scalable framework for quantitative mapping of subcellular phenotypes and provides a new avenue for investigating the neurocellular basis of disease.

## Introduction

Organellomics, the systematic study of subcellular organellar architecture (the organellome), may be a useful strategy to directly phenotype cells by their subcellular compartments. This is conceivable since the quantification of RNAs, lipids, metabolites and proteins that compose organelles cannot be directly translated into complex organelle properties. Furthermore, organelles are interconnected by functional crosstalk and inter-organelle interdependency ^1,2^, further supporting the value of organellomics-level insights for cell biology.

The analysis of multiple organelles poses substantial challenges due to the diversity of their localizations and morphologies, which must be integrated and evaluated in parallel. Systematic quantification of organellar proteomic content using mass spectrometry–based approaches is highly informative ^3–10^. However, mass spectrometry does not provide spatial or morphological context and depends on designated biochemical specializations. In addition, imaging-based approaches have been developed ^11–23^; however, these are largely limited to non-polarized cultured cells.

The study of cellular organization has been traditionally focused on membrane-restricted organelles, such as the nucleus and mitochondria. However, in recent years, certain biomolecular condensates have been identified as membraneless organelles with specialized functions ^24–28^. Therefore, the study of both membrane-restricted and membraneless organelles is important for gaining a comprehensive view of cellular biology.

The dysfunction of organelles is a central driver of disease. Disorders of membrane-bound organelles such as mitochondria, lysosomes, or autophagosomes have been comprehensively characterized and are associated with distinctive clinical pictures ^29–34^. In addition, compelling evidence suggests a role for membraneless organellar dysfunction in neurodegeneration ^28,35,36^.

We hypothesized that comprehensively capturing the localization and morphological patterns across the entire organellome would allow unbiased mapping of organellar responses to perturbations using a unified scale. To test this hypothesis, we employed an image representation learning approach, particularly based on Vision Transformers (ViT ^37,38^) tailored for perturbation learning ^21,39–45^, to capture high-level abstraction features from fluorescent confocal images. Deep representation learning enables automatic feature extraction, which is unbiased and does not require a pre-defined object or task. Hence it describes any localization and morphological pattern in the image, surpassing widely used approaches to extract objects from images based on a-priori assumptions ^13,46,47^.

ViTs have emerged as powerful models for analyzing microscopy images. Unlike traditional convolutional neural networks (CNNs), ViTs offer enhanced flexibility in capturing global relationships within images by processing the input as a sequence of image patches, and leveraging self-attention to compute dependencies between patches ^37^. This capability is crucial for identifying minute, yet meaningful, alterations in neuronal structure and organization that can arise from specific perturbations. This characteristic makes ViTs well-suited for complex tasks such as deep phenotypic learning in diverse cellular environments, where quantification of spatial relationships of subtle organellar variations is critical.

Here, we introduce Neuronal Organellomics Vision Atlas (NOVA). The contrastive learning framework employed to minimize natural variation, forced the model to focus on genuine perturbation effects through learning transitive relationships between conditions. NOVA takes as input immunofluorescence images of a given cellular organelle in cultured cells and quantifies perturbational changes. NOVA was first pre-trained on massive amounts of OpenCell data ^48^ to learn to encode for subcellular localization patterns of 1,311 proteins. Then, the pre-trained model was fine-tuned for a perturbation learning task on data from highly polarized human neuronal cells incorporating diverse chemical or disease-relevant genetic perturbations. NOVA neither requires detection of the cellular borders (segmentation) nor a pre-defined task, overcoming a limitation in neuron image analysis, and is sensitive enough to depict organellome changes associated with single nucleotide genetic variants. To ensure reporting of reproducible results, NOVA enables mapping of organellar responses to perturbation by quantifying combined effect across experimental repeats using mixed-effect meta-analysis. Our organellomics approach introduces an AI-powered imaging tool that interrogates neuro-cellular biology by capturing phenotypes defined by organelle localization and morphology. We reveal functional interactions between mislocalized cytoplasmic TDP-43 - a hallmark of ALS - and P-bodies, which regulate mRNA stability, validated in patient-derived neurons. This approach offers the foundation for future hypothesis generation in neurobiology and proposes new biomarkers for human pathology.

## Results

### Generation of comprehensive imaging datasets to study the organellome of human neurons

The complexity of the organellome necessitates a modeling strategy that inspects unbiased, quantitative data of multiple organelles. Here, we devised a set of 25 markers for key membrane-bound and membraneless organelles (**Figure 1A** and **Table 1**). Images of neurons with specific dyes or immune-staining (image sites or ‘sites’) were captured by a spinning disc confocal microscope and subjected to QC and preprocessing steps, resulting in single cell tile images (1.5 neurons per tile on average, **Supplementary Figure 2** and **Supplementary Materials**). A comprehensive training dataset was generated, composed of ∼3.2 million image tiles of human iPSC-derived neurons, under different experimental conditions (i.e., 1 chemical, 7 genetic perturbations, and WT; **Supplementary Figure 1**). This data was used for model training (2.24M images), validation (0.48M images), and test (0.48M images). The training dataset was generated from two independent biological differentiations (referred to as “experimental repeats”), with each differentiation including two to eight technical repeats per condition (**Supplementary Figure 1**).

**Figure 1.**
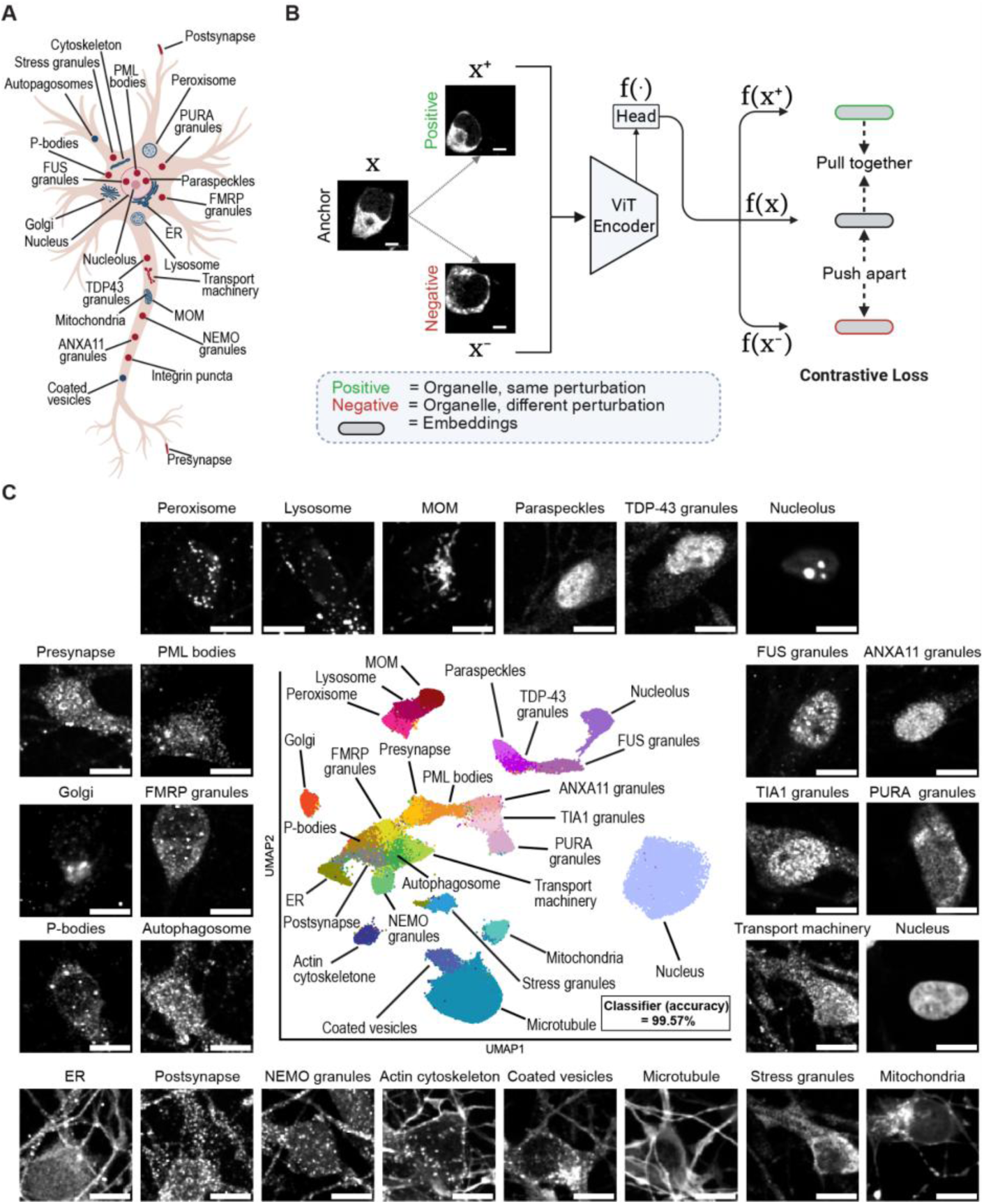
Vision transformer-based perturbation learning model enables organellomic studies in neurons. **(A)** Diagram of a neuron with the 25 organelles used for sequential fine-tuned training in NOVA. **(B)** Perturbation learning workflow with contrastive learning. An input image (anchor) is encoded into 192 latent features (embeddings). Each anchor is randomly matched with other images of the same organelle under the same perturbation (“positives,” similar effects) and with images of the same organelle under different perturbations (“negatives,” dissimilar or no effect). During training, the model converges to establish meaningful representations via contrastive loss: pulling together positives in feature space, while pushing apart negatives based on the distinct differences learned from the data. Perturbations with no apparent effect result in feature representations similar to unperturbed cells. To prevent the model overfitting to potential experimental repeat or well effects, positives and negatives were sampled from different experimental repeats and wells (**Supplementary Figure 6** and **Methods** for details). **(C)** UMAP visualization of jointly projected three experimental repeats, showing the organellar landscape of day 8 human iPSC-derived neurons under basal growth conditions. 26 organelles form distinct topographic patterns (classifier accuracy 99.57%; see **Supplementary Figure 7** and **Methods**). ∼6k image tiles per organelle. Representative images displayed. Scale bar, 5 µm.

**Table 1.** Key 26 organelles characterized in this study, respective abbreviations and markers.

| Organelle | Abbr. | Marker | Literature |
| --- | --- | --- | --- |
| Nucleus |  | Hoechst 33342 | 65 |
| Mitochondria |  | Mitotracker | 66 |
| Mitochondria Outer Membrane | MOM | TOMM20 | 67 |
| Golgi Apparatus | Golgi | GM130 | 68 |
| Endoplasmic Reticulum | ER | Calreticulin | 69 |
| Actin Cytoskeleton | Cytoskeleton | Phalloidin | 70 |
| Microtubule Cytoskeleton | Microtubule | Tubulin or TUJ1 (beta Tubulin 3) | 71–73 |
| Autophagosomes |  | SQSTM1 | 74 |
| Lysosomes |  | LAMP1 | 75 |
| Peroxisomes |  | PEX14 | 76 |
| Coated Vesicles |  | CLTC | 77 |
| Nucleolus |  | NCL | 78 |
| Paraspeckles |  | NONO | 79,80 |
| TDP-43 granules |  | TDP-43 | 81 |
| FUS granules |  | FUS | 82 |
| Promyelocytic Leukaemia Nuclear Bodies | PML bodies | PML | 83 |
| P-Bodies |  | DCP1A or LSM14A | 84,85 |
| Stress granules |  | G3BP1 | 86 |
| ANXA11 granules |  | ANXA11 | 87 |
| PURA granules |  | PURA | 88,89 |
| NEMO granules |  | NEMO | 57 |
| Microtubule-Associated Transport Machinery | Transport machinery | KIF5A | 90 |
| Presynaptic Terminals | Presynapse | SNCA | 91,92 |
| Post-Synaptic Subcompartments | Postsynapse | PSD95 | 93 |
| FMRP granules |  | FMRP | 94 |
| Nuclear speckles |  | SON | 95 |

All 6 independent datasets described below, encompassing a total of ∼11.56M tile images, of which ∼2.55M were used solely for inference (held-out from training). Furthermore, these datasets originated from different experiments in different labs, spanned different cell lines, and included unseen organelles (**Supplementary Materials**).

### ViT for encoding organelle localization patterns in perturbed human neurons

Vision Transformers (ViTs), with their self-attention mechanism, excel at modeling long-range spatial relationships across image patches, making them particularly effective for capturing fine-grained cellular details in microscopy images ^37^. This is particularly valuable for neurons, whose morphology is far more complex than that of simple cells. Neurons extend processes over large spatial scales, with organelles distributed across axons, dendrites, and soma. Thus, we decided to develop a VIT model to recognize intricate localization patterns and understand perturbation microscopic effects.

To examine organellar changes in neuronal cells upon disease-associated perturbations, we developed NOVA, a sequential fine-tuned model based on ViTs and contrastive learning (**Figure 1B**). First, we trained a ViT model with a multi-class classification head and cross-entropy loss on a publicly available dataset of 1,311 OpenCell proteins in human embryonic kidney cells that were cultured in basal growth conditions (HEK293T, 1,100,253 image tiles^48^). By learning to predict the protein identity in the image (loss curves in **Supplementary Figure 3A**), the model learns to encode a wide variety of organellar morphologies and localizations (referred here as topographies), achieving mean accuracy of 87% in identification of organelle localization categories (ROC AUC curves, confusion matrix and UMAP visualization of localization categories in **Supplementary Figure 3B-D**).

We next extended the model to perform a perturbation learning task by continuing the training with images of neurons with perturbations (**Methods**). To retain the knowledge learned from the localization encoding task, a partial fine-tuning step was performed, using an angle-metric ^49^ optimization approach on the model’s layers (**Supplementary Figure 4** and **Methods**). Specifically, the angle-metric quantified how much each layer changed between the localization and perturbation tasks. Layers showing minimal deviation were kept frozen to preserve general knowledge, while layers with deviations larger than the median were retrained to adapt the model to neuron-specific and perturbation-related features.

The perturbation learning head and loss function were devised to perform contrastive learning, using Information Noise Contrastive Estimation loss (InfoNCE, **Figure 1B** and **Supplementary Figures 5**, **6**). InfoNCE loss encourages the model to implicitly learn to focus on similarities or dissimilarities that correspond to true perturbation effects. Hence, impactful perturbations are pushed apart in the latent space, diverging their feature representations away from the baseline, control condition. In parallel, perturbations without effect generated similar representations, ensuring that the model doesn’t overfit irrelevant non-biological differences or technical artifacts. Together, NOVA performs two tasks: (1) the encoding of neuron images for feature representations, and (2) detecting true perturbation effects.

### Full neuronal organelle landscape in unperturbed neurons

As an initial characterization of the neuronal organellome, we sought to establish proof of feasibility for the identification of organelles in the basal, unperturbed state. Therefore, we performed inference on images from three experimental repeats of unperturbed neurons acquired in the NIH CARD (using a Ti2 Spinning Disk confocal microscope, 8 technical repeats per experimental repeat, 296,776 neuron images post QC). 26 organelles were analyzed, including microtubules (TUJ1) and TIA1 granules, two organelles not previously seen by the model in the training set.

Images of unperturbed neurons were encoded using NOVA, and the three experimental repeats were jointly projected into a single Uniform Manifold Approximation and Projection (UMAP)^50^ plot (**Figure 1C**). The full neuronal organelle landscape revealed distinct clusters corresponding to most organelles, indicating that the learned embeddings capture organelle-specific features, and that experimental repeat effects did not dominate the latent space. Furthermore, the resulting image representations were evaluated with a post-hoc classifier for neuronal organelle identification (**Methods**) and achieved 99.57% accuracy in correctly calling each of the 26 organelle averaged across all experimental repeats (**Supplementary Figure 7** and **Methods**).

We benchmarked neuronal organelle identification by comparing performance of a classifier trained on representations extracted by NOVA versus Cytoself model^11^ or CellProfiler^13,47^ features. The classifier trained on NOVA’s embeddings achieved the highest sensitivity performance (sensitivity 90.26%, accuracy 99.57%, **Figure 1C**), outperforming Cytoself (sensitivity 83.66%, accuracy 99.28%) and CellProfiler (sensitivity 79.88%, accuracy 98.62%). Organellar maps generated with Cytoself or CellProfiler appeared more diffuse than those from NOVA (**Supplementary Figure 8**), likely because NOVA has learned images of neurons and is trained to minimize experimental repeat-related variability.

To quantify relationships between organelles, we computed the pairwise Euclidean distances between organelles, using the full latent representations, and rescaled by min-max normalization to [0,1] (distance noted as *d*, **Supplementary Figure 9**). Small distances in the latent space indicate that the model has captured organelle similarities in their morphology (size, shape, texture) or spatial context (co-occurrence within the same subcellular regions).

As expected, pairwise distance analysis revealed neighborhoods of organelles with evident physical and/or functional crosstalk. Mitochondria-outer-membrane (MOM), peroxisomes, and lysosomes clustered closely, with a mean intra-community distance of *d*=0.21, compared to a mean inter-community distance of *d*=0.74 (**Supplementary Figure 9**). This is consistent with known communication through membrane contact sites, shared functions and linked dysfunction in Parkinson’s disease^51–56^. Likewise, the cytoskeleton-associated organelles, actin cytoskeleton and microtubules, presented an intra-community distance of *d*=0.22, as also apparent in the UMAP projection (**Figure 1C**). Unexpectedly, Mitotracker-labeled mitochondria, which preferentially mark active mitochondria, were distant from MOM (*d* = 0.8, **Supplementary Figure 9**), likely reflecting functional or structural differences between these subcompartments which are of biological interest. We note that one of the lowest pairwise distances (*d* = 0.11) was between NEMO granules, essential for NF-κB signaling^57^, and the postsynapse (PSD-95; **Supplementary Figure 9**), suggesting potential new functional links. Finally, the low pairwise distances of TDP-43 granules to paraspeckles (*d* = 0.22) and ANXA11 granules (*d* = 0.27) align with reported interactions in ALS and FTD^58–64^. Taken together, the relative concordance between multiple experimental repeats UMAP and latent-space distances supports the robustness of NOVA, whose embedding space captures organelle neighborhoods and reveals potential new biological relationships.

### A ranking system for quantification of the organellar response across experimental repeats

For each perturbation under examination during inference, the NOVA-derived embeddings of control images versus perturbed are utilized to estimate an effect size for each organelle across experimental repeats (biological differentiations or human subjects). Effect size is designed to capture the degree to which a perturbation shifts organelle topography from the baseline state, and is estimated for each combination of (perturbation × organelle × experimental repeat). Then, effect sizes are combined in mixed-effects meta-analytic analysis to estimate the combined effect size (denoted *μ̂*), while accounting for heterogeneity across experimental repeats **(Figure 2A, Methods)**. Latent space distances provide a single-metric quantification of the response to perturbations. This approach models both the sampling variance within repeats (uncertainty) and the heterogeneity across repeats (random effect), allowing to statistically test whether an organelle is consistently affected (reproducibility). Effects are presented in organelle scoring forest plots; organelles are ranked based on their combined effect size and multiple hypothesis correction is applied.

**Figure 2.**
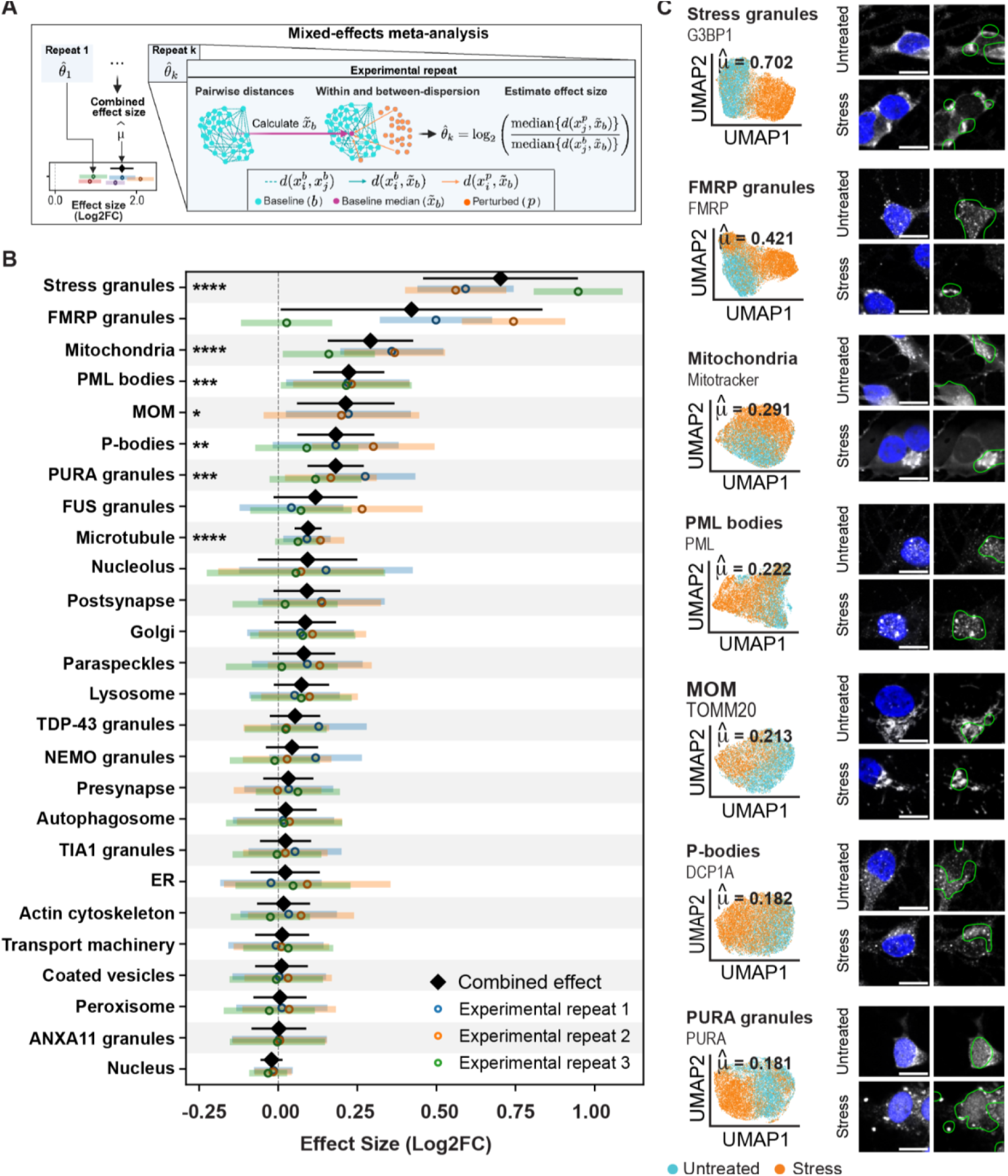
Neuronal organelle response to oxidative stress in human neurons. **(A)** Overview of meta-analytic framework for perturbation effect size estimation. Effect size is defined as the shift in organelle topography (localization/morphology) under perturbation, quantified as the distance in NOVA embedding space between perturbed and baseline states. Effect sizes are computed independently per experimental repeat and integrated using mixed-effects meta-analysis, accounting for within- and between-repeat variance, yielding a combined estimate with confidence intervals that reflect biological and technical variability. **(B)** Stacked forest plot of organelle effect size (log2 fold change) in day 8 neurons in response to oxidative stress (NaAsO₂, 0.25 mM, 30 min), relative to untreated controls. Three experimental repeats. Colored dots represent the experimental repeat-wise effect estimate, with horizontal bars indicating the 95% CI variance within experimental repeats. The combined effect estimate (*μ̂*) is shown in black diamond with its 95% meta-analytic CI, which accounts for variance between experimental repeats when present. Adjusted p-value: * < 0.05, ** < 0.01, *** < 0.001, **** < 0.0001. ∼800-2800 image tiles per condition/experimental repeat. **(C)** Joint UMAP projection of three experimental repeats, highlighting top organelles by combined effect size and FDR-corrected meta-analysis significance. Point - single image tile, ∼7k images/condition. Untreated (blue) vs oxidative stress (orange). *μ̂*- the combined effect estimate. Representative images with nucleus (hoechst 33342) and overlaid model’s attention maps. Lens ×63; scale bar, 10 µm.

To evaluate the ability of NOVA to detect organellar changes in response to perturbations we introduced sodium arsenite (0.5 mM, 30 minutes)^96,97^, which leads to oxidative stress, into day 8 human iPSC-derived neurons (three experimental repeats, >590,600 images; **Supplementary Figure 10**). A mixed-effects meta-analysis, accounting for experimental repeat reproducibility, revealed substantial alterations in the topography of multiple organelles (**Figure 2B, C**). The largest combined effect size was of stress granules (SGs), cytoplasmic biomolecular condensates that are composed of RNA-binding proteins and RNAs that are formed in response to stress (marked by G3BP1; *μ̂*=0.703, adj. p=2.22 × 10⁻⁷). A similar effect was observed for FMRP, which is localized in neurons into RNA-associated granules (“FMRP granules”), including SGs under stress^94^. However, strong heterogeneity driven by a single divergent experimental repeat limited statistical significance after FDR correction (*μ̂*=0.421, adj. p=0.074; heterogeneity: I²=95.45, τ²=0.126 (95% CI=[0.024,2.33]); **Figure 2B, C**). In addition, PURA granules effectively discriminated stressed from untreated neurons (*μ̂*=0.182, adj. p=1.98 × 10⁻^4^, **Figure 2B, C**), consistent with their association with SGs^88,89^. Reproducible and significant stress-associated changes were also observed in the mitochondria (*μ̂*= 0.292, adj. p=9.87 × 10⁻^5^), MOM (*μ̂* = 0.213, adj. p=0.012), PML bodies (*μ̂*=0.223, adj. p=2.6e × 10⁻^4^), P-bodies (*μ̂*=0.182, adj. p=0.007), and microtubules (*μ*=0.095, adj. p=9.35 × 10⁻^5^; **Figure 2B, C**). We further implemented NOVA rollout attention maps to visualize image regions that most strongly influenced the model’s output, highlighting areas of highest attention and enabling interpretation of the features driving NOVA’s output (**Figure 2C, Methods**).

Altogether, we comprehensively characterize the changes that neuronal organelles undergo in response to oxidative stress, using a single-scale system. Organelle scoring is a unified system for organellar comparison that reassuringly detects known markers of the neuronal stress response and provides an overview of significant, reproducible perturbation effects on organelles across multiple experimental repeats.

### An organellar response to ALS-associated single-nucleotide variants in neuron

The iPSC Neurodegenerative Disease Initiative (iNDI)^98^ enables controlled comparison of different neurodegeneration-associated variants within a single isogenic background. We quantified the effects of ALS-associated genetic variants on neuronal organelles in iPSC-derived iNDI neurons. The ancestral isogenic control line (Knockout Laboratory F2.1 Jackson, KOLF, wild-type) was compared to four ALS-associated variants: *FUS* (Fused in Sarcoma) R495*^99,100^, *TARDBP* (TAR DNA/RNA-Binding Protein 43; TDP-43) M337V^101,102^, *OPTN* (Optineurin) Q398E^103^, and *TBK1* (TANK Binding Kinase 1) E696K^104^ (**Figure 3A**). We generated a neuronal imaging dataset consisting of six experimental repeats with 2 technical repeats each (∼6,495,000 images post-QC, representing ∼1,250,000 neurons captured in 2–4 channels; average 1.7 nuclei per tile; **Supplementary Figure 11** and **Supplementary Materials**). Neurons were stained with the 26 organelle markers initially used (**Table 1**) and three other markers that were used for inference only (SON (nuclear speckles), hnRNPA1 (hnRNP complex), and LSM14A (P-bodies).

**Figure 3.**
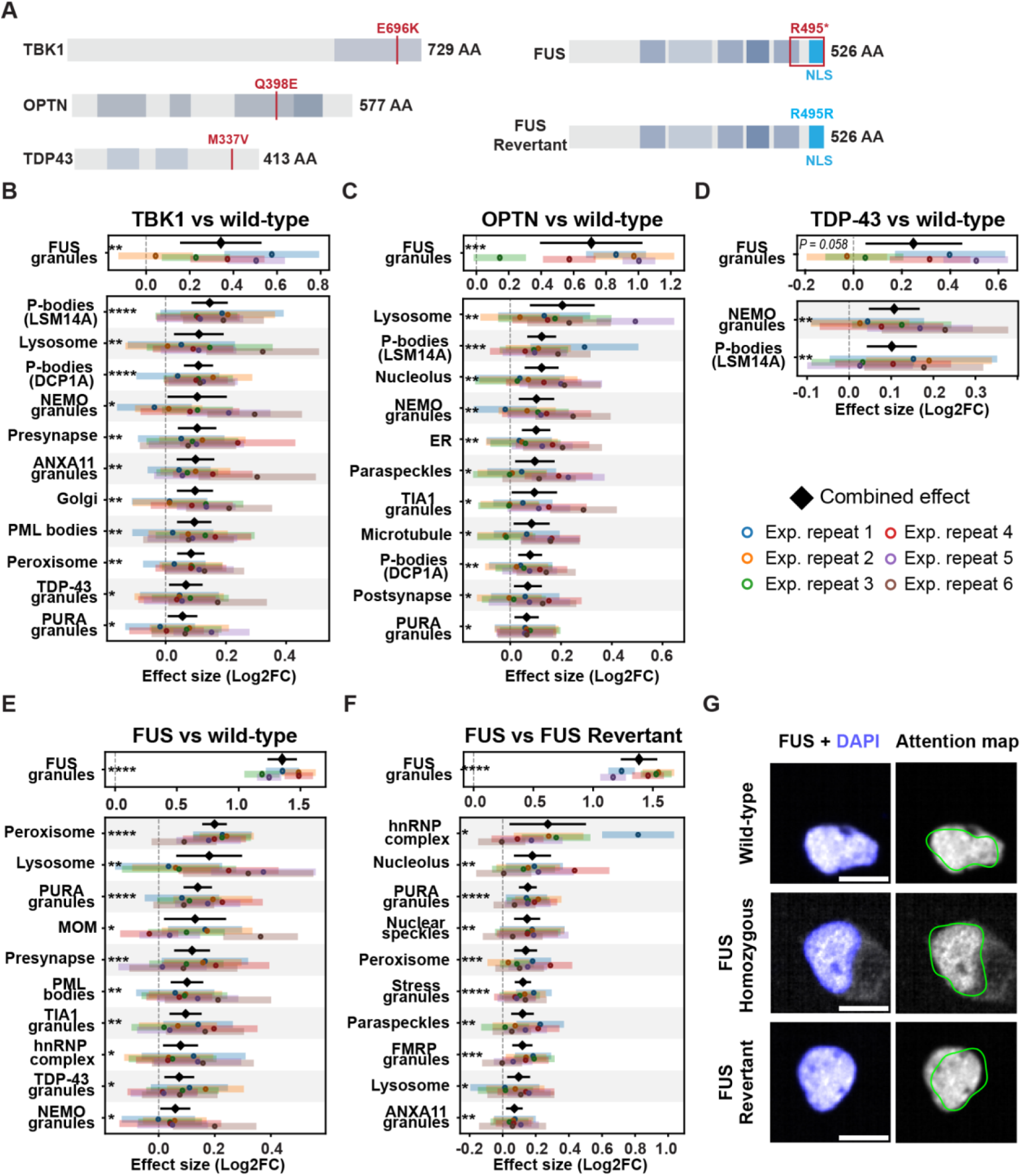
Neuronal organelle response to ALS-associated variants. **(A)** ALS-associated variants and genetically-corrected revertant for FUS variant (from stop codon at amino acid 495 to arginine; *495R) from the iPSC Neurodegenerative Disease Initiative (iNDI) ^98^. **(B-F)** Stacked forest plot of organelle effect size (log2 fold change) in day 8 neurons harboring homozygous variants of **(B)** *TBK1*, **(C)** *OPTN*, **(D)** *TARDBP* or **(E)** *FUS*, relative to wild-type control. **(F)** *FUS* homozygous variant relative to *FUS* Revertant line. Six experimental repeats. 29 organelles studied. Only significant organelles are shown. Colored dots represent the experimental repeat-wise effect estimate, with horizontal bars indicating the 95% CI variance within experimental repeats. The combined effect estimate (*μ̂*) is shown in black diamond with its 95% meta-analytic CI, which accounts for variance between experimental repeats when present. Adjusted p-value: * < 0.05, ** < 0.01, *** < 0.001, **** < 0.0001. ∼620-6800 image tiles per organelle/variant/experimental repeat. **(G)** Representative images of *FUS* allelic series with nucleus (hoechst 33342) and overlaid model’s attention maps. Lens ×63; scale bar, 10 µm

Inference on the iNDI induced neurons dataset revealed diverse and significant organellar changes in ALS-associated variants relative to wild-type. Overall, a total of 19 organelles were affected in ALS-associated lines, relative to wild-type (WT) including FUS granules, lysosome, peroxisome, PURA granules, NEMO granules, presynapse, PML bodies, TIA1 granules, P-bodies (LSM14A, DCP1A), MOM, hnRNP complex (HNRNPA1), TDP-43 granules, ANXA11 granules, Golgi, ER, paraspeckles, microtubule (Tubulin), and postsynapse (**Figure 3B-F**). The effect size of FUS granules was the largest across all lines (*OPTN* Q398E (*μ̂*=0.709, adj.p=1.17 × 10⁻^4^; heterogeneity: I²=95.41%, τ²=0.1217 (95% CI=[0.035,0.7738]))), *TBK1* E696K (*μ̂*=0.342, adj.p=1.1 × 10⁻^3^; heterogeneity: I²=83.47%, τ²=0.0377 (95% CI=[0.0068,0.264])), *TARDBP* M337V (*μ̂*=0.247, adj.p=5.9 × 10⁻2; heterogeneity: I²=88.1%, τ²=0.045 (95% CI=[0.01,0.311]; *FUS* R495* *μ̂*=1.352, adj. p=3.22 × 10⁻^15^; heterogeneity: I²=78.25%, τ²=0.0142 (95% CI=[0.0019,0.1059]) **Figure 3B-E** and **Supplementary Figure 12**).

We also compared *FUS* 495* to the revertant line (REV), in which the variant had been corrected by genome editing. The restoration of the wild-type *FUS* gene sequence was associated with relocalization of the FUS protein into the nucleus (FUS granules,REV:*μ̂*=1.385, adj.p=6.4 × 10⁻^15^; heterogeneity: I²=87.96%, τ²=0.0266 (95% CI=[0.0062,0.1801]); **Figure 3F,G**). Since the *FUS* 495* variant disrupts FUS’s steady-state nuclear localization and thereby reduces nuclear signal intensity, the model’s nuclear focus might suggest that it utilizes this loss of nuclear enrichment to distinguish *FUS* 495* from the wild-type and revertant lines. Several other organelles showed significant alterations in *FUS* R495* relative to both the wild-type (WT) and revertant (REV), namely, PURA granules (WT: *μ̂*=0.139, adj.p=3.4 × 10⁻^7^, REV: *μ̂*=0.152, adj.p=2.5 × 10⁻^7^; **Figure 3E, F**), Peroxisomes (WT: *μ̂*=0.199, adj.p=3.2 × 10⁻^15^, REV: *μ̂*=0.139, adj.p=2.9 × 10⁻^4^), lysosome (WT: *μ̂*=0.18, adj.p=5.44 × 10⁻^3^, REV: *μ̂*=0.09, adj.p=9.05 × 10⁻^3^) and hnRNP complex (WT: *μ̂*=0.078, adj.p=2.25 × 10⁻^2^ REV: *μ̂*=0.27, adj.p=2.72 × 10⁻^2^; heterogeneity: I²=88.33%, τ²=0.074 (95% CI=[0.02,0.384])). These data underscore a robust and reproducible modification across two different controls. Comparable analysis of the *FUS* R495* heterozygous variant, is described in **Supplementary Figure 13**.

Additional organelles were altered across multiple lines: P-bodies, marked by LSM14A, altered in response to variants *TBK1* E696K (*μ̂*=0.146, adj.p=2.8 × 10⁻^5^), *OPTN* Q398E (*μ̂*=0.124, adj.p=1.2 × 10⁻^4^) and *TARDBP* M337V (*μ̂*=0.101, adj.p=5.8 × 10⁻^3^; **Figure 3B-D**)), while DCP1A was also impacted in both *TBK1* E696K (*μ̂*=0.109, adj.p=7 × 10⁻^5^) and *OPTN* Q398E (*μ̂*=0.078, adj.p=3 × 10⁻^3^); **Figure 3B-D** and **Supplementary Figure 14**). Additionally, lysosomes were affected by variants in *TBK1* E696K (*μ̂*=0.11, adj.p=1.3 × 10⁻^2^), *OPTN* Q398E (*μ̂*=0.205, adj.p=3.8 × 10⁻^3^; heterogeneity: I²=74.22%, τ²=0.0179 (95% CI=[0.0028, 0.1143])) and *FUS* R495* (*μ̂*=0.18, adj.p=5.4 × 10⁻^3^; (**Figure 3B-D and Supplementary Figure 15**). NEMO granules altered by variants in *TARDBP* M337V (*μ*=0.107, adj.p=5.9 × 10⁻^3^), *OPTN* Q398E (*μ*=0.103, adj.p=7.2 × 10⁻^3^) and *TBK1* E696K (*μ̂*=0.105, adj.p=0.044; **Figure 3B-D**). Notably, TDP-43 granules were unaffected by any of the variants, consistent with the absence of TDP-43 proteinopathy in premature 8-day-old neurons ^105^.

Altogether, NOVA detected subtle yet reproducible changes associated with single nucleotide changes, uncovering multiple organelles affected across different ALS-associated mutations, including FUS granules and P-bodies. Together, these results provide a comprehensive view of shared and distinct organellar alterations, underscoring the complexity of diverse organelles in ALS.

### Cytoplasmic TDP-43 interacts with P-bodies and affect their functions

We next focused on TDP-43 cytoplasmic mislocalization, a hallmark of ALS and frontotemporal dementia (FTD) pathology ^106–108^. To this end, we used human iPSC-derived neurons overexpressing doxycycline-inducible TDP-43 (*TARDBP*) lacking a functional nuclear localization signal (NLS; TDP-43^ΔNLS^), which is therefore restricted to the cytoplasm **(Figure 4A)**. Notably, TDP-43^ΔNLS^ expression was minimal without doxycycline and, upon induction, reached ectopic levels ∼3-fold higher than endogenous TDP-43 (**** < 0.0001, One-way ANOVA with Benferroni’s multiple comparison test, **Supplementary Figure 16**). Induction of TDP-43^ΔNLS^ by doxycycline (+DOX) increased cytoplasmic TDP-43 relative to nuclear TDP-43 by ∼10-fold compared to the uninduced condition (-DOX; *β* = 0.55, p < 0.0001, (95% CI [0.54, 0.56]), fixed-effect fallback used, **Figure 4B**).

**Figure 4.**
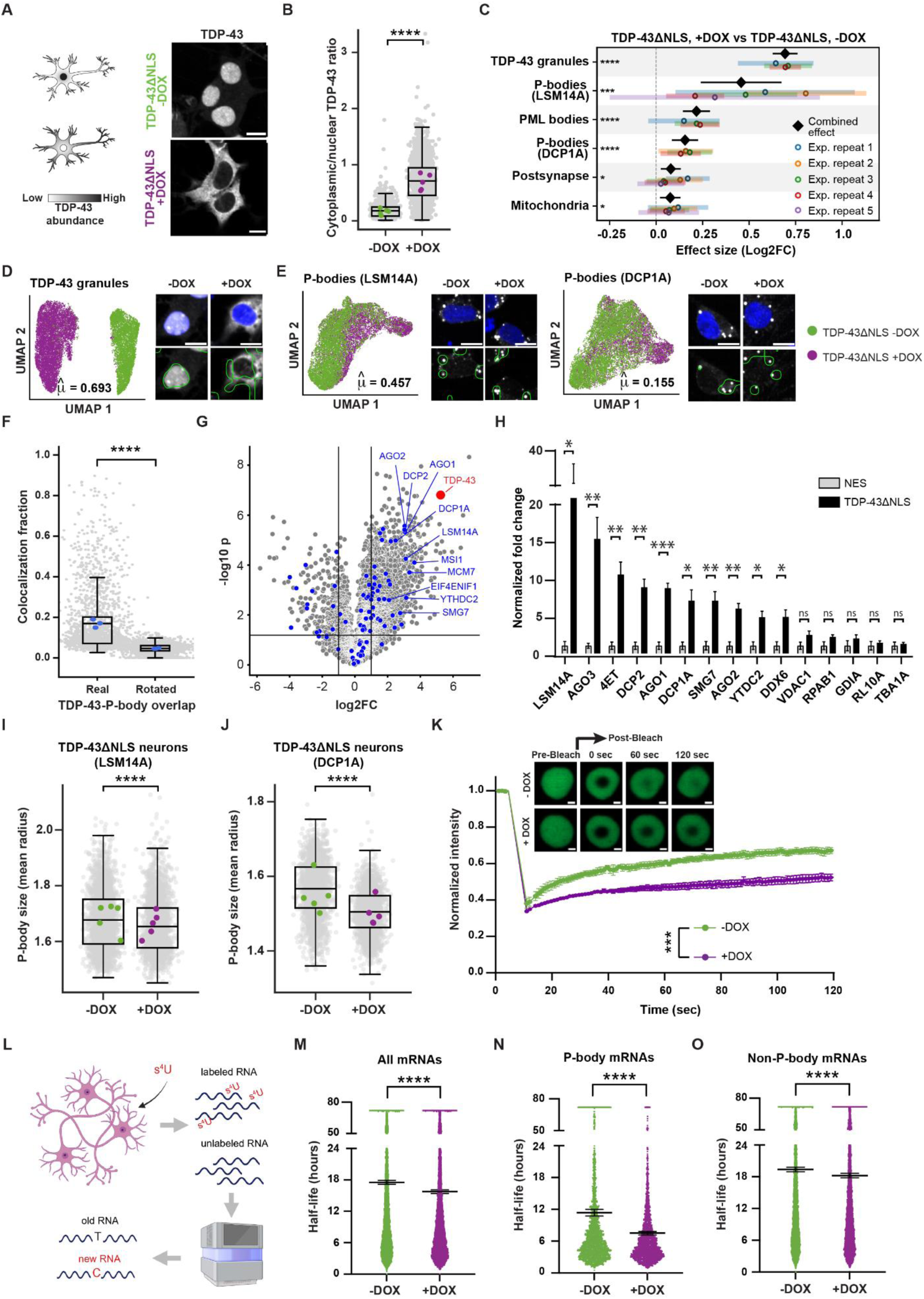
Cytoplasmic TDP-43 overexpression interacts with neuronal P-bodies, altering their function. **(A)** Diagram and representative images of day 8 neurons expressing doxycycline-inducible TDP-43 with a mutated nuclear localization signal (TDP-43^ΔNLS^; -DOX uninduced, +DOX induced). Lens ×63; scale bar, 5 µm. **(B)** Boxplots of cytoplasmic-to-nuclear TDP-43 signal ratios in TDP-43^ΔNLS^ neurons ±DOX induction (means = 0.064 (-DOX) vs. 0.615 (+DOX)). Colored dots represent experimental repeat means; gray dots - individual site images. ****p < 0.0001. Linear model with DOX as a fixed effect and experimental repeat as a covariate; random-effect variance estimated as 0. **(C)** Stacked forest plot of organelle effect sizes (log2 fold change) in day 8 TDP-43^ΔNLS^ neurons +DOX, relative to -DOX. Five experimental repeats. 29 organelles. Colored dots represent the experimental repeat effect estimate, with horizontal bars indicating the 95% CI within-experimental repeat variance. The combined effect estimate (*μ̂*) is shown in black diamond with its 95% meta-analytic CI, which accounts for between-experimental repeat variance when present. Adjusted p-value: * < 0.05, ** < 0.01, *** < 0.001, **** < 0.0001. Only significant organelles are shown. ∼280-6200 image tiles per organelle/condition/experimental repeat. Joint UMAP projection of five experimental repeats showing **(D)** TDP-43 granules (image tiles: 12,530 +DOX; ∼8,500 -DOX) and **(E)** P-bodies stained with LSM14A (left, image tiles: 9,510 +DOX; ∼8,580 -DOX) or DCP1A (right, image tiles: ∼8,610 +DOX; ∼9,690 -DOX). Neurons -DOX/+DOX (green/ purple). Points - single image tiles. *μ̂*- the combined effect estimate. Representative images with nucleus (hoechst 33342) or overlayed model’s attention maps. Lens ×63; scale bar, 10 µm. **(F)** Boxplots of P-body-positive (DCP1A) pixels that are also positive for TDP-43 (V5 tag). Colocalization fraction in real images vs. randomized (rotated) images (mean fraction overlap = 0.170 vs. 0.047, respectively). ∼1,800 site images; Wilcoxon signed-rank test, ****p < 0.0001. Blue dots - condition means; Gray dots - individual images; three experimental repeats of day 8 induced neurons. **(G)** Volcano plots of relative protein levels in induced APEX-TDP-43^ΔNLS^ relative to APEX-NES samples (x axis log2 scale), analyzed by mass spectrometry. y axis depicts the differential expression p values (−log10 scale). One of two independent experiments; 3-4 technical repeats per condition. TDP-43 (red) and selected P-body proteins (from ref.^116^, blue). Unpaired two-sided t-test. **(H)** Normalized enrichment of P-body proteins (AGO1, AGO2, DCP1A, DCP2, EIF4ENIF1, LSM14A, MCM7, MSI1, SMG7, YTHDC2) and housekeeping proteins (VDAC1, RPAB1, GDIA, RL10A, TBA1A) in the vicinity of TDP-43^ΔNLS^ vs. NES control, analyzed by targeted mass spectrometry. Unpaired two-sided t-test. CellProfiler quantification of P-body size in monoclonal TDP-43^ΔNLS^ neurons (-DOX vs. +DOX) stained by P-body markers: **(I)** LSM14A or **(J)** DCP1A (means (-DOX vs. +DOX) = 1.70 vs. 1.68 or 1.51 vs. 1.46, respectively). Colored dots - experimental repeat means; gray dots - site-level measurements. Five experimental repeats per condition. Linear mixed-effects model with DOX as a fixed effect and experimental repeat as a random intercept; fixed-effects fallback used when variance was zero. ****p < 0.0001. **(K)** Live imaging of fluorescence recovery after photobleaching (FRAP) of YFP-DCP1A in P-bodies. U2OS cells with or without DOX-induced TDP-43^ΔNLS^, normalized to pre-bleaching baseline. Images depict a single P-body; scale bar, 1 µm. Four repeats with 3-5 fields of view and 2-9 P-bodies per field. Two-way ANOVA with repeated measures and Geisser-Greenhouse correction. *** P < 0.001 **(L)** Schematic of the SLAM-seq approach for measuring RNA half-life through incorporation of 4-thiouridine (4sU; 200 μM) into newly transcribed RNA molecules ^130^. GRAND-SLAM^131^ RNA half-life estimates from 0, 2, and 3 h of 4-thiouridine labeling: **(M)** all mRNAs, **(N)** P-body–enriched mRNAs or **(O)** P-body-depleted mRNAs. Dots = individual transcripts; 2–4 technical repeats per timepoint. Mean ± 95% CI. Whiskers - SD. Mann–Whitney U test, P-value: * < 0.05, ** < 0.01, *** < 0.001, **** < 0.0001, ns = not significant.

We generated a dataset of TDP-43^ΔNLS^ neuron images across five experimental repeats (three technical repeats each) stained for 29 organelles (∼1,192,600 images post-QC, **Table 1**, **Supplementary Figure 17**). This line, unseen during training, was stained with the 26 organelle markers initially used (**Table 1**) and three other markers that were used for inference only (SON (nuclear speckles), hnRNPA1 (hnRNP complex), and LSM14A (P-bodies).

As could be expected, the topography of TDP-43 granules was the most affected by the TDP-43^ΔNLS^, relative to uninduced state (*μ̂*=0.693, adj. p=6.4 × 10^-15^; (95% CI=[0.626,0.76]), **Figure 4C, D**). P-bodies and PML nuclear bodies, two principal membraneless organelles that regulate nucleic acids in the cytoplasm and nucleus, respectively, also displayed significant topographic changes (P-bodies (LSM14A): *μ̂*=0.457, adj. p=0.0001; heterogeneity: I²=67.64, τ²=0.032 (95% CI = [0.24, 0.675]); P-bodies (DCP1A): *μ̂*=0.155, adj. p=7.11 × 10^-5^; (95% CI = [0.085, 0.226]); PML nuclear bodies (PML): *μ̂*=0.217, adj. p=3.7 × 10^-8^; (95% CI = [0.144, 0.289]); **Figure 4C,E**).

PML nuclear bodies regulate TDP-43 homeostasis and may accordingly be affected by TDP-43 overexpression ^109,110^. No discernible changes were identified in several other organelles, including stress granules markers (G3BP1, FMRP, and PURA; **Supplementary Figure 18A**), consistent with previous reports that cytosolic TDP-43 assemblies operate separately from conventional stress granules ^111^.

P-body topography changes were depicted with two independent markers (LSM14A and DCP1A). To further investigate the hypothesis that TDP-43 granules interact with P-bodies, we demonstrated that colocalization of P-bodies with TDP-43^ΔNLS^ (measured as the percent of TDP-43^ΔNLS^ signal inside P-bodies; **Supplementary Figure 18B**) was significantly greater in TDP-43^ΔNLS^ neurons than expected by chance under spatial randomization (mean real fraction overlap 0.170 vs. 0.047 in rotated; Wilcoxon signed-rank test, ****p < 0.0001; three experimental repeats, four technical repeats each, ∼200 imaging sites per condition; **Figure 4F**).

Next, using SpeedPPI, a machine-learning algorithm for structure-based protein-protein interaction prediction ^112–114^, we tested whether P-body-enriched proteins are predicted to physically interact with TDP-43. We analyzed proteins that were co-immunoprecipitated with the P-body protein DDX6 ^115^ and were either enriched (n=20 proteins), or depleted (n=18 proteins) in P-bodies by unbiased mass spectrometry ^116^. The predicted interactions of the enriched P-body proteins with TDP-43 were significantly stronger than those of the group of proteins highly depleted from P-bodies (average pDockQ score = 0.3334 vs. 0.2058; p = 0.0112; Mann–Whitney U test; **Supplementary Figure 18C** and **Supplementary Materials**), supporting the specificity of these predicted interactions with TDP-43.

We further characterized the proteome in the vicinity of cytoplasmic TDP-43 using engineered ascorbate peroxidase (APEX) proximity labeling and mass spectrometry. APEX uses hydrogen peroxide (H2O2), to generate reactive biotin radicals that label nearby biomolecules, revealing weak or transient interactions and enabling characterization of cellular compartments at nanometer resolution ^117–119^. An APEX peptide fused to TDP-43^ΔNLS^ or to NES constructs in frame, demonstrated specificity by immunostaining. The biotinylated signal was observed only when both H2O2 and biotin phenol (BP) were present in the medium and overlapped with construct localization **(Supplementary Figure 19A)**. This provides a reliable framework to capture cytoplasmic TDP-43 interactions and enable proteomic mapping of its cytoplasmic environment.

Following proximity labeling, biotinylated proteins were pulled down with streptavidin beads and analyzed by mass spectrometry (MS) in two independent experiments (3-4 technical repeats per condition each). In total, 212 proteins interacted specifically with TDP-43^ΔNLS^ relative to the NES control in both experiments (FDR < 0.01, log2FC > 2, unpaired two-sided t-test). TDP-43 itself was reassuringly among the most enriched proteins (13-fold and 38-fold relative to NES, respectively; **Figure 4G, Supplementary Figure 19B, Supplementary Materials**). Gene Ontology (GO) cellular component analysis ^120,121^ revealed strong enrichment of “P-body” proteins (fold enrichment = 4.32, FDR < 0.01; **Supplementary Figure 19C**) as well as RNA-induced silencing complex (RISC)/RNAi effector complex proteins, which often localize to P-bodies and share overlapping functions in mRNA silencing ^122,123^. We further validated 10 P-body-enriched proteins ^116^ (AGO1, AGO2, DCP1A, DCP2, EIF4ENIF1, LSM14A, MCM7, MSI1, SMG7, YTHDC2) as TDP-43^ΔNLS^ interactors in an independent targeted MS experiment, whereas five housekeeping proteins (obtained from the proteomic data, VDAC1, RPAB1, GDIA, RL10A, TBA1A) were not significantly enriched (unpaired two-sided t-test; **Figure 4H, Supplementary Materials**).

P-bodies are dynamic, phase-separated condensates, and their fluidity directly influences cellular processes ^124,125^. We hypothesized that cytoplasmic TDP-43 disrupts P-body function as indicated by P-body size (average radius). P-body size was significantly reduced in cultured neurons upon induction of TDP-43^ΔNLS^ (LSM14A: β = −0.0247, p < 0.0001, (95% CI [−0.03, −0.02]); DCP1A: β = −0.0566, p < 0.0001, (95% CI [−0.059, −0.052]); fixed-effect fallback used, **Figure 4I-J**). Effects were estimated with a linear mixed-effects model (DOX treatment as a fixed effect, experimental repeat as a random intercept). Notably, wild-type TDP-43 overexpression (containing functional NLS) in day 8 neurons, caused a slight increase in P-body size with ten-fold smaller effect size compared to the induction of TDP-43^ΔNLS^ (TDP-43^ΔNLS^: β = -0.032, p < 0.0001, (95% CI: [-0.04, -0.03]); TDP-43^WT^: β = 0.003, p = 0.025, (95% CI: [0.00, 0.01])), **Supplementary Figure 20)**, suggesting that TDP-43 mislocalization drives the observed P-body alterations.

Reduced fluidity of biomolecular condensates and a shift toward more solid-like states is a process linked to neurodegenerative diseases including ALS. To test whether cytoplasmic TDP-43 impacts P-body diffusion dynamics, we used partial fluorescence recovery after photobleaching (FRAP). In U2OS cells constitutively expressing YFP-DCP1A and transfected with inducible TDP-43^ΔNLS^, induction of cytoplasmic TDP-43 significantly delayed internal diffusion of YFP-DCP1A within P-bodies, relative to uninduced controls (p < 0.001, Two-way ANOVA with repeated measures and Geisser–Greenhouse correction; **Figure 4K**, **Supplementary Figure 21A, B**). In contrast, induction of the NES control peptide did not significantly affect P-body dynamics **(Supplementary Figure 21C–E)**. These results suggest that cytoplasmic TDP-43 reduces P-body liquidity.

We also evaluated the effect of mislocalized cytoplasmic TDP-43 on P-bodies in orthogonal model systems. In U2OS cells expressing clover-tagged TDP-43^ΔNLS^, cytoplasmic TDP-43 altered P-body size (unpaired t-test, p < 0.0001, **Supplementary Figure 21F**). Notably, this effect was observed only under oxidative stress conditions (0.25 M sodium arsenite, 30 min) and during a subsequent 60-minute recovery period, but not under basal (pre-stress) conditions. This likely reflects a previously reported phenomenon in this cellular model, in which oxidative stress induces the formation of toxic, SG-independent cytoplasmic TDP-43 foci ^126^.

A similar effect was demonstrated in wild type iPSC-derived neurons, where cytoplasmic mislocalization of endogenous TDP-43 was triggered by osmotic stress ^127,128^ (sorbitol 0.4 M, 2 hr; **Supplementary Figure 21G**). Consistently, under sorbitol-induced cytoplasmic TDP-43 conditions, DCP1A-containing P-bodies were significantly smaller (β = −0.0327, p < 0.0001, (95% CI [−0.033, −0.031]), fixed effect fallback used), whereas LSM14A-containing P-bodies were not significantly affected (**Supplementary Figure 21H**). Taken together, these results support interaction between mislocalized cytoplasmic TDP-43 and P-bodies that alter P-body properties, suggesting a role for P-bodies in ALS and FTD that was not shown previously.

Although P-body function is still highly debated, P-bodies seem to play a role in RNA storage and decay ^129^. Thus, we assessed the changes to RNA half-life in response to TDP-43^ΔNLS^ expression in neurons using the incorporation of 4-thiouridine (4sU; 200 μM) into newly transcribed RNA molecules (2 or 3 hours labeling pulse), followed by SLAM-seq and GRAND-SLAM analysis ^130–132^(**Figure 4L, Supplementary Materials**). Induction of TDP-43^ΔNLS^ caused a 10% reduction in global RNA half-life, relative to uninduced control (p-value < 0.0001, Mann–Whitney U test; **Figure 4M**). This reduction in mRNA stability increased to 30% when analyzing mRNAs that are enriched inside P-bodies ^116^ (p-value < 0.0001, Mann–Whitney U test; **Figure 4N**). In contrast, there was a milder 5% decrease in the half-life of RNAs not specifically enriched in P-bodies (p-value < 0.01, Mann–Whitney U test; **Figure 4O**). Therefore, accelerated mRNA decay is caused by cytoplasmic TDP-43 malposition and is specific to mRNAs that reside within P-bodies. Overall, we conclude that cytoplasmic TDP-43 directly interacts with P-bodies, and disrupts their biophysical properties and functions. This suggests a potential role of P-bodies in the molecular mechanisms underlying TDP-43 proteinopathy and neurodegeneration.

### TDP-43 and P-bodies in ALS patient-derived iPSC neurons and post-mortem tissues

We tested NOVA’s ability to identify neuronal phenotypes (ALS vs. healthy human subjects) in patient-derived neurons that were not previously seen by the model. We differentiated iPSCs from a cohort of 21 subjects into day 60 motor neurons ^105,133^, including healthy controls (n=6, 3 males, 3 females), *C9orf72*-ALS patients (C9ALS, n=3, 2 males, 1 female), two sporadic ALS patients without cytoplasmic TDP-43 (sALS-, n=2, 1 male, 1 female), and ten sporadic ALS patients with cytoplasmic TDP-43 (sALS+, n=10, 6 males, 4 females). This cohort (∼9,160 image tiles post-QC) was immunofluorescently stained for 4 organellar markers: TDP-43 granules, P-bodies (DCP1A), nucleus (Hoechst 33342) and microtubule (MAP2; **Figure 5A**). Lines derived from different human subjects are defined as experimental repeats, in accordance with the MDAR (Materials Design Analysis Reporting) framework guidelines ^134^.

**Figure 5.**
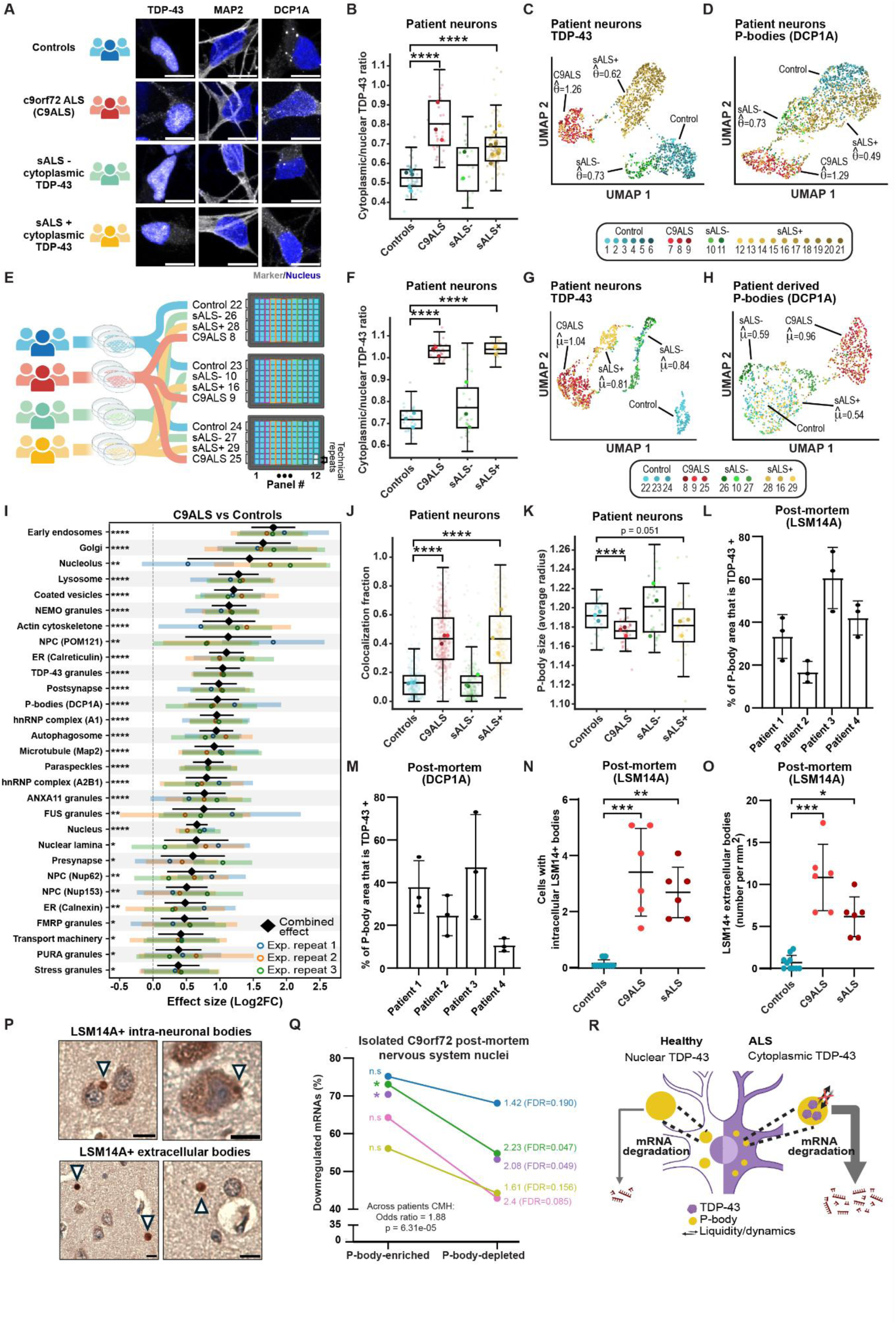
Cytoplasmic TDP-43 and P-body function in ALS patient-derived iPSC neurons and post-mortem tissues. **(A)** Representative images of TDP-43 granules (green), Cytoskeleton (red), P-bodies (magenta), and nucleus (blue) in day 60 iPSC-derived motor neurons from human subjects. Healthy control (blue, n=6), *C9orf72*-ALS (C9ALS, red, n=3), sporadic ALS (sALS) without cytoplasmic TDP-43 (sALS-, green, n=2), or sALS with cytoplasmic TDP-43 (sALS+, yellow, n=10). Lines derived from different human subjects are defined as experimental repeats, in accordance with the MDAR (Materials Design Analysis Reporting) framework guidelines^134^. Scale bar - 10 µm. **(B)** Boxplots of cytoplasmic-to-nuclear TDP-43 signal ratios in C9ALS, sALS- or sALS+ vs. healthy controls (means: 0.802, 0.591 or 0.686 vs. 0.526, respectively). **** p < 0.0001. Linear fixed-effects model with human subject-clustered standard errors (fallback used). Joint UMAP projection of day 60 motor neurons from a cohort of 21 human subjects, showing **(C)** TDP-43 granules and **(D)** P-bodies (DCP1A). Points - single image tiles. ∼60-160 image tiles per human subject/organelle. (*θ̂*) the effect size estimate. **(E)** Experimental design for an expanded marker study of 32 organelles in day 60 motor neurons. Three plates, each containing one human subject from every group. Three experimental repeats (human subjects) for each group in the meta-analysis. **(F)** Cytoplasmic-to-nuclear TDP-43 signal ratios in C9ALS, sALS- or sALS+ vs. healthy controls (means: 1.032, 0.772 or 1.038 vs. 0.717, respectively, **** p < 0.0001, linear mixed-effects model). Joint UMAP projections of **(G)** TDP-43 granules and **(H)** P-bodies (DCP1A). Individual human subject IDs denoted. Subjects 8, 9, 10 and 16 shared between initial and expanded marker cohorts. 30-90 image tiles per human subject/organelle. (*μ̂*) the combined effect size estimate. **(I)** Stacked forest plot of organelle effect size (log2 fold change) in day 60 human subject motor neurons from three C9ALS patients relative to three healthy controls. Colored dots represent the effect estimate per experimental repeat (human subject), with horizontal bars indicating the 95% CI of within-study variance. Paired C9ALS and healthy control within plate and meta-analysis of three plates. The combined effect estimate (*μ̂*) is shown in black diamond with its 95% meta-analytic CI, which accounts for between experimental repeat (human subject) variance. Adjusted p-value: * < 0.05, ** < 0.01, **** < 0.0001. ∼50-170 image tiles per organelle/human subject. **(J)** Boxplots of P-body-positive (DCP1A) pixels that are also positive for endogenous TDP-43. Colocalization fraction in human subjects day 60 motor neuron images in C9ALS, sALS- or sALS+ vs. healthy controls (mean fraction overlap = 0.433, 0.13 or 0.433 vs. 0.128, respectively). ∼70-170 image tiles per human subject; Linear fixed-effects model with human subject-clustered standard errors (fallback used). **** p < 0.0001. Three human subjects per group. **(K)** CellProfiler quantification of P-body size (mean radius) in C9ALS, sALS- or sALS+ vs. healthy controls (means: 1.17, 1.20 or 1.18 vs. 1.19, respectively). Three human subjects per group. Linear fixed-effects model with human subject-clustered standard errors (fallback used). **** p < 0.0001. In all box plots, the black line indicates group mean. Large, saturated dots - human subject means; small, washed dots - individual site images. Percentage of DAB chromogenic immunohistochemistry signal for P-body proteins, **(L)** LSM14A or **(M)** DCP1A, colocalized with TDP-43 aptamer^139^. Each point represents the average colocalized area (%) across three regions of interest (n=3 ROIs) in post-mortem frontal lobe punches from four C9ALS patients (1–4). Mean ± SD, quantified using ImageJ. Quantification of **(N)** intra-cellular (means: 0.07, 3.33, 2.61) and **(O)** extracellular (means: 0.70, 10.83, 6.17) LSM14A-positive(+) bodies in brain sections and **(P)** their representative images (arrowheads), from controls (blue, n=10), C9ALS (red; n=6) or sporadic ALS (dark brown, n=6). Data points - mean number of cells containing intracellular bodies or number of extracellular bodies per mm^2^; Mean ± SD; Kruskal-Wallis test with Dunn’s multiple comparisons; Adjusted p-value: * < 0.05, ** < 0.01, *** < 0.001. **(Q)** Percentage of down-regulated mRNAs in isolated neuronal nuclei without TDP-43 relative to uninvolved normal neuronal nuclei analyzed in five *C9orf72* post-mortem human nervous systems ^140^. Bins of P-body-enriched/depleted mRNAs (by FDR ≤ 0.05 and enrichment ≥ 1 or ≤ −1, respectively in Ref. ^116^). Odds-ratio and one-sided Fisher’s exact test and Benjamini–Hochberg FDR; * p < 0.05. n.s. non-significant. Lower left: Cochran-Mantel-Haenszel (CMH) test for overall association based on per-subject contingency 2×2 tables and controlling for subject as a stratum. χ² statistic, **** p (one-sided) < 0.0001, and 95% Cl [1.38, 2.56]. **(R)** Model for mechanism and consequences of cytoplasmic TDP-43 interactions with P-bodies in ALS.

C9ALS and sALS+ patient groups showed a significant increase in cytoplasmic-to-nuclear TDP-43 signal ratios, relative to healthy controls (C9ALS: β = 0.276, p < 0.0001 (95% CI [0.1749, 0.3776]); sALS+: β = 0.1598, p < 0.0001, (95% CI: [0.1103, 0.2092]); fixed-effect with human subject-clustered standard errors, **Figure 5B**).

UMAP projection of NOVA-derived embeddings displayed group-specific topography of TDP-43 granules and P-bodies (**Figure 5C, D,** C9ALS: *θ̂*=1.26; sALS-: *θ̂*=0.73; sALS+: *θ̂*=0.62; C9ALS: *θ̂*=1.29; sALS-: *θ̂*=0.73; sALS+: *θ̂*=0.49, respectively, adj. p<0.05; permutation test, 100 iterations). These clusters are also reflected in the pairwise Euclidean distances computed on the NOVA-derived embeddings between pairs of human subjects (**Supplementary Figure 22A, B**). Surprisingly, subclusters could be observed in UMAPs of nucleus and microtubule topography with significant effect sizes as well (C9ALS: *θ̂*=1.39; sALS-: *θ̂*=0.77; sALS+: *θ̂*=0.41; C9ALS: *θ̂*=1.24; sALS-: *θ̂*=0.71; sALS+: *θ̂*=0.44, respectively, adj. p<0.05; permutation test, 100 iterations; **Supplementary Figure 22C-F**). Furthermore, colocalization of P-bodies with TDP-43 was significantly greater than expected in C9ALS patients and in patients with cytoplasmic TDP-43 (C9ALS: *β* = 0.063, *p* < 0.001, (95% CI [0.026, 0.1]); sALS+: *β* = 0.188, *p* < 0.0001, (95% CI [0.113, 0.263]); linear fixed-effects model with patient-clustered standard errors, fallback used; **Supplementary Figure 22G**). Therefore, NOVA detects organellar changes in patient-derived neurons, despite being trained solely on isogenic iNDI lines.

Next, we generated another dataset of day 60 neurons (n=12, three subjects per human subject group, **Figure 5E**), stained for 32 organelles (∼58K tile images post-QC). This dataset included ten markers not seen in training: NUP62, NUP98 and NUP153 (nuclear pore complex), Lamin B1 (nuclear envelope), TIA1 (stress granules), EEA1 (early endosomes), hnRNPA1 and hnRNPA2B1 (hnRNP complex), calnexin (ER) and MAP2 (microtubule). Similarly to the first cohort, C9ALS and sALS+ patient groups showed a significant increase in cytoplasmic-to-nuclear TDP-43 signal ratios, relative to healthy controls (C9ALS: β = 0.315, p < 0.0001 (95% CI [0.24012, 0.38942]); sALS+: β = 0.321, p < 0.0001, (95% CI: [0.24591, 0.39521]); linear mixed-effects model, **Figure 5F**). Consistent with the previous cohort, TDP-43 granules and P-bodies UMAPs displayed group-specific topographies (mixed-effect meta analysis: C9ALS: *μ̂*=1.04, adj. p=1.98 × 10^-14^; sALS-: *μ̂*=0.84, adj. p=4.26 × 10^-9^; sALS+: *μ̂*=0.81, adj. p=6.6 × 10^-8^; C9ALS: *μ̂*=0.96, adj. p=1.98 × 10^-8^; sALS-: *μ̂*=0.59, adj. p=8.22 × 10^-4^; sALS+: *μ̂*=0.54, adj. p=7.22 × 10^-2^, respectively; mixed effects. **Figure 5G, H**), with C9ALS forming a distinct community with a mean intra-community distance of *d*=0.02 (TDP-43 granules) and *d*=0.007 (P-bodies) based on pairwise Euclidean distance between human subjects (**Supplementary Figure 23**).

Meta-analytic effect sizes of all 32 organelles were quantified, relative to control, for C9ALS (**Figure 5I** and **Supplementary Figure 24A**); sALS+, sALS- (**Supplementary Figure 24B, C**). In C9ALS patients, 29 organellar markers displayed a change in topography that is significantly different from the controls (**Figure 5I**). Unexpectedly, the effect size of nine organelles was larger than that of TDP-43 granules (early endosomes (*μ̂*=1.8, adj. p=5.33 × 10^-15^), Golgi (*μ̂*=1.64, adj. p=2.93 × 10^-14^), nucleolus (*μ̂*=1.44, adj. p=3.78 × 10^-3^; heterogeneity: I²=82.49%, τ²=0.5422 (95% CI=[0.0369,11.71])), lysosome (*μ̂*=1.28, adj. p=5.33 × 10^-15^), coated vesicles (*μ̂*=1.2, adj. p=6.4 × 10^-15^), NEMO granules (*μ̂*=1.13, adj. p=5.33 × 10^-15^), actin cytoskeleton (*μ̂*=1.13, adj. p=4.76 × 10^-8^), NPC (*μ̂*=1.12, adj. p=1.32 × 10^-3^), and ER (*μ̂*=1.09, adj. p=5.33 × 10^-15^), **Supplementary Figure 24D**). These findings may imply a change in autophagy and endosomal–lysosomal pathways in C9ALS. ^135,136^

To investigate whether patient-derived neurons reflect the suggested crosstalk between TDP-43 and P-bodies, we modeled the fraction of TDP-43 overlap with P-bodies using a linear fixed-effects model with human subject-clustered standard errors (fallback used). Relative to healthy controls, colocalization was significantly increased in C9ALS patients (*β* = 0.305, p < 0.0001, (95% CI [0.27, 0.34]; **Figure 5J**) and sALS+ patients (*β* = 0.304, p < 0.0001, (95% CI [0.18, 0.42]); fixed effect fallback used). In contrast, sALS- patients did not differ significantly from controls (*β* = 0.001, p = 0.94) as expected given the absence of TDP-43 mislocalisation. In addition, P-bodies were significantly smaller in C9ALS patients and showed a near significant reduction in sALS+ patients relative to controls (C9ALS: *β* = - 0.0157, p < 0.0001, (95% CI [-0.0223, -0.009]); sALS+: *β* = -0.0102, p = 0.051, (95% CI [-0.020, 0.00006]); fixed-effects with patient-clustered standard errors, **Figure 5K**)

Finally, we assessed the relevance of TDP-43-P-body crosstalk in ALS pathology, characterized by cytoplasmic TDP-43 aggregates ^137,138^. Approximately 10-60% of the puncta area that was positive for either P-body protein DCP1A or LSM14A colocalized with TDP-43-specific RNA aptamer^139^ and colocalization was evident in 46-100% of the neurons in C9ALS post-mortem motor cortices (four human subjects, **Figure 5L,M** and **Supplementary Figure 25A, B**). In C9ALS post-mortem tissue, cells contained a significantly lower number of puncta containing P-body proteins (DCP1A, LSM14A, or AGO2) compared to non-neurological controls (Mann-Whitney U test, p < 0.05). In contrast, puncta area size was significantly increased for LSM14A and DCP1A, but not for AGO2 (Mann-Whitney U test, p < 0.05, **Supplementary Figure 25C-I**). Furthermore, analysis of neurons with and without TDP-43 mislocalisation to the cytoplasm (-/+ cytoplasmic TDP-43) in C9ALS post-mortem tissues, revealed a significantly lower number of puncta containing LSM14A or DCP1A in the presence of mislocalised TDP-43 (nested t-test, p < 0.01, **Supplementary Figure 25J-K**), and significantly increased P-body area size (nested t-test, p < 0.05, **Supplementary Figure 25L-M**). This indicates that these properties are directly affected by cytoplasmic TDP-43. Intriguingly, large intra-neuronal and extracellular LSM14A-positive bodies were evident in C9ALS and sporadic ALS (sALS) brains, which were nearly absent in control tissues (10 controls, 6 C9ALS, 6 sAS; Kruska-Wallis test with Dunn’s multiple comparisons test, adj. P-value for intra-neuronal bodies = 0.0006 or 0.0034; and for extracellular bodies = 0.0001 or 0.0182, C9ALS or sALS vs. controls, respectively; **Figure 5N-P**). Thus, TDP-43 pathology is associated with altered distribution of typical P-body markers.

Since cytoplasmic TDP-43 affects mRNA decay in cultured neurons, we sought supporting evidence in neuropathology. We analyzed RNA-seq data from post-mortem neuronal *C9orf72* nuclei that were sorted based on their TDP-43 content ^140^. Nuclei exhibiting a loss of TDP-43 had more severe downregulation of P-body-enriched mRNAs relative to P-body-depleted mRNAs. In two out of five patients, this increase in the proportion of downregulated P-body mRNAs was statistically significant, as well as the overall effect across patients (Cochran-Mantel-Haenszel test < 0.0001, **Figure 5Q**). These data are consistent with altered RNA metabolism in TDP-43–deficient neurons. Therefore, accelerated mRNA decay is associated with cytoplasmic TDP-43 malposition and is specific to mRNAs that reside in P-bodies.

Together, cytoplasmic TDP-43 crosstalks with P-bodies, disrupting their biophysical properties and function, proposing a role for P-bodies in the molecular mechanisms underlying TDP-43 proteinopathy and neurodegeneration (**Figure 5R**).

## Discussion

In this work, we present NOVA, a deep learning framework for microscopic phenotyping studies in neurons. We developed a ViT architecture with supervised contrastive loss to systematically encode perturbation-induced subcellular localization and morphology patterns from immunofluorescent images of membrane-bound and membraneless organelles (**Figure 1**). NOVA enables characterization of organellar responses to perturbations on a single metric scale, including responses to stress (**Figure 2**) and ALS-associated genetic variants (**Figure 3**). Our study further revealed that cytoplasmic TDP-43, a key hallmark of ALS, alters P-body biophysics and function (**Figure 4**). Deep microscopic phenotyping with NOVA can also be applied to human–derived motor neurons (**Figure 5**), overcoming inter-patient heterogeneity and sex differences^141^. The organellar changes described in experimental ALS-like models and ALS patient neurons demonstrate both unique and shared alterations across different ALS genetic backgrounds, highlighting the broader potential of organellomics to advance insight into neuro-cellular biology and prioritise functionally relevant effects.

Organellomics is a new high-throughput omics approach for comparison of any number of organelles across diverse subcellular topographies. The relevance of organellomic studies lies in the need to understand the complex regulation of cellular functions through the network of membrane-bound and membraneless organelles^87,142–149^. Our image-based organellomics strategy provides a compelling alternative to traditional biochemical methods of organellar fractionation^3–7^, addressing the inherent challenges of isolating organelles, particularly liquid-like membraneless structures^150^. The use of thoroughly characterized commercial antibodies for immunofluorescence in the method we have developed, combined with standard confocal microscopy, offers a flexible platform that is easily transferable and scalable, making it readily adoptable by most laboratories without specialized infrastructure. Thus, our approach delivers unprecedented depth to organellar studies while enabling investigations at scale.

NOVA’s ability to study perturbed cultured human neurons offers several advantages. First, the model was pre-trained on OpenCell data from 1,311 proteins, learning to encode localization patterns from a publicly available dataset. This pre-training step optimized the model for predicting organelle identity from image embeddings and enhanced its generalizability to unseen organelles. Next, we extended the model to a perturbation learning task through contrastive loss and partial fine-tuning. During fine-tuning on a dataset composed of ∼3.2 million images of perturbed cultured human neurons generated in-house, we selectively froze specific layers using the angle metric approach, and replaced the loss function with a contrastive objective that encourages the model to learn features that are robust and discriminative for perturbation effects.

Furthermore, we developed an organelle scoring framework that ranks perturbation responses and enables direct comparisons across all studied organelles, while accounting for inter-experimental repeat differences. To ensure robust and reproducible statistical inference, experimental repeats were explicitly modeled using mixed-effects modeling. To test whether organelles are consistently affected, this approach incorporates random effects to capture variability alongside fixed effects representing the biological conditions or perturbations of interest. As suggested in ^151^, this strategy separates true treatment effects from experimental repeat-specific noise, thereby improving reproducibility and interpretability. In our framework, NOVA-derived embeddings are used to estimate organelle-level effect sizes (*θ̂*) that capture the extent to which a perturbation shifts organelle topography from baseline. Effect sizes are calculated per perturbation-organelle-experimental repeat combination, relative to the natural variability observed in control embeddings. Experimental repeat-specific effect sizes (*θ̂*) are then combined through meta-analytic modeling to yield a combined effect size (*μ̂*), which accounts for heterogeneity across experimental repeats (cell line differentiation, or human subject). Organelles are ranked by their combined effect size, and multiple hypothesis correction is applied. By operating exclusively on distances in latent space, this framework provides a unified metric for quantifying organellar changes in response to perturbations. Thus, mixed-effects meta-analysis of NOVA-derived embeddings determines both effect size and reproducibility with rigorous statistical support.

NOVA’s generalizability is demonstrated by its ability to analyze data from multiple cell lines, including those not seen during training (e.g., TDP-43^ΔNLS^ and patient-derived motor neurons), across different neuronal types (day 8 cortical neurons and day 60 human subject–derived motor neurons), and using data generated by different microscopes and experimentalists across laboratories. In addition, we successfully performed inference on several neuronal markers not included in the training set. NOVA also performed well in a clinically relevant out-of-distribution task: identifying neuronal disease phenotypes (human subject–derived C9 and sporadic ALS neurons versus healthy controls), where disease-related changes outweighed individual variability. Together, these results support NOVA as a solid foundation for future AI-driven studies in neurobiology.

Moreover, NOVA offers practical advantages: (i) our in-house dataset, comprising millions of neuronal images captured under diverse perturbations, is now released and publicly available, providing a valuable resource for future research. (ii) the technology operates on standard confocal images, offering a cost-effective alternative to complex multiplexed platforms such as PhenoCycler or 4i^152,153^; (iii) it can be applied retrospectively to existing microscopy datasets and is agnostic to the number of fluorescent channels; and (iv) compared with methods based on hand-engineered features (e.g., CellPainting and CellProfiler^46,47^), NOVA’s deep representation learning captures multi-scale patterns without pre-defined objects, which was shown to generalize better across markers and conditions^154–157^. Noteworthy, Cell Painting captures standardized morphology using six dyes and offers interpretable baselines, whereas NOVA learns organelle organization directly from endogenous protein patterns, flexibly adapts to any fluorescent marker, and quantifies perturbation effects across numerous organelles.

NOVA enabled mapping of the neuronal stress response at the level of the entire organellome for the first time. This revealed that oxidative stress induces not only the expected formation of stress granules (SGs), but also topographic remodeling of diverse additional organelles. While changes in individual organelles have been reported^158,159^, we are able to demonstrate the relative topographic changes to all organelles, including nuclear PML bodies and mitochondria. In this way, NOVA highlights how stress triggers a distributed reorganization across multiple organelles. Recently, Rhoads et al., suggested organelle-level differences between neurons and astrocytes in rodent CNS cells in response to stress ^160^, suggesting another layer of cell-type specificity.

Application of NOVA to the iPSC Neurodegenerative Disease Initiative (iNDI) provided the first organellomic maps of ALS-affected human cortical neurons. This revealed unanticipated changes in FUS granules, P-bodies, and lysosomes in several ALS-associated variants. *TARDBP* M337V drove only modest alterations, consistent with literature reporting minimal mislocalization of TDP-43 in premature iPSC-derived neurons^161,162^, possibly reflecting the early developmental stage of the neurons. The *FUS* R495* variant, clinically associated with aggressive juvenile ALS and deficient in its nuclear localization signal^99,163–165^, was analyzed alongside an isogenic revertant in which the arginine at position 495 was restored. NOVA detected robust changes in FUS granules and additional compartments, including the hnRNP complex, peroxisomes, lysosomes, and PURA granules. Comparison with both wild-type and revertant controls indicated that many alterations reflected genuine FUS-driven organellar changes, noticeable even in the context of passage effects or CRISPR off-target edits. Together, these data confirm that FUS mislocalization exerts widespread, reproducible effects on organelle organization.

A prominent NOVA finding was the reproducible sensitivity of P-bodies to cytoplasmic TDP-43, in line with previous work in rat hippocampal neurons^128^. Inducible TDP-43^ΔNLS^ neurons displayed significant remodeling of P-bodies (marked by LSM14A/DCP1A) without concomitant SG changes, demonstrating that TDP-43–P-body crosstalk can be SG-independent. This is intriguing given the reported interactions between P-bodies and SGs and TDP-43 influence on P-body-SG docking, ^145,166^ but resonates with other reports about independence of cytoplasmic TDP-43 and stress granules ^126^. Orthogonal assays reinforced this conclusion: microscopy showed colocalization between P-bodies and TDP-43 inclusions; proximity labeling/MS identified P-body proteins enriched in the vicinity of cytoplasmic TDP-43; and quantitative morphology analyses revealed smaller, less fluid P-bodies, consistent with FRAP-based evidence of altered P-body biophysics. These effects extended to patient-derived neurons and C9ALS post-mortem cortex, where cytoplasmic TDP-43 aggregates co-localized with P-body markers and correlated with altered P-body morphology. Furthermore, large intracellular and extracellular LSM14A-positive inclusions were specific to human C9ALS and sporadic ALS neuropathology and absent in non-neurological controls, suggesting new potential biomarkers.

Cytoplasmic TDP-43 also increased the decay of mRNAs enriched in P-bodies, which are similarly downregulated in nuclei from *C9orf72* TDP-43 proteinopathy patients. Together, and supported by recent preprints that suggest new roles for P-bodies in ALS ^167,168^, these observations highlight P-bodies as a convergent hub in TDP-43 proteinopathies and as a potential therapeutic target. These findings implicate P-bodies in the pathobiology of C9ALS and motivate targeted studies of their dynamics, RNA-handling roles, and interactions with TDP-43.

We acknowledge notable limitations in the model. In order to be applicable to any cell type, NOVA’s design intentionally avoids whole-cell contour segmentation and the training images predominantly contain single cells. This confers substantial computational efficiency but confines use to homogeneous cultures. Analyses in more complex settings, such as co-cultures, mixed genotypes, and tissue sections, will require additional development. At present, the NOVA model has been validated on iPSC-derived cortical and motor neurons. We anticipate that it can be extended to additional disease-relevant cell types, including microglia and astrocytes. Similarly, the present focus on static topography could be broadened by quantifying organelle contact sites and microscopy-compatible functional reporters could provide complementary readouts of organelle activity. Quality control of such large and complex screening datasets, model training on new data, and statistical inference all require a degree of machine learning proficiency. Finally, translating quantified topographic shifts into mechanisms will benefit from targeted perturbations and dedicated interpretability methods, which together can provide clearer causal insight into the underlying biology.

Altogether, organellomics is a microscopy-based framework for interrogating organellar cell biology at scale. It builds on standard immunofluorescence workflows and functions robustly across diverse imaging platforms. It provides representation of the full organellar landscape, enabling quantitative comparisons of cellular responses to any type of perturbation. In doing so, organellomics move beyond compartment-specific narratives toward system-level deep phenotyping of cellular organization and its disruption in disease. The NOVA framework is broadly extensible to additional disease phenotypes, other types of patient-derived cells, and existing imaging data. Organellomics holds the potential to generate novel hypotheses in cellular and disease biology research and may foster diagnosis and precision medicine for neurological diseases.

## Supplementary Figures

**Supplementary Figure 1.**
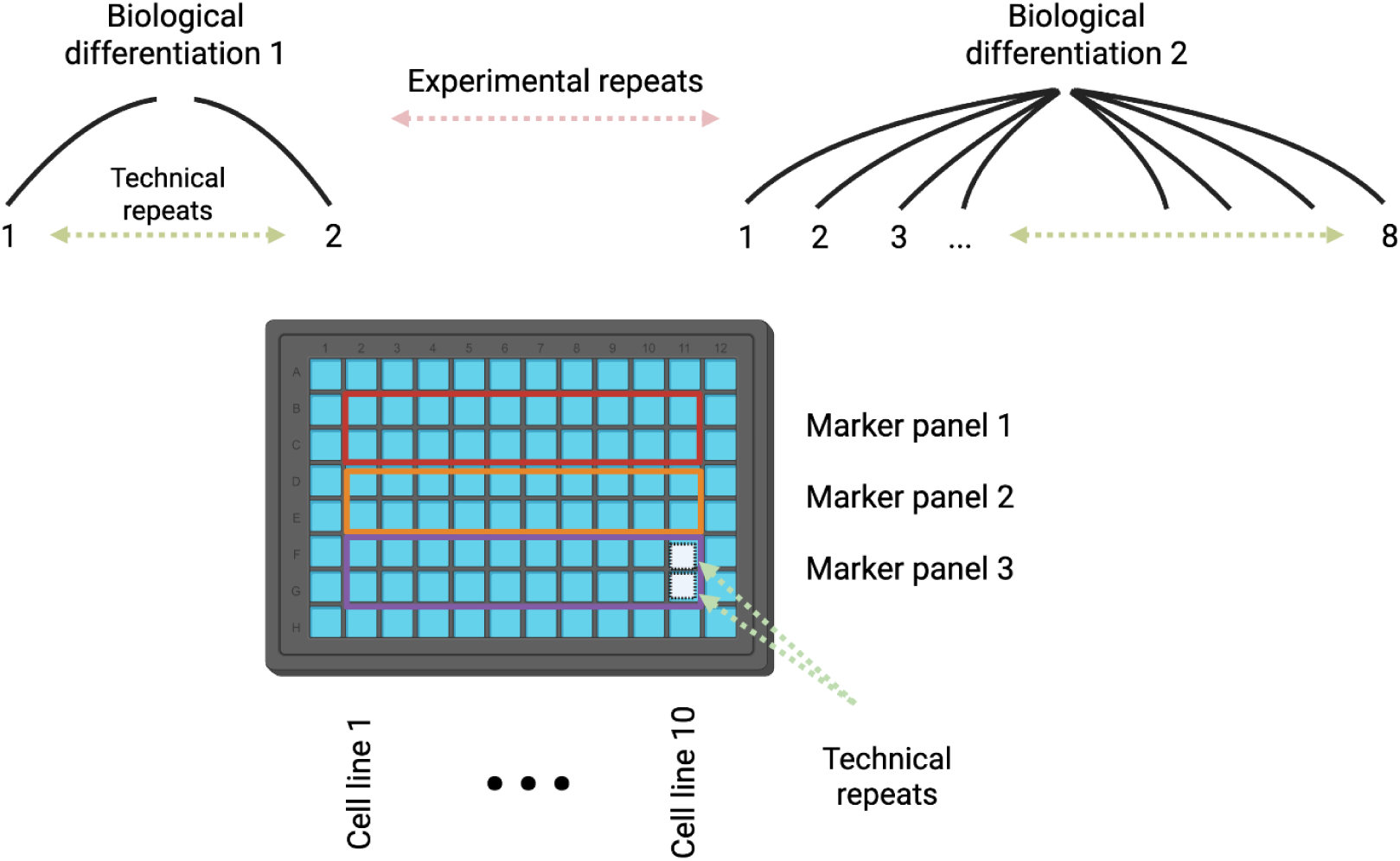
Experimental design of iPSC-derived cortical neurons used for model training. Each 96-well plate included 10 cell lines or conditions and 3 marker panels, with a total of 60 wells used per plate. A full experimental repeat for studying 25 organelles was generated using 11 marker panels across 4 different plates (see Table 1). Each well contained a distinct cell line or experimental condition, with a specific marker panel consisting of antibodies targeting 2–4 organelles. From each well, 100 site images were acquired. For each full organellomic study, two wells (technical repeats) per marker panel per plate were imaged. Bottom: an exemplary plate design from a single experimental repeat (biological differentiation).

**Supplementary Figure 2.**
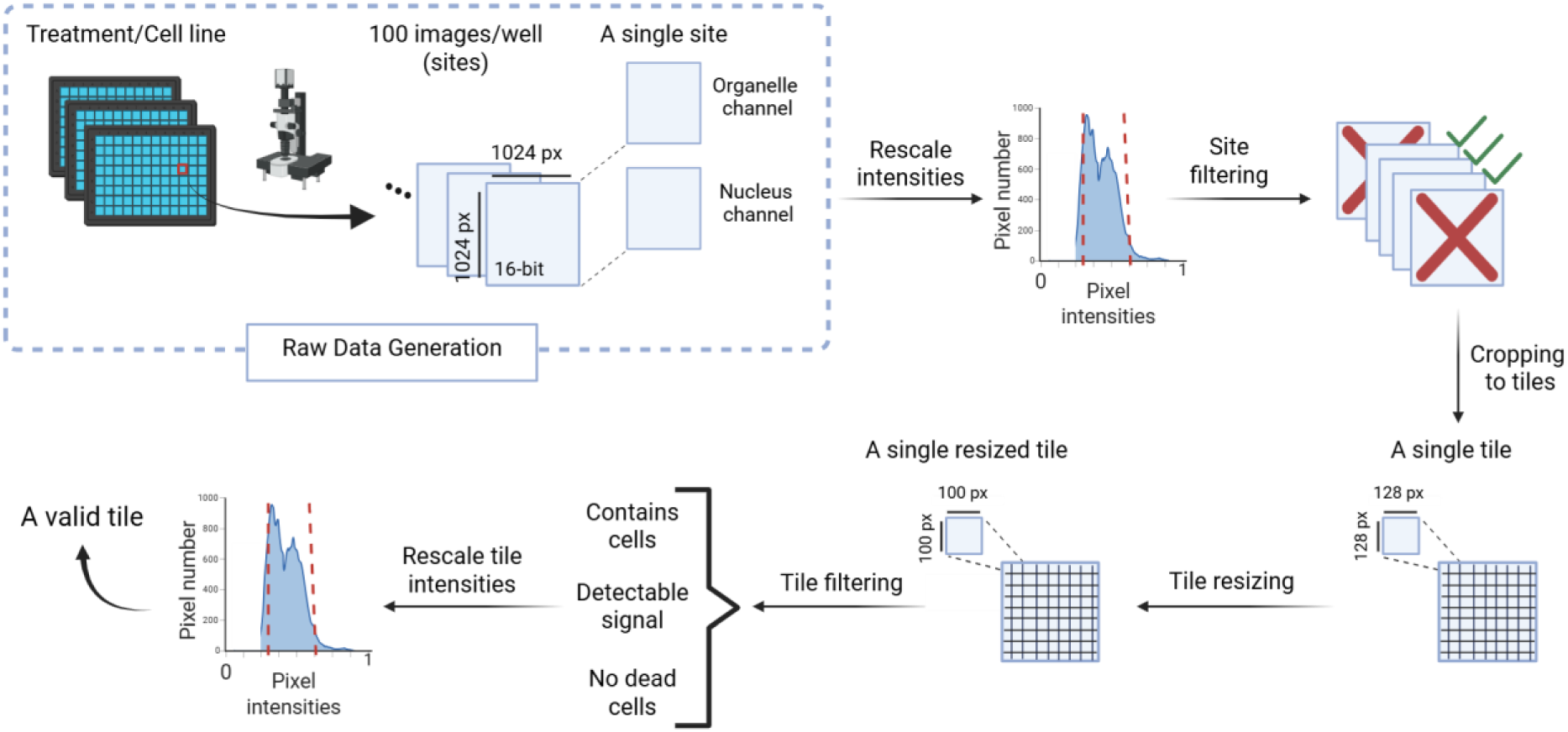
Image quality control (QC) and pre-processing. Microscopy images were acquired at defined fluorescent emission frequencies, stained for a single organelle, and collected at 16-bit resolution, 1024 × 1024 pixels (referred to as “site images”). All organelle images were paired with the corresponding nucleus channel. Site images were rescaled to [0,1] and tested for autofocus quality using the Brenner score (**Methods**). Site images that passed QC were cropped into 64 tiles of 128 × 128 pixels. Tiles then resized to 100 × 100 pixels. Automated nuclei detection was performed on the nucleus channel using CellPose, retaining tiles with at least one but no more than five nuclei. The resulting tiles were rescaled again to [0,1]. To reduce false positives, tiles lacking detectable signal in either channel or flagged as containing dead cells were excluded (**Methods**). The remaining tiles were retained for analysis.

**Supplementary Figure 3.**
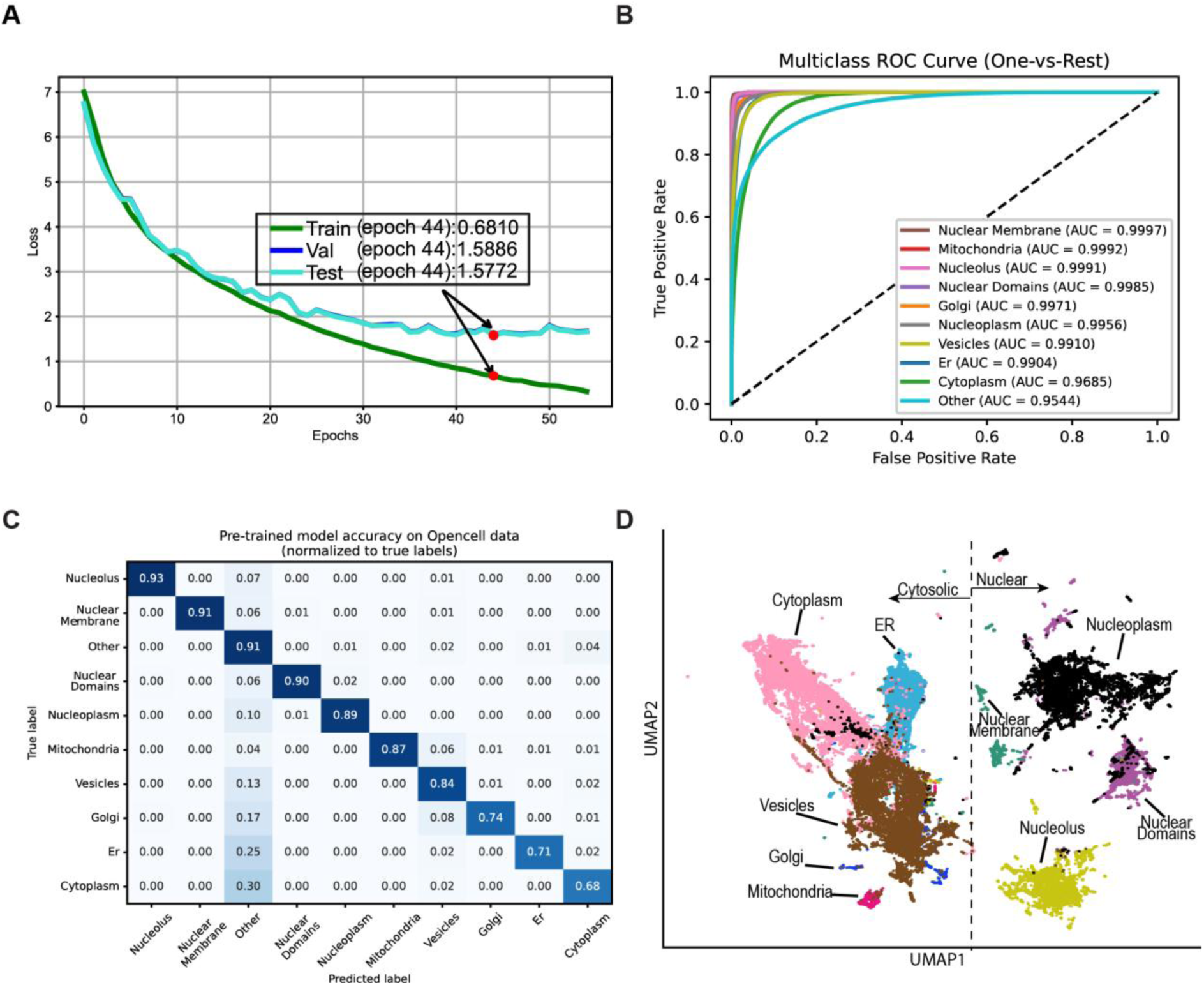
ViT-based protein classification model trained to encode organelle subcellular localization patterns of 1,311 proteins in human embryonic kidney cells. **(A)** Training curves of the model on the OpenCell dataset (train, validation, and test). The y-axis shows the loss value, and the x-axis the number of epochs. Epoch 44, marked by a red dot, was selected as the final model, as it was the last epoch with an improvement in validation loss. Exact loss values are shown in the inset. **(B)** Receiver operating characteristic (ROC) curves for each organelle class in a one-vs-rest setting. The x-axis represents the false positive rate, and the y-axis the true positive rate. Each colored curve corresponds to one organelle class, with its area under the curve (AUC) shown in the legend. AUC values near 1 indicate excellent discriminative performance. The dashed diagonal line represents random classification (AUC = 0.5). The class “Other” represents markers not associated with annotated organelles in the dataset. **(C)** Confusion matrix showing the accuracy of the pre-trained model in predicting organelle identity. The x-axis represents predicted labels, and the y-axis true labels. Values are normalized per true label to [0,1]. High diagonal values indicate correct predictions, while off-diagonal values reflect misclassifications. The model achieved strong performance across most organelles. **(D)** UMAP visualization of the OpenCell test set (held-out data), colored by protein localization categories, shows distinct organelle clustering. Only proteins with clear and exclusive localization patterns are depicted (∼7k image tiles per localization category). The list of annotated proteins is provided in the **Supplementary Materials** (OpenCell_annotated_proteins.csv).

**Supplementary Figure 4.**
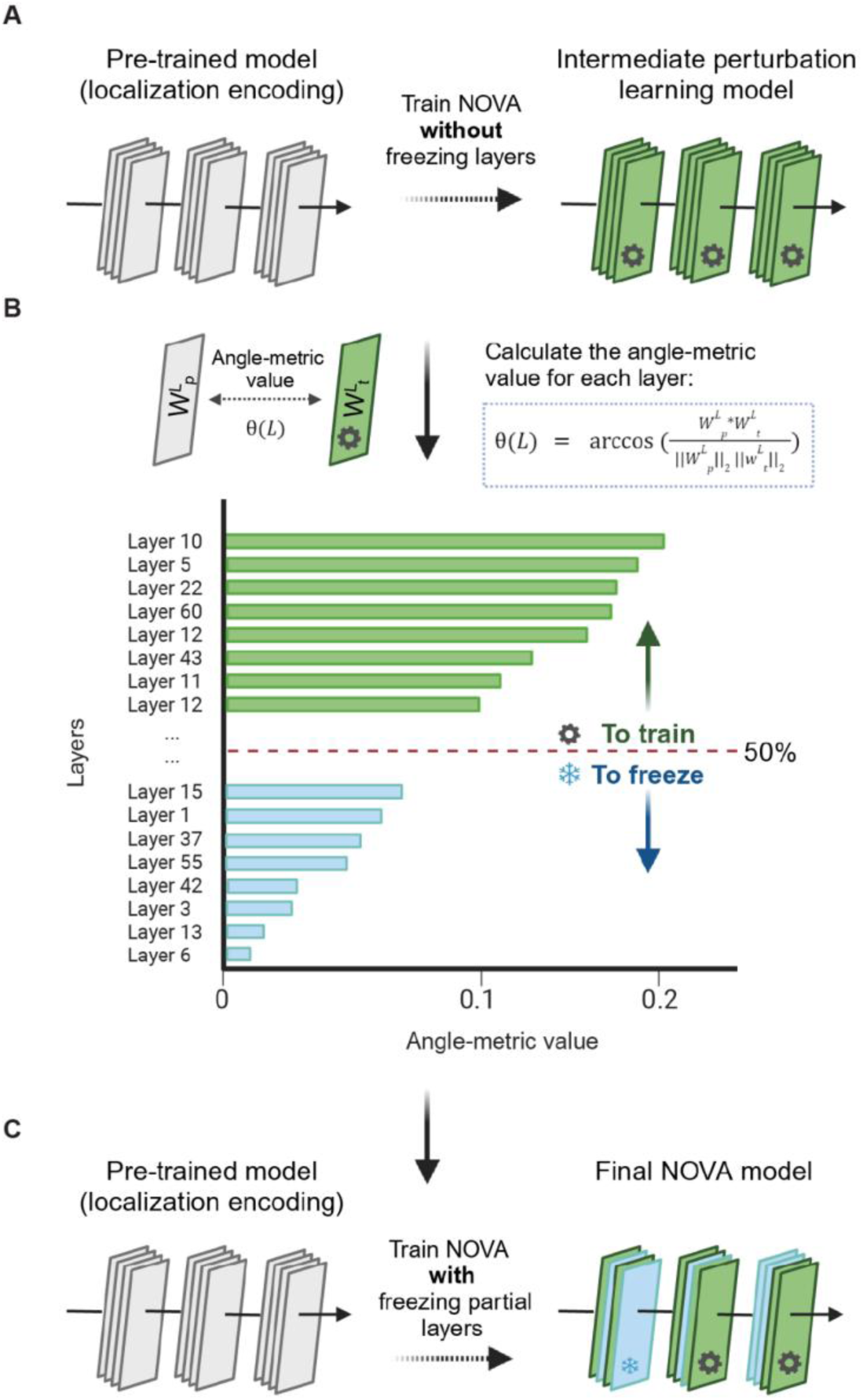
Partial fine-tuning using the angle-metric approach. Starting from the pre-trained model for localization encoding on the OpenCell dataset, a data-driven strategy was applied to identify which layers to freeze during fine-tuning for perturbation learning. Freezing a layer prevents it from updating during fine-tuning. **(A)** Initial fine-tuning with a contrastive loss function was performed on neuronal perturbation data without freezing any layers. **(B)** The angle metric, which is a vector similarity measure, was calculated between corresponding layers of the pre-trained and fine-tuned models. Layers with angle metric values below the median were frozen. **(C)** A final fine-tuning step, beginning from the same pre-trained model with the selected layers frozen, produced the NOVA model used throughout this work.

**Supplementary Figure 5.**
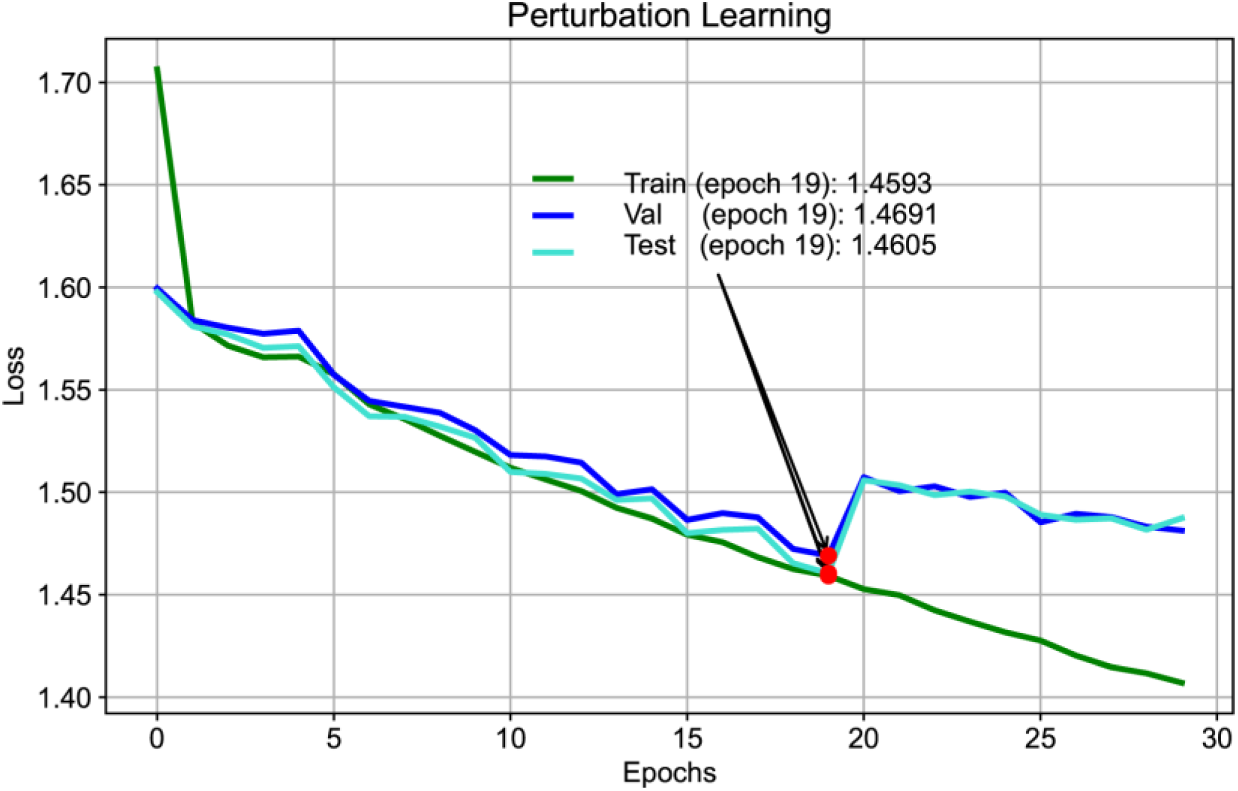
Loss curves for the perturbation learning ViT fine-tuned model. The training curves of the fine-tuned model for perturbations learning on iPSCs-derived neurons (train, val (validation), and test. Loss value of the model (y-axis) and the number of epochs (x-axis). Epoch 19, marked by a red dot, was selected as the final model, as it was the last epoch with an improvement in validation loss. Exact loss values depicted.

**Supplementary Figure 6.**
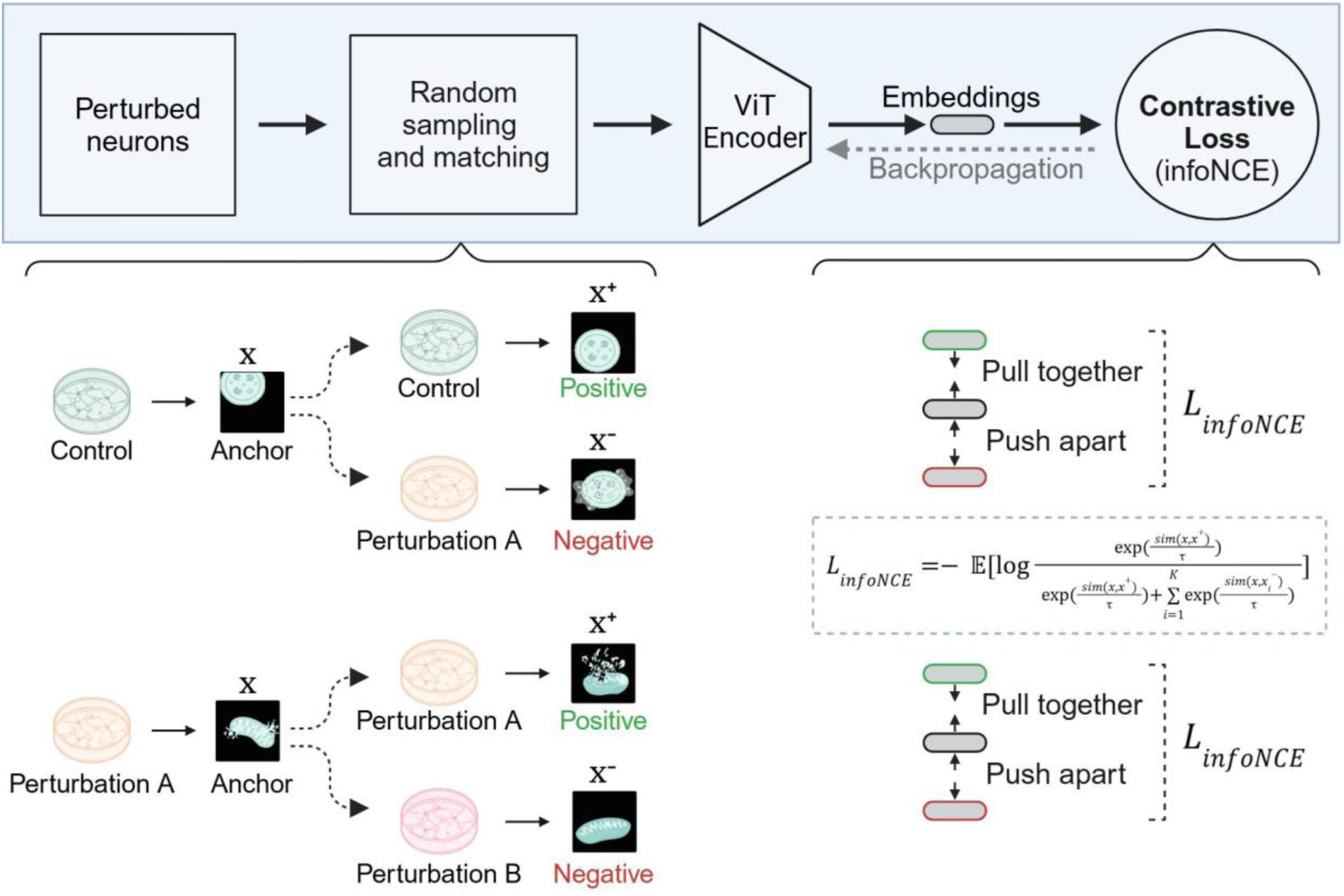
Perturbation learning workflow with contrastive learning. For each image in the training set (the “anchor”), six additional tiles were randomly sampled from different wells and experimental repeats: one positive and five negatives. Positives were tiles from the same organelle and perturbation as the anchor, while negatives were tiles from the same organelle under different perturbations. Embeddings for all seven samples were computed, and the InfoNCE contrastive loss was applied. Contrastive learning pulled together images with similar topographies (anchor and positives) and pushed apart dissimilar ones (negatives), enabling the model to learn a structured latent space. Random sampling across experimental repeats was performed to reduce overfitting to non-biological differences such as biological differentiation or plate effects. See **Methods** for details.

**Supplementary Figure 7.**
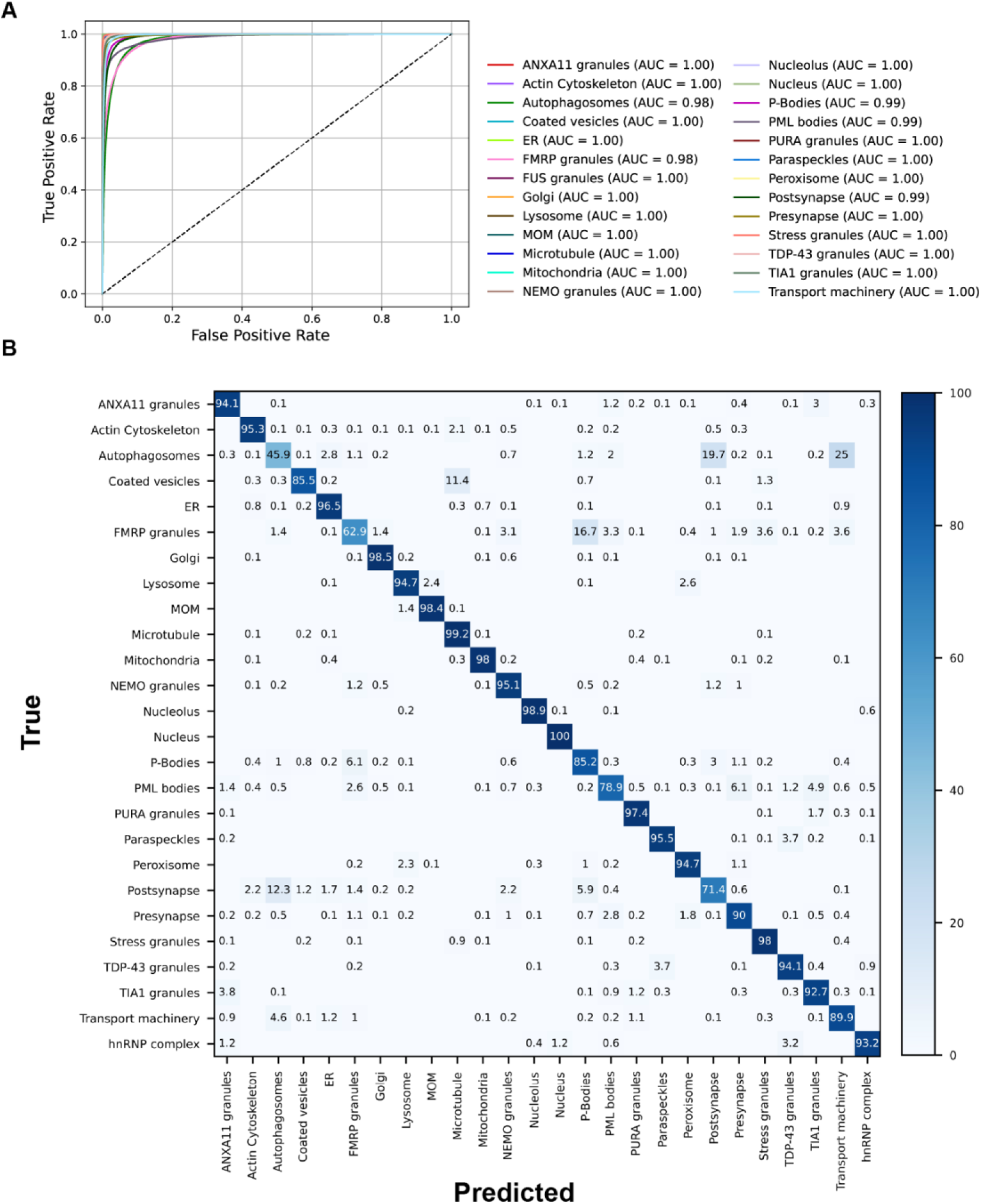
Organelle identification performance of NOVA embeddings. NOVA’s embeddings showed strong organelle identification performance on the dataset in **Figure 1C**. A classifier was trained on these embeddings (see **Methods**) and evaluated using two complementary approaches. **(A)** Receiver operating characteristic (ROC) curves for all organelles. Each curve shows the change in true positive rate relative to false positive rate as the classification threshold varied. The area under the curve (AUC) is reported, with values > 0.97 for all organelles, indicating excellent discriminative performance. **(B)** Confusion matrix of classification accuracy across organelles. Values represent the percentage of true labels assigned to each predicted label. High diagonal values indicate strong prediction accuracy, while low off-diagonal values reflect minimal misclassification.

**Supplementary Figure 8.**
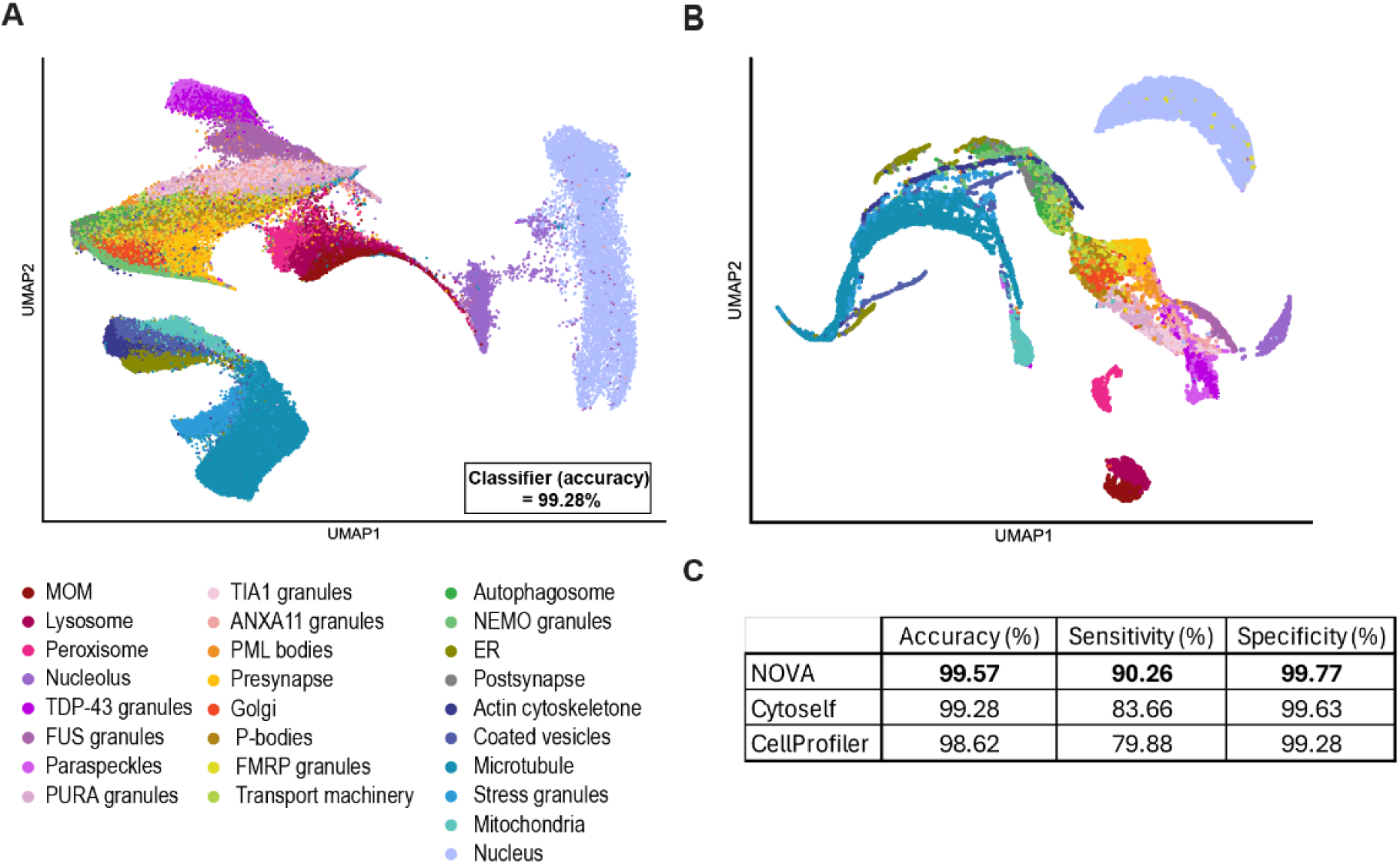
Benchmarking NOVA in neuronal organelle identification against published methods. The organelle landscape in the dataset shown in **Figure 1C** displayed better-defined clusters compared to **(A)** Cytoself-derived features (classifier accuracy = 99.28%; ∼6k image tiles per organelle) or **(B)** CellProfiler-derived features (classifier accuracy = 98.62%; ∼560 image sites per organelle to match CellProfiler’s typical site-level analysis). **(C)** Performance comparison of classifiers trained on NOVA-derived embeddings, Cytoself-derived embeddings, and CellProfiler features for organelle identification. While Cytoself and CellProfiler achieved equivalent overall accuracy, NOVA’s embeddings captured the information needed for this task with higher sensitivity (83.66% vs. 79.88% vs. 90.26%).

**Supplementary Figure 9.**
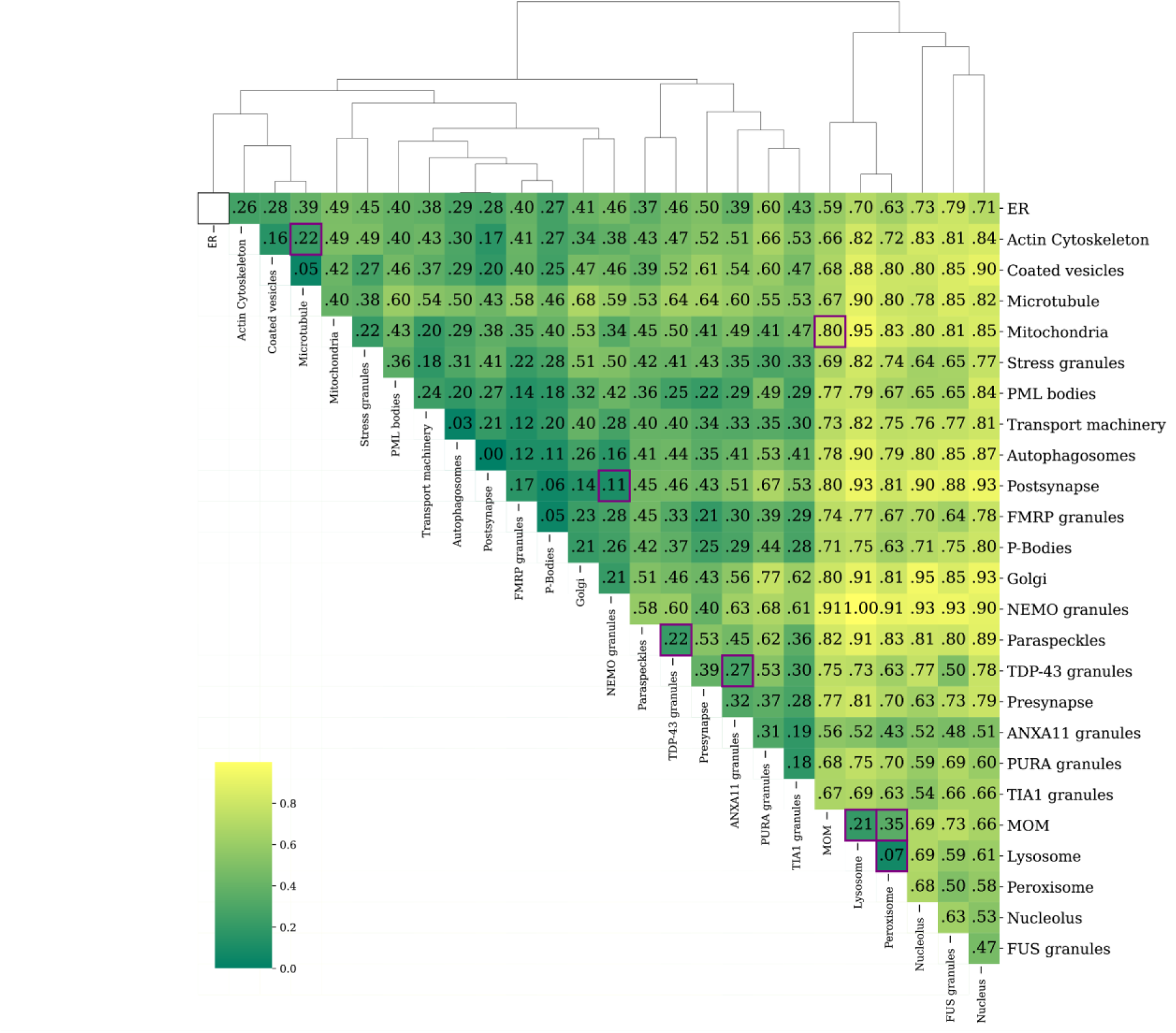
Distances across the organellar landscape of day 8 human iPSC-derived neurons. For each organelle pair in the landscape shown in **Figure 1C**, pairwise Euclidean distances were computed between NOVA-derived embeddings. Heatmap values represent the median distance, rescaled to the [0,1] interval using min–max normalization. A value of 0 indicates the closest organelles (highest similarity, darker colors), and 1 the most distant (lowest similarity, lighter colors). Rows and columns were clustered by agglomerative hierarchical clustering of the scaled distance values (Euclidean distance, average linkage). Dendrogram depicts the hierarchical relationships among organelles.

**Supplementary Figure 10.**
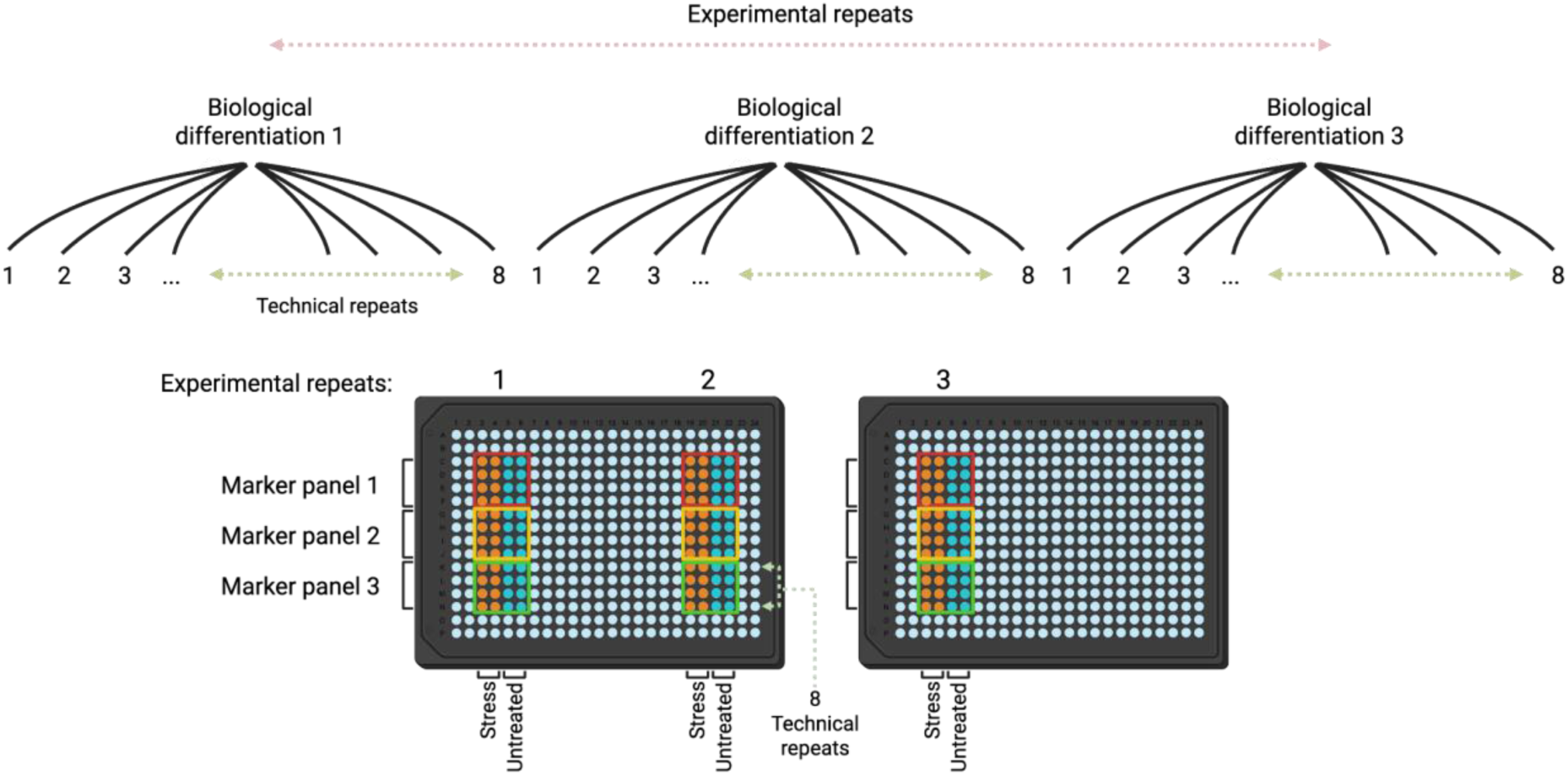
Experimental design of the NIH CARD dataset, related to. **Figures 1 and 2**. Three experimental repeats (biological differentiations), with eight technical repeats (wells) each. Twenty-five site images were acquired per well. Two columns for oxidative stress treatment and two untreated. Every four rows were assigned to a different marker panel. 12 marker panels in total with three marker panels per plate. Cells were seeded into PLO-coated 384-well plates on day 4 post-differentiation and fixed on day 8 of differentiation.

**Supplementary Figure 11.**
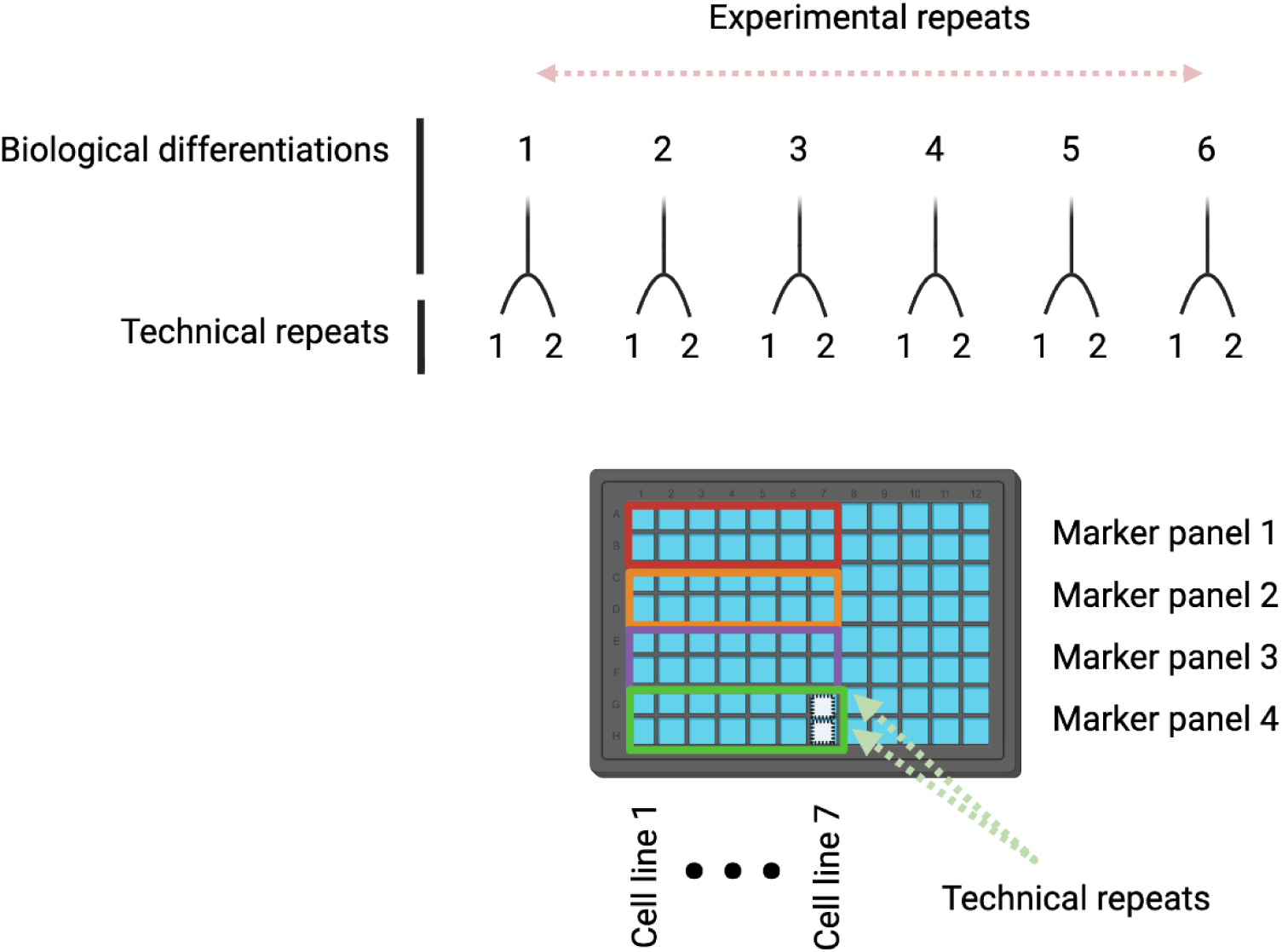
Experimental design of iNDI lines used for inference related to. **Figure 3**. Six experimental repeats (biological differentiations), with two technical repeats (wells) each. Seven iNDI lines: ancestral isogenic control line, Knockout Laboratory F2.1 Jackson (wild-type, KOLF), *FUS* R495* homozygous, *FUS* R495* heterozygous, *FUS* revertant (corrected *495R), *TARDBP* M337V homozygous, *OPTN* Q398E homozygous, *TBK1* E696K homozygous. Each line seeded into a separate column. Every two rows were assigned to a different marker panel. 12 marker panels in total with four marker panels per plate. iNDI cells were seeded into PLO-coated 96-well plates on day 4 post-differentiation and fixed on day 8 of differentiation. Two hundred fifty site images were acquired per well. An exemplary plate design from a single experimental repeat (biological differentiation) is depicted.

**Supplementary Figure 12.**
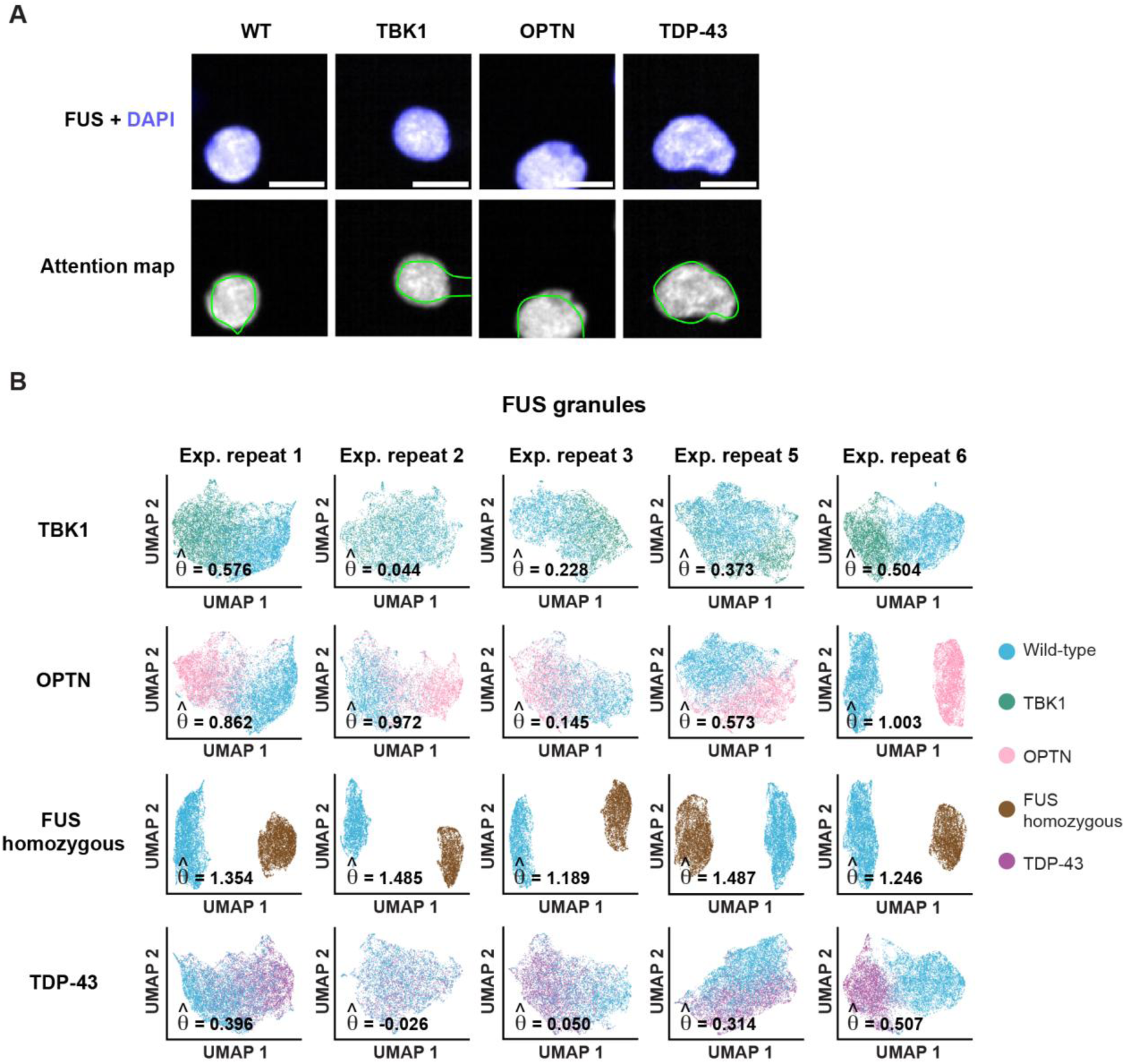
Altered topography of FUS granules in the iNDI lines. **(A)** Representative images of FUS granules across iNDI lines. Nucleus (hoechst 33342, blue) and overlaid model’s attention maps. Scale bar - 10 µm. **(B)** UMAP visualization of FUS granule changes per experimental repeat across iNDI lines. experimental repeat # 4 is missing due to insufficient valid image tiles on FUS marker (∼2.5K–7K tiles per line/experimental repeat). (*θ̂*) the effect size estimate.

**Supplementary Figure 13.**
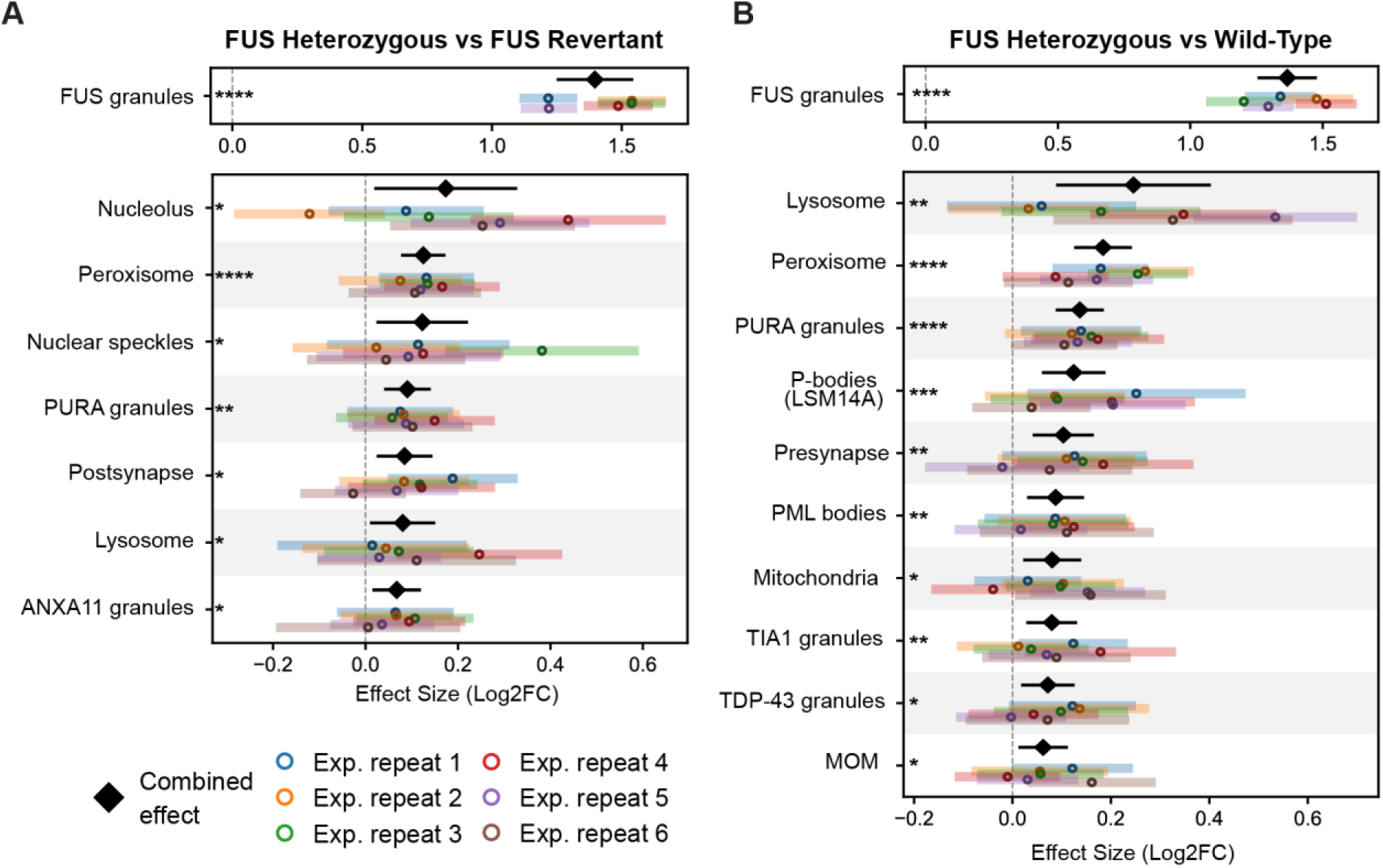
Organellar scoring of iNDI FUS lines reveals unique and shared changes, including FUS granules. Stacked forest plot of organelle effect sizes (log₂ fold change) in day 8 neurons carrying the heterozygous FUS R495* variant relative to **(A)** the genetically corrected FUS revertant line and **(B)** wild-type control. Six experimental repeats; 29 organelles analyzed. Only organelles significantly altered in mixed-effects meta-analysis are shown (**Methods**). Colored dots represent experimental repeat effect estimates, with horizontal bars indicating 95% CI. The combined effect estimate (***μ̂***) is shown as a black diamond with its 95% meta-analytic CI, accounting for between-experimental repeat variance. Adjusted p-values: * < 0.05, ** < 0.01, *** < 0.001, **** < 0.0001. ∼780-6500 image tiles per organelle/variant/experimental repeat. FUS granules were the most altered organelle (***μ̂*** = 1.366, adj. p = 6.4 × 10⁻¹⁵; I² = 74.99%, τ² = 0.0123, 95% CI = [0.0011, 0.0944]), followed by lysosomes (***μ̂*** = 0.246, adj. p = 4.9 × 10⁻³; I² = 78.95%, τ² = 0.0289, 95% CI = [0.0059, 0.1756]). P-bodies marked by LSM14A, were significantly affected in the heterozygous FUS R495* variant (***μ̂*** = 0.124, adj. p = 5.6 × 10⁻⁴) but not in the FUS R495* homozygous variant. Peroxisomes and PURA granules were significantly altered relative to both wild-type (***μ̂*** = 0.184, adj. p = 5.3 × 10⁻⁹; ***μ̂*** = 0.136, adj. p = 1.95 × 10⁻⁷, respectively) and the genetically corrected FUS revertant line (*495R, ***μ̂*** = 0.125, adj. p = 2.2 × 10⁻⁶; ***μ̂*** = 0.09, adj. p = 2 × 10⁻³), alongside FUS granules (***μ̂*** = 1.397, adj. p = 6.4 × 10⁻¹⁵; I² = 86.84%, τ² = 0.0243, 95% CI = [0.0055, 0.1664]).

**Supplementary Figure 14.**
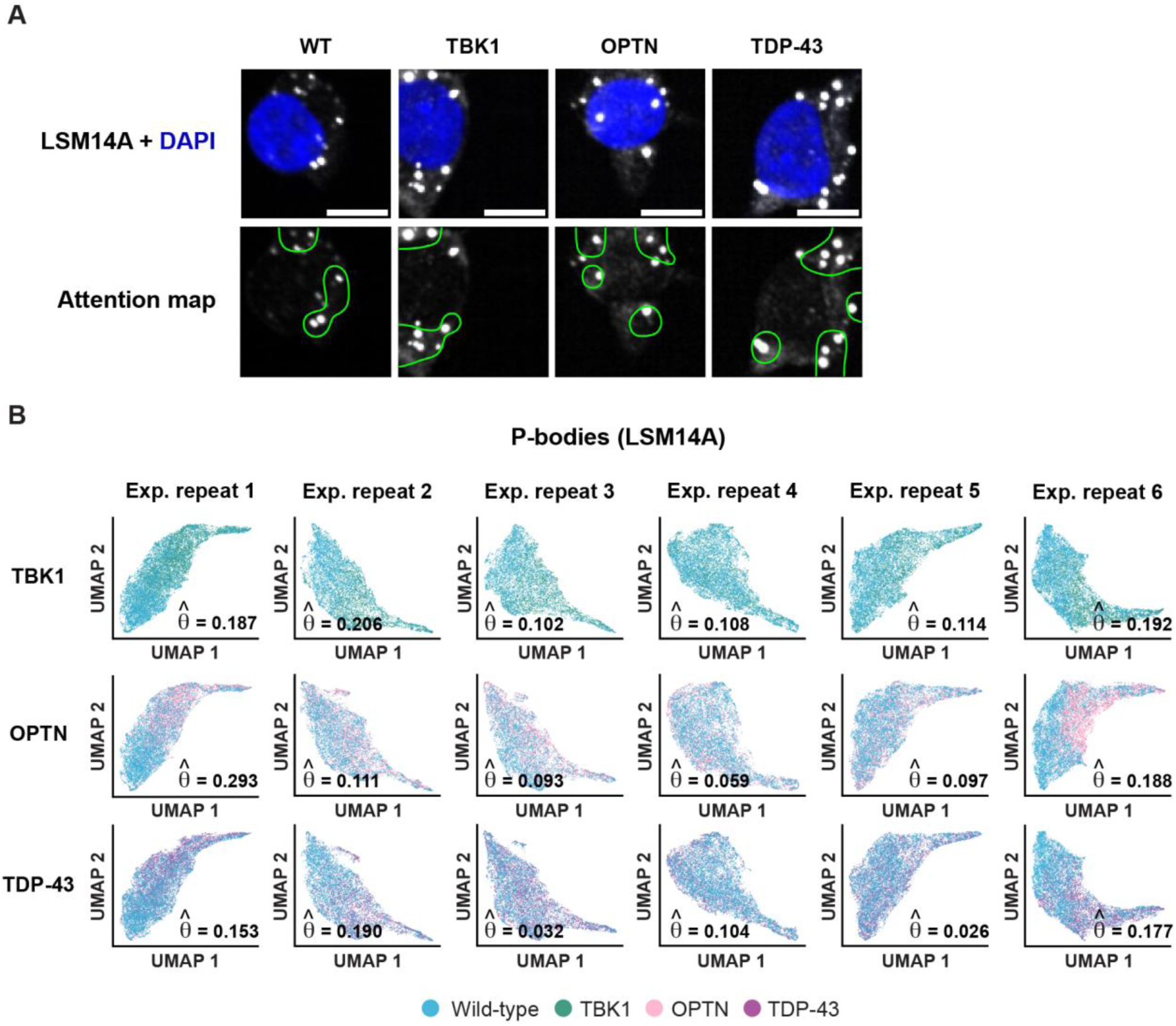
Altered topography of P-bodies marked by LSM14A in the iNDI lines. **(A)** Representative images of P-bodies marked by LSM14A across iNDI lines. Nucleus (hoechst 33342, blue) and overlaid model’s attention maps. Scale bar - 10 µm. **(B)** UMAP visualization of LSM14A changes per experimental repeat across iNDI lines. ∼3300-6600 image tiles per condition/experimental repeat. (***θ̂***) the effect size estimate.

**Supplementary Figure 15.**
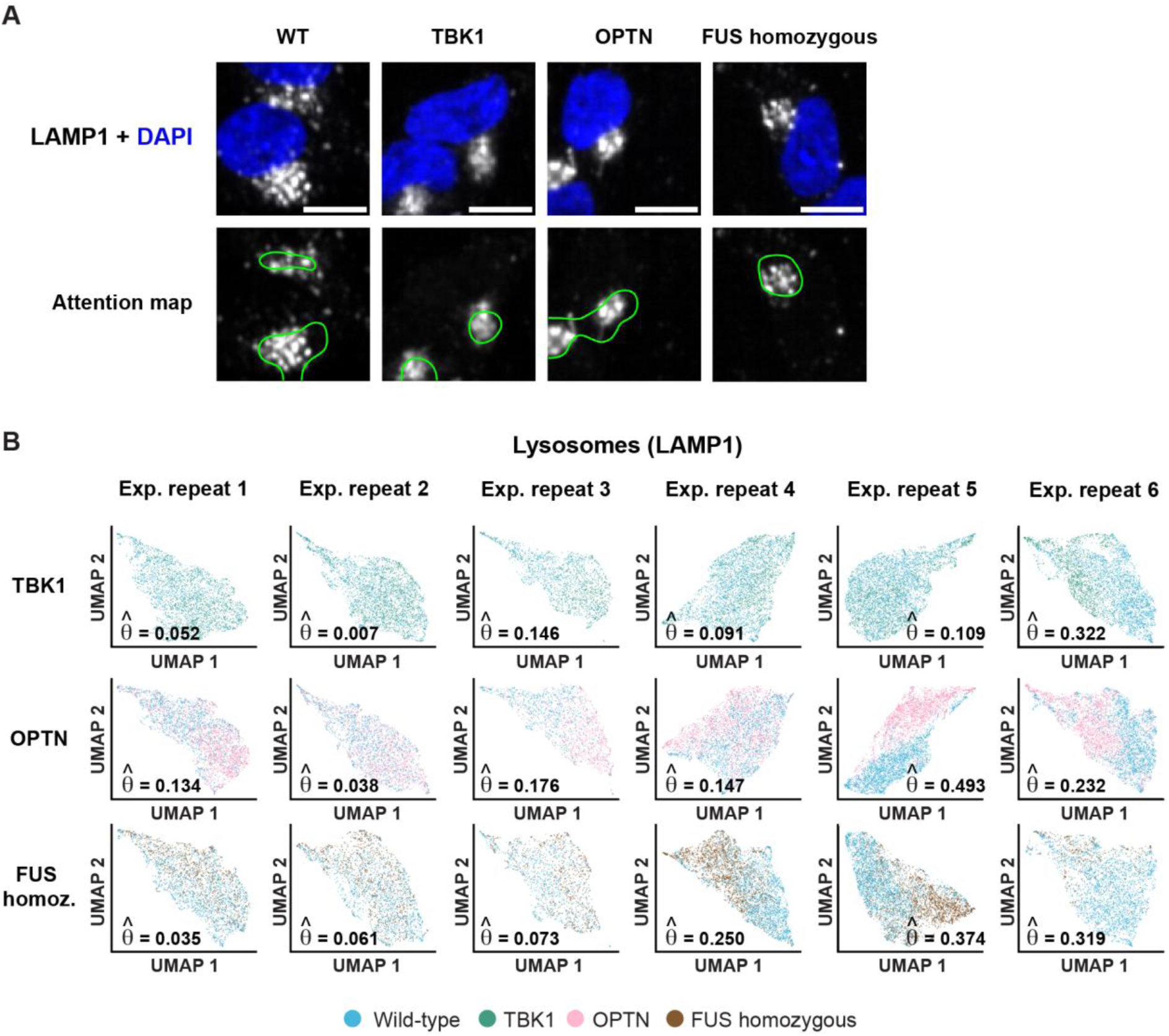
Altered topography of lysosomes in the iNDI lines. **(A)** Representative images of lysosomes marked by LAMP1 across iNDI lines. Nucleus (hoechst 33342, blue) and overlaid model’s attention maps. Scale bar - 10 µm. **(B)** UMAP visualization of LAMP1 changes per experimental repeat across iNDI lines. ∼600-2700 image tiles per condition/experimental repeat. (***θ̂***) the effect size estimate.

**Supplementary Figure 16.**
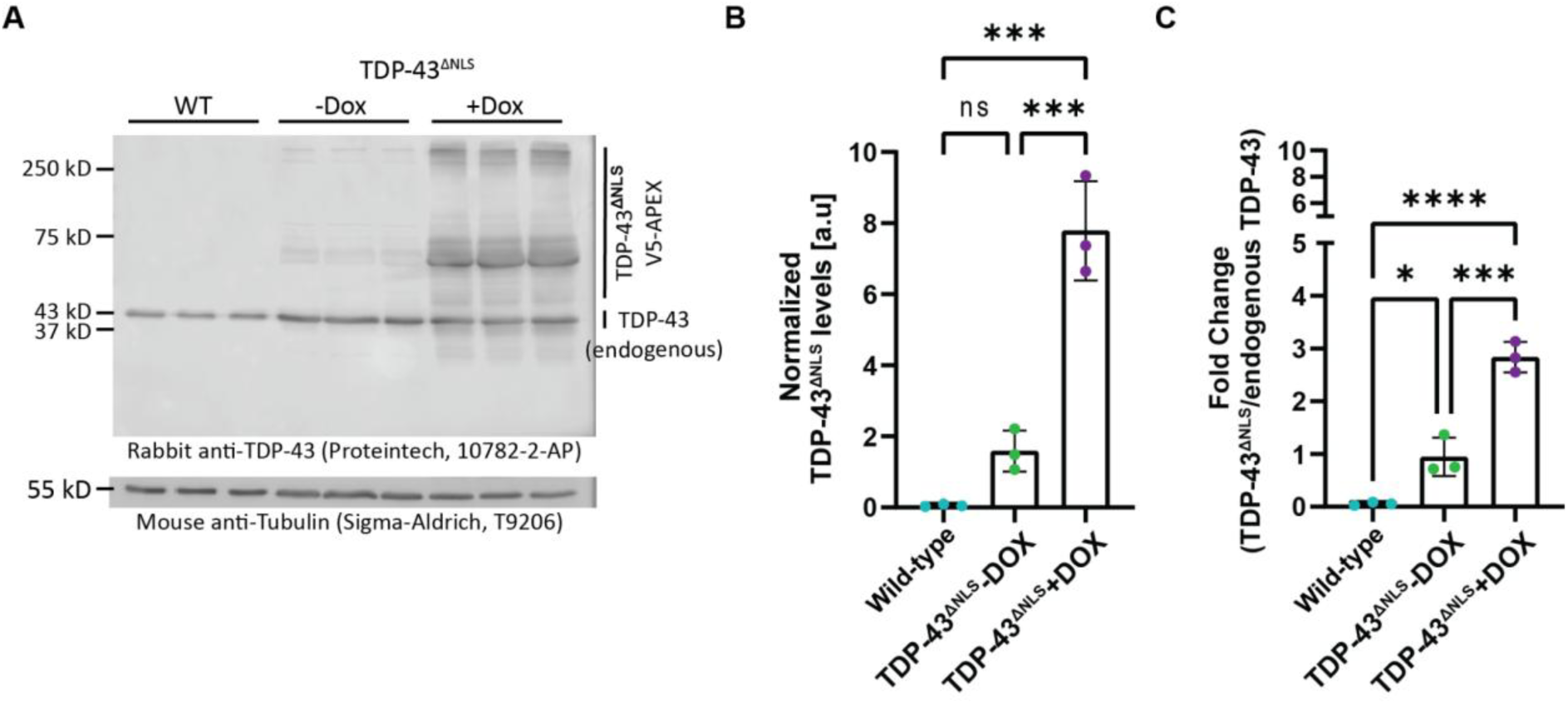
Protein levels of TDP-43 in wild-type and TDP-43^ΔNLS^-expressing neurons. **(A)** Western blot analysis of endogenous TDP-43 (∼43 KDa) and TDP-43^ΔNLS^-V5-APEX protein and protein oligomers (45-250 KDa bands) expression in 8-day-old wild-type (WT) neurons or neurons carrying TDP-43^ΔNLS^-V5-APEX, uninduced (-Dox) or after 24 hours of doxycycline induction (+Dox). Tubulin levels (55 KDa) were tested as loading control for normalization. **(B)** Quantification of TDP-43^ΔNLS^ levels. **(C)** Fold change of normalized TDP-43^ΔNLS^ levels relative to normalized endogenous TDP-43. P-value: * < 0.05, *** < 0.001, **** < 0.0001, ns = non-significant, One-way ANOVA with Benferroni’s multiple comparison test.

**Supplementary Figure 17.**
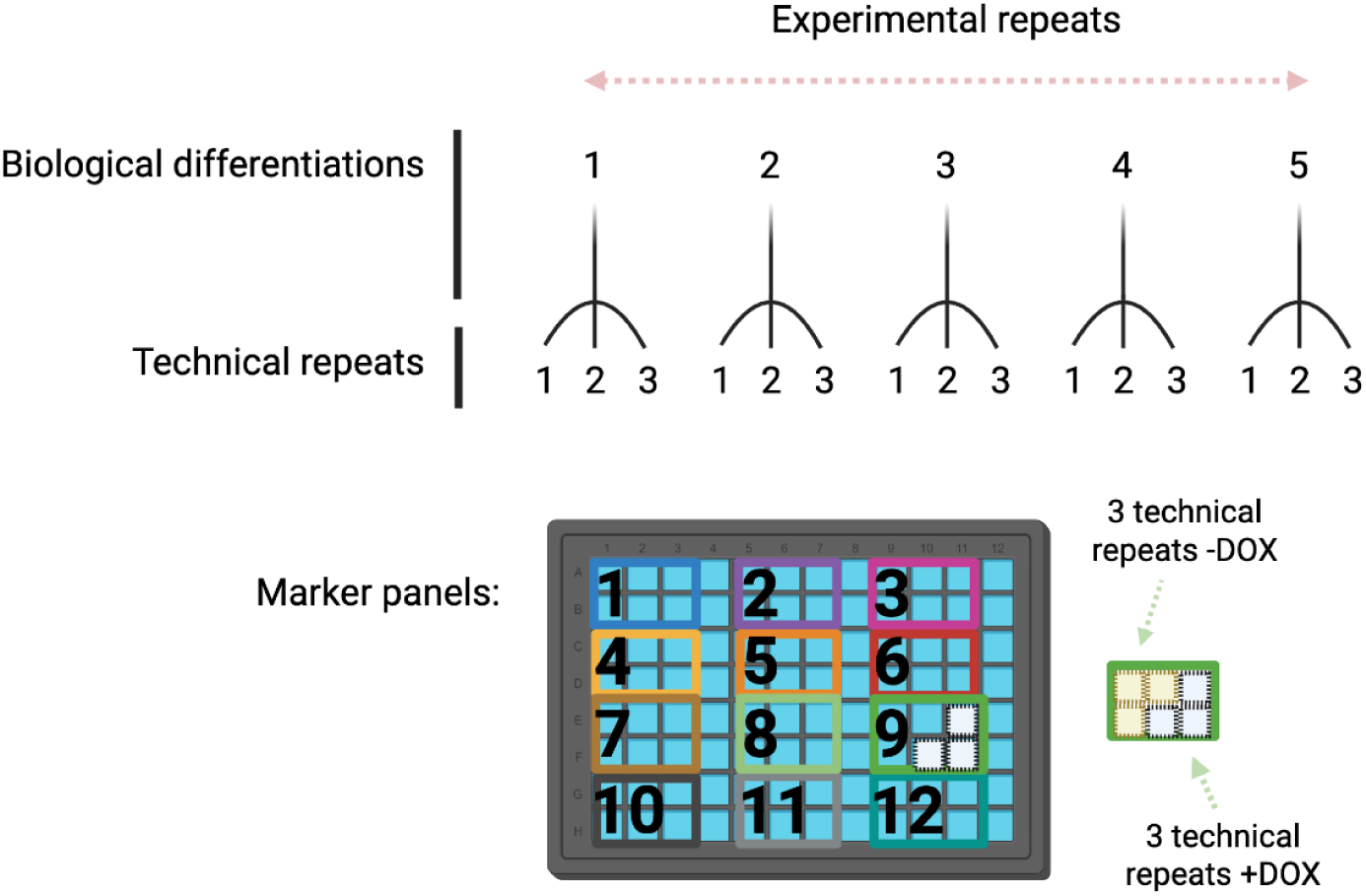
Experimental design of inducible TDP-43^ΔNLS^ related to. **Figure 4**. Five experimental repeats (biological differentiations), with three technical repeats (wells) each. TDP-43^ΔNLS^ without or with induction (±doxycycline, DOX). 12 marker panels per plate. Cells were seeded into PLO-coated 96-well plates on day 4 post-differentiation and fixed on day 8 of differentiation. 250 site images were acquired per well. An exemplary plate design from a single experimental repeat (biological differentiation) is depicted.

**Supplementary Figure 18.**
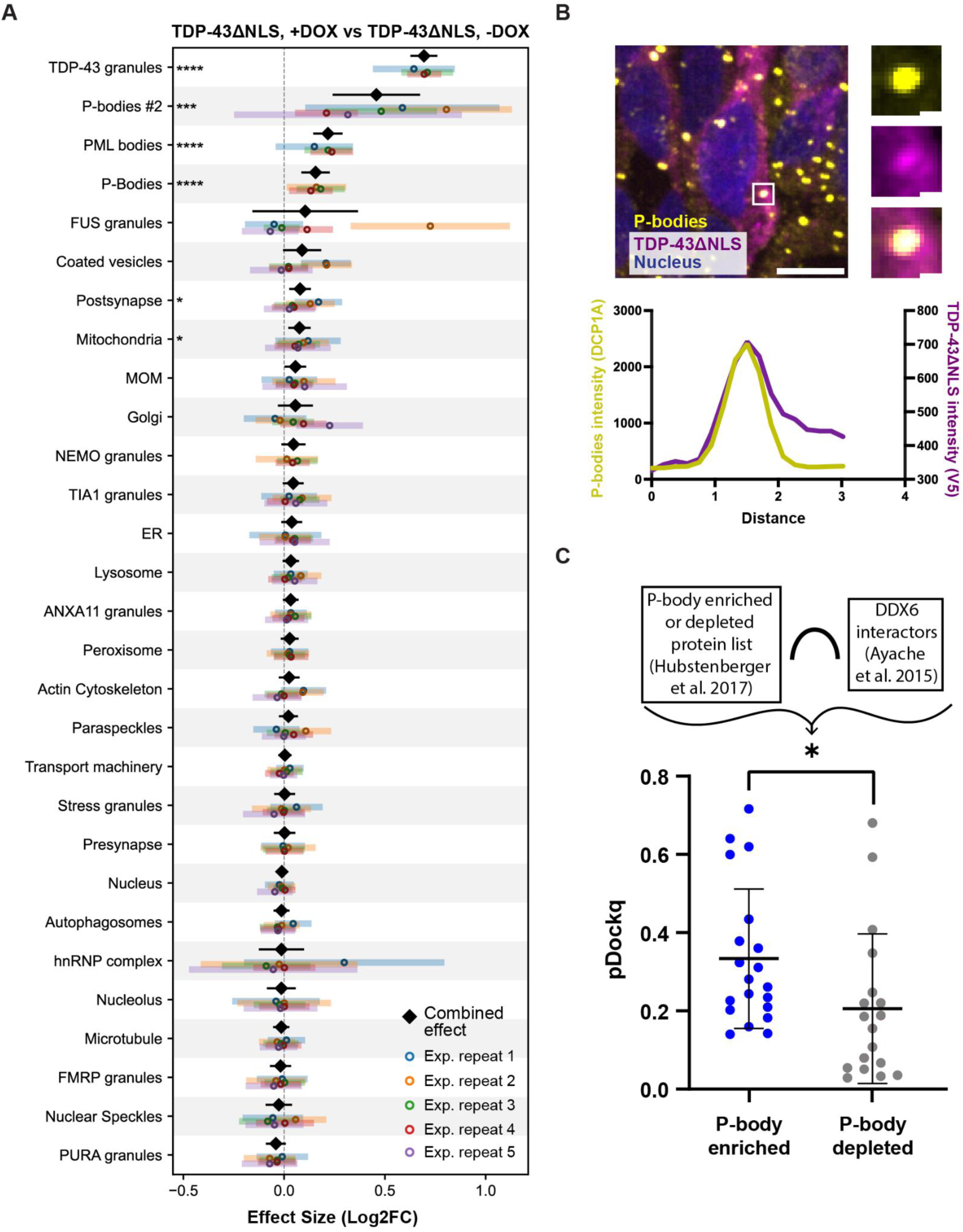
Cytoplasmic TDP-43–P-body interaction analyses in 8-day iPSC-derived inducible TDP-43^ΔNLS^ neurons. **(A)** Stacked forest plot of organelle effect sizes (log₂ fold change) in TDP-43^ΔNLS^ neurons with doxycycline (+DOX) relative to without (–DOX). Five experimental repeats; 29 organelles analyzed. Colored dots represent experimental repeat estimates, with horizontal bars showing 95% CI. The combined effect estimate (µ) is shown as a black diamond with its 95% meta-analytic CI. Adjusted p-values: * < 0.05, ** < 0.01, *** < 0.001, **** < 0.0001. ∼280-6700 image tiles per organelle/condition/experimental repeat. **(B)** Representative image showing colocalization of cytoplasmic TDP-43^ΔNLS^ (V5 tag, purple) with P-bodies (DCP1A, yellow). Lens ×63; scale bar, 10 µm. Insets show merged and separated channels of TDP-43^ΔNLS^ and a single P-body (scale bar, 1 µm) with corresponding intensity profiles. **(C)** Predicted protein–protein interactions between TDP-43 and P-body proteins using SpeedPPI. Y-axis: pDockQ interaction confidence score (Methods). P-body–enriched proteins (blue) or depleted proteins (grey) were defined by Hubstenberger et al. and identified as DDX6 interactors by immunoprecipitation. Mean: black horizontal line; error bars: SD. P-value * < 0.05, Mann–Whitney U test.

**Supplementary Figure 19.**
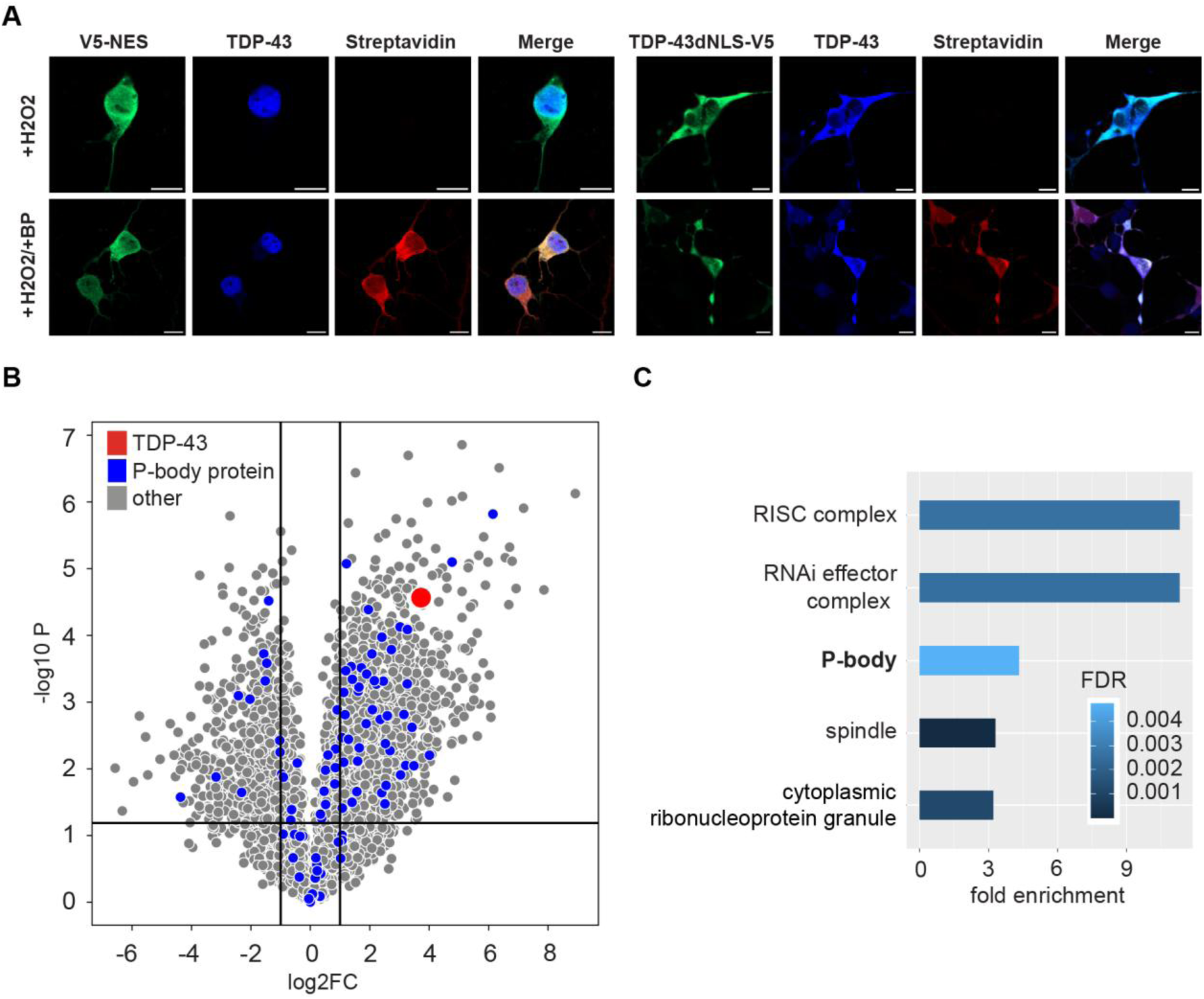
Validation of APEX construct activity in 8-day iPSC-derived neurons. **(A)** Immunostaining of neurons overexpressing APEX-V5-NES (+DOX, top) or TDP-43^ΔNLS^-V5-APEX (+DOX, bottom) without or with biotin phenol (BP) in the culture medium. V5-tagged construct (green), endogenous TDP-43 (blue), and biotinylated proteins (streptavidin, red). Lens ×63; scale bar, 10 µm. **(B)** Volcano plot of proteins enriched near TDP-43^ΔNLS^ compared to a cytoplasmic control construct (NES) after proximity labeling proteomics in 8-day neurons. Data from one of two independent experimental repeats (data of the other experiment shown in Figure 4G), each with 3–4 technical repeats per condition. P-body proteins (blue) as reported in ref. ^116^; TDP-43 as positive control (red). Unpaired two-sided t-test. **(C)** Gene Ontology (GO) cellular component analysis (see Methods) of proteins interacting with TDP-43^ΔNLS^ in two independent proximity labeling experiments. The top five GO terms are shown. X-axis: statistical significance (FDR); colors: fold change.

**Supplementary Figure 20.**
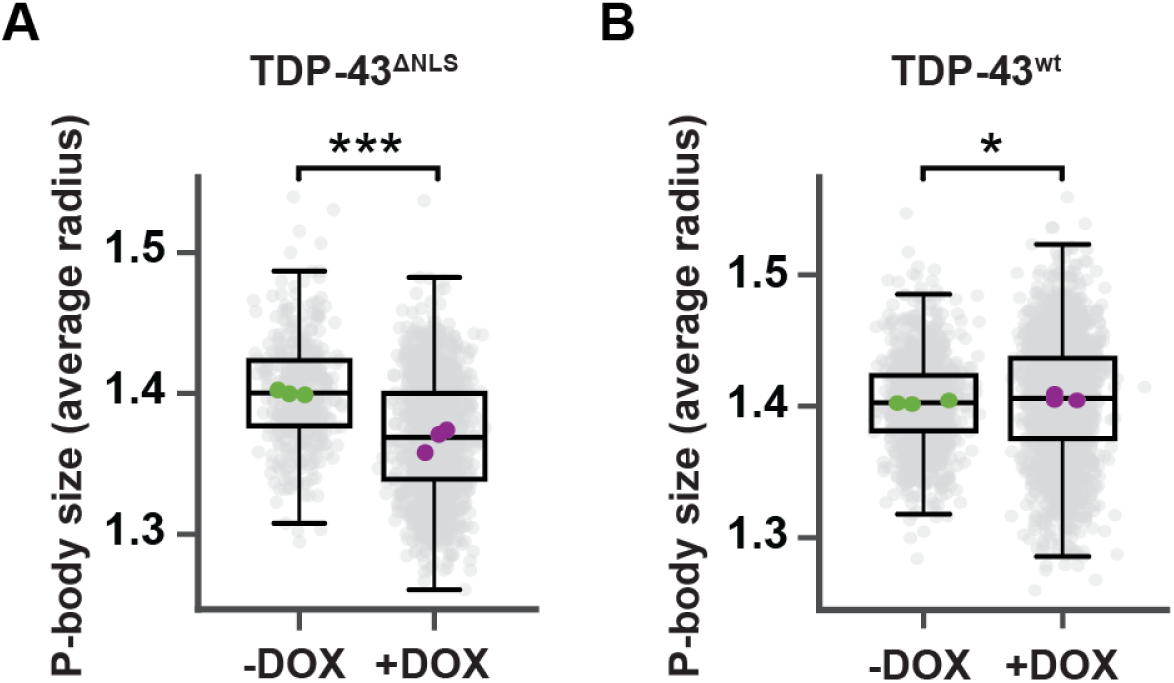
TDP-43^ΔNLS^ overexpression affects P-body size more than TDP-43^wt^ overexpression. CellProfiler quantification of P-body size in polyclonal day 8 iPSC-derived neurons inducibly overexpressing **(A)** TDP-43^ΔNLS^ or **(B)** TDP-43^wt^ (±DOX; means = 1.4 (-DOX) vs. 1.368 (+DOX) or 1.403 (-DOX) vs. 1.406 (+DOX), respectively); P-body marker - DCP1A. Colored dots - experimental repeat means (-DOX; green, +DOX; purple); gray dots - site-level measurements. Three experimental repeats per condition. *p < 0.05, ***p < 0.001. Linear mixed-effects model with DOX as a fixed effect and experimental repeat as random intercept.

**Supplementary Figure 21.**
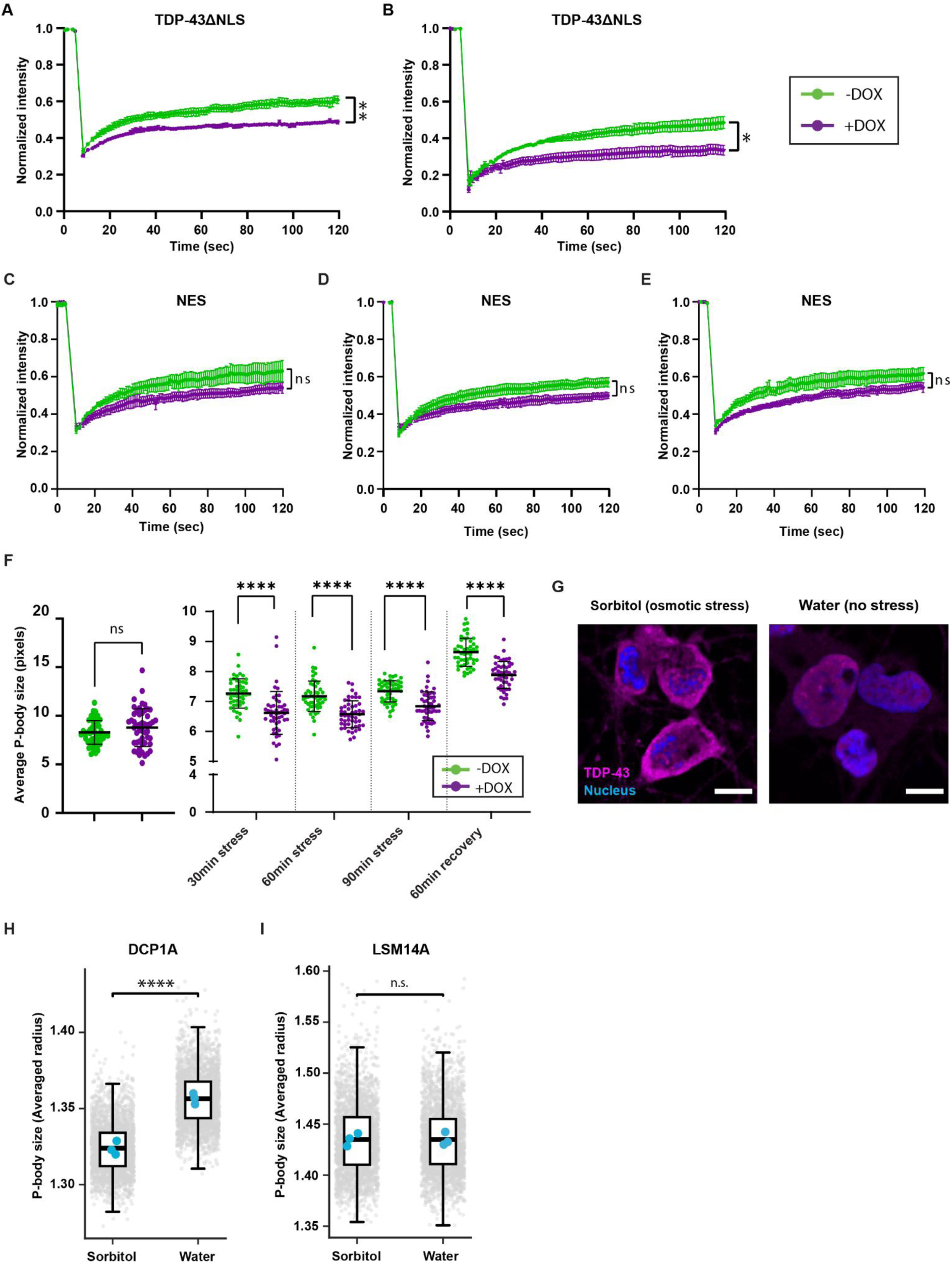
Cytoplasmic TDP-43 disrupts P-body properties under stress in different cellular models. FRAP analysis of YFP-DCP1A–labeled P-bodies in U2OS cells, partially photobleached ±DOX. **(A, B)** TDP-43^ΔNLS^ or **(C-E)** NES control. Fluorescence was normalized to baseline and recovery monitored over time (n = 4). Each data point represents the mean of 3–5 fields of view per repeat, with 2–9 P-bodies quantified per field. Two-way ANOVA with repeated measures and Geisser–Greenhouse correction; p-value * < 0.05, ** < 0.01; ns = not significant. **(F)** CellProfiler quantification of P-body size in U2OS cells expressing YFP-DCP1A with or without cytoplasmic TDP-43 (±DOX) under sodium arsenite stress (0.25 mM, 90 min) and recovery (60 min). n = 50 image fields/condition. Unpaired t-test; P-value ** < 0.01, **** < 0.0001; ns = not significant. Horizontal black lines indicate means. **(G)** Immunostaining of day 8 iPSC-derived neurons (iW11) treated with carrier (water) or sorbitol (0.4 M, 2 hr) to induce mislocalization of endogenous TDP-43 (magenta) into the cytoplasm. Nucleus stained with Hoechst 33342 (blue). Lens ×63; scale bar, 10 µm. CellProfiler quantification of P-body size per cell, using **(H)** DCP1A or **(I)** LSM14A, in iW11 neurons under basal vs. osmotic stress conditions (means (water vs. sorbitol) = 1.36 vs. 1.328 (DCP1A) or 1.439 vs. 1.439 (LSM14A)). Blue dots: experimental repeat means; gray dots: 250 site-level measurements per experimental repeat; three experimental repeats per condition. p-value **** < 0.0001. Linear model with sorbitol as fixed effect and experimental repeat as covariate; random-effect variance = 0.

**Supplementary Figure 22.**
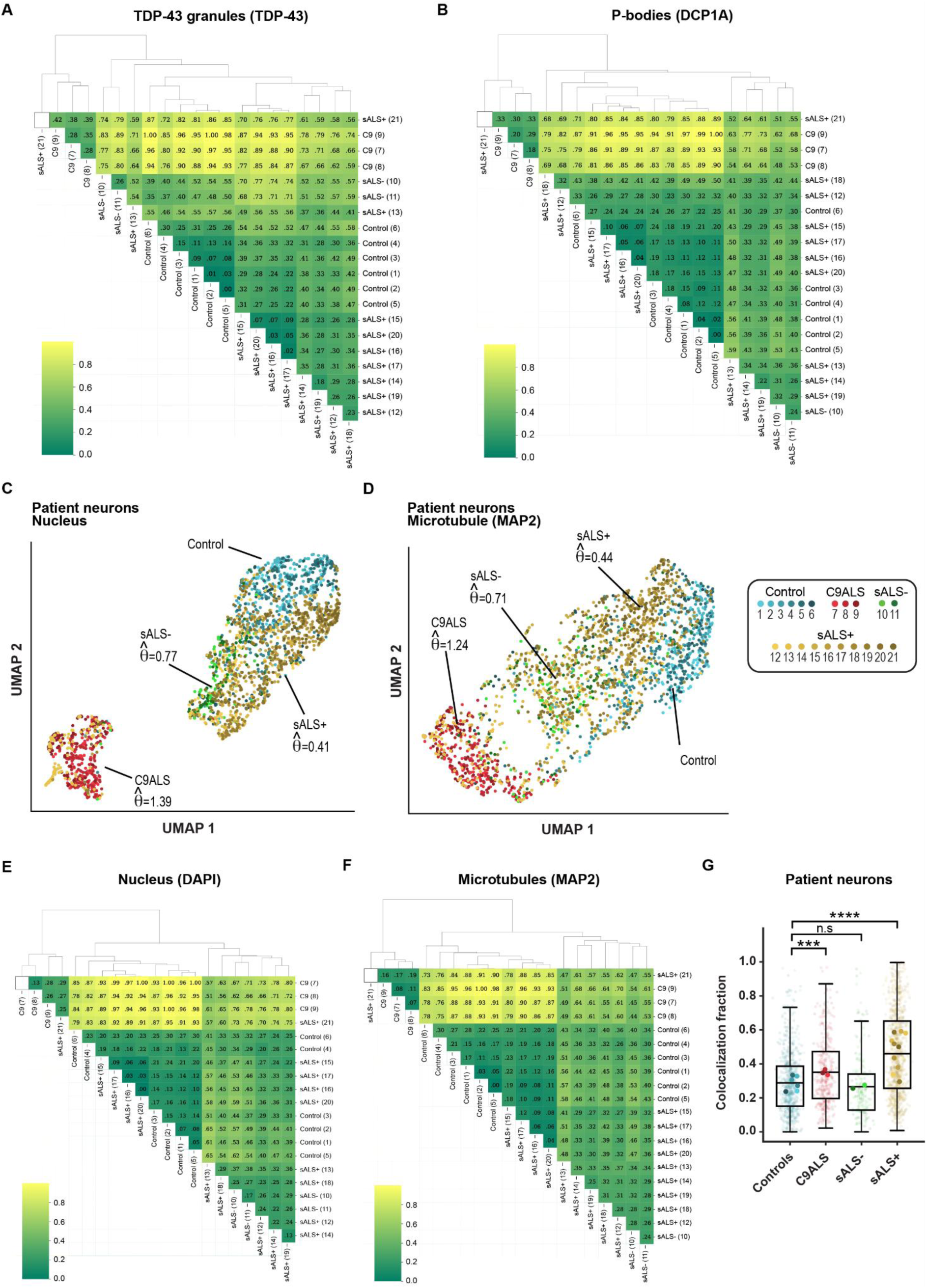
Extended study in day 60 human motor neurons. Pairwise Euclidean distances between NOVA-derived embeddings of **(A)** TDP-43 granules and **(B)** P-bodies were computed in the organellar landscapes shown in **Figure 5C**, **D** for each pair of human subjects. Heatmap values represent the median distance, rescaled to the [0,1] interval using min–max normalization. A value of 0 indicates the closest subjects (highest similarity, darker colors), and 1 the most distant (lowest similarity, lighter colors). Rows and columns were clustered by agglomerative hierarchical clustering of the scaled distance values (Euclidean distance, average linkage). Dendrograms depict the hierarchical relationships among human subjects. UMAP projections of day 60 motor neurons from a cohort of 21 human subjects, showing **(C)** the nucleus (hoechst 33342) or **(D)** Microtubule (MAP2). Points - single image tiles. (*θ̂*) the effect size estimate. Pairwise Euclidean distances between NOVA-derived embeddings of **(E)** nucleus and **(F)** microtubules. **(G)** Boxplots of P-body-positive (DCP1A) pixels that are also positive for TDP-43. Colocalization fraction in human subjects day 60 motor neuron images in C9ALS, sALS-, or sALS+ vs. healthy controls (mean fraction overlap = 0.35, 0.265 or 0.46 vs. 0.288, respectively). ∼60-160 image tiles per human subject; Linear fixed-effects model with human subject-clustered standard errors (fallback used); *** p < 0.001, **** p < 0.0001. Large dots - condition mean per experimental repeat; small dots - individual images; Three human subjects per group.

**Supplementary Figure 23.**
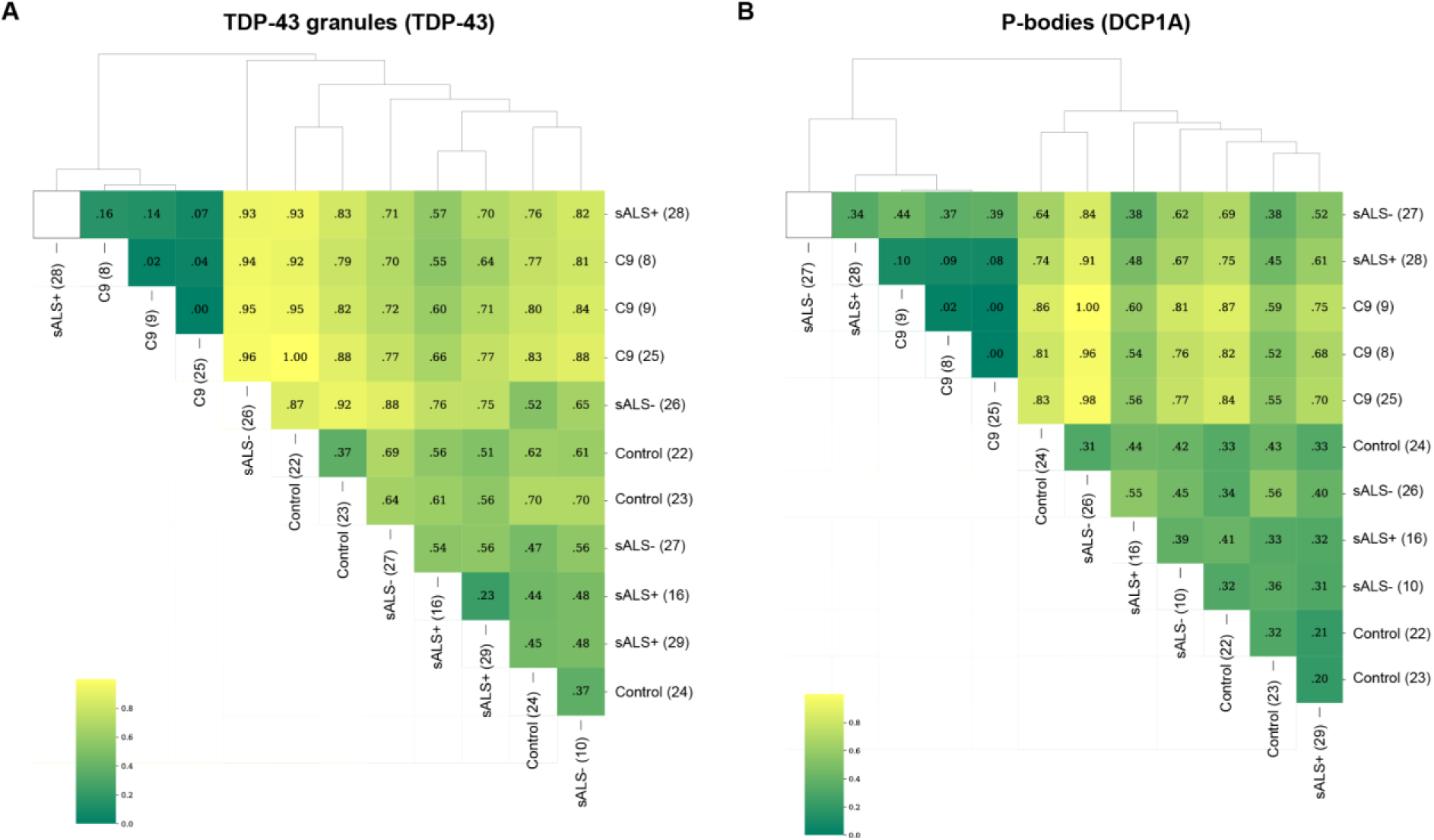
Distances for TDP-43 granules and P-bodies across the human subjects in the extended study of day 60 human motor neurons. Pairwise Euclidean distances between NOVA-derived embeddings of **(A)** TDP-43 granules and **(B)** P-bodies were computed in the organellar landscapes shown in **Figure 5G, H** for each pair of human subjects. Heatmap values represent the median distance, rescaled to the [0,1] interval using min–max normalization. A value of 0 indicates the closest subjects (highest similarity, darker colors), and 1 the most distant (lowest similarity, lighter colors). Rows and columns were clustered by agglomerative hierarchical clustering of the scaled distance values (Euclidean distance, average linkage). Dendrograms depict the hierarchical relationships among human subjects.

**Supplementary Figure 24.**
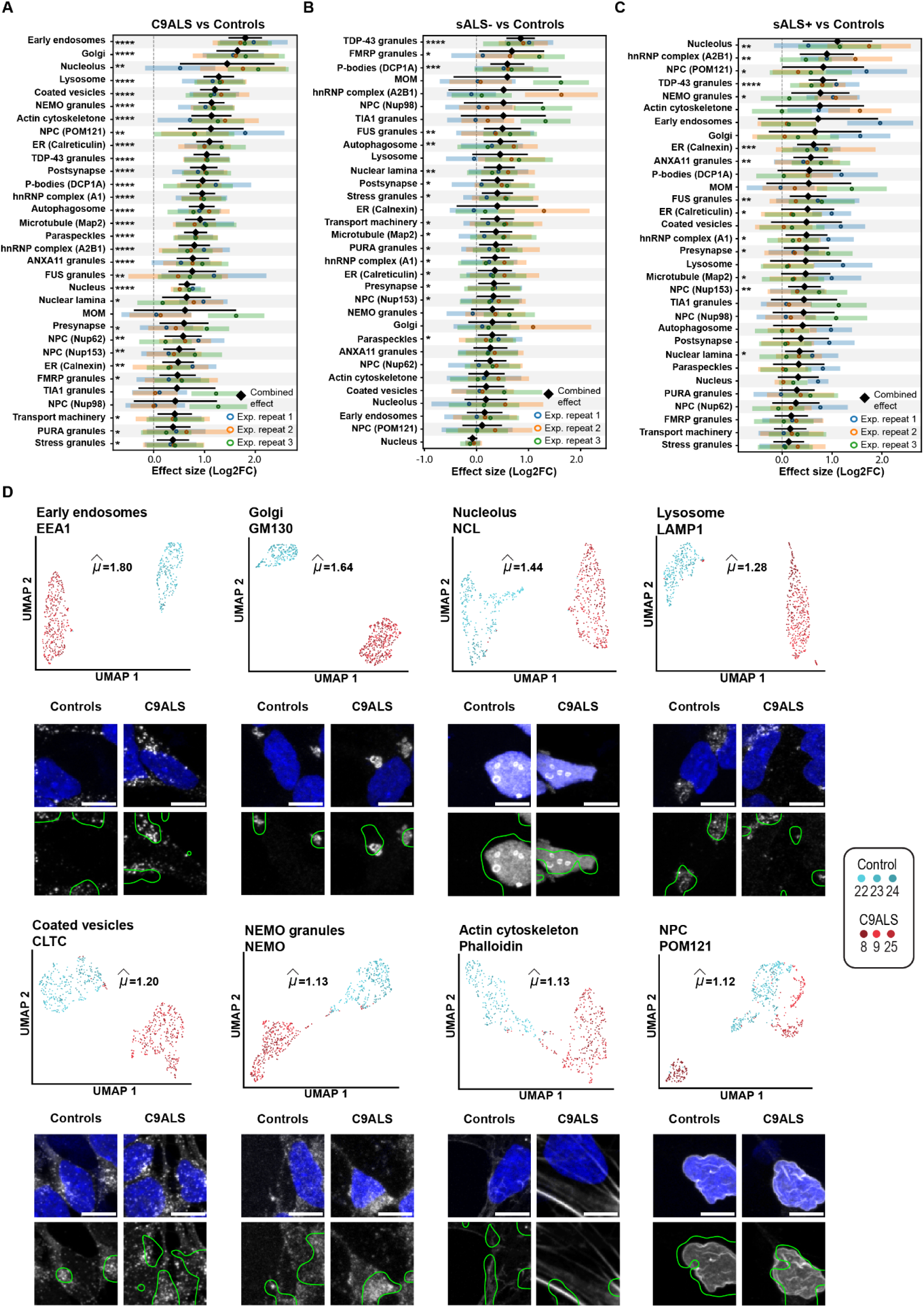
Comprehensive organellar response of day 60 human subject-derived motor neurons. Stacked forest plot of organelle effect sizes (log₂ fold change) in day 60 motor neurons from **(A)** C9ALS, **(B)** sALS without cytoplasmic TDP-43 (sALS–), and **(C)** sALS with cytoplasmic TDP-43 (sALS+), relative to healthy controls. Twelve human subjects were analyzed (three per group); 32 organelles were studied. Colored dots represent group-wise effect estimates, with horizontal bars indicating 95% CI. The combined effect estimate (*μ̂*) is shown as a black diamond with its 95% meta-analytic CI, accounting for between-experimental repeat variance. Adjusted P-values: * < 0.05, ** < 0.01, *** < 0.001, **** < 0.0001. ∼45-170 image tiles per organelle/human subject. **(D)** UMAP projection of day 60 motor neurons from C9ALS patients (n = 3) and healthy controls (n = 3), highlighting early endosomes, Golgi, nucleolus, lysosome, coated vesicles, NEMO granules, actin cytoskeleton and the nuclear pore complex (NPC), related to the autophagy and endosomal–lysosomal pathway. Each point represents a single image tile (∼55 tiles per human subject per organelle). Effect size estimates (µ) are indicated. Representative images with nuclei (Hoechst 33342) and overlaid model attention maps for each organelle/group. Lens ×63; scale bar, 10 µm.

**Supplementary Figure 25.**
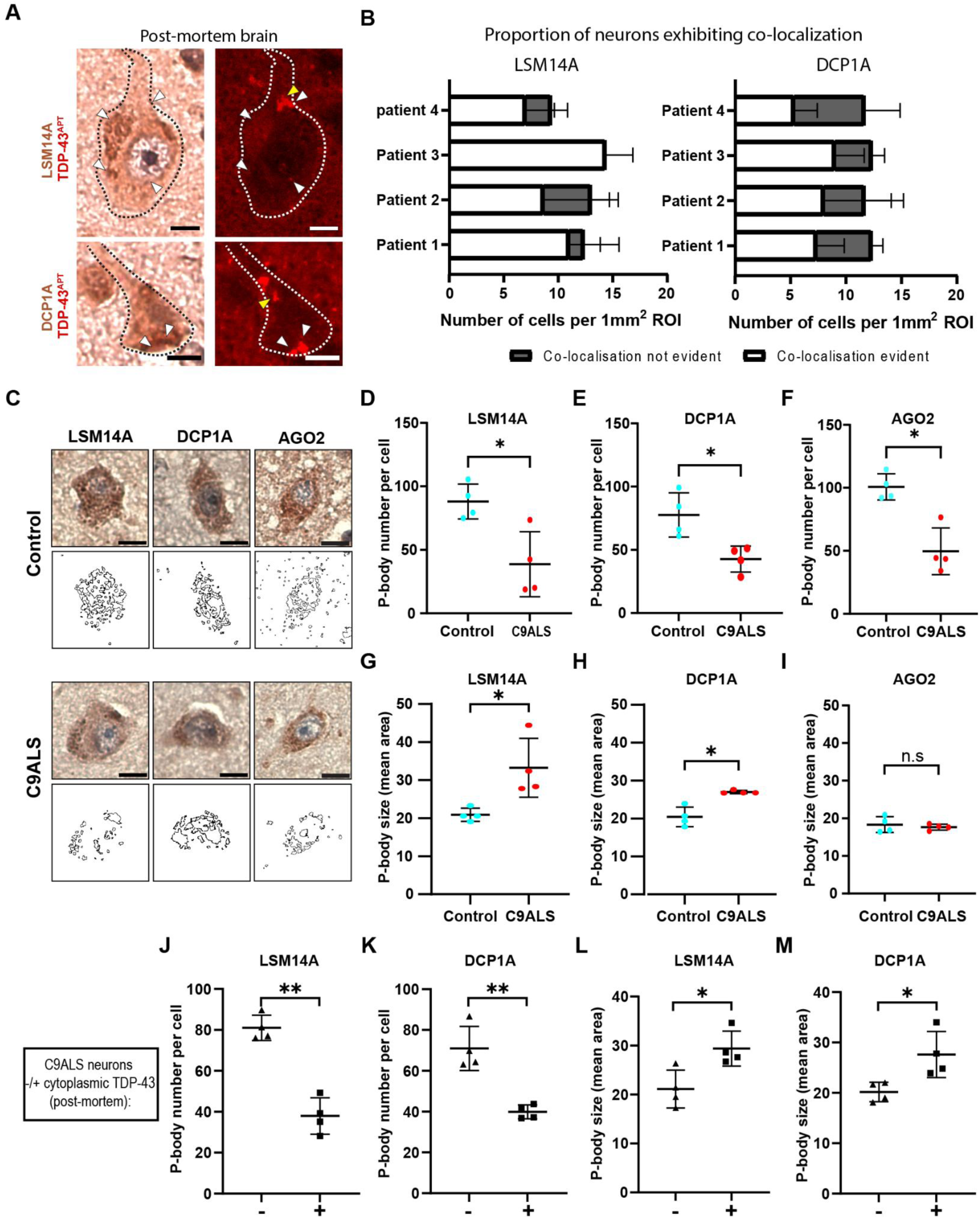
TDP-43 interacts with P-bodies in ALS post-mortem tissue. Images of C9ALS neurons showing cytoplasmic TDP-43 (aptamer^139^, red) colocalized with P-body markers LSM14A or DCP1A (chromogenic IHC). Yellow arrowheads: TDP-43 without colocalization; white arrowheads: with colocalization. Scale bar, 10 µm. **(B)** Quantification of neurons with (white bars) or without (grey bars) TDP-43-P-body colocalization (LSM14A, left; DCP1A, right) across four patients per region of interest (ROI) = 1 mm². Mean of three ROIs/patient; error bars, SD; analysis by ImageJ. **(C)** Representative DAB IHC of P-body proteins (DCP1A, LSM14A, AGO2) in C9ALS motor cortex (lower panels) vs. controls (upper panels). Scale bar, 10 µm. Quantification of mean P-body number **(D–F)** and size **(G–I)** per cell per subject (4 C9ALS patients (red) compared to 4 controls (blue)). Median P-body number (control vs C9ALS) = 85.95 vs 31.20 (LSM14A) or 75.15 vs 45.50 (DCP1A) or 98.20 vs 43.90 (AGO2); median P-body size (control vs C9ALS) = 20.60 vs 30.35 (LSM14A) or 20.00 vs 26.80 (DCP1A) or 18.00 vs 17.75 (AGO2). Mann–Whitney U test; *p < 0.05; ns = not significant. Scatter plots show means (black lines) ± SD. Each point represents one subject. Quantification of mean number **(J, K)** and size area **(L, M)** of P-bodies (LSM14A or DCP1A) in neurons without (-, triangle) or with (+, square) cytoplasmic TDP-43 within each of the C9ALS patients (mean P-body number (- vs. +) = 81.02 vs. 37.252 (LSM14A) or 68.04 vs. 39.9 (DCP1A); mean P-body size area (- vs. +) = 21.12 vs. 30 (LSM14A) or 20.19 vs. 26.99 (DCP1A)). Nested t-test; *p < 0.05, **p < 0.01. Scatter plots show means (black lines) ± SD. Each point represents one subject (n = 4).

## Material & Methods

### Cell lines

1. **Cancer cell lines and iNDI lines**

2. **Patient-derived iPSC lines including demographic Information**

| # | Cell line | Reference / gift |
| --- | --- | --- |
| 1 | KOLF2.1J human iPSC | 169 |
| 2 | TDP-43<sup>ΔNLS</sup>-V5-APEX NGN2-iPSC | This study |
| 3 | TDP-43<sup>wt</sup>-V5-APEX NGN2-iPSC | This study |
| 4 | NES-V5-APEX NGN2-iPSC | This study |
| 5 | U2OS stably expressing YFP-DCP1A | A gift from Nancy Kedersha<br>Pavel Ivanov and Paul<br>Anderson, Harvard Medical<br>School |
| 6 | U2OS stably expressing UBAP2L-ruby and /inducibly<br>expressing TDP-43<sup>ΔNLS</sup>-clover | <sup>111</sup>A gift from Don W<br>Cleveland, University of<br>California, San Diego |
| 7 | *TARDBP* M337V SNV/SNV | 98 |
| 8 | *OPTN* Q398E SNV/SNV | <sup>98</sup> Gift from Michael Ward,<br>National Institutes of Health |
| 9 | *TBK1* E696K SNV/SNV | 98 |
| 10 | *FUS* R495* SNV/SNV | 98 |
| 11 | *FUS* R495* SNV/WT | 98 |
| 12 | *FUS* R495* Rev/WT | 98 |
| 13 | *SNCA* E46K SNV/SNV | 98 |

**2. Patient-derived iPSC lines including demographic Information**
| # | iPSC Line<br>Name | Source | Clinical Diagnosis | Age at<br>Time of | Sex | Origin |
| --- | --- | --- | --- | --- | --- | --- |

|  |  |  |  | Collection |  |  |
| --- | --- | --- | --- | --- | --- | --- |
| 1 | CS0002 | Cedars-Sinai | Non-neurologic control | 51 | Male | PBMC |
| 2 | CS0201 | Cedars-Sinai | Non-neurologic control | 56 | Female | PBMC |
| 3 | CS1ATZ | Cedars-Sinai | Non-neurologic control | 60 | Male | PBMC |
| 4 | CS2AE8 | Cedars-Sinai | Non-neurologic control | 50 | Female | PBMC |
| 5 | CS8PAA | Cedars-Sinai | Non-neurologic control | 58 | Female | PBMC |
| 6 | CS9XH7 | Cedars-Sinai | Non-neurologic control | 53 | Male | PBMC |
| 7 | EDi022-A | Cedars-Sinai | Non-neurologic control | 79 | Male | PBMC |
| 8 | EDi029-A | Cedars-Sinai | Non-neurologic control | 80 | Male | PBMC |
| 9 | EDi037-A | Cedars-Sinai | Non-neurologic control | 79 | Male | PBMC |
| 10 | CS0JGZ | Cedars-Sinai | sALS | 56 | Male | PBMC |
| 11 | CS3EPR | Cedars-Sinai | sALS | 60 | Female | PBMC |
| 12 | CS3XLK | Cedars-Sinai | sALS | 55 | Female | PBMC |
| 13 | CS4PFR | Cedars-Sinai | sALS | 72 | Female | PBMC |
| 14 | CS4ZCD | Cedars-Sinai | sALS | 53 | Male | PBMC |
| 15 | CS5JPF | Cedars-Sinai | sALS | 55 | Female | PBMC |
| 16 | CS5ZHY | Cedars-Sinai | sALS | 64 | Male | PBMC |
| 17 | CS6PYD | Cedars-Sinai | sALS | 69 | Female | PBMC |
| 18 | CS8JGP | Cedars-Sinai | sALS | 61 | Male | PBMC |
| 19 | CS9GXD | Cedars-Sinai | sALS | 68 | Male | PBMC |
| 20 | CS2FN3 | Cedars-Sinai | sALS | 58 | Male | PBMC |
| 21 | CS7TN6 | Cedars-Sinai | sALS | 33 | Male | PBMC |
| 22 | CS0JPP | Cedars-Sinai | sALS | 61 | Male | PBMC |
| 23 | CS4MLT | Cedars-Sinai | sALS | 54 | Male | PBMC |
| 24 | CS6ZU8 | Cedars-Sinai | sALS | 61 | Male | PBMC |
| 25 | CS0ANK | Cedars-Sinai | sALS | 65 | Male | PBMC |
| 26 | CS0BUU | Cedars-Sinai | *C9orf72* | 63 | Female | PBMC |
| 27 | CS2YNL | Cedars-Sinai | *C9orf72* | 60 | Male | PBMC |
| 28 | CS7VCZ | Cedars-Sinai | *C9orf72* | 64 | Male | PBMC |
| 29 | CS8RFT | Cedars-Sinai | *C9orf72* | 62 | Male | PBMC |

### Human post-mortem brain tissue samples

| Case Number | GUID/Block number | Source | Clinical Diagnosis | Age at Death | Sex |
| --- | --- | --- | --- | --- | --- |
| GBB-19-14 | NEUNC276RND | Target ALS post-mortem core | Control | 67 | Male |
| GBB-16-10 | NEUCG551GV1 | Target ALS post-mortem core | Control | 57 | Female |
| GBB-18-13 | NEUGV102NYT | Target ALS post-mortem core | Control | 63 | Female |
| GBB-17-22 | NEUGJ583BEE | Target ALS post-mortem core | Control | 70 | Male |
| LP005/2007 | AD | Sheffield Brain bank | Control | 63 | Male |
| LP085/2007 | C | Sheffield Brain bank | Control | 59 | Female |
| LP098/2007 | C | Sheffield Brain bank | Control | 68 | Male |
| PA135/2018 | A1 | Sheffield Brain bank | Control | 85 | Female |
| PA058/2022 | A19 | Sheffield Brain bank | Control | 68 | Female |
| PA154/2022 | A13 | Sheffield Brain bank | Control | 97 | Female |
| GBB-18-07<br>(Georgetown Case) | NEUGU437VNF | Target ALS post-mortem core | *C9orf72* | 66 | Female |
| GWF15-09 | NEUTP019MY9 | Target ALS post-mortem core | *C9orf72* | 61 | Female |
| GWF18-31 | NEUKN179PH8 | Target ALS post-mortem core | *C9orf72* | 62 | Female |
| GBB-18-17<br>(Georgetown case) | NEUBE992ZD1 | Target ALS post-mortem core | *C9orf72* | 68 | Male |
| LP045/2006 | A | Sheffield Brain bank | *C9orf72* | 46 | Male |
| LP066/2008 | B | Sheffield Brain bank | *C9orf72* | 59 | Female |
| LP005/2011 | C | Sheffield Brain bank | sALS | 76 | Female |
| NA009/1996 | L | Sheffield Brain bank | sALS | 78 | Female |
| LP020/2013 | A2 | Sheffield Brain bank | sALS | 78 | Male |
| LP024/2008 | B | Sheffield Brain bank | sALS | 63 | Female |
| LP027/2008 | C | Sheffield Brain bank | sALS | 70 | Female |
| LP039/2005 | B | Sheffield Brain bank | sALS | 57 | Female |

### Primary antibodies used

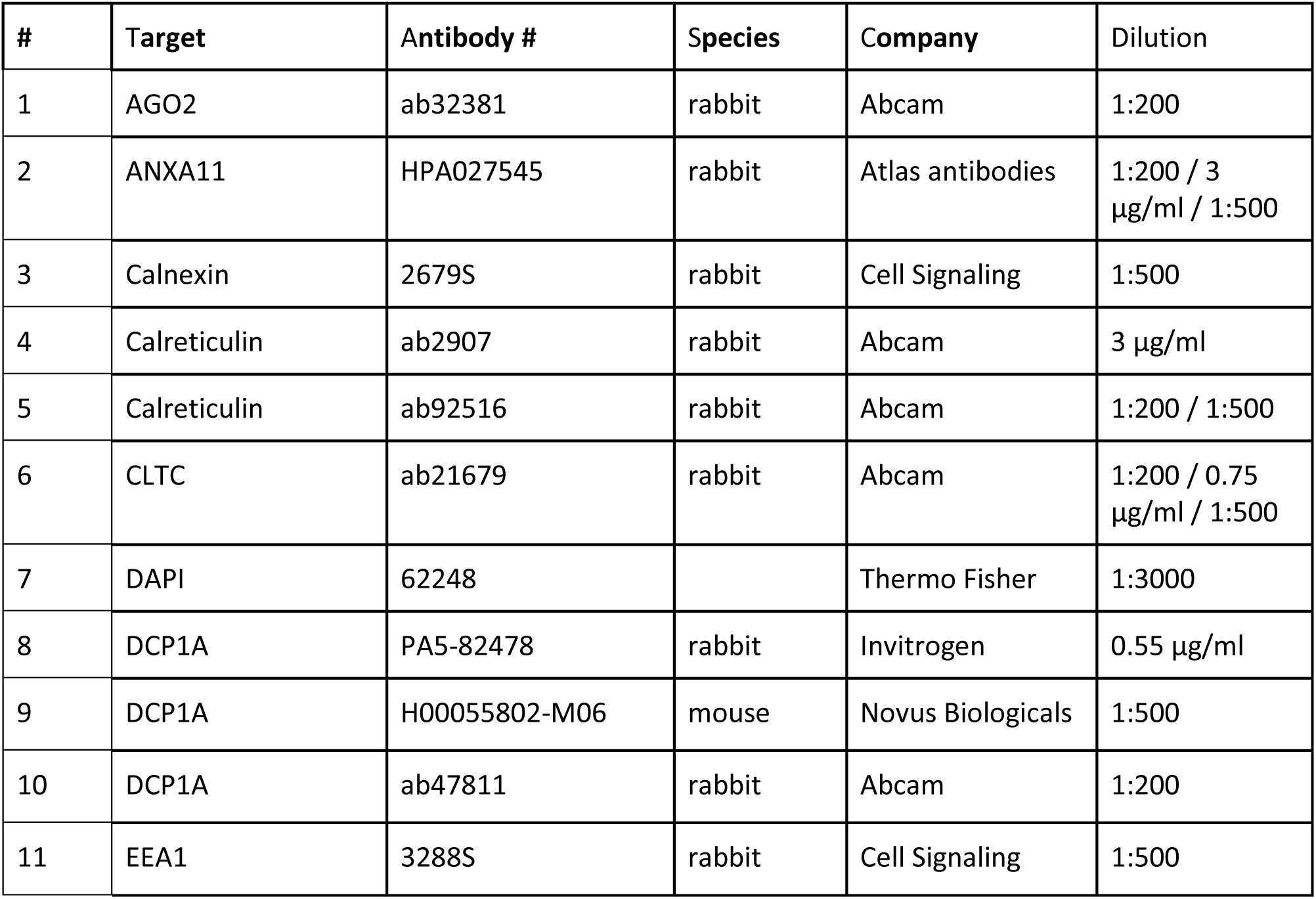
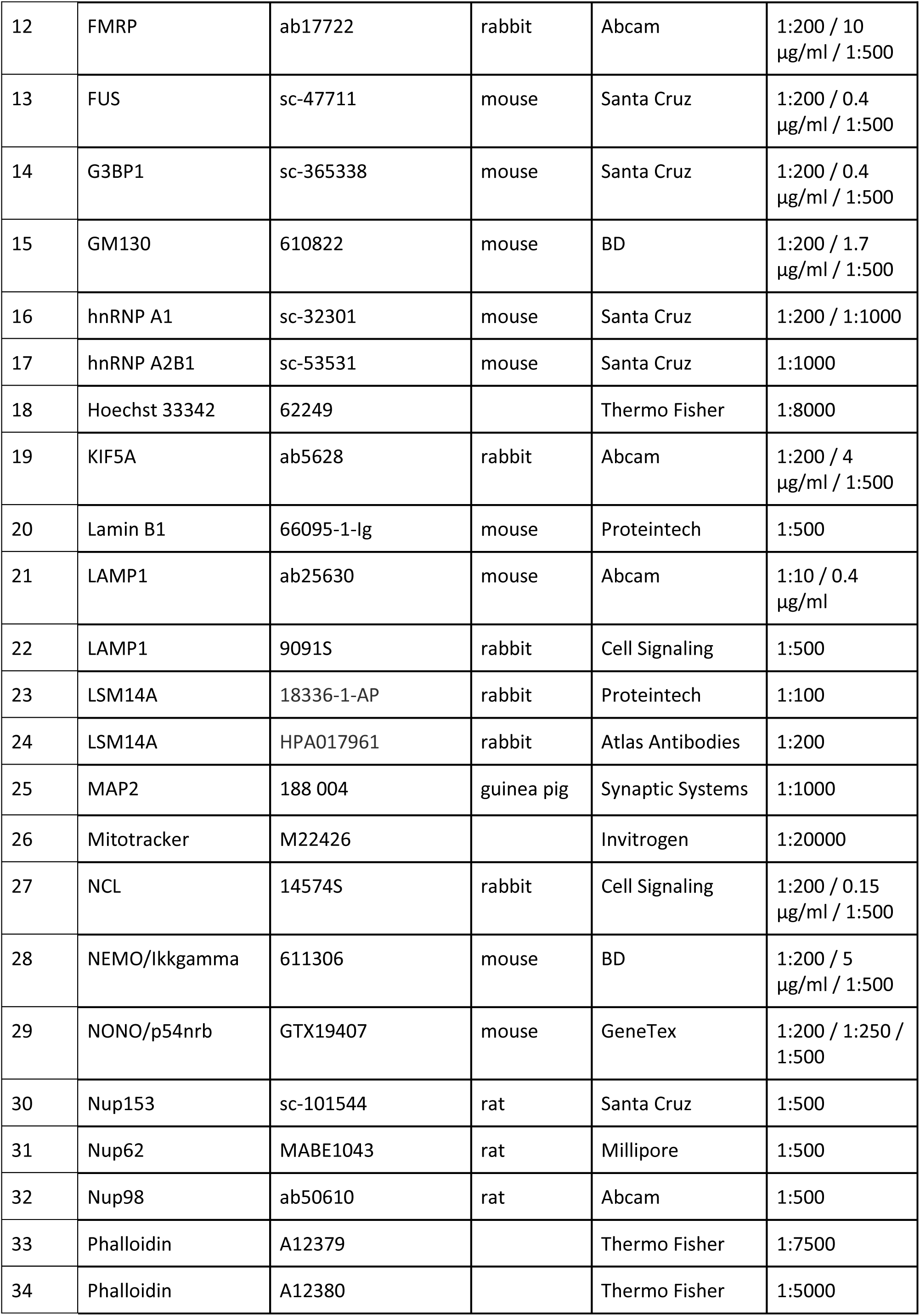
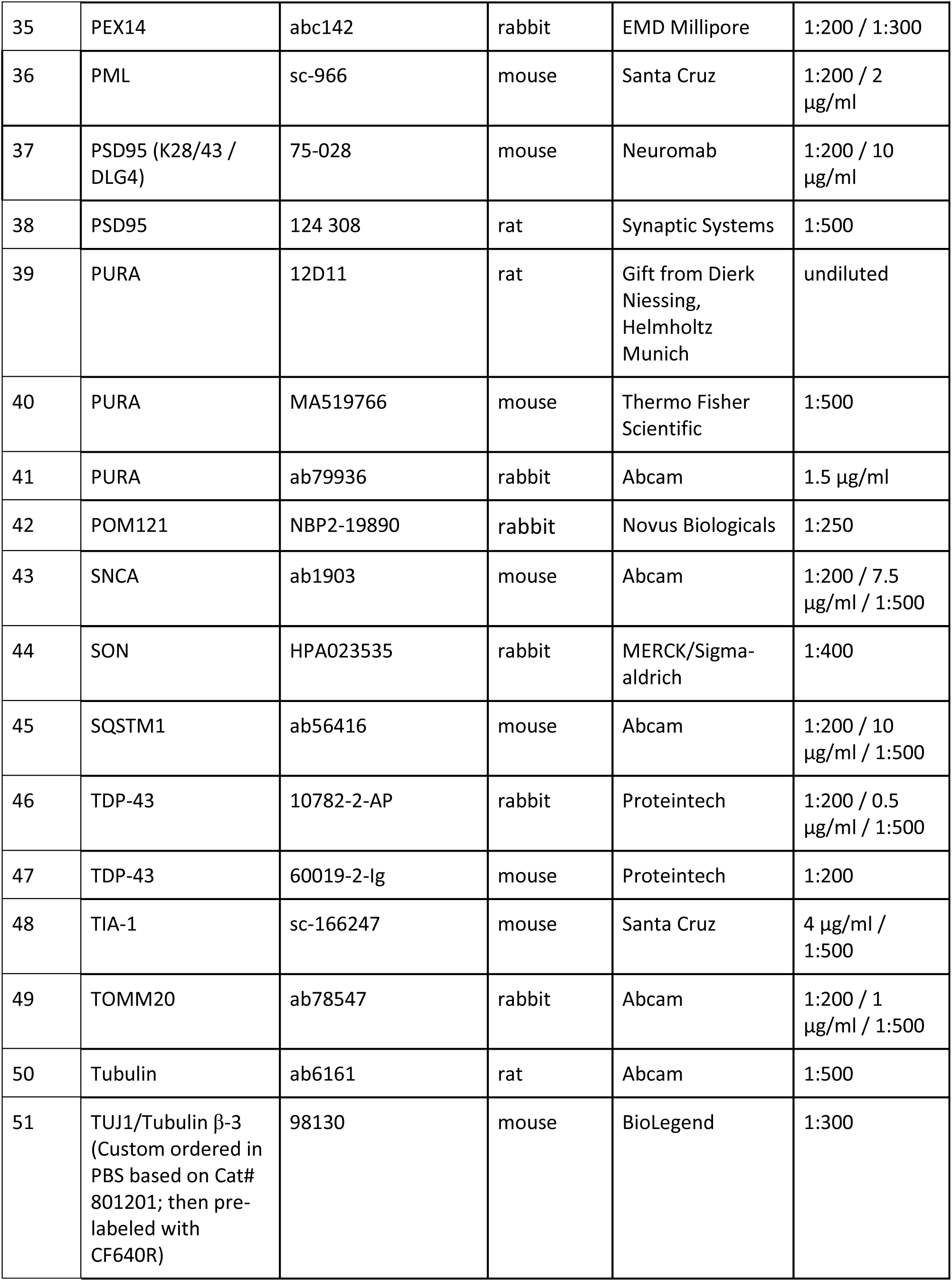

### Secondary antibodies used

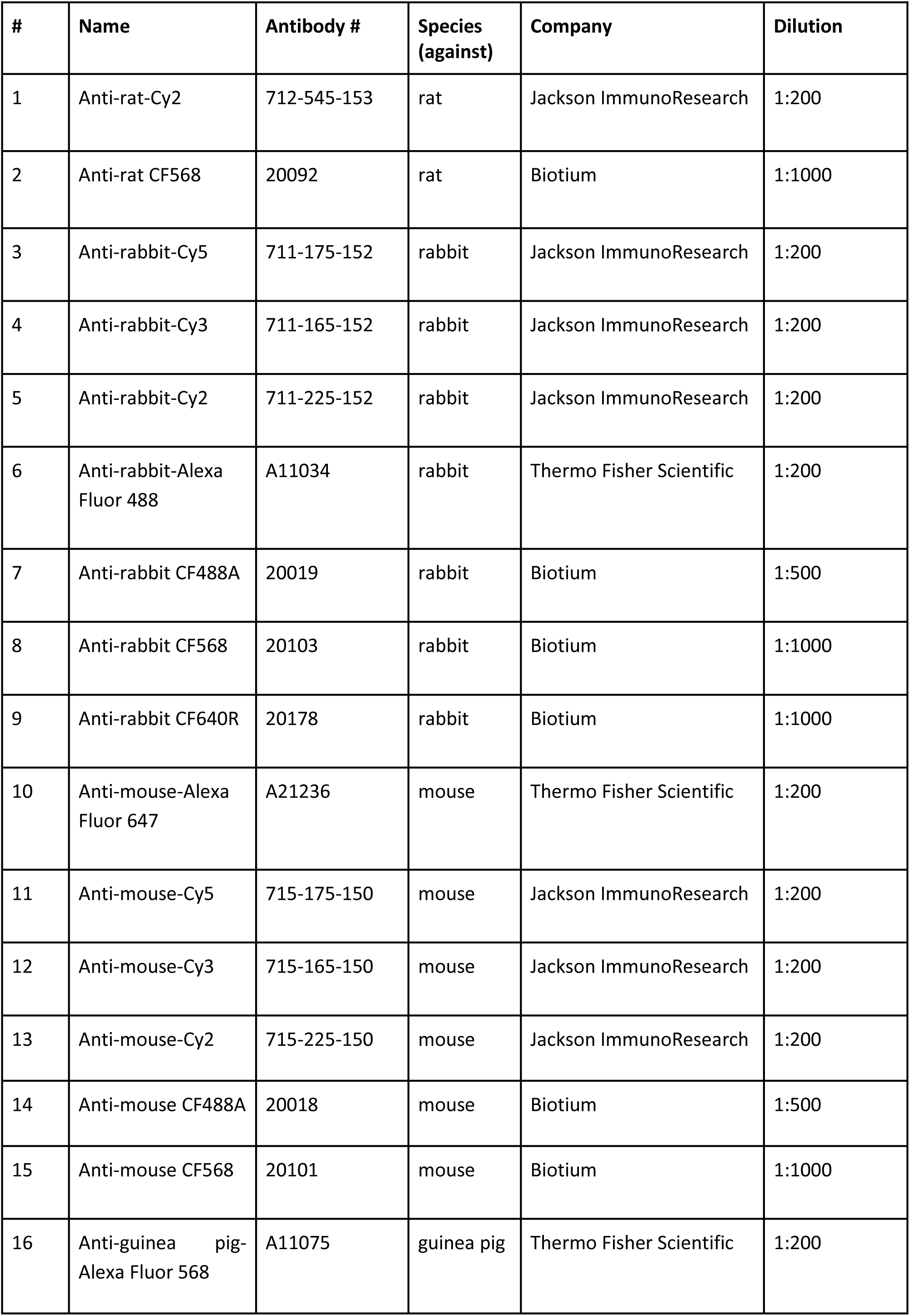
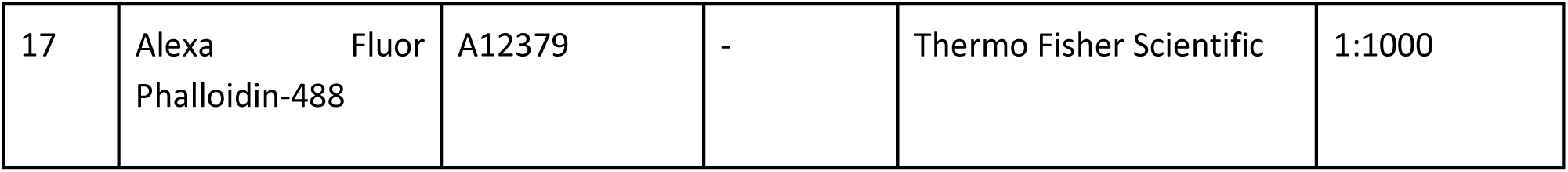

| # | Name | Antibody # | Species (against) | Company | Dilution |
| --- | --- | --- | --- | --- | --- |
| 1 | Anti-rat-Cy2 | 712-545-153 | rat | Jackson ImmunoResearch | 1:200 |
| 2 | Anti-rat CF568 | 20092 | rat | Biotium | 1:1000 |
| 3 | Anti-rabbit-Cy5 | 711-175-152 | rabbit | Jackson ImmunoResearch | 1:200 |
| 4 | Anti-rabbit-Cy3 | 711-165-152 | rabbit | Jackson ImmunoResearch | 1:200 |
| 5 | Anti-rabbit-Cy2 | 711-225-152 | rabbit | Jackson ImmunoResearch | 1:200 |
| 6 | Anti-rabbit-Alexa Fluor 488 | A11034 | rabbit | Thermo Fisher Scientific | 1:200 |
| 7 | Anti-rabbit CF488A | 20019 | rabbit | Biotium | 1:500 |
| 8 | Anti-rabbit CF568 | 20103 | rabbit | Biotium | 1:1000 |
| 9 | Anti-rabbit CF640R | 20178 | rabbit | Biotium | 1:1000 |
| 10 | Anti-mouse-Alexa Fluor 647 | A21236 | mouse | Thermo Fisher Scientific | 1:200 |
| 11 | Anti-mouse-Cy5 | 715-175-150 | mouse | Jackson ImmunoResearch | 1:200 |
| 12 | Anti-mouse-Cy3 | 715-165-150 | mouse | Jackson ImmunoResearch | 1:200 |
| 13 | Anti-mouse-Cy2 | 715-225-150 | mouse | Jackson ImmunoResearch | 1:200 |
| 14 | Anti-mouse CF488A | 20018 | mouse | Biotium | 1:500 |
| 15 | Anti-mouse CF568 | 20101 | mouse | Biotium | 1:1000 |
| 16 | Anti-guinea pig-Alexa Fluor 568 | A11075 | guinea pig | Thermo Fisher Scientific | 1:200 |
| 17 | Alexa Fluor<br>Phalloidin-488 | A12379 | - | Thermo Fisher Scientific | 1:1000 |

### Primary antibody labeling kit

| Kit name | Provider | Catalog number |
| --- | --- | --- |
| CF640R antibody labeling kit | Biotium | 92278 |

## Experimental methods

### Generation of polyclonal NGN2 iNDI lines

Prior to transfection, 0.4-0.5 million iPSCs were seeded into a well of a 6-well plate, allowing for 12-24 hours for cell attachment. PG-TO-hNGN2 plasmid (Gift from Michael Ward, Addgene plasmid #172115) was transfected with TransfeX Transfection Reagent (ATCC® ACS-4005™) along with piggyBac™ transposase vector. Plasmid DNA and Opti-MEM I Reduced-Serum Medium (Gibco 31985-047) were equilibrated to room temperature and were gently mixed to avoid foaming or damaging the reagents. A total of 2μg of DNA was introduced per well, with a donor-to-transposase ratio of 2:1. Next, 100ul of Opti-MEM I Reduced-Serum Medium was dispensed into a sterile microcentrifuge tube, along with the donor and transposase plasmids. Then, 3ul of TransfeX Transfection Reagent ACS-4005 was incorporated into the DNA mixture and was mixed thoroughly using gentle pipetting. A mini centrifuge was used briefly to consolidate the tube’s contents. The resultant lipo:DNA complexes were allowed to incubate at room temperature for 15-30 minutes, promoting the formation of transfection-ready complexes. The complexes were carefully added drop-wise to the iPSCs, ensuring diverse coverage within the well for optimal transfection efficiency. The cells were then incubated at 37°C with 5% CO2 for 24 hours. At 24 hours post-transfection, a complete media change was made and 0.4 ug/ml of puromycin (InvivoGen ant-pr-1) was added to the media. Media supplemented with 0.4 ug/ml of puromycin was replaced every day for 2-3 days until only BFP-expressing cells remained in the culture.

### Generation of inducible TDP-43**^Δ^**^NLS^ and TDP-43^wt^ iPS cell lines

We generated iPS cell lines based on iW11-NGN2 (gift from Michael Ward ^170^) featuring a doxycycline-inducible APEX construct ^171^ fused to either wild-type (TDP-43^wt^) or NLS-mutated TDP-43 (TDP-43^ΔNLS^) (Addgene plasmid #229081), and nuclear export signal (NES) (Addgene plasmid #229082) as cytoplasmic control. A V5 tag was included to enable visualization by microscopy. After cloning these constructs into the PG-TO-hNGN2 plasmid in the place of NGN2, the cassette was integrated into the genome via the piggyBac transposon system described above. After introducing the plasmid, we carefully selected clones and cultivated them according to established protocols. The differentiation process followed the described methodology, with the induction of cytoplasmic TDP-43^ΔNLS^ achieved by adding doxycycline (2 ug/ml) for 24 hours.

### Human iPSC culture (NGN2 lines)

iPSCs were grown at 37°C with 5% CO2 on tissue culture dishes coated with Geltrex (Gibco, A1413302) for at least 1 hour at 37°C. Cell culture media was exchanged daily with mTeSR1 medium (STEMCELL Technologies, 85850). For passaging, iPSCs were dissociated with accutase (ThermoFisher Scientific, A1110501) and seeded into mTeSR1 supplemented with ROCK inhibitor Y-27632 (Tocris Bioscience, 1254). Cells were frozen in mTeSR1 supplemented with 30% KnockOut Serum Replacement (ThermoFisher Scientific, 1082802) and 10% DMSO

### Differentiation of iPSCs to day 8 neurons (NGN2 lines)

Differentiation was performed as in ^170^ with minor adjustments. Briefly, after thawing, cells were transferred to tubes with DPBS, centrifuged, and resuspended in culture medium before seeding onto one Geltrex (Gibco, A1413302) coated well of a 6-well plate. Media was supplemented with 10 μM ROCK inhibitor (Y-27632, Tocris Bioscience 1254) for cell survival. During expansion, cells were cultured with daily media replacements of mTeSR1 medium (STEMCELL Technologies, 85850). Upon reaching 70%-80% confluency, cells were transferred to larger plates in preparation for induction. On the day of differentiation induction, cells were seeded with ROCK inhibitor (Tocris Bioscience, 1254, 10 μM) supplemented to induction medium comprising DMEM/F12, HEPES (Gibco, 11330032), N2 supplement (Gibco, 17502048), NEAA (Gibco, 11140050), L-glutamine (Gibco, 25030081), and freshly prepared doxycycline solution (2mg/ml stock, Sigma D9891). Based on plate size, either 1.5 million or 2.5 million cells were seeded, targeting 5 million and 15 million neurons, respectively. On induction days 1-3 induction media was replaced daily. For maturation starting from day 3 onwards, overnight PDL-coated (sigma, P7405, 0.04mg/ml) or Poly-L-Ornithine (PLO)-coated (Sigma Aldrich, P4957) 96-well plates (Brooks, MGB096-1-2-LG-L or Revvity, 6055302) were prepared, and approximately 60,000 day 3 neurons were seeded per well in maturation medium containing BrainPhys (STEMCELL Technologies, 05790), B27 supplement (Gibco, 17504044), BDNF (PeproTech, 450-02), NT3 (PeproTech, 450-03), and Laminin (Gibco, 23017-015). On the first maturation day, 40mM BrdU (Sigma, B9285) was added to the maturation medium. Neurons matured with half-medium replacements every 3 days. For staining mitochondria, cells were incubated with mitotracker (M22426, 1:20,000 dilution) for 30 min prior to fixation. For stress experiments, cells were incubated with sodium arsenite (500 μM, 30 minutes, Sigma Aldrich, S7400) or D-sorbitol (sorbitol, 0.4 M, 2 hr, Sigma Aldrich, S7547) prior to fixation. For the experiments of wild-type (KOLF) line without treatment or under oxidative stress, the cells were seeded in PLO-coated 384-well plates (Revvity, 6057300)

### Differentiation of iPSCs into day 60 spinal motor neurons

Sporadic ALS (sALS), *C9orf72*, and non-neurological control human subject iPSC lines were acquired from the Answer ALS repository at Cedars Sinai ^172^ (see Materials list). All iPSCs were maintained in 6 well plates coated with growth factor reduced Matrigel (Corning) and mTeSR Plus Media (StemCell Technologies). iPSC lines were passaged at a ratio of 1:10 with ReLeSR (StemCell Technologies). As shown in Supplemental Figure 15, spinal motor neuron cultures were generated by differentiating iPSCs with a modified version of the direct induced motor neuron (diMNs) protocol that has been recently described ^105,133^. Briefly, iPSCs were cultured and expanded for 3 weeks to facilitate acclimation and stabilization post-thaw. On day 0, iPSCs were dissociated with ReLeSR and plated in 6 well plates (coated with growth factor reduced Matrigel) at a density of 500,000 cells/well in stage 1 media (47.5% IMDM (Gibco), 47.5% F12 (Gibco), 1% NEAA (Gibco), 1% Pen/Strep (Gibco), 2% B27 (Gibco), 1% N2 (Gibco), 0.2 mM LDN193189 (Stemgent), 10 mM SB431542 (StemCell Technologies), and 3 mM CHIR99021 (Sigma Aldrich)) supplemented with 20 mM ROCKi (Y-27632, Millipore). The next day, media was exchanged and replaced with stage 1 media without ROCKi. A full volume of stage 1 media was subsequently exchanged on days 3 and 5 of differentiation. On day 6 of differentiation, cells were dissociated with accutase and plated in T25 flasks (coated with growth factor reduced Matrigel) at a density of 2×10^6^ cells/flask in stage 2 media (47.5% IMDM, 47.5% F12, 1% NEAA, 1% Pen/Strep, 2% B27, 1% N2, 0.2 mM LDN193189, 10 mM SB431542, 3 mM CHIR99021, 0.1 mM all-trans RA (Sigma Aldrich), and 1 mM SAG (Cayman Chemicals)) supplemented with 20 mM ROCKi. The next day, media was exchanged and replaced with stage 2 media without ROCKi. A full volume of stage 2 media was subsequently exchanged on days 9 and 11 of differentiation. On day 12 of differentiation, cells were dissociated with Trypsin and plated in 6 well plates (coated with growth factor reduced Matrigel) at a density of 5×10^6^ cells/well in stage 3 media (47.5% IMDM, 47.5% F12, 1% NEAA, 1% Pen/Strep, 2% B27, 1% N2, 0.1 mM Compound E (Millipore), 2.5 mM DAPT (Sigma Aldrich), 0.1 mM db-cAMP (Millipore), 0.5 mM all-trans RA, 0.1 mM SAG, 200 ng/mL Ascorbic Acid (Sigma Aldrich), 10 ng/mL BDNF (PeproTech), and 10 ng/mL GDNF (PeproTech) supplemented with 20 mM ROCKi. The next day, media was exchanged and replaced with stage 3 media without ROCKi. A full volume of stage 3 media was exchanged on day 16 of differentiation. On day 18 of differentiation, iPSNs were dissociated with accutase and plated in 6 well plates (coated with growth factor reduced Matrigel) at a density of 5×10^6^ cells/well in stage 3 media supplemented with 20 mM ROCKi. The next day, media was exchanged and replaced with stage 3 media supplemented with 20 mM AraC. The next day, media was exchanged and replaced with stage 3 media without AraC. A full volume of stage 3 media was subsequently exchanged every 3-4 days until day 32 of differentiation. Beginning on day 32 of differentiation, a half volume of stage 3 media was manually exchanged with a P1000 pipette every 3-4 days until day 53 of differentiation. On day 53 of differentiation, iPSNs were dissociated with accutase and plated in glass bottom 24 well plates (coated with growth factor reduced Matrigel) at a density of 350,000 cells per well in stage 3 media supplemented with 20 mM ROCKi. The next day, media was exchanged and replaced with stage 3 media without ROCKi. A half volume of stage 3 media was subsequently exchanged on day 57 of differentiation. All iPSC lines used in this study were differentiated in a single experimental repeat. All iPSCs and iPSNs were maintained at 37°C with 5% CO_2_ and routinely tested negative for mycoplasma.

### Immunofluorescence

Cells in 96-well or 384-well plates were treated with 4% PFA in 1X PBS++ for a fixed duration of 15 minutes. Post-fixation, cells were rinsed with PBS and were then exposed to 0.1% TritonX in PBS for a permeabilization step for 10-15 minutes. Blocking was achieved by incubating the cells with CasBlock (Thermo Fisher, 008120) reagent for 10 minutes at RT. Primary antibodies (listed in Materials) in CasBlock were prepared, and cells were submerged in 35-70 µl of this mix overnight at 4°C. In the case of the oxidative stress experiment, antibodies were diluted in permeabilization/blocking buffer (DPBS supplemented with 0.2% Saponin + 1.5% BSA) and incubated overnight at 4°C. The following day, cells were washed twice in 100 µl PBS for 5 minutes. Secondary antibodies (listed in Materials), diluted in PBS, were added to the cells, ensuring a 35-70 µl coverage for an hour while minimizing light exposure. Hoechst, diluted in PBS at a 1:8000 ratio, served as a DNA stain, and cells were treated with 75-100 µl of this mix for 10-30 minutes at room temperature. Before proceeding with imaging, the Hoechst solution was replaced with PBS. It was crucial to ensure the cells remained submerged in PBS, particularly during extended imaging sequences to prevent desiccation.

For day 60 motor neurons, iPSNs were fixed and immunostained on day 60, as previously described ^173,174^. Briefly, iPSNs were fixed in 4% PFA for 15 mins and washed 3X 5 mins with 1X PBS prior to permeabilization with 0.1% Triton X-100 in PBS for 15 mins. iPSNs were then incubated in 10% goat serum/1X PBS blocking solution for 30 mins. Primary antibodies were diluted in blocking solution and iPSNs were incubated with the resulting primary antibody solution for 2 hours at room temperature. iPSNs were then washed 3X 10 mins with 1X PBS and incubated with secondary antibody solution (1:1000 Goat Anti-Rabbit Alexa Fluor 488, 1:1000 Goat Anti-Guinea Pig Alexa Fluor 568, 1:1000 Goat Anti-Mouse Alexa Fluor 647; Thermo Fisher) diluted in blocking solution for 1 hour at room temperature. iPSNs were washed 3X 10 mins with 1X PBS, incubated with 1:1000 Hoechst/1X PBS solution for 5 mins at room temperature, and washed 2X 5 mins with 1X PBS prior to mounting with Prolong Gold and sealing of wells with a 15 mm glass coverslip. Agitation was not used at any step in the immunostaining protocol.

### Definitions and experimental design of imaging studies

**Experimental repeat** - an experiment with an independent differentiation, or cells derived from different human subjects, according to the MDAR (Materials Design Analysis Reporting) framework guidelines (https://osf.io/xfpn4)^134^.

**Technical repeat** - multiple wells within the same experimental repeat.

For our AI-based imaging studies, we generated several datasets of iPSC-derived neurons under different conditions (**Table 2**). For each dataset, we created two or more experimental repeats that arose from separate differentiations. Each of those experimental repeats encompasses two or more technical repeats, with cells from the same differentiation plated into a different well. An experimental repeat was imaged uninterruptedly. Human subject lines defined as experimental repeats, were generated in 2 separate experimental repeats (differentiations).

**Table 2:**
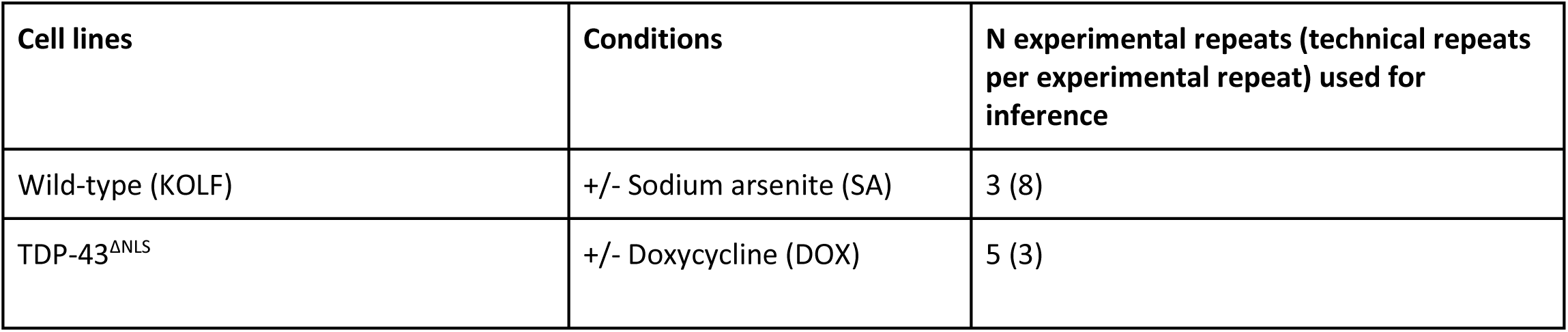
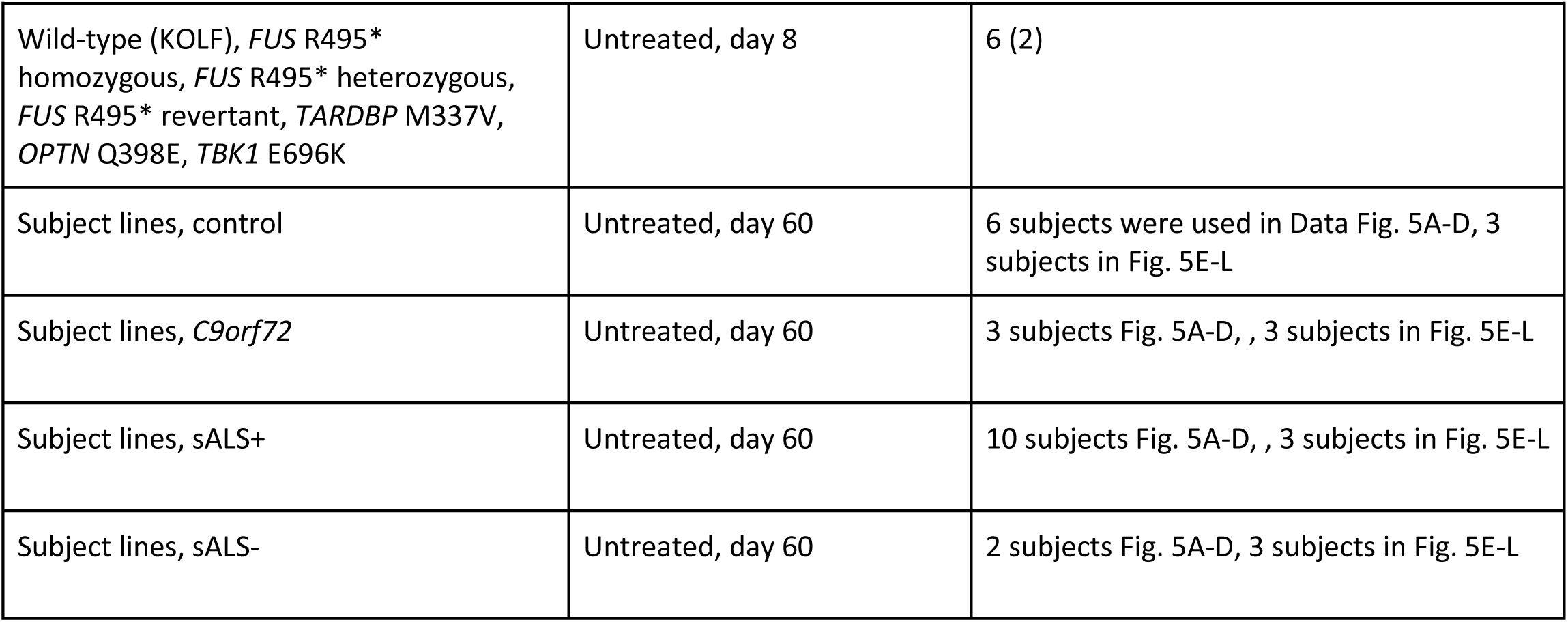
Detailed experimental datasets of iPSC-derived neurons for organellomics studies. The table below depicts the overview of experimental setups generated across this study.

| Cell lines | Conditions | N experimental repeats (technical repeats per experimental repeat) used for inference |
| --- | --- | --- |
| Wild-type (KOLF) | +/- Sodium arsenite (SA) | 3 (8) |
| TDP-43<sup>ΔNLS</sup> | +/- Doxycycline (DOX) | 5 (3) |
| Wild-type (KOLF), *FUS* R495* homozygous, *FUS* R495* heterozygous, *FUS* R495* revertant, *TARDBP* M337V, *OPTN* Q398E, *TBK1* E696K | Untreated, day 8 | 6 (2) |
| Subject lines, control | Untreated, day 60 | 6 subjects were used in Data Fig. 5A-D, 3 subjects in Fig. 5E-L |
| Subject lines, *C9orf72* | Untreated, day 60 | 3 subjects Fig. 5A-D, , 3 subjects in Fig. 5E-L |
| Subject lines, sALS+ | Untreated, day 60 | 10 subjects Fig. 5A-D, , 3 subjects in Fig. 5E-L |
| Subject lines, sALS- | Untreated, day 60 | 2 subjects Fig. 5A-D, 3 subjects in Fig. 5E-L |

### Microscopy

The iPSC-derived neurons (iNDI lines, day 8) that were used for training, were assessed using the Visitron Systems laser and laser control, complemented by the spinning disk confocal scanner unit from Yokogawa. The Olympus IX83 served as the primary microscope in this configuration, images were acquired at X60 using oil immersion. We used a filter wheel system featuring EF460/50m, E525/50m, FF01-609/54, and ET700/75m filters, along with the DM405/488/561/640 dichroic mirror. To optimize image acquisition, we employed specific parameters, including binning at 2 for pixel averaging and heightened sensitivity, an exposure time of 300ms, and dynamic adjustments to laser power (range: 0.1-100%) for optimal imaging of distinct fluorophores. Our high-throughput imaging strategy encompassed the capture of 10×10 images per well within a 96-well plate, resulting in a total of 100 images per well, without overlap between adjacent images. This systematic approach was applied to all cell lines in a consecutive run. Each image has 2-4 channels of a panel of 2-4 antibodies. The entire process was controlled through VisiView Software, ensuring seamless microscope control and efficient image acquisition.

The inference dataset of 8-day-old iPSC-derived neurons with or without oxidative stress was collected through automated imaging in the Nikon Bio-Pipeline equipped with a Ti2 Spinning Disk confocal microscope using a X60 water immersion lens. All other inference datasets of 8-day-old iPSC-derived neurons (iNDI collection; ALS lines or wild-type with or without osmotic stress, and the TDP-43^ΔNLS^ line) were assessed using the Opera Phenix Plus HCS system REVVITY, HH14001000). Images were acquired at X63 using automatic water immersion using two cameras, four channels (splitted between the cameras). Here, we also employed specific parameters, including binning at 2 for pixel averaging and heightened sensitivity. However, the laser power (Intensity) and time exposure (ms) parameters were set for each channel and marker specifically. All images were taken with 4-6 Z-stacks and max projected together with help from REVVITY’s expert.

Day 60 motor neurons were imaged with a X63 objective with X1 digital zoom on a Zeiss LSM980 confocal in 24 well plates. Identical imaging parameters (e.g. laser power, gain) were used to acquire images. Ten images per well, 4 antibodies (see above), without overlap between adjacent images were taken.

### Western blot

For western blot analysis of TDP-43 protein levels in wild-type and TDP-43^ΔNLS^-V5-APEX-expressing neurons, iPSC-derived neurons were grown to day 8 and construct expression was induced as described above. Protein content was harvested by scraping cells directly in RIPA lysis buffer supplemented with Complete Protease Inhibitor Cocktail (Roche, 4693116001) and PhosphoSTOP (Roche, 4906837001) and centrifugation of lysates at 15,000 × g for 10 min at 4°C. Protein concentration was then quantified with Bio-Rad Protein Assay Dye Reagent (Bio-Rad, 500-0006). 30 ug of protein per well was loaded and gel electrophoresis was performed in 10% SDS-PAGE at 100 V for ∼70 min. Next, proteins were transferred to nitrocellulose membranes (Whatman; 10401383) at 250 mA for 70 min followed by membrane blocking with 3% bovine albumin fraction V (MPBio; 160069) in PBS supplemented with 0.05% Tween-20 (PBST) for 1 h at R.T. Membranes were then incubated at 4°C overnight on a shaker with rabbit anti-TDP-43 antibody (1:5000, Proteintech, 10782-2-AP) and mouse anti-Tubulin (1:4000; Sigma-Aldrich, T9026) in antibody solution (5% albumin, 0.02% sodium azide and five drops of phenol red in 0.05% PBST). The following day, membranes were washed with 0.05% PBST three times for 5 min before proceeding to incubation with horseradish peroxidase-conjugated secondary antibodies (anti-rabbit, Jackson ImmunoResearch, 711-035-152; anti-mouse, Jackson ImmunoResearch, 715-035-150). Membranes were then washed with 0.05% PBST three times for 5 min and imaged using the Westar NOVA 2.0 Chemiluminescence kit (Cyanagen, XLS071) and ImageQuant LAS 4000 (GE Healthcare Life Sciences). Band intensities were quantified using ImageJ software and averaged across three experimental repeats for plotting.

### Fluorescence recovery after photobleaching (FRAP)

U2OS cells overexpressing DCP1A-YFP (yellow fluorescent protein, a gift from Dr. Nancy Kadersha and Prof. Paul Tylor) were maintained as described above (section of: U2OS cells). For FRAP of P-bodies under uninduced or induced cytoplasmic TDP-43 or Nuclear export signal (NES) expression, ∼250K U2OS cells were seeded in individual 6-wells with coverslips. Next, the cells were co-transfected with two different plasmids: piggyBac transposase plasmid and PG-TO-hNGN2 that carries doxycycline-inducible NES-V5-APEX (as negative control) or TDP-43^ΔNLS^-V5-APEX (see above, expresses BFP; Blue Fluorescent Protein) in the place of NGN2, using Transfex reagent (ATCC, ACS-4005). The inducible expression constructs were integrated into the genome via the piggyBac transposon system. Twenty-four hours post-transfection, the cells were incubated overnight with doxycycline (DOX; 1ug/ml) for NES or cytoplasmic TDP-43 induction or only with media (as control; -DOX; uninduced) and with sodium arsenite (250 uM, Sigma, S7400) 60 min before the FRAP measurement. After 60 min, FRAP image sequences were obtained during 30 min on a Zeiss LSM 900 inverted scanning confocal microscope equipped with a heated chamber and a Plan-Apochromat 40× water objective (Carl Zeiss, Jena, Germany). First, we looked for cells expressing BFP to verify the successful uptake of the inducible constructs (NES- or TDP-43^ΔNLS^-V5-APEX. Those cells were scanned using a 488 laser for the detection of YFP-labeled P-bodies. For analysis of the fluorescence recovery, an internal area within a few P-bodies in a field of view was bleached and the fluorescence recovery was measured every ∼1 sec for 2 min. A single P-body and background area without bleaching were also measured per field of view (a site in a well). In each experiment, for each condition (uninduced or induced), 3-5 sites were analyzed in each of four 6-wells (technical repeats). In each site, between two to nine P-bodies were measured and the intensities for each time point were averaged. FRAP data were normalized and calculated by subtracting the background signal and then normalized to the P-body that was not bleached as reference. Measured P-bodies that were out-of-focus or exhibiting technical problems (e.g. unsuccessful bleaching) were excluded. The average of the normalized data for an experiment was synchronized by the average time point value. The final data points that are shown in the figures represent average of the 4 technical repeats as described above. For each experimental repeat, the statistical testing was two-way ANOVA with repeated measures and Geisser-Greenhouse correction, in Prism (version 10).

### APEX proximity labeling

Monoclonal iPSC lines carrying TDP-43^ΔNLS^ and NES APEX constructs were differentiated into neurons as described previously. After 5 days of maturation, construct expression was induced for 24 and 4 hours respectively to achieve comparable levels of expression. APEX activity was induced by supplementing biotin-phenol (BP, 500 μM, Iris Biotech GmbH, LS-3500) for 90 minutes and H2O2 (1 mM, J.T.Baker 7722-84-1) for 1 min. Labeling activity was halted with quenching solution (QS: sodium azide (10mM, Mallinckrodt, 1953-57), sodium ascorbate (10mM, Sigma-Aldrich, A7631) and Trolox (5mM, Sigma-Aldrich, 238813) in PBS. Then, cells were scraped directly in RIPA lysis buffer supplemented with Complete Protease Inhibitor Cocktail (Roche, 4693116001) and PhosphoSTOP (Roche, 4906837001). Lysates were centrifuged at 15,000 × g for 10 min at 4°C after which protein concentration was quantified with Bio-Rad Protein Assay Dye Reagent (Bio-Rad, 500-0006). Streptavidin-coated magnetic beads (Pierce Streptavidin Magnetic Beads, Thermo-Fisher, 88816) were incubated at a ratio of 100 μl beads per 500 μg of sample with rotation overnight at 4°C. The next day, bead samples were washed with a series of buffers, 1 ml per wash, for 3 minutes at 1100 rpm at room temp to remove non-specific binders: twice with RIPA lysis buffer, once with 1 M KCl (MERCK, 104936), once with 0.1 M Na2CO3 (Sigma-Aldrich, S7795), and once with 2 M Urea solution containing 10 mM Tris-HCl, pH 8.0 (Invitrogen, 15568–025).

### Sample preparation for mass spectrometry

All chemicals are from Sigma-Aldrich Aldrich, unless stated otherwise. Samples were subjected to on-bead tryptic digestion followed by a desalting step. Briefly, washed beads after pull-down were suspended in 8M urea, 0.1M Tris HCl buffer, pH 7.6, and incubated at room temp for 30 min. Proteins were then reduced with 100 mM DTT (5mM final conc) at room temp for 1 hr and then alkylated using iodoacetamide (10 mM final concentration) for 45 min in the dark at room temp. Urea concentration was diluted 5-fold by addition of 50mM ammonium bicarbonate. 250 ng trypsin was then added and samples were incubated at 37°C overnight, followed by a second trypsin digestion (250 ng) for 4 hours at 37°C. Digested peptides were then collected by centrifugation of the beads and transferring the supernatant into a clean tube. Peptides were acidified with trifluoroacetic acid, desalted using HBL Oasis (Waters 094225), speed vac to dryness and stored in -80°C until analysis.

### Liquid chromatography and mass spectrometry

Digested proteins were loaded and analyzed using split-less nano-Ultra Performance Liquid Chromatography (10 kpsi nanoAcquity; Waters, Milford, MA, USA). The mobile phase was: A) H2O + 0.1% formic acid and B) acetonitrile + 0.1% formic acid. Desalting of the samples was performed online using a Symmetry C18 reversed-phase trapping column (180 µm internal diameter, 20 mm length, 5 µm particle size; Waters). The peptides were then separated using a T3 HSS nano-column (75 µm internal diameter, 250 mm length, 1.8 µm particle size; Waters) at 0.35 µL/min. Peptides were eluted from the column into the mass spectrometer using the following gradient: 4% to 26%B in 155 min, 26% to 90%B in 5 min, maintained at 90%B for 5 min and then back to initial conditions. The nanoUPLC was coupled online through a nanoESI emitter (10 μm tip; New Objective; Woburn, MA, USA) to a quadrupole orbitrap mass spectrometer (Q Exactive Plus, Thermo Scientific) using a FlexIon nanospray apparatus (Proxeon).

For targeted MS, samples were analyzed by targeted analysis while monitoring relevant peptides. The targeted analysis data were analyzed using the Skyline algorithm and manually curated for confident identifications and accurate quantification. The data were also searched against the human protein database using the Proteome Discoverer software to increase confidence in the assignment of the relevant peaks.

For unbiased, proteome-wide MS, data was acquired in data dependent acquisition (DDA) mode, using a Top10 method. MS1 resolution was set to 70,000 (at 200m/z) and maximum injection time was set to 60 msec. MS2 resolution was set to 15,000 (at 200m/z) and maximum injection time of 60 msec.

### Raw proteomic data analysis of unbiased MS

Raw data was analyzed using MaxQuant (v1.6.6.0) ^175^. The raw data was searched against the human protein database downloaded from UniprotKB appended with 125 common laboratory contaminant proteins. Enzyme specificity was set to trypsin and up to two missed cleavages were allowed. Fixed modification was set to carbamidomethylation of cysteines and variable modifications were set to oxidation of methionines, asparagine and glutamine deamidation, and protein N-terminal acetylation. Peptide precursor ions were searched with a maximum mass deviation of 4.5 ppm and fragment ions with a maximum mass deviation of 20 ppm. Peptide, protein and site identifications were filtered at an FDR of 1% using the decoy database strategy. The minimal peptide length was 7 amino-acids and the minimum Andromeda score for modified peptides was 40. Peptide identifications were propagated across samples using the match-between-runs option checked. Searches were performed with the label-free quantification option selected.

### Proteomics statistical analysis

MaxQuant output table ProteinGroups was imported to Perseus ^176^ and analyzed as follows. Reverse proteins, proteins identified only based on a modified peptide and contaminants were excluded before log2-transforming the data. Rows were filtered to keep only those with at least 6 valid values, and missing values were replaced by low values from a normal distribution. Two-sample t-test comparing samples with biotin to samples without biotin was performed to remove non-specific binders (FDR < 0.05, log2FC > 0). Samples were separated based on stress condition before filtering to remain only those proteins with a minimum of 2 valid values in at least one experimental group (TDP-43^ΔNLS^ or NES). Missing values were imputed again using a normal distribution. Two-sample t-test using log2-transformed LFQ values was performed to gain proteins enriched in TDP-43^ΔNLS^.

For targeted MS, peptide intensities were normalized against the Total Ion Current Area per peptide for each protein. These values were averaged to get a single value per protein. Then, data from both groups (TDP-43^ΔNLS^ and NES) were normalized to the average of the NES group per protein before comparing between conditions to get fold changes.

### Human post-mortem tissue staining and quantification

Motor cortical tissues from *C9orf72* or sALS cases and age-and sex-matched controls were from the Target ALS post-mortem core or from the Sheffield Brain bank under the following ethical approvals: SBTB REC number 08/MRE00/103+5, SBTB application no: 23/007. Cases were included in triplicate in a tissue microarray (TMA) format (TMA available through the Target ALS Post-mortem core). For P-body immunohistochemistry analysis, sections were cut at 4 µm thickness onto superfrost charged slides and deparaffinized. Sections then underwent heat mediated antigen retrieval in citric acid (pH=6) for 15 minutes in a pressure cooker. Following this antibodies were incubated for 90 minutes at the following concentrations: DCP1A 1:100 (Abcam ab47811), LSM14A 1:100 (Proteintech 18336-1-AP) and AGO2 1:200 (Abcam ab32381) for 1 hr at room temperature. Sections were stained using the Novolink Polymer detection system and 3,3’-Diaminobenzidine (DAB) chromogen was used and counterstaining was performed with haematoxylin, according to standard operating procedures. TDP-43 aptamer dual chromogenic stains were performed using our standard operating protocol freely available online ^177^ using the following concentrations: DCP1A 1:100, LSM14A 1:100, secondary anti-biotin Alkaline phosphatase antibody at 1:100 (30 minutes incubation). Digital analysis was performed using ImageJ analyze particle function. Images were batch processed as follows: Image>8-bit. Process>Sharpen. Image>Adjust>Brightness & Contrast>minimum = 53, maximum = 144. Set measurements> area and limit to threshold. Image>Adjust>Threshold>48-103. Analyze>Analyze Particles>Size=5-Infinity, Circularity = 0-1, Show = Bare Outlines, Display results, Add to Manager.

### SLAM-seq and mRNA stability analysis

For metabolic RNA labeling of TDP-43^ΔNLS^-expressing iPSC-derived neurons, the maturation medium was replaced with a medium containing 4sU (200 μM, T4509, Sigma). Since multiple timepoints are recommended, and 3 hours of labeling was shown to be sufficient to reveal changes in RNA half-life ^131^, cells were incubated with 4sU for 0, 2 or 3 hours. After labeling, cells were harvested with Tri-reagent (Sigma) followed by RNA extraction under reducing conditions and treatment with iodoacetamide (A3221, Sigma) as previously described ^130^. RNA-seq libraries were prepared at the Crown Genomics Institute of the Nancy and Stephen Grand Israel National Center for Personalized Medicine, Weizmann Institute of Science. A bulk adaptation of the MARS-Seq protocol ^178,179^ was used to generate RNA-Seq libraries for SLAM-seq. Briefly, 30 ng of input RNA from each sample was barcoded during reverse transcription and pooled. Following Agencourt Ampure XP beads cleanup (Beckman Coulter), the pooled samples underwent second-strand synthesis and were linearly amplified by T7 in vitro transcription. The resulting RNA was fragmented and converted into a sequencing-ready library by tagging the samples with Illumina sequences during ligation, RT, and PCR. Libraries were quantified by Qubit and TapeStation as well as by qPCR for Human GAPDH gene as previously described ^178,179^. Sequencing was done on a Nextseq 500, using Nextseq High Output 75 cycles kit mode, allocating 400M reads in total (Illumina). RNA-seq reads were aligned to the human genome (version GRCh38.99) with STAR (version 2.7.3a) ^180^ using parameters “--outFilterMismatchNmax 20 --outFilterScoreMinOverLread 0.4 --outFilterMatchNminOverLread 0.4 --outSAMattributes nM MD NH --alignEndsType Extend5pOfRead1 --quantMode GeneCounts”. Bam files from STAR were processed with GRAND-SLAM (version 2.0.7b) ^131^ with trimming of 20 nucleotides in the 5’ end due to higher mismatch rates across all nucleotides in this region. The error rate correction linear model was set to AT instead of the default TA, because the latter showed a higher error rate and variability across samples. Unlabeled samples without 4sU were used for the model estimation. The main output table of GRAND-SLAM ({prefix}.tsv) was used as input for analysis with grandR (version 0.2.5) ^132^ for half-life estimations as previously explained ^181^. We based analysis of P-body-enriched RNAs, on data from ^116^ (FDR < 0.05, log2FC > 1). Final statistical testing was done with Prism (version 10).

## Computational methods

### Preprocessing pipeline of microscopic images

Day 8 iPSC-derived neurons (used for model training): Immunofluorescence confocal images were taken using a Spinning disc confocal microscope. Each image (1,024x1,024 pixels, also referred as “site image”) is dual-channel, holding the target marker as the first channel and the nucleus (Hoechst 33342) as the second. First, raw pixels were scaled to a range of 0 to 1 using the rescale_intensity function from skimage python package (V0.21.0), with the in_range parameter sets to 0.5 percentile and 99.9 percentile of the pixels’ intensity distribution and out_range sets to numpy.float32. Corrupted/out-of-focus sites were filtered (see **Methods**). Site image was then cropped to 64 equal-sized tiles (128x128 pixels). Each tile was resized to 100×100 pixels (using the resize function from skimage package V0.19.3, with anti_aliasing=True). Tiles containing at least 80% of a single nucleus and not more than 5 nuclei were considered valid. See also **Supplementary Figure 2**.

Day 8 iPSC-derived neurons in inference (**Figures 1-4**): Immunofluorescence confocal images were taken using an Opera Phenix Plus microscope (Revvity, **Methods**). Site images of 1,080x1,080 pixels were cropped into 1024×1024 pixels by cropping 28 pixels from each edge. The same pipeline as described before was applied here. In addition, pixel intensity of each tile was scaled to a range of 0 to 1 with the rescale_intensity function using the same parameters as applied to the full site. Tiles containing dead cells or lacking detectable signal in any channel were excluded. Since images of neurons are highly sparse, all tiles were subjected to QC process, where we kept only valid tiles. See also **Supplementary Figure 2**.

Immunofluorescent confocal images of sALS patient-derived motor neurons (day 60) (1,024x1,024 pixels) were taken using Zeiss LSM 980 confocal microscope (Zeiss), with a larger zoom magnitude (X63) than the rest of the data (X60). Hence, each image was first cropped to 1022×1022 by removing one pixel from each side, then cropped into 49 equal-sized tiles (146x146 pixels), which were subsequently resized to 100×100 pixels as described before. Neither intensity rescaling nor filtering of unfocused images were needed. Tiles containing dead cells or lacking detectable signal in at least one channel were excluded.

A detailed summary of the datasets is provided in the **Supplementary Materials**.

### Quality control

Site images (both the target organelle and the nucleus channels) that contained debris or were out of focus were filtered using Brenner’s Gradient score ^182^ (applied on both horizontal and vertical dimensions), an approach for focus evaluation. Site images that passed filtering were then uniformly cropped to tiles of size 100×100. A tile considered valid if it contained 0.8 of the nuclei, and if both its maximum pixel intensity and variance exceeded predefined thresholds as follows.

To determine whether a tile contained at least 80% of a nucleus, we performed nuclei detection using Cellpose^183^ (V2.0.3, parameters: model type=’nuclei’, diameter=60) on the nucleus channel convolved with [[-1,-1,-1], [-1,25,-1], [-1,-1,-1]] for edges enhancement. For each detected nucleus, a polygon was computed using the Python package Shapely (version 2.0.1). We then calculated the ratio between the area of the polygon within the tile and the total polygon area outside the tile. Tiles with a ratio greater than 0.8 were considered to contain a sufficiently complete nucleus. Nuclei intersecting the site boundaries, where the full nuclear size could not be determined and the 80% threshold could therefore not be tested, were excluded from the nuclei count. Intersection with the site boundaries was detected by computing a polygon for the entire site boundary, minus a padding of 1 unit on each side. If the nucleus polygon was not completely contained within the site’s polygon, the nucleus was disqualified (**Supplementary Figure 2**). The neuron count in our data, therefore, reflects the number of nuclei for which at least 80% of the area was captured within valid tile.

To detect tiles containing dead cells, we leveraged the observation that dead cells typically exhibit elevated nuclear pixel intensities and smaller nuclei than living cells. Accordingly, we evaluated the median intensity of each nucleus within the rescaled intensity tile. Instance segmentation of nuclei was performed in two steps: first, pixels belonging to nuclei were identified by applying Otsu thresholding (using the threshold_otsu function from scikit-image), and second, individual nuclei were delineated by detecting connected components (using the label function from scipy.ndimage, v1.12.0). This procedure yielded a mask for each nucleus, from which we computed the median pixel intensity. Nuclei with median intensity above 0.95 (on a scale where 1 represents the maximum possible intensity) were classified as dead cells, except for the sALS patient-derived motor neuron (Day 60) full dataset, where a lower threshold of 0.8 was used. For the iPSC-derived neuron datasets, an additional size-based criterion was applied to improve dead-cell detection, since dead cells typically display smaller nuclei than living cells. For nuclei not touching the tile frame, the area was measured as the number of pixels within the nucleus mask. Nuclei with an area below 800 pixels were classified as dead.

For detecting tiles lacking a detectable signal, we applied two criteria. First, we tested whether the maximum pixel intensity of the tile (prior to intensity rescaling) was below a predefined threshold (0.2 for both channels). Second, we evaluated whether the variance of the rescaled pixel intensities fell below a predefined threshold (0.0001 for the target channel, and 0.03 for the nucleus). Tiles meeting either condition were excluded.

### Localization encoding ViT-based model

A ViT model with linear head and cross-entropy loss was trained from scratch on the publicly available OpenCell dataset (1,100,253 cropped images ^48^). The data was split into 70%, 15%, 15% for training, validation, and testing data, respectively. Model parameters: embed_dim=192, patch_size=14, in_chans=2, output_dim=1311. The input for the model are dual-channel tiles of 100×100 (a 3D vector sized (2,100,100)). Model training converged after 44 epochs (**Supplementary Figure 3A**). In training, the following augmentations were applied with a probability of 0.5 for each: horizontal flip, vertical flip, and rotation of 90, 180, or 270 degrees. Train parameters: we used the AdamW optimizer with adjustable weight_decay for non bias nor Norm parameters. A cosine scheduler was used for controlling the weight decay (base value=0.04, final value=0.4, number of epochs=300) and the learning rate (base value=0.0008*experimental repeat_size/256, final value= 0.000001, number of epochs=300, number of warmup epochs=5). Experimental repeat comprised of 350 site images, each containing 7 valid tiles on average. Early stopping was applied after ten epochs in which the validation loss showed no improvement. A grad scaler (torch.cuda.amp.GradScaler) was used for calculating the gradients. Unscaled gradients’ norms were clipped (torch.nn.utils.clip_grad_norm_) with max_norm=3.

### Perturbation learning ViT-based model using InfoNCE contrastive loss

For perturbation learning, we extended the pre-trained model as follows. We first initialized a ViT model with weights from the pre-trained model (that was trained for localization encoding). To better capture non-linear relationships in the data, the original linear ViT head was replaced with a multi-layer architecture (*Linear*(*192, 768*) *-> BatchNorm1*(*768*) *-> GELU -> Linear*(*768, 384*) *-> BatchNorm1d*(*384*) *-> GELU -> Linear*(*384, 128*) *-> LayerNorm*(*128*)). During fine-tuning, we partially froze selected layers (see **Methods**, **Supplementary Figure 4**) and the model was trained in a supervised contrastive framework using the infoNCE^4445^ loss (**Figure 1B**, **Supplementary Figure 6**).

The contrastive learning task was implemented as following: for each anchor tile *x* during training (100x100 pixels, two channels), six additional images of the same organelle were randomly sampled: one positive (*x*^+^) and five negatives (*x*^−^). Positives corresponded to image tiles from the same organelle under the same perturbation, whereas negatives were tiles under different perturbations. Notably, sampling is done with replacement, and matches were allowed to originate from different experimental repeats. Pulling anchors and positives together across repeats reduced repeat-specific variations, thereby mitigating overfitting to technical artifacts and encouraging the model to learn repeat-invariant representations. Conversely, pushing anchors away from negatives refined representations towards true perturbation-specific effects.

The InfoNCE loss was calculated as follows:

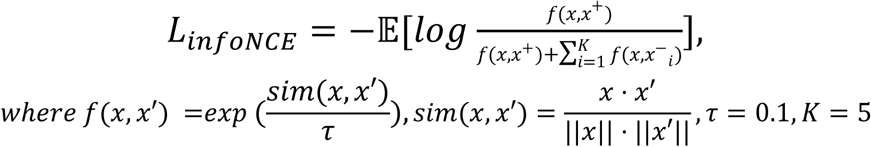

Since sampling was performed with replacement, each tile could also serve as an anchor, positive or negative across different matched triplets, enhancing generalization. This transitive structure of contrastive learning reinforces relationships across samples, reducing overfitting to specific image pairs and improving robustness. By sharing anchors among multiple comparisons, the model learns a consistent representation space that better captures true perturbation effects.

Unlike standard supervised classification, this contrastive formulation models *weak supervision*: perturbation labels indicate potential, but not guaranteed, phenotypic effects. Because not all perturbations yield consistent organellar changes, relying solely on label differences could introduce spurious relationships. Incorporating negatives within the contrastive framework mitigates this risk by emphasizing reproducible, perturbation-specific signals over random or noise-driven variation.

Training followed the same procedure as the pre-trained model, with the exception of a reduced output dimension of 128 and an experimental repeat size of 750. The dataset used for training was composed of 3,213,360 dual-channel images of human iPSC-derived neurons spanning 25 organelles (Nucleus, ER, Nucleolus, TDP43 granules, Paraspeckles, ANXA11 granules, Golgi, Lysosome, FUS granules, Peroxisome, P-bodies, Integrin puncta, Autophagosomes, PML bodies, Presynapse, NEMO granules, Postsynapse, Transport machinery, Coated vesicles, MOM, Mitochondria, PURA granules, Stress granules, Actin Cytoskeleton, FMRP granules) across 9 different cell lines/conditions (WT, WT treated with stress, FUS heterozygous, FUS homozygous, FUS revertant, TBK1, OPTN, TDP43 and SNCA). The data was generated from two independent biological differentiations (referred to as “experimental repeats”), with each differentiation including two to eight technical repeats per condition. It was split into train (70%), validation (15%), and test (15%) sets. The model converged after 19 epochs (**Supplementary Figure 5**). All reported results were obtained on held-out datasets and were not used during model development.

### Partial fine-tuning: the angle-metric approach for freezing layers in perturbation learning

Fine-tuning is the adaptation of a pre-trained model, initially trained on massive amounts of labeled data, to perform well on a different but related problem. This involves selectively updating the model’s parameters to capture the nuances of the new task. ^184^. The “localization encoding model” is extended to perform “perturbation learning” in neurons (**Figure 1B**) using partial fine-tuning (**Supplementary Figure 4**). All weights of the transformer encoder are copied to the new model, except for the head, but not all model layers were re-trained with data of neurons. The training process is done by utilizing a data-drive approach for selecting which layers to freeze and which to fine-tune with our perturbed neurons. Frozen layers are not being updated during the fine-tuning process. For selecting which layers to freeze we first trained a perturbation learning model (see **Methods**) without freezing any layer. Then, we calculated for each layer how much it deviated from the pre-trained model weights using the angle metric ^49^:

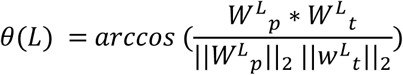

where *W*^*L*^_*p*_ are the weights of layer L in the pre-trained model, and *W*^*L*^_*t*_ are the weights of the same layer L in the model after no-freezing fine-tuning.

Finally, since the tasks are different, yet share basic commonalities, we picked the least deviated layers (50% of the layers) to be the ones we freeze.

### Image encoding

While our training pipeline runs the training consecutively (the localization encoding model and then the perturbation learning model), the inference step is done only with the perturbation learning model. During inference, a 100×100 pixels image (target and nucleus channel) is fed into the model to generate a 192-dimensional numerical representation from the second-to-last layer, normalized with L2 norm.

### Combined effect size via random-effects meta-analysis

To quantify the effect of a perturbation on an organelle topography (relative to a baseline), we define a distance-based “effect size” metric on NOVA-derived embeddings (full latent space). This measure captures the degree to which a perturbation shifts organelle topography away from a baseline state, while accounting for natural variability within the baseline population.

Since we seek for perturbational changes that are consistent across experimental repeats (i.e., biological differentiations or human subjects within the same gene group, etc.), our meta-analytic approach considers each experimental repeat as a “study” with effect size, and performs a meta-analysis with mixed-effects for each organelle–perturbation pair. This results in an estimate of “combined effect size” across experimental repeats, while modeling both the sampling variance within each repeat and the heterogeneity across repeats.

Let *K* be the number of experimental repeats. For each combination of (perturbation × organelle × experimental repeat), we compute the observed effect size and variance for experimental repeat (steps 1-4), that are subsequently aggregated across experimental repeats using mixed-effects meta-analysis to yield a final combined effect estimate (step 5).

Denote,

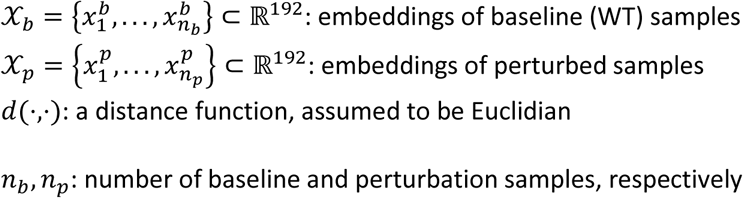

∀*k* ∈ 1, . . , *K* We compute the per-experimental repeat estimates, namely effect size *θ̂*_*k*_ and *Var*(*θ̂*_*k*_) as follows:

**Step 1: Baseline median**

Let 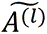 ∈ ℝ^192^ be the median of the baseline group in the latent space.

**Step 2: Within-group dispersion**

Median distance of baseline samples to baseline median:

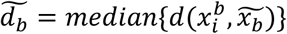

**Step 3: Between-group dispersion**

Median distance of perturbed samples to baseline median:

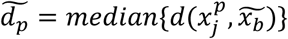

**Step 4: Effect size and bootstrapped sampling variance per experimental repeat**

Effect size *θ̂* is defined as the “relative distance shifts” (how much a perturbation shifts a distribution in embedding space), and computed as the log fold change between the “between-group dispersion” and “within-group dispersion”:

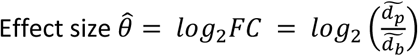

Estimate the empirical variance *Var*(*θ̂*): assess the sampling uncertainty of the effect size across samples by bootstrapping: (i) Resample (with replacement) tiles from different sites from 80% of the perturbation and baseline samples. (ii) Recompute the effect size *θ̂* for each bootstrap resample. (iii) Repeat *B* times, and trim extreme outliers (iv) Estimate variance (*Var*(*θ̂*)) from the bootstrap distribution.

**Step 5: Random-effects meta-analysis**

Given the per-repeat estimates from steps 1-4 above (experimental repeat effect sizes *θ̂*_1_, . . , *θ̂*_*K*_ and *Var*(*θ̂*_1_), . . , *Var*(*θ̂*_*K*_)), the combined (pooled) effect size *μ̂* and heterogeneity measures *τ*^2^ (namely “between-studies variance”), *Q* and *I*^2^, are computed as follows:

Fixed-effects weights:

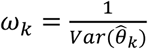

Q statistics (heterogeneity):

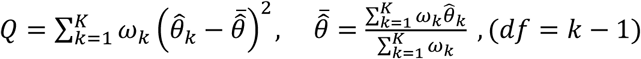

Estimate between-experimental repeat variance *τ*^2^:

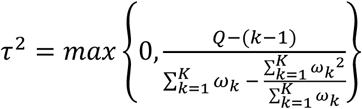

Random-effects weights: each experimental repeat is weighted by the inverse of its total variance

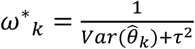

Combined (pooled) effect size:

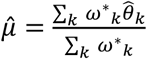

The *I*^2^ statistic, a measure of heterogeneity across repeats:

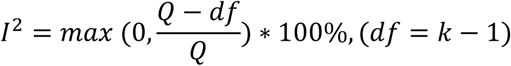

*I*^2^ranges from 0% (no observed heterogeneity) to 100% (all variation due to heterogeneity). Values above 75% are typically interperted as indicating considerable heterogeneity.

Notably, when estimated heterogeneity (*τ*^2^) was zero, the combined effect size and CI coincides with the fixed-effect model.

**Step 6 (optional): confidence interval for** *τ*^**2**^

Confidence intervals were calculated using a profile likelihood approach. Log-likelihood was defined as follow:

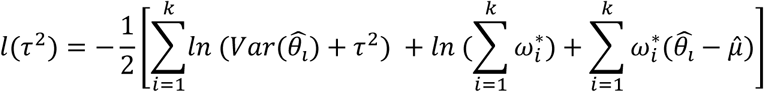

If *τ*^2^ = 0, the lower bound was set to 0. Otherwise the lower bound was obtained by solving for the value of ^*τ̂*2^ between 0 and *τ*^2^ where the 2 ∗ *l*(*τ*^2^) − *l*(^*τ̂*2^) equalled the chi-square cutoff (*χ*^2^). For the upper bound, the search range was iteratively expanded from *τ*^2^until the log-likelihood dropped below the chi-square threshold, after which the precise upper bound was located by root-finding within this interval, also using optimize.brentq.

Effect sizes were tested for significance using z-scores, calculated as the ratio between the combined effect estimate and its standard error. One-tailed p-values were obtained under the null hypothesis of *ratio* <= 0, (*α* = 0.05). To control for multiple testing across organelles, p-values were corrected using the Benjamini–Hochberg procedure (FDR).

Inclusion of an experimental repeat in the meta-analysis required meeting a predefined minimum sample size (see **Supplementary Materials** for the threshold for each dataset).

Combined effect size and its confidence intervals were calculated using the combine_effects function from the statsmodels python package (v0.14.2). Confidence interval for *τ*^2^(profile likelihood approach) were computed using optimize.brentq function from the scipy (v1.12.0) python package.

### Organelle scoring forest plot

For each organelle × perturbation pair, the organellome was ranked based on the combined effect size *μ̂*. The ranking is visualized with a stacked forest plot with multiple organelle × perturbation pairs, presenting as a group both experimental repeat-level effects and the combined (pooled) inference for every organelle. Each repeat is represented by a dot (experimental repeat-wise effect estimate) and a light-colored horizontal line (error bar width indicating within-experimental repeat CIs). The experimental repeat-level (within-study) CIs are computed using experimental repeat-level variance and effect (effect ± 1.96·SE, Wald intervals). The black diamond in each group represents the combined effect estimate across experimental repeats (*μ̂*), shown with its 95% meta-analytic CIs (black horizontal line), which accounts for between-experimental repeat variance (*τ*^2^) when present. For perturbations where *τ*^2^ = 0, the combined effect and CI corresponds to a fixed-effect model.

### Pixel-level colocalization analysis of TDP-43 and P-body

To assess spatial colocalization between TDP-43 and P-body, we implemented a pixel-wise overlap analysis using multiplexed site images co-stained for TDP-43 (V5 tag) and P-body marker (DCP1A) in 8 day iPSC-derived neurons expressing TDP-43^ΔNLS^. Site images (1024×1024) were acquired from three experimental repeats, each containing four technical repeats and ∼200 sites (fields of view) per condition. Pixel-level colocalization was quantified using a conditional probability approach: for each image, the fraction of DCP1A-positive pixels that were also positive for TDP-43 was calculated, yielding the colocalization score:

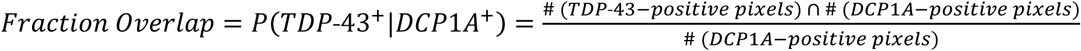

To define “positive” signal pixels, binary masks for each marker were generated by applying per-image intensity thresholds. For TDP-43, the threshold was defined as the mean pixel intensity plus two standard deviations (mean+2×SD), and for DCP1A, the threshold was mean+2.5×SD. Only pixels exceeding these thresholds were considered “positive.”

To determine whether observed colocalization exceeded what would be expected by random spatial proximity, we implemented a rotation-based null model. For each image, the TDP-43 channel was rotated 90° counterclockwise and re-aligned to the original DCP1A channel. This procedure preserved pixel intensities and marginal marker distributions, while disrupting spatial alignment. Real and rotated scores were computed for each image, yielding one colocalization value and one null control per image. A paired Wilcoxon signed-rank test was used to test whether real colocalization was significantly greater than rotated (null) colocalization.

A similar analysis was performed on site images (2275x2284) of 60 day iPSC-derived neurons from subjects (healthy controls or ALS patients) stained for TDP-43 (TDP-43 antibody) and P-body marker (DCP1A) using thresholds of mean+0.5×SD for TDP-43 and mean+3×SD for DCP1A. To estimate the level of colocalization in each human subject group, we applied a linear fixed-effects model with human subject-clustered standard errors (fallback used).

All analyses were performed using Python, with custom utilities implemented in the tools.channels_colocalization.pixel_colocalization_utils module. Visualization and statistical tests were conducted using matplotlib, seaborn, and scipy.stats.

### Post-hoc classifier for neuronal organelle identification: Model training and accuracy

To evaluate model performance, classification was performed on the wild-type group of the day 8 iPSC-derived neurons across 26 organelles spanning three experimental repeats. A three-fold cross-validation scheme was used, where in each fold one experimental repeat was used for training and the remaining two for testing. The classifier was a linear support vector machine (LinearSVC, C=1.0, max_iter=1000, random_state=42) obtained from the *scikit-learn* python package (v1.3.0).

For each fold, accuracy, sensitivity, and specificity were computed per class. Macro-level statistics were obtained by averaging the per-class metrics within each fold, and the overall performance was calculated as the mean of the fold-level macro statistics.

ROC curves were computed in a one-vs.-rest manner for each class in every fold using the classifier decision scores. To obtain curves across folds, decision scores from all folds were concatenated and the ROC curves were recalculated from these combined arrays.

Confusion matrices were generated for each fold and then aggregated across folds. For visualization, the aggregated confusion matrix was row-wise normalized.

### Rollout attention maps

For visualization of attention maps, self-attention tensors were extracted from each attention layer of the ViT model for every image, resulting in a tensor of shape (12,3,50,50), corresponding to 12 attention layers, 3 attention heads per layer and 50*x*50 patch-to-patch attention maps. Each tensor was reduced to shape (12,50,50) by averaging over the attention heads.

The attention mechanism is defined as:

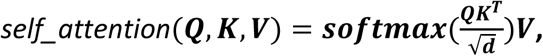

where *Q*, *K*, *V* ∈ *R*^*n*×*d*^ are the query, key, and value tensors for *n* patches with embedding dimension *d*, obtained from the output of the previous layer of the model. The softmax term produces an attention weight matrix *A* ∈ *R*^*n*×*n*^, with entries *a*_*ij*_ representing the contribution of patch *j* to *i*.

To capture the aggregated contribution of all layers, we applied an attention rollout ^185^. Rollout propagates attention across all layers by recursively multiplying the attention matrices:

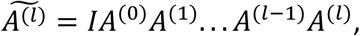

where *A*^(*l*)^ is the attention tensor at layer *l*, and 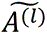 represents the cumulative attention up to layer *l*. This yields a final rollout attention tensor of shape (50,50).

From the resulting tensor, we extracted the values corresponding to the CLS token (the first patch) as its attention distribution over the remaining 49 patches. The resulting values were min-max scaled to the range [0,1].

Then, to match the resolution of the input image, the attention tensor was multiplied by 255 and cast to uint8 to enable resizing with the PIL library. We applied bicubic interpolation to resize the attention tensor to (100,100), matching the image dimensions. Finally, we divided it by 255 to restore the [0,1] range, yielding a resized attention tensor that highlights regions of high model focus with larger values and low model focus with lower values. For visualization purposes, we applied a threshold to display only values exceeding 0.5.

### UMAP, dimensionality reduction of latent space and 2D clustering

To cluster images of organelles from different experimental perturbations, the image representations are subjected to dimensionality reduction using the Uniform Manifold Approximation and Projection (UMAP) algorithm (umap-learn v0.5.3, random_state=1). The first 2 UMAP components were plotted in a scatter plot to visualize clusters and colored according to their experimental perturbation. Our work presents different types of UMAPs based on the model output: single organelle plots (**Figure 2C**, **Figure 4D,E**, **Figure 5C,D,G,H**, **Supplementary Figure 12B**, **Supplementary Figure 14B, Supplementary Figure 15B**, **Supplementary Figure 22C,D**, **Supplementary Figure 24D**), and multiple organelles plots (**Figure 1C**, **Supplementary Figure 3D**, **Supplementary Figure 8**)

### Distances between organelles or human subjects

To quantify the relationships between groups of embeddings shown in a UMAP (organelles / human subjects), pairwise Euclidean distances *d* were computed in the full latent space using 192-dimensional embeddings per image. The median of these pairwise distances was used as a robust summary statistic reflecting their similarity in the embedding space. For visualization, distances were shown in a heatmap after min–max normalized to the range [0, 1], where 0 indicates the closest and 1 the furthest relationship.

### Unbiased CellProfiler analysis

For benchmarking organellome-wide analysis, CellProfiler (version 4.2.1; jdk version 14.0.1; python version 3.8) ^47^ objects were identified in images in three different size ranges to cover all possible organellar stainings; in addition, ‘secondary object’ identification was used for more diffuse cellular stains. Entire site images (1,024x1,024) were used as input images. All main object-related CellProfiler features relating to area, intensity, distribution, neighbors and texture were then extracted. Object measurements were averaged per image and concatenated across object types (using pandas, version 1.4.4). After removing constant features and features containing missing values, the remaining 347 features calculated with Cellprofiler as described above were used as features for classification and UMAP plotting as described above.

### CellProfiler quantification of P-body size

A CellProfiler pipeline for calculating P-body size was applied. The pipeline identified P-bodies in the correct size range and extracted object features with ‘MeasureObjectSizeShape’. Object-level measurements were extracted and averaged per site image. Effect size and statistical significance was calculated with linear mixed-effects models. Linear fixed-effects fallback used when variance was zero. In the case of TDP-43^ΔNLS^, DOX was set as a fixed effect and experimental repeat as a random intercept. In the case of day 60 human subject-derived neurons, effect was estimated per group (C9ALS, sALS+, etc.) while accounting for within-human subject dependences, by modeling the group variable as a fixed effect with human subject-clustered SEs (to handle collinearity of subject ID with group).

### CellProfiler quantification of cytoplasmic/nuclear TDP-43 ratio

A CellProfiler pipeline was applied to quantify the cytoplasmic TDP-43 signal relative to the nuclear TDP-43 signal. First, nuclei were identified with ‘IdentifyPrimaryObjects’. Using nuclei as input objects, the cytoplasm was then identified with ‘IdentifySecondaryObjects’ using the MAP2 channel and the ‘Propagation’ method in case of day 60 human subject-derived neurons. For TDP-43^ΔNLS^ neurons, the TDP-43 channel and the ‘Distance -N’ method were used as input for secondary object identification. Since TDP-43 is also present in the nucleus, a third step was applied to subtract the nucleus from the secondary object using ‘IdentifyTertiaryObjects’, resulting in the cytoplasm. Intensity of TDP-43 staining was then measured in both nucleus and cytoplasm, and the final ratio of cytoplasmic/nuclear TDP-43 was calculated with the ‘CalculateMath’ module.

Effect size and statistical significance was calculated with linear mixed-effects models. Linear fixed-effects fallback used when variance was zero. In the case of TDP-43^ΔNLS^, DOX was set as a fixed effect and experimental repeat as a covariate. In the case of day 60 human subject-derived neurons, effect was estimated per group (C9ALS, sALS+, etc.) by linear mixed-effects models (modeling the human subject group as a fixed effect and human subject as random intercept).

### Prediction of protein-protein interactions

SpeedPPI ^186^ was implemented to predict interaction strength between TDP-43 and P-body proteins. Briefly, the pipeline generates multiple sequence alignments (MSAs) for TDP-43 and its potential interactors using HHlibs. It offers multiple operational modes, including “all-vs-all” and “some-vs-some”; we utilized the “some-vs-some” mode to compare TDP-43 against a defined list of target proteins. The MSAs are then subjected to structure prediction and validation using AlphaFold2 ^113^, providing high-confidence structural models for each protein. These validated structures are fed into the SpeedPPI predictor, a machine learning-based model that integrates several features. The final output is a structural modeling prediction that illustrates the potential interaction interfaces between TDP-43 and its interactors. In addition, a pDockQ score, a probabilistic score ranging from 0 to 1, is generated to estimate the confidence of the predicted interaction. This score integrates structural features such as interface size and pLDDT confidence. Notably, the list of the P-body-enriched and - depleted proteins was based on Hubstenberger et al. ^116^. The list was also determined based on the direction of the most extreme fold-change, with only proteins meeting a p-value < 0.05 without applying a strict fold-change cutoff, considered significant. In addition, proteins that were not also identified as interactors of DDX6 (a key resident protein of P-bodies) as characterized in the literature ^115^ were filtered out. Accordingly, positive controls correspond to FC > 1.58, while negative controls correspond to FC < -0.71.

### Analysis of post-mortem patient RNA-seq data for depleted and enriched P-body genes

Data sources:

● Bulk RNA-seq data from Edward Lee, “Loss of Nuclear TDP-43 Is Associated with Decondensation of LINE Retrotransposons”^140^.
● List of enriched and depleted genes from Hubstenberger et al, “P-Body Purification Reveals the Condensation of Repressed mRNA Regulons”^116^.

Bulk RNA-seq count tables (raw counts and TPM) from the GSE126542 dataset were downloaded, and gene identifiers were mapped to gene symbols using a GRCh38 annotation. To restrict analyses to expressed genes, we computed the median raw count per gene across samples and retained genes with median > 5 in the raw matrix. Per subject, log2 fold change (L2FC) was calculated as log2(TPM_NEG / TPM_POS).

For P-body gene set construction, we retained genes with median count ≥ 5 P-body–enriched genes were defined by FDR ≤ 0.05 and enrichment ≥ 1, and P-body–depleted genes by FDR ≤ 0.05 and enrichment ≤ −1.

Per patient, variable genes were defined as those with absolute L2FC ≥ 1. We then counted up-regulated (L2FC > 0) and down-regulated (L2FC < 0) genes and computed the percentage down-regulated. This was done separately within the enriched and depleted P-body subsets. A 2×2 contingency table per-patient was constructed. One-sided Fisher’s exact tests were performed, with the alternative hypothesis being greater down-regulation in P-body–enriched genes compared to depleted genes. P-values across subjects were adjusted by Benjamini–Hochberg FDR. To combine evidence across subjects while controlling for subject as a stratum, we performed a Cochran–Mantel–Haenszel (CMH) test on the set of per-subject 2×2 tables. The analysis reported the CMH χ² statistic and p-value, the Mantel–Haenszel common odds ratio with a 95% confidence interval obtained via the Mantel–Haenszel method, and a one-sided p-value in the expected direction derived from the two-sided CMH p-value.

## Data availability

Proximity labeling proteomics data are available in ProteomeXchange (PXD056534). SLAM-seq RNA-seq data are available in GEO (GSE279980).

The imaging dataset^187^ can be downloaded via Amazon S3 API. Bucket name: organellomics, region: eu-west-1. (https://organellomics.s3.amazonaws.com). A script for downloading the data is available on our Github repository. To browse the data we suggest using Cyberduck (https://cyberduck.io/). Detailed instructions are provided in the README file on our Github repository.

## Software and hardware

All deep-learning architectures were implemented in Pytorch version 2.2.2^188^, cuda version 11.8 on Python v3.9.19. The training was performed on an NVIDIA A40 (48GB GPU) card.

## Code availability

The code can be found on our Github page: https://github.com/Sagykri/NOVA^189^

## Extendend statistical values

Detailed tables of statistical parameters and their values for each dataset are provided in the supplementary materials files.

## Figure design

We placed and organized all the figures by using Adobe Illustrator software, which was also used for generating Figure 1A. The graphs were generated either by Prism (version 5.9.1 or 10) or via python (v3.9.19; matplotlib (v3.8.4) and seaborn (v0.13.2)). Representative confocal images were generated by FiJi software v. 1.52p. or ZEN software. We generated the graphical abstract, Figures 1B, **2A**, **4L**, **5A,E,R** and **Supplementary** Figures 1**,2,4,6,10,11,** and **17** by Biorender.com.

## Acknowledgments

We wish to thank Leeat Keren (WIS), Maya Schuldiner (WIS), Eran Segal (WIS) Sasha Devore (WIS), Ernest Fraenkel (MIT), Hirofumi Kobayashi (Chan Zuckerberg Biohub), Assaf Zaritzky (BGU), Tomer Lapidot (UPENN), Ophir Shalem (UPENN) and Johnathan Cooper Knock (U Sheffield) for advice during project development and on the manuscript draft. We thank Michael Ward (NINDS, NIH) and Bill Skarnes (Jackson labs) for early access to iNDI lines Nancy Kadersha Paul Anderson and Pavel Ivanov (HMS) for G3BP1 and YFP-DCP1A U2OS cells. Shai Bagon (WIS) and Ron Rotkofp (WIS) for advice in statistical and computational sides. Alon Savidor (WIS) and Yishai Levin (WIS) for Mass spectrometry. For the RNA-seq procedure for SLAM-seq, we wish to thank Inbal Bolocan Nachman (WIS). We thank Chen Eitan, Aviad Siany, Jazz Lubliner, Yahel Cohen, Emmanuel Amzalleg, and Lior Lin for preliminary experiments that are not in the scope of the final manuscript.

EH is the Mondry Family Professorial Chair and Head of the Andrea L. and Lawrence A. Wolfe Family Center for Research on Neuroimmunology and Neuromodulation. Funding for research: Binational Science Foundation (BSF); Association Francaise Contre les Myopathies (AFM); Amyotrophic Lateral Sclerosis Association (ALSA); Target ALS; Israel Science Foundation (ISF 3497/21, 424/22, 494/24); ALS Canada; Minna-James-Heineman Stiftung through Minerva, Minerva Foundation, with funding from the Federal German Ministry for Education and Research; Robert Packard Center for ALS Research at Johns Hopkins; McGill University; EU - ERA-Net; Radala Foundation for ALS Research; Additional support generously provided by the Kekst Family Institute for Medical Genetics. Weizmann SABRA - Yeda-Sela - WRC Program, the Estate of Emile Mimran, and The Maurice and Vivienne Wohl Biology Endowment. Nella and Leon Benoziyo Center for Neurological Diseases. Goldhirsh-Yellin Foundation. Dr. Sydney Brenner and friends. Weizmann - Center for Research on Neurodegeneration. Redhill Foundation – Sam and Jean Rothberg Charitable Trust Dr. Dvora and Haim Teitelbaum Endowment Fund. This research was supported in part by the Intramural research Program of the NIH, National institute on Aging and National Institutes of Neurological Diseases and Stroke. LM is funded by a Minerva Postdoctoral Fellowship. YDM was funded by the Weizmann - CNRS (Centre National de la Recherche Scientifique) Collaboration Program.

## References

1. Gottschling, D. E. & Nyström, T. The Upsides and Downsides of Organelle Interconnectivity. Cell 169, 24–34 (2017).

2. Zung, N. & Schuldiner, M. New horizons in mitochondrial contact site research. Biol. Chem. 401, 793–809 (2020).

3. Itzhak, D. N. et al. A Mass Spectrometry-Based Approach for Mapping Protein Subcellular Localization Reveals the Spatial Proteome of Mouse Primary Neurons. Cell Rep. 20, 2706–2718 (2017).

4. Borner, G. H. H. Organellar Maps Through Proteomic Profiling - A Conceptual Guide. Mol. Cell. Proteomics 19, 1076–1087 (2020).

5. Schessner, J. P., Albrecht, V., Davies, A. K., Sinitcyn, P. & Borner, G. H. H. Deep and fast label-free Dynamic Organellar Mapping. Nat. Commun. 14, 5252 (2023).

6. Itzhak, D. N., Schessner, J. P. & Borner, G. H. H. Dynamic organellar maps for spatial proteomics. Curr. Protoc. Cell Biol. 83, e81 (2019).

7. Thul, P. J. et al. A subcellular map of the human proteome. Science 356, (2017).

8. Youn, J.-Y. et al. High-Density Proximity Mapping Reveals the Subcellular Organization of mRNA-Associated Granules and Bodies. Mol Cell 69, 517–532.e11 (2018).

9. Go, C. D. et al. A proximity-dependent biotinylation map of a human cell. Nature 595, 120–124 (2021).

10. Hein, M. Y. et al. Global organelle profiling reveals subcellular localization and remodeling at proteome scale. Cell 188, 1137–1155.e20 (2025).

11. Kobayashi, H., Cheveralls, K. C., Leonetti, M. D. & Royer, L. A. Self-supervised deep learning encodes high-resolution features of protein subcellular localization. Nat. Methods 19, 995–1003 (2022).

12. Spitzer, H., Berry, S., Donoghoe, M., Pelkmans, L. & Theis, F. J. Learning consistent subcellular landmarks to quantify changes in multiplexed protein maps. Nat. Methods 20, 1058–1069 (2023).

13. Carpenter, A. E. et al. CellProfiler: image analysis software for identifying and quantifying cell phenotypes. Genome Biol. 7, R100 (2006).

14. Heinrich, L. et al. Whole-cell organelle segmentation in volume electron microscopy. Nature 599, 141–146 (2021).

15. Lu, A. X., Kraus, O. Z., Cooper, S. & Moses, A. M. Learning unsupervised feature representations for single cell microscopy images with paired cell inpainting. PLoS Comput. Biol. 15, e1007348 (2019).

16. Husain, S. S. et al. Single-cell subcellular protein localisation using novel ensembles of diverse deep architectures. Commun Biol 6, 489 (2023).

17. Long, W., Yang, Y. & Shen, H.-B. ImPLoc: a multi-instance deep learning model for the prediction of protein subcellular localization based on immunohistochemistry images. Bioinformatics 36, 2244–2250 (2020).

18. Kim, K.-M., Son, K. & Palmore, G. T. R. Neuron Image Analyzer: Automated and Accurate Extraction of Neuronal Data from Low Quality Images. Sci. Rep. 5, 17062 (2015).

19. Ascoli, G. A., Donohue, D. E. & Halavi, M. NeuroMorpho.Org: a central resource for neuronal morphologies. J. Neurosci. 27, 9247–9251 (2007).

20. Oberlaender, M., Bruno, R. M., Sakmann, B. & Broser, P. J. Transmitted light brightfield mosaic microscopy for three-dimensional tracing of single neuron morphology. J. Biomed. Opt. 12, 064029 (2007).

21. Moshkov, N. et al. Learning representations for image-based profiling of perturbations. Nat. Commun. 15, 1594 (2024).

22. Serrano, E., et al. Progress and new challenges in image-based profiling. *arXiv [q-bio.QM]* (2025) doi:10.48550/ARXIV.2508.05800.

23. Seal, S. et al. Cell Painting: a decade of discovery and innovation in cellular imaging. Nat Methods 22, 254–268 (2025).

24. Hirose, T., Ninomiya, K., Nakagawa, S. & Yamazaki, T. A guide to membraneless organelles and their various roles in gene regulation. Nat. Rev. Mol. Cell Biol. (2022) doi:10.1038/s41580-022-00558-8.

25. Banani, S. F., Lee, H. O., Hyman, A. A. & Rosen, M. K. Biomolecular condensates: organizers of cellular biochemistry. Nat. Rev. Mol. Cell Biol. 18, 285–298 (2017).

26. Brangwynne, C. P. Phase transitions and size scaling of membrane-less organelles. J. Cell Biol. 203, 875–881 (2013).

27. Lyon, A. S., Peeples, W. B. & Rosen, M. K. A framework for understanding the functions of biomolecular condensates across scales. Nat. Rev. Mol. Cell Biol. 22, 215–235 (2021).

28. Alberti, S. & Hyman, A. A. Biomolecular condensates at the nexus of cellular stress, protein aggregation disease and ageing. Nat. Rev. Mol. Cell Biol. 22, 196–213 (2021).

29. Platt, F. M., d’Azzo, A., Davidson, B. L., Neufeld, E. F. & Tifft, C. J. Lysosomal storage diseases. Nat Rev Dis Primers 4, 27 (2018).

30. Cox, T. M. & Cachón-González, M. B. The cellular pathology of lysosomal diseases. J. Pathol. 226, 241–254 (2012).

31. Gorman, G. S. et al. Mitochondrial diseases. Nat Rev Dis Primers 2, 16080 (2016).

32. Schapira, A. H. V. Mitochondrial disease. Lancet 368, 70–82 (2006).

33. Levine, B. & Kroemer, G. Autophagy in the pathogenesis of disease. Cell 132, 27–42 (2008).

34. Mizushima, N. & Levine, B. Autophagy in Human Diseases. N. Engl. J. Med. 383, 1564–1576 (2020).

35. Nedelsky, N. B. & Taylor, J. P. Bridging biophysics and neurology: aberrant phase transitions in neurodegenerative disease. Nat. Rev. Neurol. 15, 272–286 (2019).

36. Alberti, S. & Dormann, D. Liquid-Liquid Phase Separation in Disease. Annu. Rev. Genet. 53, 171–194 (2019).

37. Dosovitskiy, A., et al. An image is worth 16×16 words: Transformers for image recognition at scale. *arXiv [cs.CV]* (2020).

38. Kim, V., Adaloglou, N., Osterland, M., Morelli, F. & Zapata, P. A. M. Self-supervision advances morphological profiling by unlocking powerful image representations. (2023) doi:10.1101/2023.04.28.538691.

39. Doron, M., et al. Unbiased single-cell morphology with self-supervised vision transformers*. bioRxiv* (2023).

40. Kraus, O., et al. Masked autoencoders for microscopy are scalable learners of cellular biology*. arXiv [cs.CV]* (2024).

41. Atmaramani, R., et al. Deep Learning Analysis on Images of iPSC-derived Motor Neurons Carrying fALS-genetics Reveals Disease-Relevant Phenotypes. *bioRxiv* (2024).

42. Natekar, P., Wang, Z., Arora, M., Hakozaki, H. & Schöneberg, J. Self-supervised deep learning uncovers the semantic landscape of drug-induced latent mitochondrial phenotypes. bioRxiv (2023) doi:10.1101/2023.09.13.557636.

43. Perakis, A. et al. Contrastive learning of single-cell phenotypic representations for treatment classification. in Machine Learning in Medical Imaging 565–575 (Springer International Publishing, Cham, 2021).

44. van den Oord, A., Li, Y. & Vinyals, O. Representation learning with Contrastive Predictive Coding. arXiv [cs.LG] (2018).

45. Khosla, P., et al. Supervised Contrastive Learning*. arXiv [cs.LG]* (2020).

46. Bray, M.-A. et al. Cell Painting, a high-content image-based assay for morphological profiling using multiplexed fluorescent dyes. Nat. Protoc. 11, 1757–1774 (2016).

47. Stirling, D. R. et al. CellProfiler 4: improvements in speed, utility and usability. BMC Bioinformatics 22, 433 (2021).

48. Cho, N. H. et al. OpenCell: Endogenous tagging for the cartography of human cellular organization. Science 375, eabi6983 (2022).

49. Ye, P., et al. Partial Fine-tuning: A successor to full fine-tuning for vision transformers. *arXiv [cs.CV]* (2023).

50. McInnes, L., Healy, J. & Melville, J. UMAP: Uniform Manifold Approximation and Projection for Dimension Reduction. *arXiv [stat.ML]* (2018).

51. Shai, N. et al. Systematic mapping of contact sites reveals tethers and a function for the peroxisome-mitochondria contact. Nat Commun 9, 1761 (2018).

52. Wong, Y. C., Ysselstein, D. & Krainc, D. Mitochondria-lysosome contacts regulate mitochondrial fission via RAB7 GTP hydrolysis. Nature 554, 382–386 (2018).

53. DiGiovanni, L. F. et al. ROS transfer at peroxisome-mitochondria contact regulates mitochondrial redox. Science 389, 157–162 (2025).

54. Chu, B.-B. et al. Cholesterol transport through lysosome-peroxisome membrane contacts. Cell 184, 289 (2021).

55. Deus, C. M., Yambire, K. F., Oliveira, P. J. & Raimundo, N. Mitochondria-lysosome crosstalk: From physiology to neurodegeneration. Trends Mol. Med. 26, 71–88 (2020).

56. Kim, S., Wong, Y. C., Gao, F. & Krainc, D. Dysregulation of mitochondria-lysosome contacts by GBA1 dysfunction in dopaminergic neuronal models of Parkinson’s disease. Nat Commun 12, 1807 (2021).

57. Du, M., Ea, C.-K., Fang, Y. & Chen, Z. J. Liquid phase separation of NEMO induced by polyubiquitin chains activates NF-κB. Mol. Cell 82, 2415–2426.e5 (2022).

58. West, J. A. et al. Structural, super-resolution microscopy analysis of paraspeckle nuclear body organization. J. Cell Biol. 214, 817–830 (2016).

59. Naganuma, T. et al. Alternative 3’-end processing of long noncoding RNA initiates construction of nuclear paraspeckles. EMBO J. 31, 4020–4034 (2012).

60. Arseni, D. et al. Heteromeric amyloid filaments of ANXA11 and TDP-43 in FTLD-TDP Type C. Nature (2024) doi:10.1038/s41586-024-08024-5.

61. Robinson, J. L. et al. Annexin A11 aggregation in FTLD-TDP type C and related neurodegenerative disease proteinopathies. Acta Neuropathol 147, 104 (2024).

62. Miedema, S. S. M. et al. Proteomics of the temporal cortex in semantic dementia reveals brain-region specific molecular pathology and regulation of the TDP-43-ANXA11 interactome. Acta Neuropathol Commun 13, 162 (2025).

63. Hodgson, R. et al. TDP-43 is a Master Regulator of Paraspeckle Condensation. (2024) doi:10.2139/ssrn.4721338.

64. Nishimoto, Y. et al. The long non-coding RNA nuclear-enriched abundant transcript 1_2 induces paraspeckle formation in the motor neuron during the early phase of amyotrophic lateral sclerosis. Mol Brain 6, 31 (2013).

65. Chazotte, B. Labeling nuclear DNA with hoechst 33342. Cold Spring Harb. Protoc. 2011, db.prot5557 (2011).

66. Chazotte, B. Labeling mitochondria with MitoTracker dyes. Cold Spring Harb. Protoc. 2011, 990–992 (2011).

67. Wurm, C. A. et al. Nanoscale distribution of mitochondrial import receptor Tom20 is adjusted to cellular conditions and exhibits an inner-cellular gradient. Proc. Natl. Acad. Sci. U. S. A. 108, 13546–13551 (2011).

68. Turner, D. L., Korneev, D. V., Purdy, J. G., de Marco, A. & Mathias, R. A. The host exosome pathway underpins biogenesis of the human cytomegalovirus virion. Elife 9, e58288 (2020).

69. Nava, C. et al. Hypomorphic variants of cationic amino acid transporter 3 in males with autism spectrum disorders. Amino Acids 47, 2647–2658 (2015).

70. Chazotte, B. Labeling cytoskeletal F-actin with rhodamine phalloidin or fluorescein phalloidin for imaging. Cold Spring Harb. Protoc. 2010, db.prot4947 (2010).

71. Brinkley, B. R., Fistel, S. H., Marcum, J. M. & Pardue, R. L. Microtubules in cultured cells; indirect immunofluorescent staining with tubulin antibody. Int. Rev. Cytol. 63, 59–95 (1980).

72. Zocchi, R., Compagnucci, C., Bertini, E. & Sferra, A. Deciphering the tubulin language: Molecular determinants and readout mechanisms of the tubulin code in neurons. Int. J. Mol. Sci. 24, 2781 (2023).

73. Lauria, G. et al. Tubule and neurofilament immunoreactivity in human hairy skin: markers for intraepidermal nerve fibers. Muscle Nerve 30, 310–316 (2004).

74. Kageyama, S. et al. p62/SQSTM1-droplet serves as a platform for autophagosome formation and anti-oxidative stress response. Nat. Commun. 12, 16 (2021).

75. Gowrishankar, S., Wu, Y. & Ferguson, S. M. Impaired JIP3-dependent axonal lysosome transport promotes amyloid plaque pathology. J. Cell Biol. 216, 3291–3305 (2017).

76. Koyano, F. et al. Parkin-mediated ubiquitylation redistributes MITOL/March5 from mitochondria to peroxisomes. EMBO Rep. 20, e47728 (2019).

77. Tsygankova, O. M. & Keen, J. H. A unique role for clathrin light chain A in cell spreading and migration. J. Cell Sci. 132, (2019).

78. Srivastava, M., Fleming, P. J., Pollard, H. B. & Burns, A. L. Cloning and sequencing of the human nucleolin cDNA. FEBS Lett. 250, 99–105 (1989).

79. Fox, A. H., Bond, C. S. & Lamond, A. I. P54nrb forms a heterodimer with PSP1 that localizes to paraspeckles in an RNA-dependent manner. Mol. Biol. Cell 16, 5304–5315 (2005).

80. Fox, A. H. et al. Paraspeckles: a novel nuclear domain. Curr. Biol. 12, 13–25 (2002).

81. Tsuji, H. et al. Epitope mapping of antibodies against TDP-43 and detection of protease-resistant fragments of pathological TDP-43 in amyotrophic lateral sclerosis and frontotemporal lobar degeneration. Biochem. Biophys. Res. Commun. 417, 116–121 (2012).

82. Yang, L., Gal, J., Chen, J. & Zhu, H. Self-assembled FUS binds active chromatin and regulates gene transcription. Proc. Natl. Acad. Sci. U. S. A. 111, 17809–17814 (2014).

83. Weis, K. et al. Retinoic acid regulates aberrant nuclear localization of PML-RAR alpha in acute promyelocytic leukemia cells. Cell 76, 345–356 (1994).

84. Eystathioy, T. et al. The GW182 protein colocalizes with mRNA degradation associated proteins hDcp1 and hLSm4 in cytoplasmic GW bodies. RNA 9, 1171–1173 (2003).

85. Yang, W.-H., Yu, J. H., Gulick, T., Bloch, K. D. & Bloch, D. B. RNA-associated protein 55 (RAP55) localizes to mRNA processing bodies and stress granules. RNA 12, 547–554 (2006).

86. Tourrière, H. et al. The RasGAP-associated endoribonuclease G3BP assembles stress granules. J. Cell Biol. 160, 823–831 (2003).

87. Liao, Y.-C. et al. RNA Granules Hitchhike on Lysosomes for Long-Distance Transport, Using Annexin A11 as a Molecular Tether. Cell 179, 147–164.e20 (2019).

88. Daigle, J. G. et al. Pur-alpha regulates cytoplasmic stress granule dynamics and ameliorates FUS toxicity. Acta Neuropathol. 131, 605–620 (2016).

89. Molitor, L. et al. Depletion of the RNA-binding protein PURA triggers changes in posttranscriptional gene regulation and loss of P-bodies. Nucleic Acids Res. 51, 1297–1316 (2023).

90. Liu, M. et al. KIF5A-dependent axonal transport deficiency disrupts autophagic flux in trimethyltin chloride-induced neurotoxicity. Autophagy 17, 903–924 (2021).

91. Ray, S. et al. α-Synuclein aggregation nucleates through liquid-liquid phase separation. Nat. Chem. 12, 705–716 (2020).

92. Bayer, T. A. et al. Neural expression profile of alpha-synuclein in developing human cortex. Neuroreport 10, 2799–2803 (1999).

93. Sampedro, M. N., Bussineau, C. M. & Cotman, C. W. Postsynaptic density antigens: preparation and characterization of an antiserum against postsynaptic densities. J. Cell Biol. 90, 675–686 (1981).

94. Lai, A., Valdez-Sinon, A. N. & Bassell, G. J. Regulation of RNA granules by FMRP and implications for neurological diseases. Traffic 21, 454–462 (2020).

95. Sharma, A., Takata, H., Shibahara, K.-I., Bubulya, A. & Bubulya, P. A. Son is essential for nuclear speckle organization and cell cycle progression. Mol. Biol. Cell 21, 650–663 (2010).

96. Protter, D. S. W. & Parker, R. Principles and Properties of Stress Granules. Trends Cell Biol. 26, 668–679 (2016).

97. Anderson, P. & Kedersha, N. RNA granules: post-transcriptional and epigenetic modulators of gene expression. Nat. Rev. Mol. Cell Biol. 10, 430–436 (2009).

98. Ramos, D. M., Skarnes, W. C., Singleton, A. B., Cookson, M. R. & Ward, M. E. Tackling neurodegenerative diseases with genomic engineering: A new stem cell initiative from the NIH. Neuron 109, 1080–1083 (2021).

99. Waibel, S., Neumann, M., Rabe, M., Meyer, T. & Ludolph, A. C. Novel missense and truncating mutations in FUS/TLS in familial ALS. Neurology 75, 815–817 (2010).

100. Bosco, D. A. et al. Mutant FUS proteins that cause amyotrophic lateral sclerosis incorporate into stress granules. Hum Mol Genet 19, 4160–4175 (2010).

101. Rutherford, N. J. et al. Novel mutations in TARDBP (TDP-43) in patients with familial amyotrophic lateral sclerosis. PLoS Genet. 4, e1000193 (2008).

102. Sreedharan, J. et al. TDP-43 mutations in familial and sporadic amyotrophic lateral sclerosis. Science 319, 1668–1672 (2008).

103. Kenna, K. P. et al. Delineating the genetic heterogeneity of ALS using targeted high-throughput sequencing. J. Med. Genet. 50, 776–783 (2013).

104. Freischmidt, A. et al. Haploinsufficiency of TBK1 causes familial ALS and fronto-temporal dementia. Nat. Neurosci. 18, 631–636 (2015).

105. Rothstein, J. D. et al. Sporadic ALS induced pluripotent stem cell derived neurons reveal hallmarks of TDP-43 loss of function. Nat Commun 16, 7092 (2025).

106. Neumann, M. et al. Ubiquitinated TDP-43 in frontotemporal lobar degeneration and amyotrophic lateral sclerosis. Science 314, 130–133 (2006).

107. Balendra, R. et al. Amyotrophic lateral sclerosis caused by TARDBP mutations: from genetics to TDP-43 proteinopathy. Lancet Neurol. 24, 456–470 (2025).

108. Birsa, N., Bentham, M. P. & Fratta, P. Cytoplasmic functions of TDP-43 and FUS and their role in ALS. Semin Cell Dev Biol 99, 193–201 (2020).

109. Wagner, K. et al. Induced proximity to PML protects TDP-43 from aggregation via SUMO-ubiquitin networks. Nat Chem Biol 21, 1408–1419 (2025).

110. Antoniani, F. et al. Loss of PML nuclear bodies in familial amyotrophic lateral sclerosis-frontotemporal dementia. Cell Death Discov 9, 248 (2023).

111. Gasset-Rosa, F. et al. Cytoplasmic TDP-43 De-mixing Independent of Stress Granules Drives Inhibition of Nuclear Import, Loss of Nuclear TDP-43, and Cell Death. Neuron 102, 339–357.e7 (2019).

112. Bryant, P. & Noé, F. Improved protein complex prediction with AlphaFold-multimer by denoising the MSA profile. PLoS Comput. Biol. 20, e1012253 (2024).

113. Bryant, P., Pozzati, G. & Elofsson, A. Improved prediction of protein-protein interactions using AlphaFold2. Nat. Commun. 13, 1265 (2022).

114. Burke, D. F. et al. Towards a structurally resolved human protein interaction network. Nat. Struct. Mol. Biol. 30, 216–225 (2 2023).

115. Ayache, J. et al. P-body assembly requires DDX6 repression complexes rather than decay or Ataxin2/2L complexes. Mol Biol Cell 26, 2579–2595 (2015).

116. Hubstenberger, A. et al. P-Body Purification Reveals the Condensation of Repressed mRNA Regulons. Mol. Cell 68, 144–157.e5 (10 2017).

117. Marmor-Kollet, H. et al. Spatiotemporal Proteomic Analysis of Stress Granule Disassembly Using APEX Reveals Regulation by SUMOylation and Links to ALS Pathogenesis. Mol. Cell 80, 876–891.e6 (2020).

118. Lam, S. S. et al. Directed evolution of APEX2 for electron microscopy and proximity labeling. Nat. Methods 12, 51–54 (1 2014).

119. Hung, V. et al. Spatially resolved proteomic mapping in living cells with the engineered peroxidase APEX2. Nat. Protoc. 11, 456–475 (2016).

120. Ashburner, M. et al. Gene Ontology: tool for the unification of biology. Nature Genetics 2000 25:1 25, 25–29 (5 2000).

121. Consortium, T. G. O. et al. The Gene Ontology knowledgebase in 2023. Genetics 224, (5 2023).

122. Liu, J., Valencia-Sanchez, M. A., Hannon, G. J. & Parker, R. MicroRNA-dependent localization of targeted mRNAs to mammalian P-bodies. Nat Cell Biol 7, 719–723 (2005).

123. Sen, G. L. & Blau, H. M. Argonaute 2/RISC resides in sites of mammalian mRNA decay known as cytoplasmic bodies. Nat Cell Biol 7, 633–636 (2005).

124. Standart, N. & Weil, D. P-bodies: Cytosolic droplets for coordinated mRNA storage. Trends Genet. 34, 612–626 (2018).

125. Jain, S. & Parker, R. The discovery and analysis of P Bodies. Adv. Exp. Med. Biol. 768, 23–43 (2013).

126. Gasset-Rosa, F. et al. Cytoplasmic TDP-43 De-mixing Independent of Stress Granules Drives Inhibition of Nuclear Import, Loss of Nuclear TDP-43, and Cell Death. Neuron vol. 102 339–357.e7 Preprint at 10.1016/j.neuron.2019.02.038 (4 2019).

127. Dewey, C. M. et al. TDP-43 is directed to stress granules by sorbitol, a novel physiological osmotic and oxidative stressor. Mol. Cell. Biol. 31, 1098–1108 (2011).

128. Lee, Y.-B. et al. Cytoplasmic TDP-43 is involved in cell fate during stress recovery. Hum. Mol. Genet. 31, 166–175 (2021).

129. Luo, Y., Na, Z. & Slavoff, S. A. P-Bodies: Composition, Properties, and Functions. Biochemistry 57, 2424–2431 (5 2018).

130. Herzog, V. A. et al. Thiol-linked alkylation of RNA to assess expression dynamics. Nat. Methods 14, 1198–1204 (12 2017).

131. Jürges, C., Dölken, L. & Erhard, F. Dissecting newly transcribed and old RNA using GRAND-SLAM. Bioinformatics 34, i218–i226 (2018).

132. Rummel, T., Sakellaridi, L. & Erhard, F. grandR: a comprehensive package for nucleotide conversion RNA-seq data analysis. Nat. Commun. 14, 3559 (2023).

133. Baskerville, V., Rapuri, S., Mehlhop, E. & Coyne, A. N. SUN1 facilitates CHMP7 nuclear influx and injury cascades in sporadic amyotrophic lateral sclerosis. Brain 147, 109–121 (2024).

134. Macleod, M., et al. The MDAR (Materials Design Analysis Reporting) Framework for transparent reporting in the life sciences. Proc Natl Acad Sci U S A 118, (2021).

135. Almeida, S. & Gao, F.-B. Lost & found: C9ORF72 and the autophagy pathway in ALS/FTD. EMBO J 35, 1251–1253 (2016).

136. Beckers, J., Tharkeshwar, A. K. & Van Damme, P. C9orf72 ALS-FTD: recent evidence for dysregulation of the autophagy-lysosome pathway at multiple levels. Autophagy 17, 3306–3322 (2021).

137. Trist, B. G. et al. Co-deposition of SOD1, TDP-43 and p62 proteinopathies in ALS: evidence for multifaceted pathways underlying neurodegeneration. Acta Neuropathol. Commun. 10, 122 (2022).

138. Guise, A. J. et al. TDP-43-stratified single-cell proteomics of postmortem human spinal motor neurons reveals protein dynamics in amyotrophic lateral sclerosis. Cell Rep. 43, 113636 (2024).

139. Spence, H. et al. RNA aptamer reveals nuclear TDP-43 pathology is an early aggregation event that coincides with STMN-2 cryptic splicing and precedes clinical manifestation in ALS. Acta Neuropathol. 147, 50 (2024).

140. Liu, E. Y. et al. Loss of Nuclear TDP-43 Is Associated with Decondensation of LINE Retrotransposons. Cell Rep 27, 1409–1421.e6 (2019).

141. Workman, M. J. et al. Large-scale differentiation of iPSC-derived motor neurons from ALS and control subjects. Neuron 111, 1191–1204.e5 (2023).

142. Bussi, C. et al. Stress granules plug and stabilize damaged endolysosomal membranes. Nature 623, 1062–1069 (2023).

143. Amen, T. & Kaganovich, D. Stress granules inhibit fatty acid oxidation by modulating mitochondrial permeability. Cell Rep. 35, 109237 (2021).

144. Qin, W. et al. Dynamic mapping of proteome trafficking within and between living cells by TransitID. Cell 186, 3307–3324.e30 (2023).

145. Stoecklin, G. & Kedersha, N. Relationship of GW/P-bodies with stress granules. Adv. Exp. Med. Biol. 768, 197–211 (2013).

146. Cohen, S., Valm, A. M. & Lippincott-Schwartz, J. Interacting organelles. Curr. Opin. Cell Biol. 53, 84–91 (2018).

147. Donahue, E. K. F., Ruark, E. M. & Burkewitz, K. Fundamental roles for inter-organelle communication in aging. Biochem. Soc. Trans. 50, 1389–1402 (2022).

148. Rossini, M., Pizzo, P. & Filadi, R. Better to keep in touch: investigating inter-organelle cross-talk. FEBS J. 288, 740–755 (2021).

149. Schrader, M., Kamoshita, M. & Islinger, M. Organelle interplay-peroxisome interactions in health and disease. J. Inherit. Metab. Dis. 43, 71–89 (2020).

150. Mitrea, D. M. et al. Methods for Physical Characterization of Phase-Separated Bodies and Membrane-less Organelles. J. Mol. Biol. 430, 4773–4805 (2018).

151. Shin, M.-G., et al. RMeDPower for Biology: guiding design, experimental structure and analyses of repeated measures data for biological studies. *bioRxiv* 2022.07.18.500490 (2022) doi:10.1101/2022.07.18.500490.

152. Black, S. et al. CODEX multiplexed tissue imaging with DNA-conjugated antibodies. Nat. Protoc. 16, 3802–3835 (2021).

153. Gut, G., Herrmann & Pelkmans, L. Multiplexed protein maps link subcellular organization to cellular states. Science 361, (2018).

154. Singh, S. P. et al. 3D Deep Learning on Medical Images: A Review. Sensors 20, (2020).

155. Caicedo, J. C. et al. Data-analysis strategies for image-based cell profiling. Nat. Methods 14, 849–863 (2017).

156. Sun, J., Tárnok, A. & Su, X. Deep learning-based single-cell optical image studies. Cytometry A 97, 226–240 (2020).

157. Lavitt, F., Rijlaarsdam, D. J., van der Linden, D., Weglarz-Tomczak, E. & Tomczak, J. M. Deep learning and transfer learning for automatic cell counting in microscope images of human cancer cell lines. Appl. Sci. 11, 4912 (2021).

158. Sahin, U. et al. Oxidative stress-induced assembly of PML nuclear bodies controls sumoylation of partner proteins. J Cell Biol 204, 931–945 (2014).

159. Chen, J. et al. Oxidative stress disrupts the cytoskeleton of spinal motor neurons. Brain Behav 13, e2870 (2023).

160. Rhoads, S. N. et al. Neurons and astrocytes have distinct organelle signatures and responses to stress. Cell Rep 44, 116280 (2025).

161. Mutihac, R. et al. TARDBP pathogenic mutations increase cytoplasmic translocation of TDP-43 and cause reduction of endoplasmic reticulum Ca^2+^ signaling in motor neurons. Neurobiol. Dis. 75, 64–77 (2015).

162. Hulme, A. J., Maksour, S., St-Clair Glover, M., Miellet, S. & Dottori, M. Making neurons, made easy: The use of Neurogenin-2 in neuronal differentiation. Stem Cell Reports 17, 14–34 (2022).

163. Hofweber, M. et al. Phase Separation of FUS Is Suppressed by Its Nuclear Import Receptor and Arginine Methylation. Cell 173, 706–719.e13 (2018).

164. Yoshizawa, T. et al. Nuclear Import Receptor Inhibits Phase Separation of FUS through Binding to Multiple Sites. Cell 173, 693–705.e22 (2018).

165. Dormann, D. et al. ALS-associated fused in sarcoma (FUS) mutations disrupt Transportin-mediated nuclear import. EMBO J. 29, 2841–2857 (2010).

166. Aulas, A. et al. G3BP1 promotes stress-induced RNA granule interactions to preserve polyadenylated mRNA. J. Cell Biol. 209, 73–84 (2015).

167. Ye, Y., et al. DCPS modulates TDP-43 mediated neurodegeneration through P-body regulation*. bioRxiv* (2025) doi:10.1101/2025.06.13.659508.

168. Xie, L., et al. Context-dependent Interactors Regulate TDP-43 Dysfunction in ALS/FTLD. *bioRxiv* (2025) doi:10.1101/2025.04.07.646890.

169. Pantazis, C. B. et al. A reference human induced pluripotent stem cell line for large-scale collaborative studies. Cell Stem Cell 29, 1685–1702.e22 (2022).

170. Fernandopulle, M. S. et al. Transcription Factor-Mediated Differentiation of Human iPSCs into Neurons. Curr. Protoc. Cell Biol. 79, e51 (2018).

171. Lee, S.-Y. et al. APEX Fingerprinting Reveals the Subcellular Localization of Proteins of Interest. Cell Rep. 15, 1837–1847 (2016).

172. Baxi, E. G. et al. Answer ALS, a large-scale resource for sporadic and familial ALS combining clinical and multi-omics data from induced pluripotent cell lines. Nat Neurosci 25, 226–237 (2022).

173. Rothstein, J. D., Warlick, C. & Coyne, A. N. Highly variable molecular signatures of TDP-43 loss of function are associated with nuclear pore complex injury in a population study of sporadic ALS patient iPSNs*. bioRxiv* (2023) doi:10.1101/2023.12.12.571299.

174. Coyne, A. N. et al. Nuclear accumulation of CHMP7 initiates nuclear pore complex injury and subsequent TDP-43 dysfunction in sporadic and familial ALS. Sci. Transl. Med. 13, (2021).

175. Cox, J. & Mann, M. MaxQuant enables high peptide identification rates, individualized p.p.b.-range mass accuracies and proteome-wide protein quantification. Nat. Biotechnol. 26, 1367–1372 (2008).

176. Tyanova, S. et al. The Perseus computational platform for comprehensive analysis of (prote)omics data. Nature Methods 2016 13:9 13, 731–740 (6 2016).

177. Waldron, F. M., Rifai, O. & Gregory, J. Antibody and TDP-43 RNA aptamer dual staining to detect patterns of co-pathology in FFPE-preserved human t. (2024).

178. Jaitin, D. A. et al. Massively parallel single-cell RNA-seq for marker-free decomposition of tissues into cell types. Science 343, 776–779 (2014).

179. Keren-Shaul, H. et al. MARS-seq2.0: an experimental and analytical pipeline for indexed sorting combined with single-cell RNA sequencing. Nat. Protoc. 14, 1841–1862 (2019).

180. Dobin, A. et al. STAR: ultrafast universal RNA-seq aligner. Bioinformatics 29, 15–21 (2013).

181. Finkel, Y. et al. SARS-CoV-2 uses a multipronged strategy to impede host protein synthesis. Nature 594, 240–245 (2021).

182. Brenner, J. F. et al. An automated microscope for cytologic research a preliminary evaluation. J. Histochem. Cytochem. 24, 100–111 (1976).

183. Stringer, C., Wang, T., Michaelos, M. & Pachitariu, M. Cellpose: a generalist algorithm for cellular segmentation. Nat. Methods 18, 100–106 (2021).

184. Bozinovski,S. Reminder of the first paper on transfer learning in neural networks, 1976. Informatica (Ljubl.) 44, (2020).

185. Abnar, S. & Zuidema, W. Quantifying attention flow in transformers*. arXiv [cs.LG]* (2020) doi:10.48550/ARXIV.2005.00928.

186. Bryant, P. & Noé, F. Rapid protein-protein interaction network creation from multiple sequence alignments with Deep Learning. bioRxiv 2023.04.15.536993 (4 2023).

187. Krispin, S. & Van Zuiden, W., Danino, Y., Rudberg, N., Molitor, L., Abarbanel, B., Aviram, G., Wolf, G., Bar, C., Siani, A., Meimoun, T., Coyne, A., Santiana, M., Weller, C., Cookson, M., Ward, M., Waldron, F., Gregory, J., Fisher, T., Nachshon, A., Stern-Ginossar, N., Yacovzada, N., Hornstein, E. Organellomics: AI-driven deep organellar phenotyping reveals novel ALS mechanisms in human neurons (Dataset). Zenodo. 10.5281/zenodo.18152964.

188. Paszke, A. et al. Pytorch: An imperative style, high-performance deep learning library. Adv. Neural Inf. Process. Syst. 32, (2019).

189. Krispin, S. & Van Zuiden, W., Danino, Y., Rudberg, N., Molitor, L., Abarbanel, B., Aviram, G., Wolf, G., Bar, C., Siani, A., Meimoun, T., Coyne, A., Santiana, M., Weller, C., Cookson, M., Ward, M., Waldron, F., Gregory, J., Fisher, T., Nachshon, A., Stern-Ginossar, N., Yacovzada, N., Hornstein, E. Organellomics: AI-driven deep organellar phenotyping reveals novel ALS mechanisms in human neurons (Source Code). Zenodo. 10.5281/zenodo.18152732.

