## Supplementary material for "Organellomics: AI-driven deep organellar phenotyping reveals novel ALS mechanisms in human neurons": Data Summary.docx

**Day 8 iPSC-derived cortical neurons (used for model training)**

Number of cells per site (on average): 17
Number of tiles per site: 64
Number of valid tiles per site (on average): 10
Number of nuclei per tile (on average): 1.5
Number of valid tiles: 3,213,360
Number of neurons: ~689,149

**Day 8 iPSC-derived cortical neurons (used for model inference, Figures 1,2)**

Number of cells per site (on average): 26
Number of tiles per site: 64
Number of valid tiles per site (on average): 12
Number of nuclei per tile (on average): 1.7
Number of valid tiles: 590,634
Number of neurons: ~110,373
Minimum #tiles per organelle perturbation for meta analysis: 80

**Day 8 iPSC-derived cortical neurons (used for model inference, Figure 3)**

Number of cells per site (on average): 23
Number of tiles per site: 64
Number of valid tiles per site (on average): 8.5
Number of nuclei per tile (on average): 1.7
Number of valid tiles: 6,495,286
Number of neurons: ~1,250,175
Minimum #tiles per organelle perturbation for meta analysis: 500

**TDP-43^ΔNLS^ iPSC-derived neurons**

Number of cells per site (on average): 23
Number of tiles per site: 64
Number of valid tiles per site (on average): 5
Number of nuclei per tile (on average): 2
Number of valid tiles: 1,192,604
Number of neurons: ~485,916
Minimum #tiles per organelle perturbation for meta analysis: 80

**sALS patient-derived motor neurons (day 60) (4 organelles)**

Number of cells per site (on average): 20
Number of tiles per site: 49
Number of valid tiles per site (on average): 10
Number of nuclei per tile (on average): 2.5
Number of valid tiles: 9,152
Number of neurons: ~1,986
Minimum #tiles per organelle perturbation for meta analysis: 200

**sALS patient-derived motor neurons (day 60) (32 organelles)**

Number of cells per site (on average): 20
Number of tiles per site: 49
Number of valid tiles per site (on average): 10
Number of nuclei per tile (on average): 3
Number of valid tiles: 58,060
Number of neurons: ~11,647
Minimum #tiles per organelle perturbation for meta analysis: 8
