## Supplementary material for "Organellomics: AI-driven deep organellar phenotyping reveals novel ALS mechanisms in human neurons": Day 8 iPSC-derived cortical neurons (used for model inference, Fig 2) - QC Report.pdf

#### Processed Files Validation

Number of sites that survived pre-processing

(See next page)

### Batch 1

| | Rep | WT stress | WT Untreated |
|---|---|---|---|
| ANAX11 | rep1 | 22 | 25 |
| ANAX11 | rep2 | 25 | 25 |
| ANAX11 | rep3 | 25 | 25 |
| ANAX11 | rep4 | 25 | 25 |
| ANAX11 | rep5 | 24 | 25 |
| ANAX11 | rep6 | 24 | 24 |
| ANAX11 | rep7 | 25 | 25 |
| ANAX11 | rep8 | 25 | 25 |
| SNCA | rep1 | 22 | 24 |
| SNCA | rep2 | 25 | 25 |
| SNCA | rep3 | 24 | 25 |
| SNCA | rep4 | 25 | 25 |
| SNCA | rep5 | 23 | 25 |
| SNCA | rep6 | 24 | 24 |
| SNCA | rep7 | 24 | 23 |
| SNCA | rep8 | 25 | 17 |
| TUJ1 | rep1 | 193 | 200 |
| TUJ1 | rep2 | 200 | 200 |
| TUJ1 | rep3 | 198 | 200 |
| TUJ1 | rep4 | 200 | 200 |
| TUJ1 | rep5 | 199 | 198 |
| TUJ1 | rep6 | 199 | 199 |
| TUJ1 | rep7 | 199 | 199 |
| TUJ1 | rep8 | 200 | 200 |
| DCP1A | rep1 | 25 | 21 |
| DCP1A | rep2 | 25 | 19 |
| DCP1A | rep3 | 24 | 14 |
| DCP1A | rep4 | 25 | 11 |
| DCP1A | rep5 | 25 | 21 |
| DCP1A | rep6 | 24 | 22 |
| DCP1A | rep7 | 25 | 23 |
| DCP1A | rep8 | 25 | 23 |
| NEMO | rep1 | 25 | 25 |
| NEMO | rep2 | 25 | 25 |
| NEMO | rep3 | 24 | 24 |
| NEMO | rep4 | 25 | 25 |
| NEMO | rep5 | 25 | 25 |
| NEMO | rep6 | 25 | 25 |
| NEMO | rep7 | 25 | 25 |
| NEMO | rep8 | 25 | 25 |
| CLTC | rep1 | 25 | 20 |
| CLTC | rep2 | 16 | 23 |
| CLTC | rep3 | 11 | 24 |
| CLTC | rep4 | 11 | 21 |
| CLTC | rep5 | 23 | 24 |
| CLTC | rep6 | 25 | 17 |
| CLTC | rep7 | 14 | 24 |
| CLTC | rep8 | 14 | 25 |
| PSD95 | rep1 | 15 | 15 |
| PSD95 | rep2 | 20 | 10 |
| PSD95 | rep3 | 18 | 22 |
| PSD95 | rep4 | 23 | 25 |
| PSD95 | rep5 | 11 | 19 |
| PSD95 | rep6 | 21 | 23 |
| PSD95 | rep7 | 21 | 21 |
| PSD95 | rep8 | 22 | 19 |
| P54 | rep1 | 14 | 25 |
| P54 | rep2 | 25 | 25 |
| P54 | rep3 | 25 | 25 |
| P54 | rep4 | 25 | 24 |
| P54 | rep5 | 25 | 25 |
| P54 | rep6 | 24 | 24 |
| P54 | rep7 | 25 | 25 |
| P54 | rep8 | 25 | 23 |
| TDP43 | rep1 | 5 | 25 |
| TDP43 | rep2 | 25 | 25 |
| TDP43 | rep3 | 25 | 25 |
| TDP43 | rep4 | 15 | 24 |
| TDP43 | rep5 | 25 | 25 |
| TDP43 | rep6 | 25 | 20 |
| TDP43 | rep7 | 25 | 25 |
| TDP43 | rep8 | 25 | 25 |
| GM130 | rep1 | 24 | 12 |
| GM130 | rep2 | 21 | 25 |
| GM130 | rep3 | 23 | 25 |
| GM130 | rep4 | 24 | 25 |
| GM130 | rep5 | 24 | 21 |
| GM130 | rep6 | 23 | 25 |
| GM130 | rep7 | 25 | 24 |
| GM130 | rep8 | 25 | 25 |
| Phalloidin | rep1 | 23 | 19 |
| Phalloidin | rep2 | 18 | 23 |
| Phalloidin | rep3 | 24 | 22 |
| Phalloidin | rep4 | 21 | 22 |
| Phalloidin | rep5 | 24 | 25 |
| Phalloidin | rep6 | 24 | 22 |
| Phalloidin | rep7 | 25 | 24 |
| Phalloidin | rep8 | 24 | 23 |
| TOMM20 | rep1 | 24 | 21 |
| TOMM20 | rep2 | 23 | 25 |
| TOMM20 | rep3 | 22 | 25 |
| TOMM20 | rep4 | 15 | 25 |
| TOMM20 | rep5 | 23 | 25 |
| TOMM20 | rep6 | 18 | 25 |
| TOMM20 | rep7 | 25 | 24 |
| TOMM20 | rep8 | 25 | 19 |
| Calreticulin | rep1 | 25 | 25 |
| Calreticulin | rep2 | 25 | 25 |
| Calreticulin | rep3 | 25 | 25 |
| Calreticulin | rep4 | 25 | 25 |
| Calreticulin | rep5 | 25 | 24 |
| Calreticulin | rep6 | 25 | 25 |
| Calreticulin | rep7 | 25 | 24 |
| Calreticulin | rep8 | 25 | 25 |

| | Rep | WT stress | WT Untreated |
|---|---|---|---|
| LAMP1 | rep1 | 25 | 25 |
| LAMP1 | rep2 | 24 | 25 |
| LAMP1 | rep3 | 25 | 25 |
| LAMP1 | rep4 | 25 | 25 |
| LAMP1 | rep5 | 25 | 25 |
| LAMP1 | rep6 | 25 | 24 |
| LAMP1 | rep7 | 25 | 25 |
| LAMP1 | rep8 | 25 | 25 |
| FMRP | rep1 | 25 | 25 |
| FMRP | rep2 | 25 | 25 |
| FMRP | rep3 | 23 | 23 |
| FMRP | rep4 | 25 | 25 |
| FMRP | rep5 | 23 | 25 |
| FMRP | rep6 | 23 | 22 |
| FMRP | rep7 | 24 | 25 |
| FMRP | rep8 | 24 | 24 |
| SQSTM1 | rep1 | 17 | 19 |
| SQSTM1 | rep2 | 25 | 25 |
| SQSTM1 | rep3 | 21 | 20 |
| SQSTM1 | rep4 | 23 | 21 |
| SQSTM1 | rep5 | 17 | 25 |
| SQSTM1 | rep6 | 20 | 24 |
| SQSTM1 | rep7 | 19 | 21 |
| SQSTM1 | rep8 | 20 | 18 |
| G3BP1 | rep1 | 25 | 25 |
| G3BP1 | rep2 | 25 | 25 |
| G3BP1 | rep3 | 24 | 25 |
| G3BP1 | rep4 | 25 | 25 |
| G3BP1 | rep5 | 25 | 25 |
| G3BP1 | rep6 | 24 | 25 |
| G3BP1 | rep7 | 25 | 25 |
| G3BP1 | rep8 | 25 | 25 |
| KIF5A | rep1 | 25 | 25 |
| KIF5A | rep2 | 25 | 24 |
| KIF5A | rep3 | 24 | 21 |
| KIF5A | rep4 | 25 | 24 |
| KIF5A | rep5 | 15 | 25 |
| KIF5A | rep6 | 24 | 25 |
| KIF5A | rep7 | 25 | 19 |
| KIF5A | rep8 | 25 | 21 |
| PEX14 | rep1 | 24 | 25 |
| PEX14 | rep2 | 25 | 25 |
| PEX14 | rep3 | 25 | 24 |
| PEX14 | rep4 | 24 | 25 |
| PEX14 | rep5 | 25 | 25 |
| PEX14 | rep6 | 25 | 23 |
| PEX14 | rep7 | 24 | 24 |
| PEX14 | rep8 | 25 | 25 |
| PML | rep1 | 24 | 25 |
| PML | rep2 | 22 | 23 |
| PML | rep3 | 25 | 24 |
| PML | rep4 | 25 | 21 |
| PML | rep5 | 20 | 20 |
| PML | rep6 | 25 | 22 |
| PML | rep7 | 24 | 20 |
| PML | rep8 | 24 | 24 |
| PURA | rep1 | 24 | 25 |
| PURA | rep2 | 25 | 25 |
| PURA | rep3 | 25 | 25 |
| PURA | rep4 | 25 | 25 |
| PURA | rep5 | 25 | 23 |
| PURA | rep6 | 25 | 25 |
| PURA | rep7 | 25 | 25 |
| PURA | rep8 | 25 | 25 |
| TIA1 | rep1 | 18 | 25 |
| TIA1 | rep2 | 25 | 25 |
| TIA1 | rep3 | 25 | 25 |
| TIA1 | rep4 | 25 | 25 |
| TIA1 | rep5 | 25 | 23 |
| TIA1 | rep6 | 25 | 25 |
| TIA1 | rep7 | 25 | 25 |
| TIA1 | rep8 | 25 | 25 |
| FUS | rep1 | 24 | 25 |
| FUS | rep2 | 25 | 25 |
| FUS | rep3 | 25 | 25 |
| FUS | rep4 | 25 | 25 |
| FUS | rep5 | 24 | 25 |
| FUS | rep6 | 24 | 25 |
| FUS | rep7 | 25 | 25 |
| FUS | rep8 | 25 | 25 |
| MitoTracker | rep1 | 24 | 25 |
| MitoTracker | rep2 | 25 | 25 |
| MitoTracker | rep3 | 25 | 25 |
| MitoTracker | rep4 | 25 | 25 |
| MitoTracker | rep5 | 24 | 25 |
| MitoTracker | rep6 | 25 | 25 |
| MitoTracker | rep7 | 25 | 25 |
| MitoTracker | rep8 | 25 | 25 |
| NCL | rep1 | 24 | 25 |
| NCL | rep2 | 25 | 25 |
| NCL | rep3 | 25 | 25 |
| NCL | rep4 | 25 | 25 |
| NCL | rep5 | 24 | 25 |
| NCL | rep6 | 24 | 25 |
| NCL | rep7 | 25 | 25 |
| NCL | rep8 | 25 | 25 |
| DAPI | rep1 | 264 | 271 |
| DAPI | rep2 | 274 | 275 |
| DAPI | rep3 | 273 | 275 |
| DAPI | rep4 | 275 | 275 |
| DAPI | rep5 | 272 | 273 |
| DAPI | rep6 | 273 | 273 |
| DAPI | rep7 | 274 | 274 |
| DAPI | rep8 | 275 | 275 |

### Batch 2

| | Rep | WT stress | WT Untreated |
|---|---|---|---|
| ANAX11 | rep1 | 23 | 25 |
| ANAX11 | rep2 | 24 | 22 |
| ANAX11 | rep3 | 25 | 25 |
| ANAX11 | rep4 | 25 | 25 |
| ANAX11 | rep5 | 25 | 25 |
| ANAX11 | rep6 | 25 | 24 |
| ANAX11 | rep7 | 24 | 24 |
| ANAX11 | rep8 | 25 | 25 |
| SNCA | rep1 | 19 | 25 |
| SNCA | rep2 | 24 | 17 |
| SNCA | rep3 | 20 | 24 |
| SNCA | rep4 | 24 | 22 |
| SNCA | rep5 | 25 | 23 |
| SNCA | rep6 | 25 | 18 |
| SNCA | rep7 | 18 | 22 |
| SNCA | rep8 | 25 | 25 |
| TUJ1 | rep1 | 198 | 200 |
| TUJ1 | rep2 | 195 | 199 |
| TUJ1 | rep3 | 198 | 200 |
| TUJ1 | rep4 | 198 | 200 |
| TUJ1 | rep5 | 200 | 200 |
| TUJ1 | rep6 | 198 | 200 |
| TUJ1 | rep7 | 198 | 200 |
| TUJ1 | rep8 | 198 | 200 |
| DCP1A | rep1 | 25 | 22 |
| DCP1A | rep2 | 25 | 25 |
| DCP1A | rep3 | 23 | 24 |
| DCP1A | rep4 | 24 | 23 |
| DCP1A | rep5 | 25 | 25 |
| DCP1A | rep6 | 24 | 23 |
| DCP1A | rep7 | 25 | 25 |
| DCP1A | rep8 | 24 | 25 |
| NEMO | rep1 | 25 | 25 |
| NEMO | rep2 | 24 | 22 |
| NEMO | rep3 | 25 | 25 |
| NEMO | rep4 | 24 | 25 |
| NEMO | rep5 | 25 | 25 |
| NEMO | rep6 | 25 | 25 |
| NEMO | rep7 | 25 | 25 |
| NEMO | rep8 | 24 | 25 |
| CLTC | rep1 | 24 | 24 |
| CLTC | rep2 | 22 | 25 |
| CLTC | rep3 | 23 | 25 |
| CLTC | rep4 | 25 | 25 |
| CLTC | rep5 | 25 | 25 |
| CLTC | rep6 | 24 | 25 |
| CLTC | rep7 | 24 | 25 |
| CLTC | rep8 | 25 | 25 |
| PSD95 | rep1 | 12 | 24 |
| PSD95 | rep2 | 22 | 12 |
| PSD95 | rep3 | 15 | 23 |
| PSD95 | rep4 | 24 | 19 |
| PSD95 | rep5 | 11 | 11 |
| PSD95 | rep6 | 20 | 16 |
| PSD95 | rep7 | 20 | 13 |
| PSD95 | rep8 | 21 | 12 |
| P54 | rep1 | 19 | 15 |
| P54 | rep2 | 23 | 25 |
| P54 | rep3 | 25 | 25 |
| P54 | rep4 | 23 | 25 |
| P54 | rep5 | 24 | 25 |
| P54 | rep6 | 20 | 25 |
| P54 | rep7 | 24 | 24 |
| P54 | rep8 | 17 | 24 |
| TDP43 | rep1 | 19 | 21 |
| TDP43 | rep2 | 25 | 25 |
| TDP43 | rep3 | 25 | 25 |
| TDP43 | rep4 | 25 | 25 |
| TDP43 | rep5 | 25 | 25 |
| TDP43 | rep6 | 25 | 25 |
| TDP43 | rep7 | 25 | 24 |
| TDP43 | rep8 | 25 | 24 |
| GM130 | rep1 | 25 | 25 |
| GM130 | rep2 | 24 | 25 |
| GM130 | rep3 | 22 | 24 |
| GM130 | rep4 | 25 | 25 |
| GM130 | rep5 | 24 | 25 |
| GM130 | rep6 | 25 | 25 |
| GM130 | rep7 | 23 | 25 |
| GM130 | rep8 | 24 | 25 |
| Phalloidin | rep1 | 24 | 23 |
| Phalloidin | rep2 | 23 | 24 |
| Phalloidin | rep3 | 24 | 24 |
| Phalloidin | rep4 | 24 | 24 |
| Phalloidin | rep5 | 23 | 25 |
| Phalloidin | rep6 | 25 | 24 |
| Phalloidin | rep7 | 23 | 25 |
| Phalloidin | rep8 | 24 | 24 |
| TOMM20 | rep1 | 8 | 25 |
| TOMM20 | rep2 | 18 | 25 |
| TOMM20 | rep3 | 1 | 25 |
| TOMM20 | rep4 | 0 | 25 |
| TOMM20 | rep5 | 12 | 25 |
| TOMM20 | rep6 | 23 | 25 |
| TOMM20 | rep7 | 15 | 25 |
| TOMM20 | rep8 | 7 | 25 |
| Calreticulin | rep1 | 2 | 22 |
| Calreticulin | rep2 | 8 | 25 |
| Calreticulin | rep3 | 14 | 17 |
| Calreticulin | rep4 | 5 | 25 |
| Calreticulin | rep5 | 25 | 25 |
| Calreticulin | rep6 | 9 | 25 |
| Calreticulin | rep7 | 15 | 21 |
| Calreticulin | rep8 | 22 | 25 |

| | Rep | WT stress | WT Untreated |
|---|---|---|---|
| LAMP1 | rep1 | 23 | 25 |
| LAMP1 | rep2 | 23 | 25 |
| LAMP1 | rep3 | 24 | 25 |
| LAMP1 | rep4 | 17 | 25 |
| LAMP1 | rep5 | 21 | 25 |
| LAMP1 | rep6 | 22 | 25 |
| LAMP1 | rep7 | 25 | 25 |
| LAMP1 | rep8 | 25 | 24 |
| FMRP | rep1 | 23 | 25 |
| FMRP | rep2 | 25 | 22 |
| FMRP | rep3 | 21 | 24 |
| FMRP | rep4 | 23 | 23 |
| FMRP | rep5 | 25 | 23 |
| FMRP | rep6 | 25 | 24 |
| FMRP | rep7 | 21 | 24 |
| FMRP | rep8 | 23 | 23 |
| SQSTM1 | rep1 | 9 | 19 |
| SQSTM1 | rep2 | 23 | 13 |
| SQSTM1 | rep3 | 13 | 15 |
| SQSTM1 | rep4 | 22 | 15 |
| SQSTM1 | rep5 | 20 | 17 |
| SQSTM1 | rep6 | 23 | 16 |
| SQSTM1 | rep7 | 9 | 9 |
| SQSTM1 | rep8 | 12 | 16 |
| G3BP1 | rep1 | 23 | 25 |
| G3BP1 | rep2 | 25 | 25 |
| G3BP1 | rep3 | 25 | 25 |
| G3BP1 | rep4 | 25 | 25 |
| G3BP1 | rep5 | 25 | 25 |
| G3BP1 | rep6 | 25 | 25 |
| G3BP1 | rep7 | 25 | 25 |
| G3BP1 | rep8 | 25 | 25 |
| KIF5A | rep1 | 11 | 24 |
| KIF5A | rep2 | 24 | 17 |
| KIF5A | rep3 | 24 | 21 |
| KIF5A | rep4 | 25 | 22 |
| KIF5A | rep5 | 25 | 25 |
| KIF5A | rep6 | 25 | 23 |
| KIF5A | rep7 | 17 | 25 |
| KIF5A | rep8 | 24 | 25 |
| PEX14 | rep1 | 25 | 25 |
| PEX14 | rep2 | 25 | 25 |
| PEX14 | rep3 | 25 | 25 |
| PEX14 | rep4 | 25 | 25 |
| PEX14 | rep5 | 24 | 25 |
| PEX14 | rep6 | 25 | 25 |
| PEX14 | rep7 | 24 | 25 |
| PEX14 | rep8 | 24 | 25 |
| PML | rep1 | 24 | 22 |
| PML | rep2 | 25 | 22 |
| PML | rep3 | 23 | 20 |
| PML | rep4 | 24 | 24 |
| PML | rep5 | 23 | 22 |
| PML | rep6 | 25 | 19 |
| PML | rep7 | 22 | 22 |
| PML | rep8 | 21 | 23 |
| PURA | rep1 | 25 | 25 |
| PURA | rep2 | 24 | 25 |
| PURA | rep3 | 25 | 25 |
| PURA | rep4 | 24 | 25 |
| PURA | rep5 | 24 | 25 |
| PURA | rep6 | 25 | 25 |
| PURA | rep7 | 25 | 25 |
| PURA | rep8 | 25 | 25 |
| TIA1 | rep1 | 24 | 25 |
| TIA1 | rep2 | 24 | 25 |
| TIA1 | rep3 | 25 | 25 |
| TIA1 | rep4 | 24 | 25 |
| TIA1 | rep5 | 25 | 23 |
| TIA1 | rep6 | 25 | 25 |
| TIA1 | rep7 | 25 | 25 |
| TIA1 | rep8 | 25 | 25 |
| FUS | rep1 | 25 | 25 |
| FUS | rep2 | 25 | 25 |
| FUS | rep3 | 24 | 25 |
| FUS | rep4 | 25 | 25 |
| FUS | rep5 | 25 | 25 |
| FUS | rep6 | 24 | 25 |
| FUS | rep7 | 24 | 25 |
| FUS | rep8 | 24 | 25 |
| MitoTracker | rep1 | 25 | 25 |
| MitoTracker | rep2 | 25 | 25 |
| MitoTracker | rep3 | 23 | 25 |
| MitoTracker | rep4 | 25 | 25 |
| MitoTracker | rep5 | 20 | 25 |
| MitoTracker | rep6 | 23 | 25 |
| MitoTracker | rep7 | 24 | 25 |
| MitoTracker | rep8 | 25 | 25 |
| NCL | rep1 | 25 | 25 |
| NCL | rep2 | 25 | 25 |
| NCL | rep3 | 24 | 25 |
| NCL | rep4 | 24 | 25 |
| NCL | rep5 | 25 | 25 |
| NCL | rep6 | 25 | 25 |
| NCL | rep7 | 24 | 25 |
| NCL | rep8 | 25 | 25 |
| DAPI | rep1 | 268 | 273 |
| DAPI | rep2 | 272 | 274 |
| DAPI | rep3 | 273 | 275 |
| DAPI | rep4 | 274 | 275 |
| DAPI | rep5 | 274 | 275 |
| DAPI | rep6 | 275 | 275 |
| DAPI | rep7 | 270 | 275 |
| DAPI | rep8 | 272 | 274 |

### Batch 3

| | Rep | WT stress | WT Untreated |
|---|---|---|---|
| ANAX11 | rep1 | 23 | 24 |
| ANAX11 | rep2 | 24 | 22 |
| ANAX11 | rep3 | 24 | 25 |
| ANAX11 | rep4 | 25 | 24 |
| ANAX11 | rep5 | 23 | 24 |
| ANAX11 | rep6 | 22 | 21 |
| ANAX11 | rep7 | 25 | 25 |
| ANAX11 | rep8 | 24 | 23 |
| SNCA | rep1 | 25 | 25 |
| SNCA | rep2 | 25 | 22 |
| SNCA | rep3 | 25 | 25 |
| SNCA | rep4 | 25 | 25 |
| SNCA | rep5 | 25 | 24 |
| SNCA | rep6 | 24 | 24 |
| SNCA | rep7 | 24 | 25 |
| SNCA | rep8 | 25 | 21 |
| TUJ1 | rep1 | 184 | 200 |
| TUJ1 | rep2 | 198 | 191 |
| TUJ1 | rep3 | 190 | 199 |
| TUJ1 | rep4 | 197 | 191 |
| TUJ1 | rep5 | 188 | 200 |
| TUJ1 | rep6 | 189 | 196 |
| TUJ1 | rep7 | 199 | 199 |
| TUJ1 | rep8 | 200 | 197 |
| DCP1A | rep1 | 23 | 24 |
| DCP1A | rep2 | 24 | 24 |
| DCP1A | rep3 | 24 | 21 |
| DCP1A | rep4 | 24 | 20 |
| DCP1A | rep5 | 23 | 19 |
| DCP1A | rep6 | 24 | 23 |
| DCP1A | rep7 | 24 | 24 |
| DCP1A | rep8 | 25 | 23 |
| NEMO | rep1 | 23 | 25 |
| NEMO | rep2 | 25 | 24 |
| NEMO | rep3 | 24 | 23 |
| NEMO | rep4 | 22 | 25 |
| NEMO | rep5 | 24 | 25 |
| NEMO | rep6 | 24 | 25 |
| NEMO | rep7 | 23 | 25 |
| NEMO | rep8 | 24 | 25 |
| CLTC | rep1 | 24 | 25 |
| CLTC | rep2 | 25 | 25 |
| CLTC | rep3 | 22 | 25 |
| CLTC | rep4 | 25 | 22 |
| CLTC | rep5 | 25 | 25 |
| CLTC | rep6 | 25 | 23 |
| CLTC | rep7 | 25 | 25 |
| CLTC | rep8 | 25 | 25 |
| PSD95 | rep1 | 24 | 25 |
| PSD95 | rep2 | 22 | 25 |
| PSD95 | rep3 | 15 | 23 |
| PSD95 | rep4 | 24 | 20 |
| PSD95 | rep5 | 20 | 25 |
| PSD95 | rep6 | 25 | 16 |
| PSD95 | rep7 | 22 | 20 |
| PSD95 | rep8 | 24 | 25 |
| P54 | rep1 | 25 | 24 |
| P54 | rep2 | 25 | 24 |
| P54 | rep3 | 23 | 24 |
| P54 | rep4 | 25 | 25 |
| P54 | rep5 | 15 | 22 |
| P54 | rep6 | 17 | 15 |
| P54 | rep7 | 25 | 25 |
| P54 | rep8 | 25 | 25 |
| TDP43 | rep1 | 25 | 24 |
| TDP43 | rep2 | 24 | 24 |
| TDP43 | rep3 | 25 | 24 |
| TDP43 | rep4 | 25 | 25 |
| TDP43 | rep5 | 21 | 25 |
| TDP43 | rep6 | 24 | 24 |
| TDP43 | rep7 | 25 | 25 |
| TDP43 | rep8 | 21 | 25 |
| GM130 | rep1 | 24 | 25 |
| GM130 | rep2 | 25 | 23 |
| GM130 | rep3 | 24 | 24 |
| GM130 | rep4 | 24 | 24 |
| GM130 | rep5 | 25 | 25 |
| GM130 | rep6 | 25 | 25 |
| GM130 | rep7 | 25 | 24 |
| GM130 | rep8 | 20 | 25 |
| Phalloidin | rep1 | 24 | 25 |
| Phalloidin | rep2 | 25 | 25 |
| Phalloidin | rep3 | 22 | 24 |
| Phalloidin | rep4 | 25 | 25 |
| Phalloidin | rep5 | 25 | 24 |
| Phalloidin | rep6 | 25 | 25 |
| Phalloidin | rep7 | 25 | 25 |
| Phalloidin | rep8 | 25 | 23 |
| TOMM20 | rep1 | 4 | 24 |
| TOMM20 | rep2 | 8 | 25 |
| TOMM20 | rep3 | 1 | 25 |
| TOMM20 | rep4 | 0 | 25 |
| TOMM20 | rep5 | 9 | 25 |
| TOMM20 | rep6 | 3 | 25 |
| TOMM20 | rep7 | 7 | 25 |
| TOMM20 | rep8 | 5 | 25 |
| Calreticulin | rep1 | 22 | 25 |
| Calreticulin | rep2 | 23 | 24 |
| Calreticulin | rep3 | 24 | 25 |
| Calreticulin | rep4 | 23 | 24 |
| Calreticulin | rep5 | 24 | 25 |
| Calreticulin | rep6 | 14 | 25 |
| Calreticulin | rep7 | 24 | 25 |
| Calreticulin | rep8 | 24 | 25 |

| | Rep | WT stress | WT Untreated |
|---|---|---|---|
| LAMP1 | rep1 | 18 | 25 |
| LAMP1 | rep2 | 23 | 24 |
| LAMP1 | rep3 | 19 | 25 |
| LAMP1 | rep4 | 22 | 25 |
| LAMP1 | rep5 | 23 | 25 |
| LAMP1 | rep6 | 20 | 25 |
| LAMP1 | rep7 | 21 | 25 |
| LAMP1 | rep8 | 25 | 25 |
| FMRP | rep1 | 22 | 19 |
| FMRP | rep2 | 23 | 19 |
| FMRP | rep3 | 22 | 22 |
| FMRP | rep4 | 23 | 22 |
| FMRP | rep5 | 21 | 25 |
| FMRP | rep6 | 21 | 22 |
| FMRP | rep7 | 16 | 22 |
| FMRP | rep8 | 24 | 24 |
| SQSTM1 | rep1 | 20 | 19 |
| SQSTM1 | rep2 | 23 | 20 |
| SQSTM1 | rep3 | 23 | 21 |
| SQSTM1 | rep4 | 24 | 21 |
| SQSTM1 | rep5 | 20 | 25 |
| SQSTM1 | rep6 | 20 | 19 |
| SQSTM1 | rep7 | 16 | 22 |
| SQSTM1 | rep8 | 23 | 24 |
| G3BP1 | rep1 | 23 | 25 |
| G3BP1 | rep2 | 25 | 25 |
| G3BP1 | rep3 | 25 | 25 |
| G3BP1 | rep4 | 25 | 25 |
| G3BP1 | rep5 | 25 | 25 |
| G3BP1 | rep6 | 25 | 24 |
| G3BP1 | rep7 | 25 | 25 |
| G3BP1 | rep8 | 25 | 25 |
| KIF5A | rep1 | 23 | 25 |
| KIF5A | rep2 | 25 | 25 |
| KIF5A | rep3 | 25 | 24 |
| KIF5A | rep4 | 25 | 25 |
| KIF5A | rep5 | 25 | 25 |
| KIF5A | rep6 | 25 | 24 |
| KIF5A | rep7 | 25 | 25 |
| KIF5A | rep8 | 25 | 23 |
| PEX14 | rep1 | 17 | 24 |
| PEX14 | rep2 | 25 | 16 |
| PEX14 | rep3 | 18 | 23 |
| PEX14 | rep4 | 23 | 19 |
| PEX14 | rep5 | 25 | 25 |
| PEX14 | rep6 | 25 | 25 |
| PEX14 | rep7 | 25 | 24 |
| PEX14 | rep8 | 25 | 25 |
| PML | rep1 | 18 | 17 |
| PML | rep2 | 25 | 12 |
| PML | rep3 | 15 | 19 |
| PML | rep4 | 21 | 18 |
| PML | rep5 | 21 | 24 |
| PML | rep6 | 23 | 24 |
| PML | rep7 | 23 | 23 |
| PML | rep8 | 24 | 24 |
| PURA | rep1 | 23 | 25 |
| PURA | rep2 | 25 | 23 |
| PURA | rep3 | 25 | 25 |
| PURA | rep4 | 24 | 23 |
| PURA | rep5 | 19 | 24 |
| PURA | rep6 | 24 | 23 |
| PURA | rep7 | 24 | 25 |
| PURA | rep8 | 25 | 24 |
| TIA1 | rep1 | 24 | 25 |
| TIA1 | rep2 | 24 | 23 |
| TIA1 | rep3 | 25 | 23 |
| TIA1 | rep4 | 24 | 23 |
| TIA1 | rep5 | 14 | 25 |
| TIA1 | rep6 | 20 | 24 |
| TIA1 | rep7 | 25 | 25 |
| TIA1 | rep8 | 25 | 24 |
| FUS | rep1 | 24 | 25 |
| FUS | rep2 | 25 | 25 |
| FUS | rep3 | 24 | 25 |
| FUS | rep4 | 25 | 25 |
| FUS | rep5 | 22 | 25 |
| FUS | rep6 | 25 | 25 |
| FUS | rep7 | 25 | 25 |
| FUS | rep8 | 25 | 25 |
| MitoTracker | rep1 | 24 | 25 |
| MitoTracker | rep2 | 25 | 25 |
| MitoTracker | rep3 | 24 | 25 |
| MitoTracker | rep4 | 25 | 25 |
| MitoTracker | rep5 | 22 | 25 |
| MitoTracker | rep6 | 25 | 25 |
| MitoTracker | rep7 | 24 | 25 |
| MitoTracker | rep8 | 25 | 25 |
| NCL | rep1 | 24 | 25 |
| NCL | rep2 | 25 | 25 |
| NCL | rep3 | 24 | 25 |
| NCL | rep4 | 25 | 25 |
| NCL | rep5 | 22 | 25 |
| NCL | rep6 | 25 | 25 |
| NCL | rep7 | 25 | 25 |
| NCL | rep8 | 25 | 25 |
| DAPI | rep1 | 260 | 274 |
| DAPI | rep2 | 273 | 265 |
| DAPI | rep3 | 263 | 273 |
| DAPI | rep4 | 273 | 267 |
| DAPI | rep5 | 261 | 275 |
| DAPI | rep6 | 269 | 270 |
| DAPI | rep7 | 274 | 274 |
| DAPI | rep8 | 275 | 272 |

#### **Total Tile Count**

#### Batch1

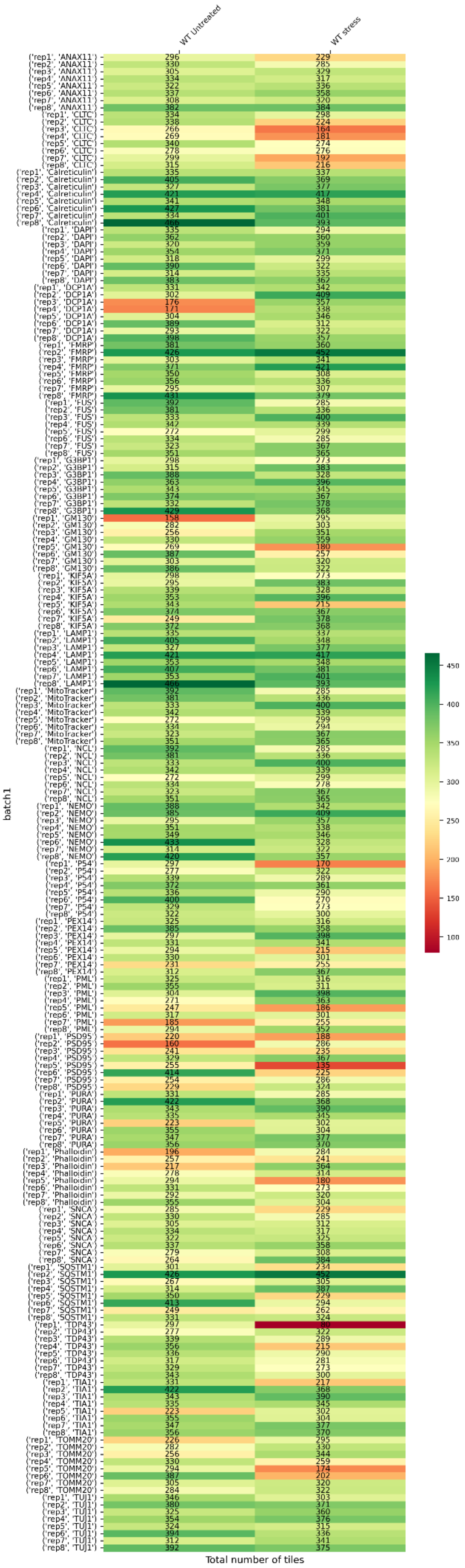

### Batch 2

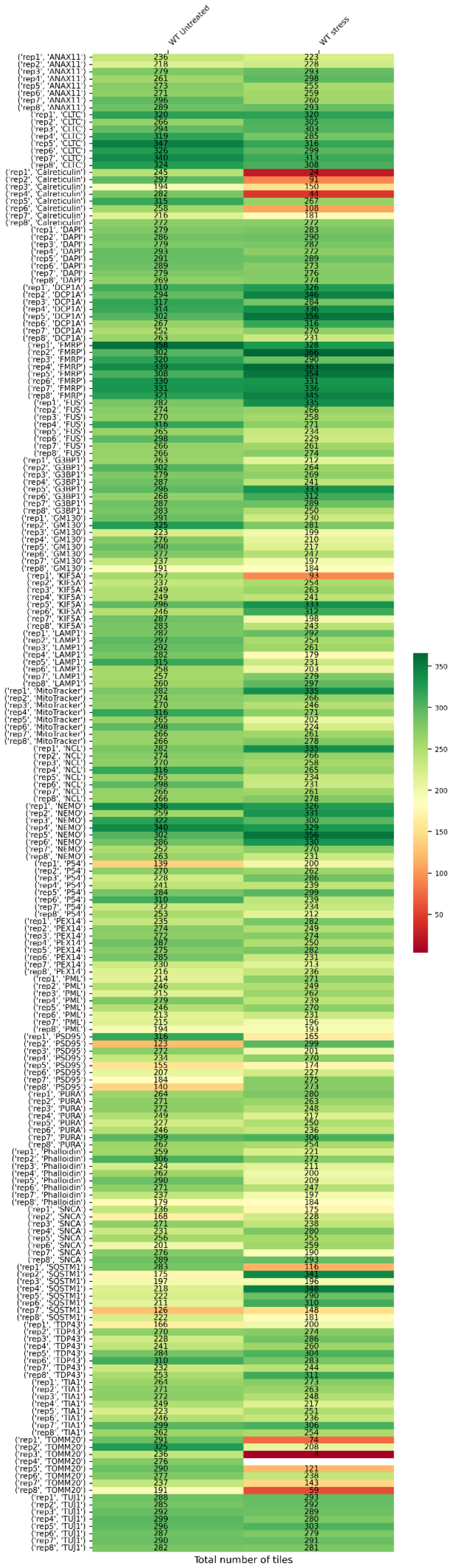

WT Untreated
WT stress
batch2
Total number of tiles
350
300
250
200
150
100
50

### Batch 3

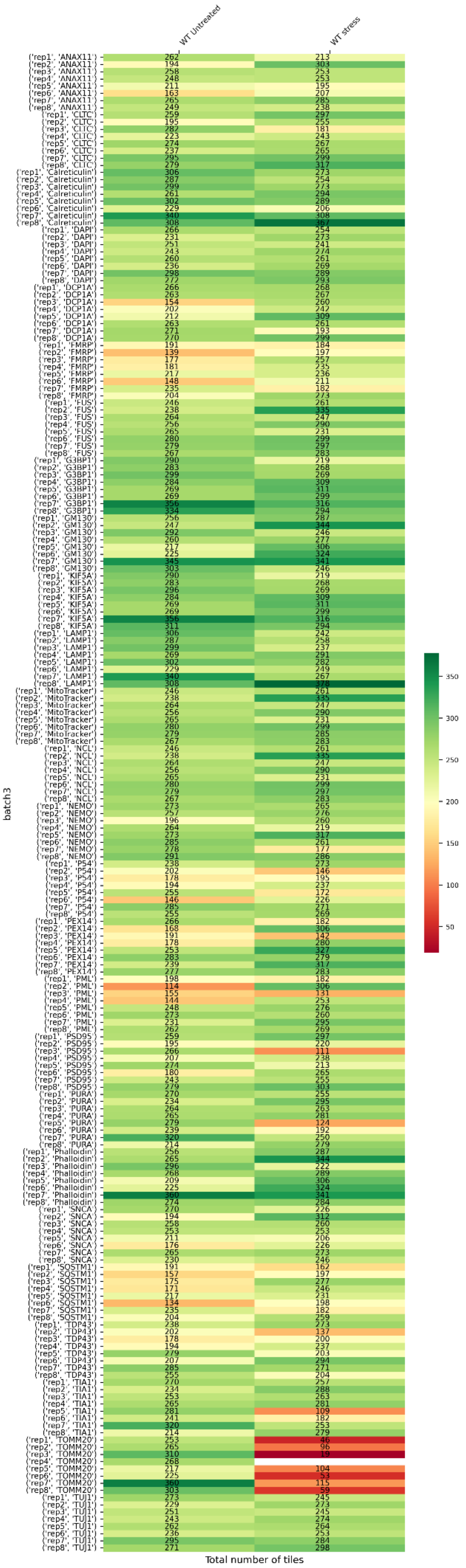

WT Untreated
WT stress
batch3
Total number of tiles
350
300
250
200
150
100
50

### Whole Cell Count

#### Batch 1

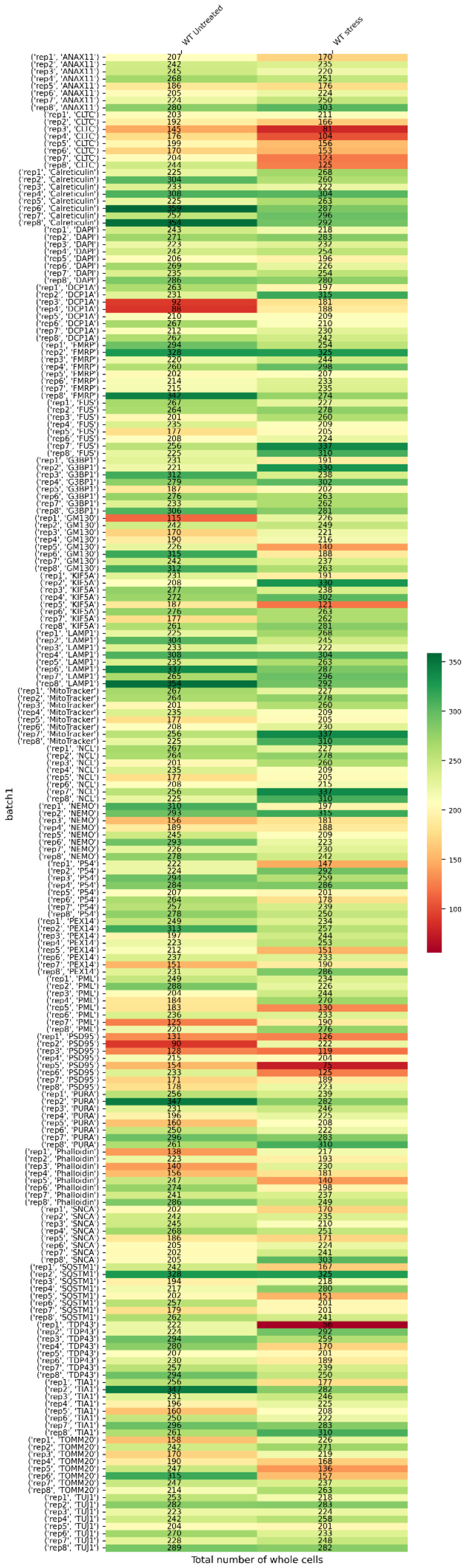

### Batch 2

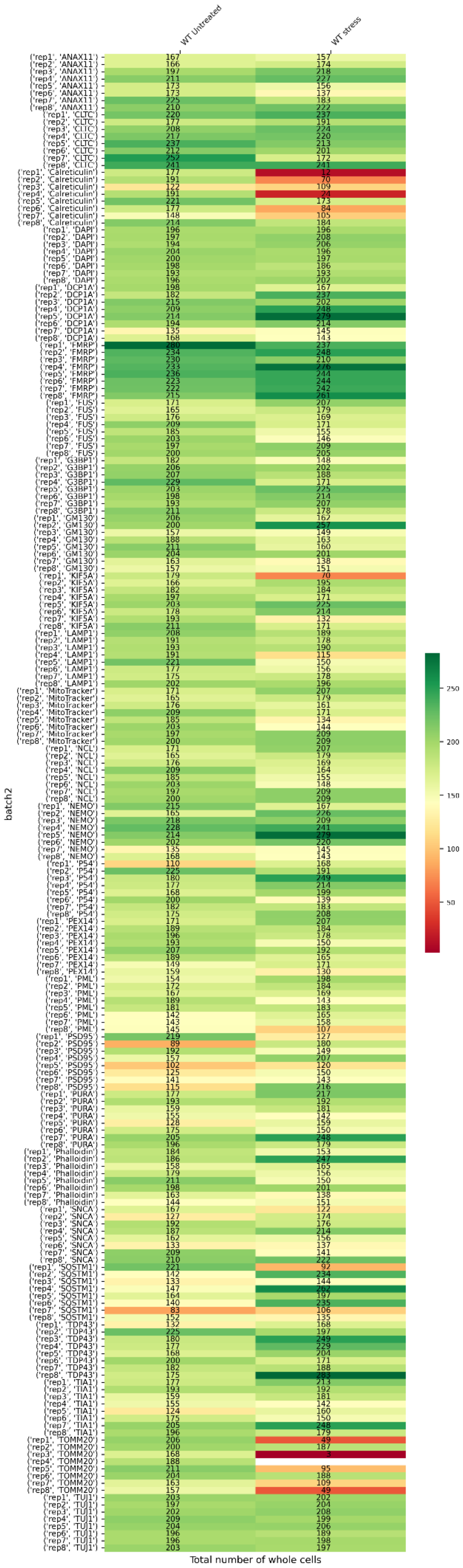
WT Untreated
WT stress
batch2
('rep1', 'ANAX11') 167 157
('rep2', 'ANAX11') 166 174
('rep3', 'ANAX11') 197 218
('rep4', 'ANAX11') 211 227
('rep5', 'ANAX11') 173 156
('rep6', 'ANAX11') 173 137
('rep7', 'ANAX11') 225 183
('rep8', 'ANAX11') 210 222
('rep1', 'CLTC') 220 237
('rep2', 'CLTC') 177 191
('rep3', 'CLTC') 208 224
('rep4', 'CLTC') 217 220
('rep5', 'CLTC') 237 213
('rep6', 'CLTC') 212 201
('rep7', 'CLTC') 252 172
('rep8', 'CLTC') 241 241
('rep1', 'Calreticulin') 177 12
('rep2', 'Calreticulin') 191 70
('rep3', 'Calreticulin') 122 109
('rep4', 'Calreticulin') 191 24
('rep5', 'Calreticulin') 221 173
('rep6', 'Calreticulin') 177 84
('rep7', 'Calreticulin') 148 105
('rep8', 'Calreticulin') 214 184
('rep1', 'DAPI') 196 196
('rep2', 'DAPI') 197 208
('rep3', 'DAPI') 194 206
('rep4', 'DAPI') 204 196
('rep5', 'DAPI') 200 197
('rep6', 'DAPI') 198 186
('rep7', 'DAPI') 193 193
('rep8', 'DAPI') 196 202
('rep1', 'DCP1A') 198 167
('rep2', 'DCP1A') 182 237
('rep3', 'DCP1A') 215 202
('rep4', 'DCP1A') 209 248
('rep5', 'DCP1A') 214 279
('rep6', 'DCP1A') 194 214
('rep7', 'DCP1A') 135 145
('rep8', 'DCP1A') 168 143
('rep1', 'FMRP') 280 237
('rep2', 'FMRP') 234 248
('rep3', 'FMRP') 230 210
('rep4', 'FMRP') 233 276
('rep5', 'FMRP') 236 244
('rep6', 'FMRP') 223 244
('rep7', 'FMRP') 222 242
('rep8', 'FMRP') 215 261
('rep1', 'FUS') 171 207
('rep2', 'FUS') 165 179
('rep3', 'FUS') 176 169
('rep4', 'FUS') 209 171
('rep5', 'FUS') 185 155
('rep6', 'FUS') 203 146
('rep7', 'FUS') 197 209
('rep8', 'FUS') 200 205
('rep1', 'G3BP1') 182 148
('rep2', 'G3BP1') 206 202
('rep3', 'G3BP1') 207 188
('rep4', 'G3BP1') 229 171
('rep5', 'G3BP1') 203 225
('rep6', 'G3BP1') 198 214
('rep7', 'G3BP1') 193 207
('rep8', 'G3BP1') 211 178
('rep1', 'GM130') 206 162
('rep2', 'GM130') 200 257
('rep3', 'GM130') 157 149
('rep4', 'GM130') 188 163
('rep5', 'GM130') 211 160
('rep6', 'GM130') 204 201
('rep7', 'GM130') 163 138
('rep8', 'GM130') 157 151
('rep1', 'KIF5A') 179 70
('rep2', 'KIF5A') 166 195
('rep3', 'KIF5A') 182 184
('rep4', 'KIF5A') 197 171
('rep5', 'KIF5A') 203 225
('rep6', 'KIF5A') 178 214
('rep7', 'KIF5A') 193 132
('rep8', 'KIF5A') 211 171
('rep1', 'LAMP1') 208 189
('rep2', 'LAMP1') 191 178
('rep3', 'LAMP1') 193 190
('rep4', 'LAMP1') 191 115
('rep5', 'LAMP1') 221 150
('rep6', 'LAMP1') 177 156
('rep7', 'LAMP1') 175 178
('rep8', 'LAMP1') 202 196
('rep1', 'MitoTracker') 171 207
('rep2', 'MitoTracker') 165 179
('rep3', 'MitoTracker') 176 161
('rep4', 'MitoTracker') 209 171
('rep5', 'MitoTracker') 185 134
('rep6', 'MitoTracker') 203 144
('rep7', 'MitoTracker') 197 209
('rep8', 'MitoTracker') 200 209
('rep1', 'NCL') 171 207
('rep2', 'NCL') 165 179
('rep3', 'NCL') 176 169
('rep4', 'NCL') 209 164
('rep5', 'NCL') 185 155
('rep6', 'NCL') 203 148
('rep7', 'NCL') 197 209
('rep8', 'NCL') 200 209
('rep1', 'NEMO') 215 167
('rep2', 'NEMO') 165 226
('rep3', 'NEMO') 218 209
('rep4', 'NEMO') 228 241
('rep5', 'NEMO') 214 279
('rep6', 'NEMO') 202 220
('rep7', 'NEMO') 135 145
('rep8', 'NEMO') 168 143
('rep1', 'P54') 110 168
('rep2', 'P54') 225 191
('rep3', 'P54') 180 249
('rep4', 'P54') 177 214
('rep5', 'P54') 168 199
('rep6', 'P54') 200 139
('rep7', 'P54') 182 183
('rep8', 'P54') 175 208
('rep1', 'PEX14') 171 207
('rep2', 'PEX14') 189 184
('rep3', 'PEX14') 196 178
('rep4', 'PEX14') 193 150
('rep5', 'PEX14') 207 192
('rep6', 'PEX14') 189 165
('rep7', 'PEX14') 149 171
('rep8', 'PEX14') 159 130
('rep1', 'PML') 154 198
('rep2', 'PML') 172 184
('rep3', 'PML') 167 169
('rep4', 'PML') 189 143
('rep5', 'PML') 181 183
('rep6', 'PML') 142 165
('rep7', 'PML') 143 158
('rep8', 'PML') 145 107
('rep1', 'PSD95') 219 127
('rep2', 'PSD95') 89 180
('rep3', 'PSD95') 192 149
('rep4', 'PSD95') 157 207
('rep5', 'PSD95') 102 120
('rep6', 'PSD95') 125 150
('rep7', 'PSD95') 141 143
('rep8', 'PSD95') 115 216
('rep1', 'PURA') 177 217
('rep2', 'PURA') 193 192
('rep3', 'PURA') 159 181
('rep4', 'PURA') 155 142
('rep5', 'PURA') 128 159
('rep6', 'PURA') 175 150
('rep7', 'PURA') 205 248
('rep8', 'PURA') 196 179
('rep1', 'Phalloidin') 184 153
('rep2', 'Phalloidin') 186 247
('rep3', 'Phalloidin') 158 165
('rep4', 'Phalloidin') 179 156
('rep5', 'Phalloidin') 211 150
('rep6', 'Phalloidin') 198 201
('rep7', 'Phalloidin') 163 138
('rep8', 'Phalloidin') 144 151
('rep1', 'SNCA') 167 122
('rep2', 'SNCA') 127 174
('rep3', 'SNCA') 192 176
('rep4', 'SNCA') 187 214
('rep5', 'SNCA') 162 156
('rep6', 'SNCA') 133 137
('rep7', 'SNCA') 209 141
('rep8', 'SNCA') 210 222
('rep1', 'SQSTM1') 221 92
('rep2', 'SQSTM1') 142 234
('rep3', 'SQSTM1') 133 144
('rep4', 'SQSTM1') 147 262
('rep5', 'SQSTM1') 164 197
('rep6', 'SQSTM1') 140 235
('rep7', 'SQSTM1') 83 106
('rep8', 'SQSTM1') 152 135
('rep1', 'TDP43') 132 168
('rep2', 'TDP43') 225 197
('rep3', 'TDP43') 180 249
('rep4', 'TDP43') 177 229
('rep5', 'TDP43') 168 204
('rep6', 'TDP43') 200 171
('rep7', 'TDP43') 182 188
('rep8', 'TDP43') 175 283
('rep1', 'TIA1') 177 213
('rep2', 'TIA1') 193 192
('rep3', 'TIA1') 159 181
('rep4', 'TIA1') 155 142
('rep5', 'TIA1') 124 160
('rep6', 'TIA1') 175 150
('rep7', 'TIA1') 205 248
('rep8', 'TIA1') 196 179
('rep1', 'TOMM20') 206 49
('rep2', 'TOMM20') 200 187
('rep3', 'TOMM20') 168 3
('rep4', 'TOMM20') 188
('rep5', 'TOMM20') 211 95
('rep6', 'TOMM20') 204 188
('rep7', 'TOMM20') 163 109
('rep8', 'TOMM20') 157 49
('rep1', 'TUJ1') 203 202
('rep2', 'TUJ1') 197 204
('rep3', 'TUJ1') 202 208
('rep4', 'TUJ1') 209 199
('rep5', 'TUJ1') 204 206
('rep6', 'TUJ1') 196 189
('rep7', 'TUJ1') 196 198
('rep8', 'TUJ1') 203 197
Total number of whole cells
250
200
150
100
50

### Batch 3

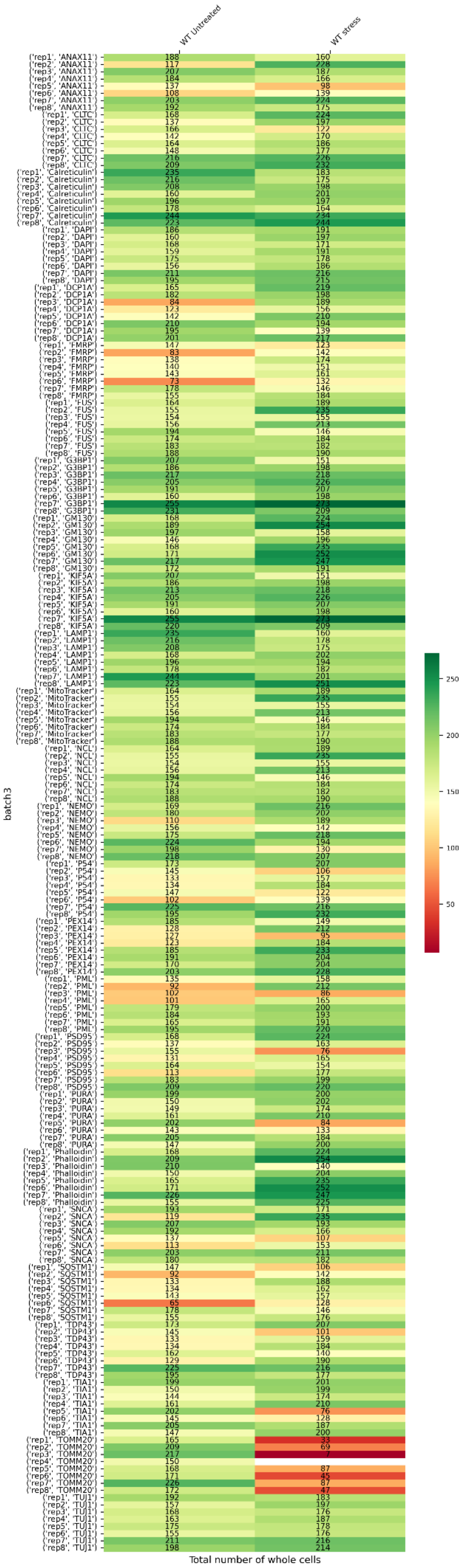

WT Untreated
WT stress
batch3
Total number of whole cells
250
200
150
100
50
