## Supplementary material for "Organellomics: AI-driven deep organellar phenotyping reveals novel ALS mechanisms in human neurons": Day 8 iPSC-derived cortical neurons (used for model inference, Fig 3) - QC Report.pdf

#### Processed Files Validation

Number of sites that survived pre-processing

Batch 1

| | Rep | FUSHomozygous | TDP43 | TBK1 | WT Untreated | FUSRevertant | OPTN | FUSHeterozygous |
|---|---|---|---|---|---|---|---|---|
| G3BP1 | rep1 | 200 | 188 | 205 | 223 | 245 | 238 | 249 |
| G3BP1 | rep2 | 222 | 205 | 183 | 179 | 231 | 229 | 247 |
| NONO | rep1 | 191 | 205 | 223 | 201 | 215 | 226 | 241 |
| NONO | rep2 | 214 | 206 | 208 | 200 | 241 | 215 | 244 |
| SQSTM1 | rep1 | 159 | 181 | 177 | 198 | 208 | 191 | 162 |
| SQSTM1 | rep2 | 124 | 175 | 159 | 195 | 185 | 192 | 164 |
| PSD95 | rep1 | 202 | 247 | 248 | 246 | 213 | 237 | 243 |
| PSD95 | rep2 | 246 | 243 | 248 | 249 | 227 | 243 | 246 |
| NEMO | rep1 | 213 | 246 | 244 | 204 | 247 | 244 | 184 |
| NEMO | rep2 | 248 | 244 | 247 | 245 | 192 | 244 | 228 |
| GM130 | rep1 | 248 | 250 | 250 | 249 | 247 | 250 | 249 |
| GM130 | rep2 | 247 | 250 | 250 | 250 | 248 | 250 | 250 |
| NCL | rep1 | 241 | 250 | 249 | 250 | 248 | 249 | 248 |
| NCL | rep2 | 250 | 250 | 249 | 250 | 247 | 250 | 250 |
| LSM14A | rep1 | 250 | 248 | 250 | 250 | 171 | 250 | 233 |
| LSM14A | rep2 | 236 | 249 | 250 | 240 | 223 | 250 | 243 |
| TDP43 | rep1 | 226 | 234 | 195 | 241 | 229 | 220 | 212 |
| TDP43 | rep2 | 188 | 225 | 189 | 235 | 220 | 222 | 225 |
| ANXA11 | rep1 | 189 | 236 | 226 | 211 | 182 | 201 | 242 |
| ANXA11 | rep2 | 188 | 249 | 212 | 236 | 245 | 214 | 238 |
| PEX14 | rep1 | 147 | 247 | 178 | 231 | 182 | 218 | 213 |
| PEX14 | rep2 | 198 | 244 | 213 | 246 | 185 | 202 | 222 |
| mitotracker | rep1 | 148 | 226 | 220 | 219 | 144 | 234 | 219 |
| mitotracker | rep2 | 227 | 242 | 234 | 230 | 196 | 245 | 198 |
| FMRP | rep1 | 186 | 152 | 194 | 208 | 242 | 233 | 248 |
| FMRP | rep2 | 201 | 167 | 193 | 181 | 227 | 212 | 243 |
| SON | rep1 | 205 | 244 | 238 | 211 | 216 | 226 | 240 |
| SON | rep2 | 224 | 225 | 226 | 218 | 239 | 217 | 248 |
| KIF5A | rep1 | 199 | 207 | 203 | 202 | 216 | 194 | 170 |
| KIF5A | rep2 | 156 | 211 | 179 | 216 | 199 | 203 | 206 |
| CLTC | rep1 | 230 | 249 | 250 | 246 | 212 | 238 | 247 |
| CLTC | rep2 | 249 | 250 | 249 | 250 | 228 | 249 | 246 |
| DCP1A | rep1 | 214 | 250 | 247 | 203 | 242 | 248 | 178 |
| DCP1A | rep2 | 250 | 250 | 250 | 249 | 185 | 249 | 227 |
| Calreticulin | rep1 | 248 | 249 | 247 | 249 | 249 | 250 | 248 |
| Calreticulin | rep2 | 246 | 248 | 249 | 250 | 248 | 250 | 248 |
| FUS | rep1 | 240 | 250 | 250 | 250 | 249 | 249 | 248 |
| FUS | rep2 | 250 | 250 | 249 | 250 | 247 | 250 | 250 |
| HNRNPA1 | rep1 | 240 | 234 | 240 | 247 | 164 | 250 | 229 |
| HNRNPA1 | rep2 | 234 | 246 | 246 | 238 | 209 | 249 | 246 |
| PML | rep1 | 230 | 248 | 240 | 246 | 221 | 244 | 217 |
| PML | rep2 | 216 | 248 | 249 | 246 | 233 | 245 | 220 |
| LAMP1 | rep1 | 81 | 77 | 77 | 87 | 60 | 113 | 66 |
| LAMP1 | rep2 | 57 | 76 | 81 | 66 | 64 | 102 | 82 |
| SNCA | rep1 | 95 | 143 | 122 | 107 | 134 | 118 | 146 |
| SNCA | rep2 | 131 | 104 | 118 | 136 | 111 | 116 | 133 |
| TIA1 | rep1 | 227 | 225 | 233 | 198 | 227 | 238 | 234 |
| TIA1 | rep2 | 200 | 228 | 218 | 207 | 205 | 227 | 218 |
| PURA | rep1 | 155 | 170 | 203 | 187 | 208 | 198 | 216 |
| PURA | rep2 | 189 | 175 | 193 | 184 | 207 | 190 | 208 |
| CD41 | rep1 | nan | nan | nan | nan | nan | nan | nan |
| CD41 | rep2 | nan | nan | nan | nan | nan | nan | nan |
| Tubulin | rep1 | 145 | 204 | 184 | 197 | 179 | 187 | 132 |
| Tubulin | rep2 | 103 | 206 | 160 | 213 | 183 | 203 | 147 |
| Phalloidin | rep1 | 209 | 233 | 233 | 230 | 188 | 225 | 170 |
| Phalloidin | rep2 | 203 | 238 | 221 | 238 | 209 | 239 | 188 |
| TOMM20 | rep1 | 203 | 242 | 238 | 227 | 234 | 235 | 236 |
| TOMM20 | rep2 | 220 | 232 | 239 | 228 | 211 | 233 | 210 |
| DAPI | rep1 | 2638 | 2961 | 2911 | 2847 | 2684 | 2868 | 2801 |
| DAPI | rep2 | 2721 | 2939 | 2882 | 2916 | 2754 | 2871 | 2903 |

#### Batch 2

| | Rep | FUSHomozygous | TDP43 | TBK1 | WT Untreated | FUSRevertant | OPTN | FUSHeterozygous |
|---|---|---|---|---|---|---|---|---|
| G3BP1 | rep1 | 240 | 245 | 240 | 241 | 216 | 242 | 244 |
| G3BP1 | rep2 | 230 | 245 | 231 | 0 | 194 | 240 | 224 |
| NONO | rep1 | 249 | 246 | 240 | 250 | 235 | 247 | 207 |
| NONO | rep2 | 249 | 208 | 250 | 249 | 244 | 248 | 202 |
| SQSTM1 | rep1 | 168 | 159 | 215 | 208 | 204 | 216 | 199 |
| SQSTM1 | rep2 | 218 | 165 | 224 | 211 | 188 | 228 | 208 |
| PSD95 | rep1 | 238 | 246 | 250 | 236 | 229 | 247 | 250 |
| PSD95 | rep2 | 249 | 248 | 249 | 248 | 226 | 250 | 249 |
| NEMO | rep1 | 244 | 247 | 248 | 242 | 141 | 245 | 130 |
| NEMO | rep2 | 238 | 173 | 244 | 241 | 200 | 243 | 216 |
| GM130 | rep1 | 245 | 244 | 248 | 221 | 192 | 197 | 247 |
| GM130 | rep2 | 248 | 237 | 249 | 184 | 214 | 250 | 249 |
| NCL | rep1 | 241 | 249 | 246 | 250 | 223 | 249 | 248 |
| NCL | rep2 | 240 | 178 | 213 | 199 | 235 | 249 | 248 |
| LSM14A | rep1 | 248 | 250 | 232 | 250 | 242 | 250 | 247 |
| LSM14A | rep2 | 243 | 246 | 248 | 248 | 231 | 250 | 240 |
| TDP43 | rep1 | 238 | 231 | 236 | 242 | 210 | 244 | 232 |
| TDP43 | rep2 | 233 | 234 | 234 | 241 | 214 | 230 | 234 |
| ANXA11 | rep1 | 227 | 237 | 229 | 241 | 233 | 230 | 235 |
| ANXA11 | rep2 | 204 | 215 | 247 | 240 | 188 | 238 | 190 |
| PEX14 | rep1 | 236 | 238 | 248 | 239 | 246 | 248 | 234 |
| PEX14 | rep2 | 233 | 236 | 245 | 247 | 32 | 243 | 223 |
| mitotracker | rep1 | 210 | 242 | 239 | 232 | 210 | 248 | 224 |
| mitotracker | rep2 | 88 | 242 | 237 | 249 | 188 | 247 | 212 |
| FMRP | rep1 | 238 | 235 | 232 | 238 | 209 | 238 | 243 |
| FMRP | rep2 | 228 | 243 | 225 | 0 | 192 | 227 | 215 |
| SON | rep1 | 249 | 247 | 240 | 250 | 238 | 250 | 207 |
| SON | rep2 | 249 | 206 | 250 | 250 | 245 | 249 | 201 |
| KIF5A | rep1 | 183 | 177 | 229 | 231 | 226 | 231 | 215 |
| KIF5A | rep2 | 236 | 183 | 232 | 224 | 209 | 236 | 221 |
| CLTC | rep1 | 237 | 247 | 250 | 236 | 229 | 248 | 250 |
| CLTC | rep2 | 249 | 249 | 250 | 250 | 227 | 250 | 249 |
| DCP1A | rep1 | 250 | 250 | 249 | 245 | 141 | 250 | 123 |
| DCP1A | rep2 | 248 | 172 | 248 | 250 | 202 | 250 | 226 |
| Calreticulin | rep1 | 250 | 244 | 249 | 215 | 201 | 196 | 248 |
| Calreticulin | rep2 | 247 | 235 | 248 | 180 | 240 | 250 | 250 |
| FUS | rep1 | 247 | 244 | 247 | 248 | 217 | 249 | 247 |
| FUS | rep2 | 249 | 170 | 210 | 196 | 238 | 249 | 248 |
| HNRNPA1 | rep1 | 244 | 249 | 228 | 249 | 235 | 250 | 249 |
| HNRNPA1 | rep2 | 241 | 245 | 246 | 247 | 225 | 249 | 242 |
| PML | rep1 | 239 | 240 | 245 | 242 | 186 | 228 | 225 |
| PML | rep2 | 216 | 242 | 247 | 246 | 241 | 248 | 209 |
| LAMP1 | rep1 | 96 | 124 | 138 | 95 | 66 | 130 | 88 |
| LAMP1 | rep2 | 83 | 130 | 148 | 92 | 81 | 80 | 99 |
| SNCA | rep1 | 124 | 116 | 114 | 152 | 138 | 140 | 98 |
| SNCA | rep2 | 152 | 142 | 100 | 165 | 72 | 104 | 136 |
| TIA1 | rep1 | 162 | 224 | 224 | 210 | 190 | 221 | 173 |
| TIA1 | rep2 | 84 | 220 | 218 | 224 | 171 | 214 | 204 |
| PURA | rep1 | 202 | 206 | 199 | 202 | 187 | 211 | 197 |
| PURA | rep2 | 193 | 202 | 184 | 0 | 157 | 185 | 188 |
| CD41 | rep1 | nan | nan | nan | nan | nan | nan | nan |
| CD41 | rep2 | nan | nan | nan | nan | nan | nan | nan |
| Tubulin | rep1 | 181 | 175 | 240 | 225 | 216 | 227 | 190 |
| Tubulin | rep2 | 223 | 159 | 222 | 230 | 209 | 234 | 224 |
| Phalloidin | rep1 | 183 | 198 | 221 | 221 | 209 | 222 | 139 |
| Phalloidin | rep2 | 203 | 203 | 222 | 218 | 210 | 229 | 156 |
| TOMM20 | rep1 | 193 | 219 | 222 | 211 | 194 | 235 | 209 |
| TOMM20 | rep2 | 72 | 226 | 233 | 229 | 193 | 234 | 204 |
| DAPI | rep1 | 2862 | 2908 | 2940 | 2918 | 2662 | 2925 | 2768 |
| DAPI | rep2 | 2761 | 2696 | 2928 | 2719 | 2580 | 2975 | 2801 |

<u>Batch 3</u>

| | Rep | FUSHomozygous | TDP43 | TBK1 | WT Untreated | FUSRevertant | OPTN | FUSHeterozygous |
|---|---|---|---|---|---|---|---|---|
| G3BP1 | rep1 | 247 | 247 | 238 | 246 | 222 | 242 | 217 |
| G3BP1 | rep2 | 220 | 237 | 194 | 235 | 161 | 200 | 204 |
| NONO | rep1 | 187 | 225 | 249 | 250 | 167 | 193 | 181 |
| NONO | rep2 | 181 | 241 | 246 | 228 | 186 | 186 | 185 |
| SQSTM1 | rep1 | 169 | 206 | 183 | 183 | 196 | 197 | 216 |
| SQSTM1 | rep2 | 158 | 214 | 220 | 234 | 179 | 219 | 163 |
| PSD95 | rep1 | 249 | 243 | 247 | 213 | 246 | 249 | 242 |
| PSD95 | rep2 | 248 | 249 | 243 | 246 | 229 | 248 | 246 |
| NEMO | rep1 | 249 | 240 | 236 | 243 | 234 | 244 | 238 |
| NEMO | rep2 | 243 | 239 | 199 | 247 | 238 | 247 | 238 |
| GM130 | rep1 | 248 | 197 | 246 | 250 | 243 | 193 | 232 |
| GM130 | rep2 | 249 | 225 | 250 | 221 | 246 | 244 | 241 |
| NCL | rep1 | 249 | 250 | 227 | 217 | 248 | 250 | 226 |
| NCL | rep2 | 248 | 245 | 245 | 250 | 250 | 250 | 247 |
| LSM14A | rep1 | 250 | 250 | 244 | 250 | 241 | 250 | 225 |
| LSM14A | rep2 | 246 | 250 | 245 | 250 | 224 | 250 | 221 |
| TDP43 | rep1 | 228 | 228 | 236 | 233 | 207 | 235 | 219 |
| TDP43 | rep2 | 195 | 235 | 232 | 237 | 220 | 238 | 207 |
| ANXA11 | rep1 | 236 | 227 | 235 | 227 | 221 | 240 | 230 |
| ANXA11 | rep2 | 236 | 219 | 232 | 238 | 230 | 242 | 178 |
| PEX14 | rep1 | 246 | 246 | 248 | 222 | 182 | 249 | 215 |
| PEX14 | rep2 | 240 | 247 | 247 | 248 | 185 | 247 | 237 |
| mitotracker | rep1 | 235 | 247 | 249 | 247 | 218 | 248 | 231 |
| mitotracker | rep2 | 240 | 240 | 245 | 249 | 234 | 249 | 231 |
| FMRP | rep1 | 244 | 234 | 229 | 235 | 208 | 226 | 205 |
| FMRP | rep2 | 216 | 207 | 182 | 226 | 154 | 189 | 194 |
| SON | rep1 | 185 | 225 | 247 | 250 | 165 | 191 | 186 |
| SON | rep2 | 180 | 249 | 249 | 227 | 185 | 188 | 185 |
| KIF5A | rep1 | 174 | 196 | 212 | 189 | 201 | 205 | 228 |
| KIF5A | rep2 | 155 | 198 | 235 | 211 | 187 | 223 | 176 |
| CLTC | rep1 | 250 | 240 | 249 | 197 | 246 | 228 | 243 |
| CLTC | rep2 | 250 | 250 | 245 | 248 | 228 | 248 | 248 |
| DCP1A | rep1 | 246 | 246 | 232 | 247 | 237 | 248 | 228 |
| DCP1A | rep2 | 246 | 249 | 183 | 243 | 246 | 247 | 238 |
| Calreticulin | rep1 | 250 | 193 | 248 | 245 | 244 | 188 | 242 |
| Calreticulin | rep2 | 247 | 218 | 247 | 215 | 249 | 239 | 245 |
| FUS | rep1 | 250 | 250 | 222 | 214 | 247 | 249 | 238 |
| FUS | rep2 | 250 | 244 | 247 | 249 | 248 | 250 | 247 |
| HNRNPA1 | rep1 | 249 | 250 | 238 | 248 | 234 | 250 | 219 |
| HNRNPA1 | rep2 | 249 | 250 | 244 | 247 | 210 | 250 | 222 |
| PML | rep1 | 214 | 244 | 243 | 194 | 232 | 245 | 187 |
| PML | rep2 | 192 | 235 | 202 | 189 | 214 | 223 | 186 |
| LAMP1 | rep1 | 68 | 57 | 53 | 64 | 83 | 59 | 100 |
| LAMP1 | rep2 | 44 | 38 | 100 | 51 | 95 | 71 | 86 |
| SNCA | rep1 | 141 | 105 | 122 | 137 | 112 | 116 | 103 |
| SNCA | rep2 | 139 | 125 | 105 | 150 | 126 | 88 | 118 |
| TIA1 | rep1 | 221 | 206 | 225 | 217 | 226 | 234 | 227 |
| TIA1 | rep2 | 224 | 214 | 221 | 222 | 217 | 222 | 211 |
| PURA | rep1 | 181 | 201 | 173 | 205 | 190 | 202 | 181 |
| PURA | rep2 | 176 | 181 | 177 | 205 | 145 | 177 | 163 |
| CD41 | rep1 | nan | nan | nan | nan | nan | nan | nan |
| CD41 | rep2 | nan | nan | nan | nan | nan | nan | nan |
| Tubulin | rep1 | 159 | 213 | 189 | 179 | 187 | 219 | 223 |
| Tubulin | rep2 | 135 | 220 | 229 | 238 | 190 | 210 | 157 |
| Phalloidin | rep1 | 222 | 230 | 225 | 230 | 205 | 188 | 210 |
| Phalloidin | rep2 | 207 | 236 | 235 | 237 | 215 | 238 | 195 |
| TOMM20 | rep1 | 235 | 239 | 232 | 235 | 227 | 238 | 229 |
| TOMM20 | rep2 | 236 | 231 | 230 | 241 | 233 | 225 | 230 |
| DAPI | rep1 | 2845 | 2898 | 2915 | 2871 | 2766 | 2866 | 2790 |
| DAPI | rep2 | 2771 | 2931 | 2865 | 2917 | 2712 | 2886 | 2691 |

#### Batch 4

| | Rep | FUSHomozygous | TDP43 | TBK1 | WT Untreated | FUSRevertant | OPTN | FUSHeterozygous |
|---|---|---|---|---|---|---|---|---|
| G3BP1 | rep1 | 184 | 214 | 211 | 219 | 184 | 220 | 196 |
| G3BP1 | rep2 | 238 | 204 | 107 | 164 | 204 | 213 | 230 |
| NONO | rep1 | 218 | 237 | 11 | 124 | 142 | 231 | 181 |
| NONO | rep2 | 219 | 238 | 220 | 227 | 211 | 241 | 138 |
| SQSTM1 | rep1 | 178 | 120 | 163 | 169 | 136 | 164 | 166 |
| SQSTM1 | rep2 | 159 | 209 | 212 | 197 | 125 | 198 | 138 |
| PSD95 | rep1 | 240 | 242 | 178 | 167 | 192 | 247 | 175 |
| PSD95 | rep2 | 191 | 248 | 227 | 243 | 149 | 248 | 80 |
| NEMO | rep1 | 206 | 231 | 237 | 187 | 216 | 239 | 158 |
| NEMO | rep2 | 213 | 230 | 238 | 186 | 211 | 238 | 140 |
| GM130 | rep1 | 249 | 250 | 250 | 250 | 250 | 250 | 250 |
| GM130 | rep2 | 250 | 236 | 249 | 249 | 250 | 250 | 250 |
| NCL | rep1 | 233 | 250 | 249 | 249 | 249 | 249 | 209 |
| NCL | rep2 | 248 | 249 | 249 | 250 | 247 | 250 | 221 |
| LSM14A | rep1 | 243 | 248 | 248 | 247 | 249 | 248 | 239 |
| LSM14A | rep2 | 245 | 249 | 250 | 248 | 248 | 249 | 243 |
| TDP43 | rep1 | 240 | 242 | 231 | 244 | 138 | 246 | 202 |
| TDP43 | rep2 | 234 | 238 | 249 | 225 | 146 | 242 | 203 |
| ANXA11 | rep1 | 246 | 155 | 171 | 238 | 206 | 230 | 218 |
| ANXA11 | rep2 | 238 | 242 | 240 | 220 | 222 | 238 | 245 |
| PEX14 | rep1 | 150 | 247 | 119 | 221 | 168 | 246 | 148 |
| PEX14 | rep2 | 246 | 236 | 221 | 169 | 174 | 244 | 239 |
| mitotracker | rep1 | 248 | 156 | 244 | 242 | 241 | 246 | 246 |
| mitotracker | rep2 | 245 | 233 | 232 | 240 | 243 | 241 | 241 |
| FMRP | rep1 | 170 | 206 | 211 | 211 | 172 | 211 | 176 |
| FMRP | rep2 | 214 | 177 | 94 | 147 | 186 | 204 | 173 |
| SON | rep1 | 236 | 242 | 13 | 165 | 196 | 235 | 225 |
| SON | rep2 | 239 | 240 | 240 | 241 | 231 | 242 | 216 |
| KIF5A | rep1 | 208 | 136 | 182 | 193 | 156 | 188 | 207 |
| KIF5A | rep2 | 204 | 226 | 229 | 208 | 160 | 224 | 185 |
| CLTC | rep1 | 249 | 242 | 176 | 247 | 193 | 247 | 206 |
| CLTC | rep2 | 197 | 249 | 227 | 250 | 149 | 249 | 155 |
| DCP1A | rep1 | 243 | 248 | 245 | 245 | 241 | 250 | 232 |
| DCP1A | rep2 | 233 | 243 | 244 | 249 | 240 | 248 | 204 |
| Calreticulin | rep1 | 242 | 250 | 247 | 241 | 238 | 248 | 220 |
| Calreticulin | rep2 | 247 | 246 | 248 | 250 | 220 | 248 | 221 |
| FUS | rep1 | 4 | 3 | 1 | 2 | 5 | 0 | 2 |
| FUS | rep2 | 1 | 0 | 4 | 2 | 0 | 1 | 1 |
| HNRNPA1 | rep1 | 246 | 247 | 249 | 229 | 232 | 246 | 217 |
| HNRNPA1 | rep2 | 247 | 246 | 248 | 223 | 236 | 249 | 235 |
| PML | rep1 | 249 | 248 | 232 | 242 | 207 | 249 | 241 |
| PML | rep2 | 249 | 246 | 249 | 250 | 243 | 249 | 248 |
| LAMP1 | rep1 | 139 | 99 | 101 | 124 | 157 | 143 | 84 |
| LAMP1 | rep2 | 123 | 122 | 209 | 79 | 151 | 186 | 149 |
| SNCA | rep1 | 88 | 104 | 95 | 90 | 88 | 51 | 112 |
| SNCA | rep2 | 128 | 125 | 151 | 108 | 130 | 128 | 179 |
| TIA1 | rep1 | 206 | 152 | 178 | 174 | 229 | 238 | 199 |
| TIA1 | rep2 | 201 | 189 | 163 | 135 | 228 | 198 | 198 |
| PURA | rep1 | 168 | 196 | 194 | 203 | 185 | 198 | 167 |
| PURA | rep2 | 205 | 181 | 102 | 209 | 185 | 193 | 197 |
| CD41 | rep1 | nan | nan | nan | nan | nan | nan | nan |
| CD41 | rep2 | nan | nan | nan | nan | nan | nan | nan |
| Tubulin | rep1 | 243 | 139 | 170 | 239 | 171 | 171 | 206 |
| Tubulin | rep2 | 240 | 226 | 234 | 239 | 168 | 212 | 185 |
| Phalloidin | rep1 | 237 | 226 | 174 | 227 | 190 | 243 | 202 |
| Phalloidin | rep2 | 180 | 237 | 220 | 222 | 143 | 243 | 218 |
| TOMM20 | rep1 | 246 | 162 | 249 | 249 | 247 | 248 | 243 |
| TOMM20 | rep2 | 245 | 245 | 236 | 246 | 241 | 249 | 246 |
| DAPI | rep1 | 2817 | 2695 | 2428 | 2880 | 2737 | 2909 | 2769 |
| DAPI | rep2 | 2916 | 2941 | 2803 | 2909 | 2747 | 2955 | 2877 |

<u>Batch 5</u>

| | Rep | FUSHomozygous | TDP43 | TBK1 | WT Untreated | FUSRevertant | OPTN | FUSHeterozygous |
|---|---|---|---|---|---|---|---|---|
| G3BP1 | rep1 | 244 | 236 | 189 | 235 | 219 | 228 | 239 |
| G3BP1 | rep2 | 234 | 219 | 184 | 218 | 216 | 214 | 222 |
| NONO | rep1 | 238 | 217 | 249 | 226 | 225 | 228 | 202 |
| NONO | rep2 | 246 | 233 | 247 | 224 | 223 | 226 | 207 |
| SQSTM1 | rep1 | 156 | 125 | 184 | 141 | 150 | 167 | 142 |
| SQSTM1 | rep2 | 183 | 180 | 172 | 169 | 184 | 159 | 164 |
| PSD95 | rep1 | 237 | 235 | 218 | 209 | 249 | 222 | 239 |
| PSD95 | rep2 | 247 | 249 | 250 | 17 | 248 | 249 | 224 |
| NEMO | rep1 | 248 | 250 | 246 | 250 | 248 | 247 | 248 |
| NEMO | rep2 | 231 | 248 | 247 | 247 | 248 | 240 | 249 |
| GM130 | rep1 | 250 | 250 | 250 | 250 | 250 | 249 | 250 |
| GM130 | rep2 | 250 | 250 | 250 | 250 | 250 | 249 | 249 |
| NCL | rep1 | 249 | 248 | 249 | 250 | 247 | 250 | 250 |
| NCL | rep2 | 250 | 249 | 248 | 250 | 240 | 250 | 248 |
| LSM14A | rep1 | 250 | 250 | 250 | 250 | 250 | 250 | 250 |
| LSM14A | rep2 | 250 | 250 | 249 | 250 | 249 | 250 | 250 |
| TDP43 | rep1 | 236 | 242 | 245 | 244 | 134 | 238 | 234 |
| TDP43 | rep2 | 243 | 248 | 237 | 248 | 213 | 235 | 236 |
| ANXA11 | rep1 | 242 | 242 | 242 | 240 | 234 | 241 | 237 |
| ANXA11 | rep2 | 242 | 241 | 233 | 237 | 228 | 241 | 232 |
| PEX14 | rep1 | 246 | 250 | 247 | 246 | 146 | 239 | 196 |
| PEX14 | rep2 | 241 | 242 | 246 | 247 | 203 | 240 | 191 |
| mitotracker | rep1 | 248 | 250 | 250 | 232 | 233 | 247 | 240 |
| mitotracker | rep2 | 246 | 241 | 236 | 246 | 247 | 243 | 245 |
| FMRP | rep1 | 233 | 212 | 182 | 214 | 200 | 224 | 201 |
| FMRP | rep2 | 212 | 197 | 172 | 200 | 192 | 205 | 196 |
| SON | rep1 | 246 | 238 | 250 | 249 | 244 | 240 | 237 |
| SON | rep2 | 250 | 243 | 249 | 242 | 244 | 225 | 221 |
| KIF5A | rep1 | 181 | 150 | 191 | 166 | 188 | 192 | 169 |
| KIF5A | rep2 | 188 | 206 | 173 | 187 | 219 | 173 | 202 |
| CLTC | rep1 | 231 | 237 | 219 | 228 | 248 | 225 | 235 |
| CLTC | rep2 | 247 | 250 | 250 | 181 | 250 | 250 | 226 |
| DCP1A | rep1 | 250 | 250 | 249 | 250 | 249 | 249 | 249 |
| DCP1A | rep2 | 231 | 250 | 249 | 247 | 246 | 249 | 249 |
| Calreticulin | rep1 | 250 | 248 | 250 | 250 | 250 | 250 | 250 |
| Calreticulin | rep2 | 250 | 249 | 248 | 250 | 250 | 250 | 249 |
| FUS | rep1 | 250 | 249 | 248 | 250 | 248 | 250 | 250 |
| FUS | rep2 | 249 | 249 | 248 | 250 | 250 | 250 | 250 |
| HNRNPA1 | rep1 | 242 | 249 | 250 | 248 | 226 | 249 | 241 |
| HNRNPA1 | rep2 | 246 | 247 | 245 | 249 | 222 | 249 | 227 |
| PML | rep1 | 248 | 239 | 224 | 189 | 219 | 250 | 224 |
| PML | rep2 | 242 | 218 | 197 | 231 | 234 | 246 | 236 |
| LAMP1 | rep1 | 168 | 112 | 176 | 145 | 164 | 124 | 142 |
| LAMP1 | rep2 | 132 | 162 | 209 | 184 | 128 | 165 | 152 |
| SNCA | rep1 | 95 | 123 | 101 | 92 | 159 | 115 | 109 |
| SNCA | rep2 | 133 | 174 | 173 | 99 | 101 | 117 | 115 |
| TIA1 | rep1 | 226 | 202 | 195 | 213 | 233 | 230 | 201 |
| TIA1 | rep2 | 205 | 197 | 215 | 161 | 240 | 214 | 213 |
| PURA | rep1 | 197 | 211 | 172 | 186 | 193 | 204 | 195 |
| PURA | rep2 | 186 | 182 | 149 | 182 | 196 | 182 | 182 |
| CD41 | rep1 | nan | nan | nan | nan | nan | nan | nan |
| CD41 | rep2 | nan | nan | nan | nan | nan | nan | nan |
| Tubulin | rep1 | 235 | 198 | 244 | 225 | 240 | 220 | 209 |
| Tubulin | rep2 | 242 | 208 | 202 | 216 | 238 | 208 | 213 |
| Phalloidin | rep1 | 230 | 214 | 205 | 230 | 232 | 220 | 212 |
| Phalloidin | rep2 | 225 | 238 | 228 | 193 | 234 | 246 | 197 |
| TOMM20 | rep1 | 239 | 246 | 250 | 247 | 232 | 246 | 193 |
| TOMM20 | rep2 | 230 | 248 | 247 | 246 | 219 | 248 | 219 |
| DAPI | rep1 | 2977 | 2934 | 2902 | 2972 | 2910 | 2939 | 2916 |
| DAPI | rep2 | 2958 | 2950 | 2887 | 2931 | 2925 | 2919 | 2873 |

#### Batch 6

| | Rep | FUSHomozygous | TDP43 | TBK1 | WT Untreated | FUSRevertant | OPTN | FUSHeterozygous |
|---|---|---|---|---|---|---|---|---|
| G3BP1 | rep1 | 240 | 232 | 228 | 238 | 232 | 238 | 242 |
| G3BP1 | rep2 | 244 | 246 | 232 | 240 | 231 | 237 | 250 |
| NONO | rep1 | 250 | 223 | 246 | 0 | 224 | 250 | 241 |
| NONO | rep2 | 250 | 228 | 206 | 250 | 230 | 248 | 240 |
| SQSTM1 | rep1 | 157 | 179 | 114 | 148 | 186 | 138 | 166 |
| SQSTM1 | rep2 | 156 | 160 | 151 | 159 | 170 | 148 | 182 |
| PSD95 | rep1 | 249 | 245 | 223 | 249 | 237 | 237 | 249 |
| PSD95 | rep2 | 247 | 246 | 249 | 245 | 248 | 230 | 245 |
| NEMO | rep1 | 250 | 249 | 249 | 248 | 249 | 248 | 249 |
| NEMO | rep2 | 249 | 250 | 249 | 248 | 246 | 250 | 249 |
| GM130 | rep1 | 249 | 250 | 249 | 250 | 250 | 0 | 250 |
| GM130 | rep2 | 250 | 249 | 250 | 250 | 250 | 249 | 250 |
| NCL | rep1 | 250 | 250 | 250 | 249 | 248 | 250 | 250 |
| NCL | rep2 | 250 | 249 | 250 | 246 | 244 | 250 | 246 |
| LSM14A | rep1 | 250 | 250 | 250 | 250 | 250 | 250 | 250 |
| LSM14A | rep2 | 250 | 250 | 250 | 249 | 250 | 250 | 250 |
| TDP43 | rep1 | 92 | 212 | 98 | 221 | 26 | 147 | 47 |
| TDP43 | rep2 | 241 | 192 | 158 | 218 | 174 | 171 | 120 |
| ANXA11 | rep1 | 58 | 178 | 48 | 156 | 132 | 138 | 70 |
| ANXA11 | rep2 | 46 | 115 | 100 | 149 | 56 | 51 | 57 |
| PEX14 | rep1 | 112 | 173 | 138 | 185 | 105 | 131 | 74 |
| PEX14 | rep2 | 91 | 147 | 148 | 216 | 140 | 144 | 90 |
| mitotracker | rep1 | 218 | 224 | 191 | 250 | 199 | 200 | 249 |
| mitotracker | rep2 | 237 | 242 | 242 | 247 | 101 | 237 | 85 |
| FMRP | rep1 | 237 | 227 | 224 | 224 | 228 | 241 | 236 |
| FMRP | rep2 | 241 | 243 | 225 | 198 | 218 | 233 | 246 |
| SON | rep1 | 249 | 225 | 248 | 4 | 242 | 249 | 246 |
| SON | rep2 | 250 | 227 | 206 | 249 | 244 | 247 | 245 |
| KIF5A | rep1 | 189 | 213 | 146 | 192 | 219 | 202 | 202 |
| KIF5A | rep2 | 189 | 193 | 185 | 189 | 220 | 187 | 228 |
| CLTC | rep1 | 250 | 249 | 222 | 250 | 240 | 237 | 250 |
| CLTC | rep2 | 249 | 244 | 249 | 244 | 250 | 236 | 250 |
| DCP1A | rep1 | 250 | 248 | 250 | 250 | 250 | 250 | 250 |
| DCP1A | rep2 | 250 | 250 | 250 | 250 | 248 | 250 | 250 |
| Calreticulin | rep1 | 247 | 250 | 249 | 250 | 247 | 0 | 247 |
| Calreticulin | rep2 | 250 | 248 | 250 | 248 | 249 | 249 | 248 |
| FUS | rep1 | 248 | 250 | 250 | 249 | 250 | 250 | 250 |
| FUS | rep2 | 249 | 250 | 250 | 247 | 250 | 250 | 250 |
| HNRNPA1 | rep1 | 247 | 248 | 249 | 244 | 250 | 247 | 235 |
| HNRNPA1 | rep2 | 248 | 249 | 249 | 241 | 229 | 250 | 235 |
| PML | rep1 | 88 | 222 | 101 | 243 | 25 | 159 | 47 |
| PML | rep2 | 242 | 195 | 160 | 245 | 178 | 158 | 155 |
| LAMP1 | rep1 | 47 | 217 | 118 | 129 | 154 | 200 | 60 |
| LAMP1 | rep2 | 43 | 118 | 113 | 122 | 89 | 92 | 50 |
| SNCA | rep1 | 69 | 111 | 97 | 103 | 71 | 100 | 61 |
| SNCA | rep2 | 49 | 77 | 95 | 137 | 108 | 103 | 74 |
| TIA1 | rep1 | 97 | 166 | 101 | 154 | 177 | 188 | 158 |
| TIA1 | rep2 | 159 | 170 | 177 | 132 | 100 | 230 | 71 |
| PURA | rep1 | 191 | 203 | 195 | 187 | 196 | 201 | 213 |
| PURA | rep2 | 190 | 180 | 200 | 214 | 200 | 217 | 183 |
| CD41 | rep1 | nan | nan | nan | nan | nan | nan | nan |
| CD41 | rep2 | nan | nan | nan | nan | nan | nan | nan |
| Tubulin | rep1 | 240 | 235 | 188 | 242 | 243 | 237 | 244 |
| Tubulin | rep2 | 196 | 175 | 201 | 212 | 237 | 222 | 246 |
| Phalloidin | rep1 | 208 | 234 | 205 | 220 | 229 | 231 | 217 |
| Phalloidin | rep2 | 222 | 233 | 214 | 190 | 240 | 230 | 221 |
| TOMM20 | rep1 | 221 | 227 | 195 | 248 | 199 | 197 | 232 |
| TOMM20 | rep2 | 244 | 249 | 248 | 249 | 106 | 248 | 79 |
| DAPI | rep1 | 2502 | 2843 | 2485 | 2661 | 2524 | 2493 | 2471 |
| DAPI | rep2 | 2613 | 2702 | 2626 | 2861 | 2551 | 2665 | 2418 |

### Total Tile Count

#### Batch1

| batch1 | FUSHeterozygous | FUSHomozygous | FUSRevertant | OPTN | TBK1 | TDP43 | WT Untreated |
|---|---|---|---|---|---|---|---|
| ('rep1', 'ANXA11') | 2189 | 1852 | 1438 | 2268 | 2625 | 2698 | 2312 |
| ('rep2', 'ANXA11') | 1862 | 1861 | 2037 | 2393 | 2697 | 2924 | 2465 |
| ('rep1', 'CD41') | 1991 | 2337 | 1204 | 2003 | 2791 | 3115 | 2485 |
| ('rep2', 'CD41') | 2226 | 2471 | 1406 | 2107 | 2656 | 2608 | 2616 |
| ('rep1', 'CLTC') | 2031 | 2278 | 1088 | 2294 | 2858 | 3017 | 2927 |
| ('rep2', 'CLTC') | 1654 | 2241 | 949 | 2419 | 2787 | 3155 | 2823 |
| ('rep1', 'Calreticulin') | 2319 | 3163 | 1933 | 2956 | 3642 | 3479 | 3420 |
| ('rep2', 'Calreticulin') | 2415 | 3095 | 2171 | 2838 | 3459 | 3502 | 3219 |
| ('rep1', 'DAPI') | 2005 | 2305 | 1539 | 2378 | 2999 | 3010 | 2669 |
| ('rep2', 'DAPI') | 2001 | 2441 | 1530 | 2494 | 2929 | 3009 | 2659 |
| ('rep1', 'DCP1A') | 1340 | 2015 | 1662 | 2459 | 3237 | 3473 | 2325 |
| ('rep2', 'DCP1A') | 2093 | 3088 | 1335 | 2747 | 3286 | 3763 | 2897 |
| ('rep1', 'FMRP') | 1802 | 1466 | 1293 | 1378 | 1238 | 1231 | 1551 |
| ('rep2', 'FMRP') | 1331 | 1803 | 861 | 1865 | 1616 | 1661 | 1127 |
| ('rep1', 'FUS') | 2250 | 2738 | 1658 | 2864 | 3433 | 3587 | 3247 |
| ('rep2', 'FUS') | 2290 | 3045 | 1615 | 2828 | 3403 | 3524 | 3340 |
| ('rep1', 'G3BP1') | 1803 | 1564 | 1299 | 1401 | 1304 | 1532 | 1651 |
| ('rep2', 'G3BP1') | 1354 | 1975 | 865 | 2033 | 1453 | 2046 | 1040 |
| ('rep1', 'GM130') | 2336 | 3163 | 1937 | 2956 | 3676 | 3486 | 3420 |
| ('rep2', 'GM130') | 2435 | 3099 | 2171 | 2838 | 3472 | 3517 | 3219 |
| ('rep1', 'HNRNPA1') | 2142 | 3070 | 1095 | 2727 | 3354 | 2788 | 3137 |
| ('rep2', 'HNRNPA1') | 2140 | 2514 | 1456 | 2708 | 3169 | 3150 | 2892 |
| ('rep1', 'KIF5A') | 1462 | 2059 | 1448 | 1804 | 2270 | 2767 | 2297 |
| ('rep2', 'KIF5A') | 1706 | 1697 | 1141 | 1963 | 1971 | 2642 | 2568 |
| ('rep1', 'LAMP1') | 611 | 740 | 470 | 1207 | 929 | 891 | 896 |
| ('rep2', 'LAMP1') | 629 | 515 | 553 | 1125 | 975 | 847 | 683 |
| ('rep1', 'LSM14A') | 2182 | 3260 | 1147 | 2727 | 3516 | 3024 | 3194 |
| ('rep2', 'LSM14A') | 2124 | 2535 | 1471 | 2725 | 3243 | 3210 | 2911 |
| ('rep1', 'NCL') | 2250 | 2751 | 1654 | 2864 | 3413 | 3587 | 3247 |
| ('rep2', 'NCL') | 2290 | 3045 | 1615 | 2828 | 3403 | 3524 | 3340 |
| ('rep1', 'NEMO') | 1353 | 2003 | 1690 | 2413 | 3200 | 3416 | 2309 |
| ('rep2', 'NEMO') | 2064 | 3048 | 1357 | 2693 | 3245 | 3677 | 2850 |
| ('rep1', 'NONO') | 1991 | 2117 | 1184 | 1962 | 2533 | 2417 | 2267 |
| ('rep2', 'NONO') | 2125 | 2350 | 1404 | 2078 | 2318 | 2228 | 2277 |
| ('rep1', 'PEX14') | 1994 | 1315 | 1406 | 2449 | 2184 | 2643 | 2371 |
| ('rep2', 'PEX14') | 1899 | 1880 | 1318 | 1927 | 2473 | 2399 | 2459 |
| ('rep1', 'PML') | 1511 | 1964 | 1468 | 2499 | 3145 | 2808 | 2444 |
| ('rep2', 'PML') | 1673 | 2266 | 2108 | 2764 | 2814 | 2854 | 2451 |
| ('rep1', 'PSD95') | 1995 | 1976 | 1089 | 2282 | 2849 | 2983 | 2927 |
| ('rep2', 'PSD95') | 1655 | 2225 | 947 | 2363 | 2780 | 3068 | 2813 |
| ('rep1', 'PURA') | 1560 | 1234 | 1109 | 1183 | 1363 | 1507 | 1443 |
| ('rep2', 'PURA') | 1155 | 1646 | 770 | 1685 | 1693 | 1799 | 1222 |
| ('rep1', 'Phalloidin') | 1331 | 2038 | 948 | 2140 | 2654 | 2822 | 2757 |
| ('rep2', 'Phalloidin') | 1239 | 1764 | 879 | 2309 | 2496 | 2995 | 2683 |
| ('rep1', 'SNCA') | 1440 | 871 | 1060 | 1319 | 1565 | 1531 | 1060 |
| ('rep2', 'SNCA') | 1204 | 1289 | 929 | 1083 | 1411 | 1068 | 1330 |
| ('rep1', 'SON') | 1991 | 2350 | 1196 | 1986 | 2769 | 3065 | 2439 |
| ('rep2', 'SON') | 2189 | 2494 | 1397 | 2104 | 2624 | 2534 | 2560 |
| ('rep1', 'SQSTM1') | 1395 | 1652 | 1426 | 1829 | 2050 | 2464 | 2253 |
| ('rep2', 'SQSTM1') | 1335 | 1399 | 1067 | 1881 | 1739 | 2241 | 2327 |
| ('rep1', 'TDP43') | 1381 | 1830 | 1476 | 2207 | 2421 | 2611 | 2380 |
| ('rep2', 'TDP43') | 1645 | 1871 | 1964 | 2429 | 2022 | 2531 | 2305 |
| ('rep1', 'TIA1') | 2154 | 2420 | 2015 | 2119 | 2543 | 2296 | 2034 |
| ('rep2', 'TIA1') | 1854 | 2082 | 1217 | 2119 | 2350 | 2348 | 1865 |
| ('rep1', 'TOMM20') | 2161 | 2054 | 2080 | 2096 | 2609 | 2456 | 2342 |
| ('rep2', 'TOMM20') | 1656 | 2175 | 1280 | 2182 | 2570 | 2384 | 2060 |
| ('rep1', 'Tubulin') | 1210 | 1595 | 1280 | 1889 | 2146 | 2797 | 2401 |
| ('rep2', 'Tubulin') | 1272 | 1294 | 1053 | 1995 | 1792 | 2648 | 2533 |
| ('rep1', 'mitotracker') | 2008 | 1382 | 1038 | 2078 | 2403 | 2297 | 2268 |
| ('rep2', 'mitotracker') | 1503 | 2235 | 1039 | 2286 | 2509 | 2484 | 2098 |

Total number of tiles

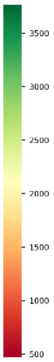

### Batch 2

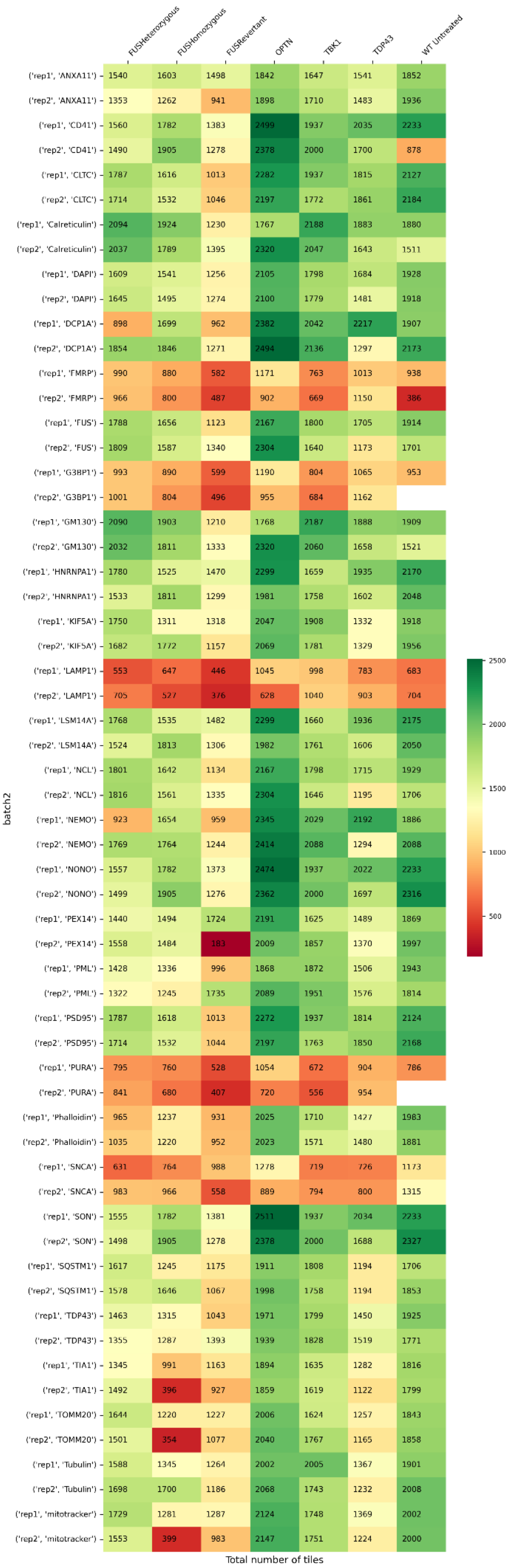

FUSHeterozygous
FUSHomozygous
FUSRevertant
OPTN
TBK1
TDP43
WT Untreated
('rep1', 'ANXA11') 1540 1603 1498 1842 1647 1541 1852
('rep2', 'ANXA11') 1353 1262 941 1898 1710 1483 1936
('rep1', 'CD41') 1560 1782 1383 2499 1937 2035 2233
('rep2', 'CD41') 1490 1905 1278 2378 2000 1700 878
('rep1', 'CLTC') 1787 1616 1013 2282 1937 1815 2127
('rep2', 'CLTC') 1714 1532 1046 2197 1772 1861 2184
('rep1', 'Calreticulin') 2094 1924 1230 1767 2188 1883 1880
('rep2', 'Calreticulin') 2037 1789 1395 2320 2047 1643 1511
('rep1', 'DAPI') 1609 1541 1256 2105 1798 1684 1928
('rep2', 'DAPI') 1645 1495 1274 2100 1779 1481 1918
('rep1', 'DCP1A') 898 1699 962 2382 2042 2217 1907
('rep2', 'DCP1A') 1854 1846 1271 2494 2136 1297 2173
('rep1', 'FMRP') 990 880 582 1171 763 1013 938
('rep2', 'FMRP') 966 800 487 902 669 1150 386
('rep1', 'FUS') 1788 1656 1123 2167 1800 1705 1914
('rep2', 'FUS') 1809 1587 1340 2304 1640 1173 1701
('rep1', 'G3BP1') 993 890 599 1190 804 1065 953
('rep2', 'G3BP1') 1001 804 496 955 684 1162
('rep1', 'GM130') 2090 1903 1210 1768 2187 1888 1909
('rep2', 'GM130') 2032 1811 1333 2320 2060 1658 1521
('rep1', 'HNRNPA1') 1780 1525 1470 2299 1659 1935 2170
('rep2', 'HNRNPA1') 1533 1811 1299 1981 1758 1602 2048
('rep1', 'KIF5A') 1750 1311 1318 2047 1908 1332 1918
('rep2', 'KIF5A') 1682 1772 1157 2069 1781 1329 1956
('rep1', 'LAMP1') 553 647 446 1045 998 783 683
('rep2', 'LAMP1') 705 527 376 628 1040 903 704
('rep1', 'LSM14A') 1768 1535 1482 2299 1660 1936 2175
('rep2', 'LSM14A') 1524 1813 1306 1982 1761 1606 2050
('rep1', 'NCL') 1801 1642 1134 2167 1798 1715 1929
('rep2', 'NCL') 1816 1561 1335 2304 1646 1195 1706
('rep1', 'NEMO') 923 1654 959 2345 2029 2192 1886
('rep2', 'NEMO') 1769 1764 1244 2414 2088 1294 2088
('rep1', 'NONO') 1557 1782 1373 2474 1937 2022 2233
('rep2', 'NONO') 1499 1905 1276 2362 2000 1697 2316
('rep1', 'PEX14') 1440 1494 1724 2191 1625 1489 1869
('rep2', 'PEX14') 1558 1484 183 2009 1857 1370 1997
('rep1', 'PML') 1428 1336 996 1868 1872 1506 1943
('rep2', 'PML') 1322 1245 1735 2089 1951 1576 1814
('rep1', 'PSD95') 1787 1618 1013 2272 1937 1814 2124
('rep2', 'PSD95') 1714 1532 1044 2197 1763 1850 2168
('rep1', 'PURA') 795 760 528 1054 672 904 786
('rep2', 'PURA') 841 680 407 720 556 954
('rep1', 'Phalloidin') 965 1237 931 2025 1710 1427 1983
('rep2', 'Phalloidin') 1035 1220 952 2023 1571 1480 1881
('rep1', 'SNCA') 631 764 988 1278 719 726 1173
('rep2', 'SNCA') 983 966 558 889 794 800 1315
('rep1', 'SON') 1555 1782 1381 2511 1937 2034 2233
('rep2', 'SON') 1498 1905 1278 2378 2000 1688 2327
('rep1', 'SQSTM1') 1617 1245 1175 1911 1808 1194 1706
('rep2', 'SQSTM1') 1578 1646 1067 1998 1758 1194 1853
('rep1', 'TDP43') 1463 1315 1043 1971 1799 1450 1925
('rep2', 'TDP43') 1355 1287 1393 1939 1828 1519 1771
('rep1', 'TIA1') 1345 991 1163 1894 1635 1282 1816
('rep2', 'TIA1') 1492 396 927 1859 1619 1122 1799
('rep1', 'TOMM20') 1644 1220 1227 2006 1624 1257 1843
('rep2', 'TOMM20') 1501 354 1077 2040 1767 1165 1858
('rep1', 'Tubulin') 1588 1345 1264 2002 2005 1367 1901
('rep2', 'Tubulin') 1698 1700 1186 2068 1743 1232 2008
('rep1', 'mitotracker') 1729 1281 1287 2124 1748 1369 2002
('rep2', 'mitotracker') 1553 399 983 2147 1751 1224 2000
batch2
Total number of tiles
2500
2000
1500
1000
500

### Batch 3

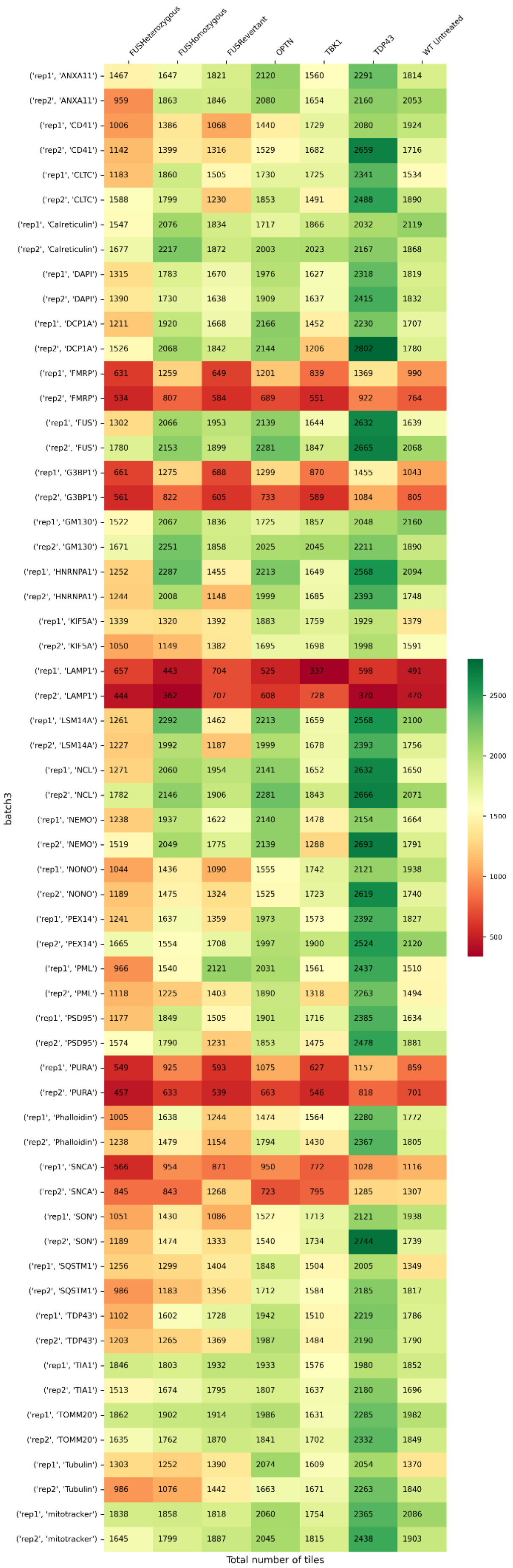

batch3
Total number of tiles
2500
2000
1500
1000
500

| batch3 | FUSHeterozygous | FUSHomozygous | FUSRevertant | OPTN | TBK1 | TDP43 | WT Untreated |
|---|---|---|---|---|---|---|---|
| ('rep1', 'ANXA11') | 1467 | 1647 | 1821 | 2120 | 1560 | 2291 | 1814 |
| ('rep2', 'ANXA11') | 959 | 1863 | 1846 | 2080 | 1654 | 2160 | 2053 |
| ('rep1', 'CD41') | 1006 | 1386 | 1068 | 1440 | 1729 | 2080 | 1924 |
| ('rep2', 'CD41') | 1142 | 1399 | 1316 | 1529 | 1682 | 2659 | 1716 |
| ('rep1', 'CLTC') | 1183 | 1860 | 1505 | 1730 | 1725 | 2341 | 1534 |
| ('rep2', 'CLTC') | 1588 | 1799 | 1230 | 1853 | 1491 | 2488 | 1890 |
| ('rep1', 'Calreticulin') | 1547 | 2076 | 1834 | 1717 | 1866 | 2032 | 2119 |
| ('rep2', 'Calreticulin') | 1677 | 2217 | 1872 | 2003 | 2023 | 2167 | 1868 |
| ('rep1', 'DAPI') | 1315 | 1783 | 1670 | 1976 | 1627 | 2318 | 1819 |
| ('rep2', 'DAPI') | 1390 | 1730 | 1638 | 1909 | 1637 | 2415 | 1832 |
| ('rep1', 'DCP1A') | 1211 | 1920 | 1668 | 2166 | 1452 | 2230 | 1707 |
| ('rep2', 'DCP1A') | 1526 | 2068 | 1842 | 2144 | 1206 | 2802 | 1780 |
| ('rep1', 'FMRP') | 631 | 1259 | 649 | 1201 | 839 | 1369 | 990 |
| ('rep2', 'FMRP') | 534 | 807 | 584 | 689 | 551 | 922 | 764 |
| ('rep1', 'FUS') | 1302 | 2066 | 1953 | 2139 | 1644 | 2632 | 1639 |
| ('rep2', 'FUS') | 1780 | 2153 | 1899 | 2281 | 1847 | 2665 | 2068 |
| ('rep1', 'G3BP1') | 661 | 1275 | 688 | 1299 | 870 | 1455 | 1043 |
| ('rep2', 'G3BP1') | 561 | 822 | 605 | 733 | 589 | 1084 | 805 |
| ('rep1', 'GM130') | 1522 | 2067 | 1836 | 1725 | 1857 | 2048 | 2160 |
| ('rep2', 'GM130') | 1671 | 2251 | 1858 | 2025 | 2045 | 2211 | 1890 |
| ('rep1', 'HNRNPA1') | 1252 | 2287 | 1455 | 2213 | 1649 | 2568 | 2094 |
| ('rep2', 'HNRNPA1') | 1244 | 2008 | 1148 | 1999 | 1685 | 2393 | 1748 |
| ('rep1', 'KIF5A') | 1339 | 1320 | 1392 | 1883 | 1759 | 1929 | 1379 |
| ('rep2', 'KIF5A') | 1050 | 1149 | 1382 | 1695 | 1698 | 1998 | 1591 |
| ('rep1', 'LAMP1') | 657 | 443 | 704 | 525 | 337 | 598 | 491 |
| ('rep2', 'LAMP1') | 444 | 362 | 707 | 608 | 728 | 370 | 470 |
| ('rep1', 'LSM14A') | 1261 | 2292 | 1462 | 2213 | 1659 | 2568 | 2100 |
| ('rep2', 'LSM14A') | 1227 | 1992 | 1187 | 1999 | 1678 | 2393 | 1756 |
| ('rep1', 'NCL') | 1271 | 2060 | 1954 | 2141 | 1652 | 2632 | 1650 |
| ('rep2', 'NCL') | 1782 | 2146 | 1906 | 2281 | 1843 | 2666 | 2071 |
| ('rep1', 'NEMO') | 1238 | 1937 | 1622 | 2140 | 1478 | 2154 | 1664 |
| ('rep2', 'NEMO') | 1519 | 2049 | 1775 | 2139 | 1288 | 2693 | 1791 |
| ('rep1', 'NONO') | 1044 | 1436 | 1090 | 1555 | 1742 | 2121 | 1938 |
| ('rep2', 'NONO') | 1189 | 1475 | 1324 | 1525 | 1723 | 2619 | 1740 |
| ('rep1', 'PEX14') | 1241 | 1637 | 1359 | 1973 | 1573 | 2392 | 1827 |
| ('rep2', 'PEX14') | 1665 | 1554 | 1708 | 1997 | 1900 | 2524 | 2120 |
| ('rep1', 'PML') | 966 | 1540 | 2121 | 2031 | 1561 | 2437 | 1510 |
| ('rep2', 'PML') | 1118 | 1225 | 1403 | 1890 | 1318 | 2263 | 1494 |
| ('rep1', 'PSD95') | 1177 | 1849 | 1505 | 1901 | 1716 | 2385 | 1634 |
| ('rep2', 'PSD95') | 1574 | 1790 | 1231 | 1853 | 1475 | 2478 | 1881 |
| ('rep1', 'PURA') | 549 | 925 | 593 | 1075 | 627 | 1157 | 859 |
| ('rep2', 'PURA') | 457 | 633 | 539 | 663 | 546 | 818 | 701 |
| ('rep1', 'Phalloidin') | 1005 | 1638 | 1244 | 1474 | 1564 | 2280 | 1772 |
| ('rep2', 'Phalloidin') | 1238 | 1479 | 1154 | 1794 | 1430 | 2367 | 1805 |
| ('rep1', 'SNCA') | 566 | 954 | 871 | 950 | 772 | 1028 | 1116 |
| ('rep2', 'SNCA') | 845 | 843 | 1268 | 723 | 795 | 1285 | 1307 |
| ('rep1', 'SON') | 1051 | 1430 | 1086 | 1527 | 1713 | 2121 | 1938 |
| ('rep2', 'SON') | 1189 | 1474 | 1333 | 1540 | 1734 | 2744 | 1739 |
| ('rep1', 'SQSTM1') | 1256 | 1299 | 1404 | 1848 | 1504 | 2005 | 1349 |
| ('rep2', 'SQSTM1') | 986 | 1183 | 1356 | 1712 | 1584 | 2185 | 1817 |
| ('rep1', 'TDP43') | 1102 | 1602 | 1728 | 1942 | 1510 | 2219 | 1786 |
| ('rep2', 'TDP43') | 1203 | 1265 | 1369 | 1987 | 1484 | 2190 | 1790 |
| ('rep1', 'TIA1') | 1846 | 1803 | 1932 | 1933 | 1576 | 1980 | 1852 |
| ('rep2', 'TIA1') | 1513 | 1674 | 1795 | 1807 | 1637 | 2180 | 1696 |
| ('rep1', 'TOMM20') | 1862 | 1902 | 1914 | 1986 | 1631 | 2285 | 1982 |
| ('rep2', 'TOMM20') | 1635 | 1762 | 1870 | 1841 | 1702 | 2332 | 1849 |
| ('rep1', 'Tubulin') | 1303 | 1252 | 1390 | 2074 | 1609 | 2054 | 1370 |
| ('rep2', 'Tubulin') | 986 | 1076 | 1442 | 1663 | 1671 | 2263 | 1840 |
| ('rep1', 'mitotracker') | 1838 | 1858 | 1818 | 2060 | 1754 | 2365 | 2086 |
| ('rep2', 'mitotracker') | 1645 | 1799 | 1887 | 2045 | 1815 | 2438 | 1903 |

### Batch 4

| | FUSHeterozygous | FUSHomozygous | FUSRevertant | OPTN | TBK1 | TDP43 | WT Untreated |
|---|---|---|---|---|---|---|---|
| ('rep1', 'ANXA11') | 1717 | 2127 | 2158 | 1973 | 954 | 1278 | 2423 |
| ('rep2', 'ANXA11') | 2261 | 2048 | 2014 | 1873 | 1562 | 1977 | 2367 |
| ('rep1', 'CD41') | 2983 | 2499 | 3035 | 2535 | 93 | 2555 | 2202 |
| ('rep2', 'CD41') | 3377 | 2898 | 3040 | 2392 | 2199 | 2353 | 3529 |
| ('rep1', 'CLTC') | 2625 | 2739 | 2156 | 2521 | 1613 | 2538 | 3219 |
| ('rep2', 'CLTC') | 1496 | 1935 | 1708 | 2298 | 2084 | 2376 | 2952 |
| ('rep1', 'Calreticulin') | 2535 | 1925 | 2567 | 2247 | 1953 | 1999 | 2934 |
| ('rep2', 'Calreticulin') | 2755 | 2309 | 2278 | 2496 | 2017 | 2046 | 3065 |
| ('rep1', 'DAPI') | 2496 | 2100 | 2456 | 2090 | 1518 | 1866 | 2690 |
| ('rep2', 'DAPI') | 2551 | 2163 | 2387 | 2104 | 1822 | 2044 | 2829 |
| ('rep1', 'DCP1A') | 2685 | 2287 | 2503 | 2053 | 1814 | 2400 | 2767 |
| ('rep2', 'DCP1A') | 2611 | 2365 | 2740 | 2365 | 2152 | 2216 | 3029 |
| ('rep1', 'FMRP') | 821 | 592 | 510 | 693 | 643 | 557 | 950 |
| ('rep2', 'FMRP') | 955 | 714 | 603 | 595 | 266 | 516 | 776 |
| ('rep1', 'FUS') | 20 | 27 | 63 | | 7 | 28 | 19 |
| ('rep2', 'FUS') | 8 | 5 | 6 | 5 | 30 | | 27 |
| ('rep1', 'G3BP1') | 914 | 615 | 535 | 735 | 605 | 583 | 968 |
| ('rep2', 'G3BP1') | 1310 | 819 | 662 | 611 | 306 | 615 | 821 |
| ('rep1', 'GM130') | 2848 | 1992 | 2717 | 2268 | 1976 | 1999 | 3037 |
| ('rep2', 'GM130') | 3077 | 2336 | 2582 | 2510 | 2023 | 1961 | 3059 |
| ('rep1', 'HNRNPA1') | 2469 | 2347 | 2581 | 2452 | 2162 | 2065 | 2542 |
| ('rep2', 'HNRNPA1') | 2226 | 2137 | 2311 | 1993 | 1701 | 1914 | 2524 |
| ('rep1', 'KIF5A') | 2434 | 2376 | 1750 | 1911 | 1588 | 1279 | 2708 |
| ('rep2', 'KIF5A') | 2206 | 2320 | 1827 | 2214 | 2256 | 2307 | 2865 |
| ('rep1', 'LAMP1') | 719 | 1170 | 1665 | 1192 | 602 | 817 | 1282 |
| ('rep2', 'LAMP1') | 1313 | 1041 | 1385 | 1464 | 1367 | 1023 | 855 |
| ('rep1', 'LSM14A') | 2831 | 2344 | 2836 | 2500 | 2174 | 2093 | 2810 |
| ('rep2', 'LSM14A') | 2350 | 2149 | 2513 | 1998 | 1729 | 1956 | 2896 |
| ('rep1', 'NCL') | 2422 | 2046 | 2660 | 2168 | 2222 | 2135 | 3118 |
| ('rep2', 'NCL') | 2639 | 2186 | 2805 | 2185 | 1940 | 2264 | 3062 |
| ('rep1', 'NEMO') | 1785 | 1877 | 2213 | 1975 | 1738 | 2232 | 2108 |
| ('rep2', 'NEMO') | 1677 | 2104 | 2386 | 2253 | 2072 | 2068 | 2270 |
| ('rep1', 'NONO') | 1971 | 2104 | 1505 | 2310 | 65 | 2438 | 1475 |
| ('rep2', 'NONO') | 1705 | 2534 | 2506 | 2344 | 1939 | 2296 | 3167 |
| ('rep1', 'PEX14') | 1306 | 1111 | 1666 | 2062 | 855 | 1987 | 2285 |
| ('rep2', 'PEX14') | 2099 | 1969 | 1652 | 2174 | 1522 | 2043 | 1726 |
| ('rep1', 'PML') | 2254 | 1897 | 2005 | 1696 | 1587 | 1750 | 2455 |
| ('rep2', 'PML') | 2185 | 2202 | 2763 | 1982 | 1857 | 1952 | 2497 |
| ('rep1', 'PSD95') | 2226 | 2643 | 2147 | 2525 | 1624 | 2538 | 2195 |
| ('rep2', 'PSD95') | 776 | 1855 | 1708 | 2303 | 2072 | 2371 | 2870 |
| ('rep1', 'PURA') | 802 | 558 | 650 | 660 | 564 | 537 | 958 |
| ('rep2', 'PURA') | 1136 | 705 | 611 | 595 | 293 | 574 | 1210 |
| ('rep1', 'Phalloidin') | 2518 | 2601 | 2119 | 2493 | 1609 | 2383 | 2959 |
| ('rep2', 'Phalloidin') | 2223 | 1816 | 1666 | 2254 | 2023 | 2259 | 2586 |
| ('rep1', 'SNCA') | 966 | 666 | 842 | 386 | 646 | 847 | 1001 |
| ('rep2', 'SNCA') | 1573 | 1084 | 1313 | 1116 | 989 | 1124 | 1187 |
| ('rep1', 'SON') | 2619 | 2333 | 2245 | 2358 | 77 | 2477 | 2146 |
| ('rep2', 'SON') | 2882 | 2827 | 2819 | 2362 | 2166 | 2304 | 3433 |
| ('rep1', 'SQSTM1') | 1961 | 2055 | 1585 | 1704 | 1384 | 1143 | 2392 |
| ('rep2', 'SQSTM1') | 1596 | 1812 | 1486 | 1953 | 2079 | 2111 | 2745 |
| ('rep1', 'TDP43') | 1753 | 1800 | 1219 | 1664 | 1577 | 1681 | 2442 |
| ('rep2', 'TDP43') | 1621 | 2001 | 1425 | 1896 | 1847 | 1814 | 2145 |
| ('rep1', 'TIA1') | 1893 | 1958 | 2456 | 2169 | 1497 | 1145 | 1848 |
| ('rep2', 'TIA1') | 2100 | 1585 | 2245 | 1682 | 1301 | 1432 | 1391 |
| ('rep1', 'TOMM20') | 2303 | 2246 | 2611 | 2248 | 2019 | 1265 | 2555 |
| ('rep2', 'TOMM20') | 2518 | 1910 | 2345 | 2086 | 1850 | 1849 | 2466 |
| ('rep1', 'Tubulin') | 2610 | 2789 | 2077 | 1872 | 1685 | 1438 | 3370 |
| ('rep2', 'Tubulin') | 2287 | 2699 | 2032 | 2161 | 2319 | 2330 | 3299 |
| ('rep1', 'mitotracker') | 2358 | 2249 | 2540 | 2235 | 1993 | 1201 | 2488 |
| ('rep2', 'mitotracker') | 2483 | 1909 | 2366 | 2019 | 1813 | 1764 | 2420 |

Total number of tiles

Color scale: 500, 1000, 1500, 2000, 2500, 3000, 3500

### Batch 5

| | FUSHeterozygous | FUSHomozygous | FUSRevertant | OPTN | TBK1 | TDP43 | WT Untreated |
|---|---|---|---|---|---|---|---|
| ('rep1', 'ANXA11') | 2174 | 2177 | 2534 | 2093 | 1535 | 1995 | 2034 |
| ('rep2', 'ANXA11') | 2190 | 2414 | 2697 | 2047 | 1490 | 2111 | 2037 |
| ('rep1', 'CD41') | 2950 | 2595 | 2642 | 2674 | 2162 | 2609 | 2718 |
| ('rep2', 'CD41') | 2510 | 2692 | 3008 | 2142 | 1997 | 2387 | 2647 |
| ('rep1', 'CLTC') | 2860 | 2494 | 2600 | 2365 | 1891 | 2288 | 2262 |
| ('rep2', 'CLTC') | 2655 | 2498 | 2418 | 2441 | 2053 | 2259 | 1688 |
| ('rep1', 'Calreticulin') | 3344 | 3216 | 2928 | 2856 | 2212 | 2684 | 3003 |
| ('rep2', 'Calreticulin') | 3328 | 3048 | 3458 | 2644 | 2294 | 2778 | 2871 |
| ('rep1', 'DAPI') | 2698 | 2483 | 2645 | 2281 | 1816 | 2270 | 2375 |
| ('rep2', 'DAPI') | 2634 | 2485 | 2704 | 2130 | 1767 | 2234 | 2347 |
| ('rep1', 'DCP1A') | 3090 | 2659 | 2612 | 2580 | 2171 | 2760 | 2948 |
| ('rep2', 'DCP1A') | 3218 | 2750 | 2933 | 2640 | 2349 | 2569 | 2823 |
| ('rep1', 'FMRP') | 866 | 995 | 626 | 846 | 440 | 897 | 1123 |
| ('rep2', 'FMRP') | 719 | 859 | 545 | 757 | 398 | 766 | 810 |
| ('rep1', 'FUS') | 3225 | 2962 | 3016 | 2694 | 2321 | 2841 | 2930 |
| ('rep2', 'FUS') | 3310 | 2970 | 3230 | 2409 | 2068 | 2493 | 2920 |
| ('rep1', 'G3BP1') | 1056 | 1047 | 677 | 862 | 446 | 981 | 1266 |
| ('rep2', 'G3BP1') | 821 | 953 | 602 | 764 | 419 | 838 | 901 |
| ('rep1', 'GM130') | 3344 | 3216 | 2928 | 2842 | 2212 | 2701 | 3003 |
| ('rep2', 'GM130') | 3332 | 3048 | 3458 | 2631 | 2308 | 2787 | 2871 |
| ('rep1', 'HNRNPA1') | 2534 | 2694 | 2541 | 2616 | 2132 | 2815 | 2689 |
| ('rep2', 'HNRNPA1') | 2293 | 2743 | 2223 | 2377 | 2005 | 2574 | 2821 |
| ('rep1', 'KIF5A') | 2198 | 2087 | 2190 | 1873 | 1620 | 1374 | 1703 |
| ('rep2', 'KIF5A') | 2371 | 2122 | 2572 | 1556 | 1387 | 2054 | 2040 |
| ('rep1', 'LAMP1') | 1260 | 1493 | 1753 | 1084 | 1094 | 899 | 1209 |
| ('rep2', 'LAMP1') | 1351 | 1274 | 1543 | 1388 | 1331 | 1408 | 1537 |
| ('rep1', 'LSM14A') | 2681 | 2834 | 2923 | 2632 | 2132 | 2830 | 2722 |
| ('rep2', 'LSM14A') | 2666 | 2813 | 2685 | 2390 | 2013 | 2609 | 2849 |
| ('rep1', 'NCL') | 3225 | 2944 | 2983 | 2694 | 2322 | 2816 | 2930 |
| ('rep2', 'NCL') | 3265 | 2976 | 3048 | 2409 | 2068 | 2493 | 2920 |
| ('rep1', 'NEMO') | 3075 | 2642 | 2601 | 2558 | 2132 | 2760 | 2948 |
| ('rep2', 'NEMO') | 3218 | 2743 | 2948 | 2555 | 2321 | 2547 | 2823 |
| ('rep1', 'NONO') | 2226 | 2492 | 2338 | 2351 | 2152 | 2318 | 2379 |
| ('rep2', 'NONO') | 2224 | 2623 | 2573 | 2108 | 1971 | 2213 | 2340 |
| ('rep1', 'PEX14') | 2093 | 2286 | 1790 | 1916 | 1702 | 2141 | 2008 |
| ('rep2', 'PEX14') | 2156 | 1972 | 2268 | 1760 | 1688 | 2144 | 2067 |
| ('rep1', 'PML') | 1734 | 2168 | 2387 | 2074 | 1208 | 1749 | 1406 |
| ('rep2', 'PML') | 2418 | 2275 | 2118 | 2313 | 1250 | 1872 | 1969 |
| ('rep1', 'PSD95') | 2916 | 2540 | 2609 | 2356 | 1887 | 2278 | 2074 |
| ('rep2', 'PSD95') | 2639 | 2498 | 2398 | 2427 | 2053 | 2249 | 190 |
| ('rep1', 'PURA') | 879 | 820 | 616 | 779 | 418 | 887 | 1040 |
| ('rep2', 'PURA') | 706 | 756 | 554 | 660 | 349 | 689 | 780 |
| ('rep1', 'Phalloidin') | 2585 | 2487 | 2474 | 2341 | 1746 | 2089 | 2285 |
| ('rep2', 'Phalloidin') | 2331 | 2273 | 2250 | 2401 | 1869 | 2152 | 1799 |
| ('rep1', 'SNCA') | 1281 | 872 | 2241 | 955 | 710 | 1039 | 703 |
| ('rep2', 'SNCA') | 1416 | 1045 | 1183 | 847 | 1190 | 1540 | 844 |
| ('rep1', 'SON') | 2770 | 2610 | 2598 | 2533 | 2162 | 2608 | 2707 |
| ('rep2', 'SON') | 2407 | 2692 | 2898 | 2100 | 1988 | 2323 | 2570 |
| ('rep1', 'SQSTM1') | 1839 | 1728 | 1717 | 1670 | 1571 | 1164 | 1452 |
| ('rep2', 'SQSTM1') | 1971 | 2006 | 2159 | 1447 | 1391 | 1802 | 1871 |
| ('rep1', 'TDP43') | 1686 | 1988 | 1090 | 1935 | 1281 | 1740 | 1690 |
| ('rep2', 'TDP43') | 2296 | 2209 | 1710 | 2177 | 1428 | 2074 | 2081 |
| ('rep1', 'TIA1') | 2397 | 2004 | 2040 | 1880 | 1365 | 1680 | 1741 |
| ('rep2', 'TIA1') | 2062 | 2083 | 2781 | 1595 | 1294 | 1603 | 1249 |
| ('rep1', 'TOMM20') | 2160 | 2048 | 1884 | 2002 | 1728 | 2021 | 1972 |
| ('rep2', 'TOMM20') | 1985 | 2280 | 2460 | 1782 | 1470 | 1996 | 1837 |
| ('rep1', 'Tubulin') | 2772 | 2735 | 2811 | 2235 | 2075 | 1923 | 2372 |
| ('rep2', 'Tubulin') | 2613 | 2650 | 2817 | 1934 | 1635 | 2121 | 2506 |
| ('rep1', 'mitotracker') | 2892 | 2156 | 2039 | 2012 | 1728 | 2052 | 1856 |
| ('rep2', 'mitotracker') | 2329 | 2446 | 2822 | 1753 | 1413 | 1930 | 1844 |

Total number of tiles

### <u>Batch 6</u>

| | FUSHeterozygous | FUSHomozygous | FUSRevertant | OPTN | TBK1 | TDP43 | WT Untreated |
|---|---|---|---|---|---|---|---|
| ('rep1', 'ANXA11') | 600 | 348 | 899 | 1186 | 308 | 1669 | 1520 |
| ('rep2', 'ANXA11') | 450 | 304 | 403 | 401 | 680 | 938 | 1551 |
| ('rep1', 'CD41') | 3240 | 2428 | 2866 | 2883 | 2135 | 2355 | 4 |
| ('rep2', 'CD41') | 3174 | 2547 | 2938 | 3119 | 1959 | 2345 | 3325 |
| ('rep1', 'CLTC') | 2989 | 2526 | 2350 | 2708 | 2068 | 2486 | 3151 |
| ('rep2', 'CLTC') | 2758 | 2570 | 2173 | 2816 | 2254 | 2538 | 3221 |
| ('rep1', 'Calreticulin') | 3020 | 2312 | 2884 | | 2030 | 2766 | 3330 |
| ('rep2', 'Calreticulin') | 3201 | 2528 | 2966 | 3030 | 2268 | 2503 | 3314 |
| ('rep1', 'DAPI') | 2305 | 1814 | 2109 | 2184 | 1714 | 2196 | 2616 |
| ('rep2', 'DAPI') | 2271 | 1934 | 2113 | 2404 | 1877 | 2115 | 2799 |
| ('rep1', 'DCP1A') | 2763 | 2245 | 2401 | 2856 | 2171 | 2479 | 3475 |
| ('rep2', 'DCP1A') | 3027 | 2458 | 2505 | 3205 | 2329 | 2660 | 3413 |
| ('rep1', 'FMRP') | 1240 | 1058 | 848 | 1373 | 780 | 1039 | 1148 |
| ('rep2', 'FMRP') | 1581 | 1238 | 769 | 1261 | 690 | 1255 | 1205 |
| ('rep1', 'FUS') | 3130 | 2422 | 3017 | 2825 | 2568 | 2524 | 3410 |
| ('rep2', 'FUS') | 3131 | 2486 | 3026 | 2857 | 2452 | 2512 | 3028 |
| ('rep1', 'G3BP1') | 1289 | 1071 | 873 | 1346 | 791 | 1058 | 1198 |
| ('rep2', 'G3BP1') | 1625 | 1252 | 824 | 1258 | 722 | 1266 | 1497 |
| ('rep1', 'GM130') | 3063 | 2324 | 2923 | | 2030 | 2766 | 3330 |
| ('rep2', 'GM130') | 3230 | 2528 | 2976 | 3022 | 2268 | 2515 | 3336 |
| ('rep1', 'HNRNPA1') | 2734 | 2522 | 2377 | 3040 | 2177 | 2413 | 3005 |
| ('rep2', 'HNRNPA1') | 2720 | 2445 | 1972 | 2763 | 2458 | 2477 | 2906 |
| ('rep1', 'KIF5A') | 2660 | 1980 | 2684 | 2526 | 1352 | 2073 | 2542 |
| ('rep2', 'KIF5A') | 2917 | 1965 | 2477 | 2268 | 1703 | 1934 | 2550 |
| ('rep1', 'LAMP1') | 581 | 321 | 1215 | 1776 | 928 | 2043 | 1351 |
| ('rep2', 'LAMP1') | 387 | 305 | 823 | 774 | 897 | 1035 | 1287 |
| ('rep1', 'LSM14A') | 2972 | 2586 | 2377 | 3096 | 2194 | 2441 | 3104 |
| ('rep2', 'LSM14A') | 2979 | 2473 | 2261 | 2763 | 2470 | 2492 | 3059 |
| ('rep1', 'NCL') | 3130 | 2436 | 2981 | 2825 | 2568 | 2524 | 3410 |
| ('rep2', 'NCL') | 3057 | 2497 | 2914 | 2857 | 2452 | 2491 | 3011 |
| ('rep1', 'NEMO') | 2752 | 2245 | 2396 | 2832 | 2162 | 2494 | 3448 |
| ('rep2', 'NEMO') | 3013 | 2448 | 2489 | 3205 | 2315 | 2660 | 3388 |
| ('rep1', 'NONO') | 3117 | 2440 | 2483 | 2897 | 2122 | 2312 | |
| ('rep2', 'NONO') | 3018 | 2547 | 2624 | 3078 | 1935 | 2339 | 3325 |
| ('rep1', 'PEX14') | 533 | 744 | 929 | 1209 | 1015 | 1385 | 1990 |
| ('rep2', 'PEX14') | 716 | 612 | 1337 | 1535 | 1233 | 1177 | 2256 |
| ('rep1', 'PML') | 355 | 545 | 165 | 1275 | 756 | 1977 | 2351 |
| ('rep2', 'PML') | 1561 | 1611 | 1322 | 1359 | 1318 | 1635 | 2498 |
| ('rep1', 'PSD95') | 2985 | 2513 | 2328 | 2708 | 2070 | 2449 | 3144 |
| ('rep2', 'PSD95') | 2689 | 2552 | 2155 | 2744 | 2255 | 2541 | 3213 |
| ('rep1', 'PURA') | 1136 | 889 | 729 | 1155 | 680 | 933 | 955 |
| ('rep2', 'PURA') | 1193 | 978 | 724 | 1169 | 611 | 923 | 1359 |
| ('rep1', 'Phalloidin') | 2574 | 2111 | 2247 | 2647 | 1938 | 2318 | 2750 |
| ('rep2', 'Phalloidin') | 2420 | 2302 | 2104 | 2726 | 1924 | 2436 | 2540 |
| ('rep1', 'SNCA') | 524 | 437 | 670 | 1041 | 704 | 886 | 1203 |
| ('rep2', 'SNCA') | 603 | 324 | 1120 | 1095 | 770 | 596 | 1420 |
| ('rep1', 'SON') | 3206 | 2428 | 2749 | 2878 | 2135 | 2349 | 4 |
| ('rep2', 'SON') | 3100 | 2547 | 2847 | 3081 | 1941 | 2324 | 3305 |
| ('rep1', 'SQSTM1') | 2225 | 1632 | 2266 | 1730 | 1037 | 1718 | 1938 |
| ('rep2', 'SQSTM1') | 2355 | 1648 | 1904 | 1803 | 1465 | 1552 | 2162 |
| ('rep1', 'TDP43') | 304 | 551 | 136 | 1073 | 695 | 1834 | 2056 |
| ('rep2', 'TDP43') | 1027 | 1584 | 1229 | 1412 | 1256 | 1591 | 2110 |
| ('rep1', 'TIA1') | 1731 | 684 | 1609 | 1936 | 857 | 1421 | 1721 |
| ('rep2', 'TIA1') | 781 | 1064 | 817 | 2256 | 1518 | 1652 | 1428 |
| ('rep1', 'TOMM20') | 2493 | 1442 | 1800 | 2020 | 1603 | 1848 | 2713 |
| ('rep2', 'TOMM20') | 814 | 1680 | 846 | 2425 | 2048 | 2329 | 2559 |
| ('rep1', 'Tubulin') | 3235 | 2476 | 2940 | 2965 | 1707 | 2274 | 3159 |
| ('rep2', 'Tubulin') | 3146 | 2134 | 2691 | 2731 | 1940 | 1797 | 2877 |
| ('rep1', 'mitotracker') | 2764 | 1417 | 1807 | 2072 | 1579 | 1818 | 2735 |
| ('rep2', 'mitotracker') | 936 | 1645 | 846 | 2313 | 2001 | 2254 | 2543 |

Total number of tiles

### Whole Cell Count

#### Batch 1

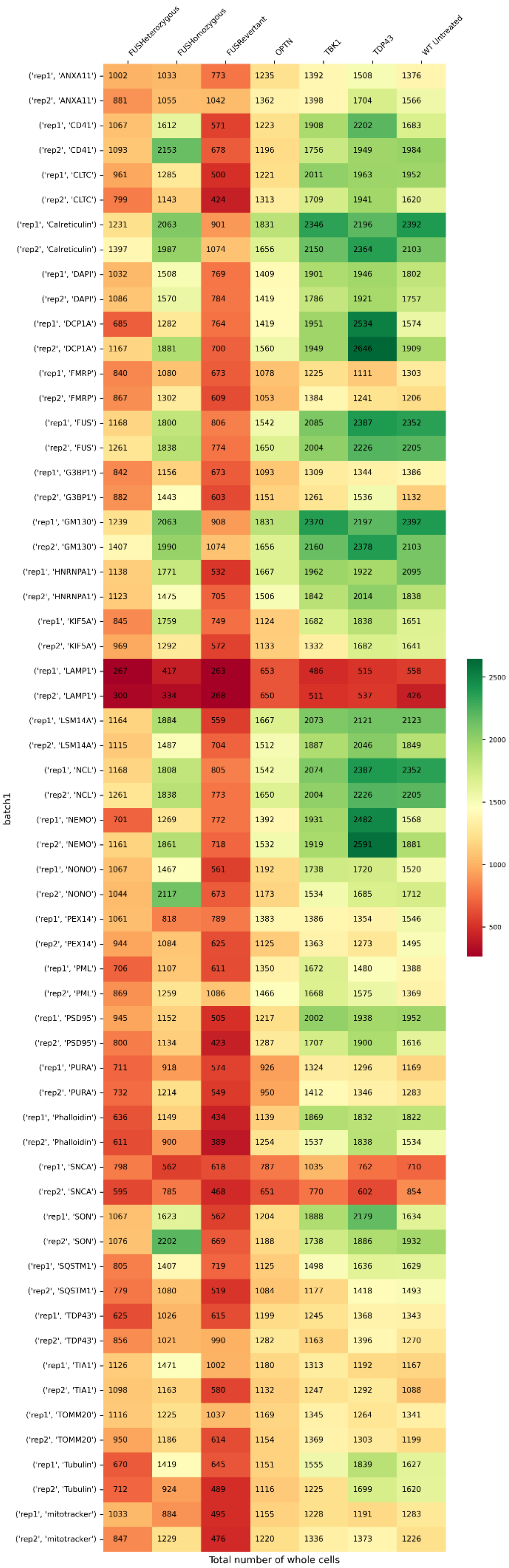

#### Batch 2

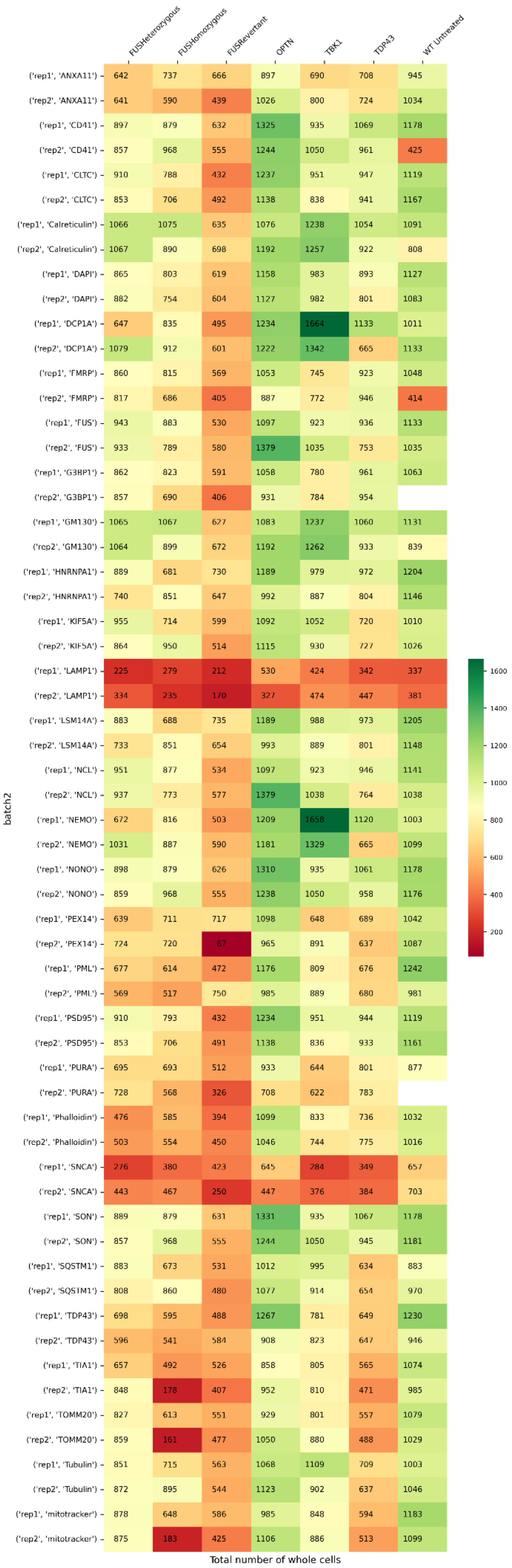
FUSHeterozygous
FUSHomozygous
FUSRevertant
OPTN
TBK1
TDP43
WT Untreated
batch2
('rep1', 'ANXA11') 642 737 666 897 690 708 945
('rep2', 'ANXA11') 641 590 439 1026 800 724 1034
('rep1', 'CD41') 897 879 632 1325 935 1069 1178
('rep2', 'CD41') 857 968 555 1244 1050 961 425
('rep1', 'CLTC') 910 788 432 1237 951 947 1119
('rep2', 'CLTC') 853 706 492 1138 838 941 1167
('rep1', 'Calreticulin') 1066 1075 635 1076 1238 1054 1091
('rep2', 'Calreticulin') 1067 890 698 1192 1257 922 808
('rep1', 'DAPI') 865 803 619 1158 983 893 1127
('rep2', 'DAPI') 882 754 604 1127 982 801 1083
('rep1', 'DCP1A') 647 835 495 1234 1664 1133 1011
('rep2', 'DCP1A') 1079 912 601 1222 1342 665 1133
('rep1', 'FMRP') 860 815 569 1053 745 923 1048
('rep2', 'FMRP') 817 686 405 887 772 946 414
('rep1', 'FUS') 943 883 530 1097 923 936 1133
('rep2', 'FUS') 933 789 580 1379 1035 753 1035
('rep1', 'G3BP1') 862 823 591 1058 780 961 1063
('rep2', 'G3BP1') 857 690 406 931 784 954
('rep1', 'GM130') 1065 1067 627 1083 1237 1060 1131
('rep2', 'GM130') 1064 899 672 1192 1262 933 839
('rep1', 'HNRNPA1') 889 681 730 1189 979 972 1204
('rep2', 'HNRNPA1') 740 851 647 992 887 804 1146
('rep1', 'KIF5A') 955 714 599 1092 1052 720 1010
('rep2', 'KIF5A') 864 950 514 1115 930 727 1026
('rep1', 'LAMP1') 225 279 212 530 424 342 337
('rep2', 'LAMP1') 334 235 170 327 474 447 381
('rep1', 'LSM14A') 883 688 735 1189 988 973 1205
('rep2', 'LSM14A') 733 851 654 993 889 801 1148
('rep1', 'NCL') 951 877 534 1097 923 946 1141
('rep2', 'NCL') 937 773 577 1379 1038 764 1038
('rep1', 'NEMO') 672 816 503 1209 1658 1120 1003
('rep2', 'NEMO') 1031 887 590 1181 1329 665 1099
('rep1', 'NONO') 898 879 626 1310 935 1061 1178
('rep2', 'NONO') 859 968 555 1238 1050 958 1176
('rep1', 'PEX14') 639 711 717 1098 648 689 1042
('rep2', 'PEX14') 724 720 67 965 891 637 1087
('rep1', 'PML') 677 614 472 1176 809 676 1242
('rep2', 'PML') 569 517 750 985 889 680 981
('rep1', 'PSD95') 910 793 432 1234 951 944 1119
('rep2', 'PSD95') 853 706 491 1138 836 933 1161
('rep1', 'PURA') 695 693 512 933 644 801 877
('rep2', 'PURA') 728 568 326 708 622 783
('rep1', 'Phalloidin') 476 585 394 1099 833 736 1032
('rep2', 'Phalloidin') 503 554 450 1046 744 775 1016
('rep1', 'SNCA') 276 380 423 645 284 349 657
('rep2', 'SNCA') 443 467 250 447 376 384 703
('rep1', 'SON') 889 879 631 1331 935 1067 1178
('rep2', 'SON') 857 968 555 1244 1050 945 1181
('rep1', 'SQSTM1') 883 673 531 1012 995 634 883
('rep2', 'SQSTM1') 808 860 480 1077 914 654 970
('rep1', 'TDP43') 698 595 488 1267 781 649 1230
('rep2', 'TDP43') 596 541 584 908 823 647 946
('rep1', 'TIA1') 657 492 526 858 805 565 1074
('rep2', 'TIA1') 848 178 407 952 810 471 985
('rep1', 'TOMM20') 827 613 551 929 801 557 1079
('rep2', 'TOMM20') 859 161 477 1050 880 488 1029
('rep1', 'Tubulin') 851 715 563 1068 1109 709 1003
('rep2', 'Tubulin') 872 895 544 1123 902 637 1046
('rep1', 'mitotracker') 878 648 586 985 848 594 1183
('rep2', 'mitotracker') 875 183 425 1106 886 513 1099
Total number of whole cells
1600
1400
1200
1000
800
600
400
200

### Batch 3

| batch3 | FUSHeterozygous | FUSHomozygous | FUSRevertant | OPTN | TBK1 | TDP43 | WT Untreated |
|---|---|---|---|---|---|---|---|
| ('rep1', 'ANXA11') | 774 | 729 | 816 | 1071 | 750 | 1208 | 990 |
| ('rep2', 'ANXA11') | 466 | 882 | 881 | 1022 | 801 | 1160 | 1085 |
| ('rep1', 'CD41') | 465 | 723 | 525 | 723 | 857 | 958 | 897 |
| ('rep2', 'CD41') | 538 | 685 | 768 | 685 | 806 | 1353 | 898 |
| ('rep1', 'CLTC') | 491 | 807 | 742 | 801 | 813 | 975 | 603 |
| ('rep2', 'CLTC') | 661 | 777 | 586 | 759 | 661 | 1103 | 742 |
| ('rep1', 'Calreticulin') | 850 | 1044 | 892 | 985 | 1089 | 1131 | 1322 |
| ('rep2', 'Calreticulin') | 925 | 1164 | 977 | 1365 | 1037 | 1240 | 1128 |
| ('rep1', 'DAPI') | 666 | 915 | 889 | 1020 | 850 | 1222 | 1000 |
| ('rep2', 'DAPI') | 692 | 874 | 861 | 1007 | 841 | 1317 | 990 |
| ('rep1', 'DCP1A') | 549 | 996 | 892 | 1109 | 732 | 1189 | 1030 |
| ('rep2', 'DCP1A') | 780 | 1024 | 973 | 1385 | 704 | 1645 | 1027 |
| ('rep1', 'FMRP') | 509 | 899 | 398 | 634 | 597 | 1003 | 742 |
| ('rep2', 'FMRP') | 468 | 666 | 426 | 533 | 483 | 797 | 706 |
| ('rep1', 'FUS') | 715 | 1043 | 1033 | 1226 | 962 | 1570 | 982 |
| ('rep2', 'FUS') | 851 | 1081 | 1003 | 1311 | 1016 | 1682 | 1250 |
| ('rep1', 'G3BP1') | 534 | 911 | 414 | 690 | 621 | 1077 | 781 |
| ('rep2', 'G3BP1') | 491 | 681 | 448 | 561 | 516 | 938 | 736 |
| ('rep1', 'GM130') | 843 | 1040 | 895 | 1008 | 1082 | 1152 | 1341 |
| ('rep2', 'GM130') | 915 | 1177 | 973 | 1390 | 1050 | 1282 | 1139 |
| ('rep1', 'HNRNPA1') | 603 | 1132 | 768 | 1107 | 923 | 1326 | 1245 |
| ('rep2', 'HNRNPA1') | 568 | 1014 | 554 | 925 | 876 | 1139 | 821 |
| ('rep1', 'KIF5A') | 633 | 795 | 772 | 899 | 999 | 830 | 682 |
| ('rep2', 'KIF5A') | 491 | 610 | 677 | 769 | 846 | 978 | 725 |
| ('rep1', 'LAMP1') | 344 | 189 | 299 | 261 | 163 | 315 | 275 |
| ('rep2', 'LAMP1') | 215 | 165 | 332 | 317 | 337 | 192 | 240 |
| ('rep1', 'LSM14A') | 604 | 1134 | 776 | 1107 | 927 | 1326 | 1246 |
| ('rep2', 'LSM14A') | 561 | 1004 | 573 | 925 | 871 | 1139 | 826 |
| ('rep1', 'NCL') | 699 | 1041 | 1036 | 1233 | 967 | 1570 | 991 |
| ('rep2', 'NCL') | 852 | 1078 | 1006 | 1311 | 1009 | 1685 | 1252 |
| ('rep1', 'NEMO') | 561 | 1010 | 861 | 1087 | 745 | 1141 | 992 |
| ('rep2', 'NEMO') | 783 | 1021 | 952 | 1382 | 754 | 1573 | 1034 |
| ('rep1', 'NONO') | 480 | 747 | 534 | 771 | 863 | 993 | 902 |
| ('rep2', 'NONO') | 561 | 724 | 770 | 690 | 818 | 1339 | 910 |
| ('rep1', 'PEX14') | 571 | 767 | 875 | 1032 | 651 | 1270 | 979 |
| ('rep2', 'PEX14') | 792 | 711 | 880 | 946 | 817 | 1330 | 1154 |
| ('rep1', 'PML') | 475 | 808 | 1092 | 965 | 748 | 1321 | 766 |
| ('rep2', 'PML') | 491 | 578 | 679 | 912 | 577 | 1140 | 780 |
| ('rep1', 'PSD95') | 487 | 802 | 742 | 863 | 808 | 989 | 637 |
| ('rep2', 'PSD95') | 655 | 777 | 586 | 759 | 655 | 1098 | 740 |
| ('rep1', 'PURA') | 439 | 661 | 366 | 572 | 456 | 870 | 659 |
| ('rep2', 'PURA') | 402 | 563 | 394 | 508 | 470 | 722 | 645 |
| ('rep1', 'Phalloidin') | 404 | 696 | 631 | 663 | 757 | 955 | 695 |
| ('rep2', 'Phalloidin') | 515 | 636 | 553 | 733 | 637 | 1045 | 704 |
| ('rep1', 'SNCA') | 261 | 447 | 580 | 520 | 294 | 547 | 613 |
| ('rep2', 'SNCA') | 426 | 371 | 703 | 343 | 343 | 676 | 705 |
| ('rep1', 'SON') | 486 | 737 | 528 | 758 | 849 | 995 | 902 |
| ('rep2', 'SON') | 561 | 723 | 779 | 692 | 824 | 1399 | 906 |
| ('rep1', 'SQSTM1') | 600 | 773 | 774 | 881 | 867 | 845 | 662 |
| ('rep2', 'SQSTM1') | 453 | 634 | 652 | 775 | 786 | 1026 | 808 |
| ('rep1', 'TDP43') | 525 | 844 | 848 | 931 | 716 | 1208 | 912 |
| ('rep2', 'TDP43') | 529 | 592 | 616 | 955 | 670 | 1107 | 918 |
| ('rep1', 'TIA1') | 891 | 890 | 1075 | 1024 | 741 | 1031 | 1016 |
| ('rep2', 'TIA1') | 767 | 793 | 973 | 957 | 864 | 1193 | 892 |
| ('rep1', 'TOMM20') | 901 | 922 | 1019 | 1062 | 767 | 1192 | 1082 |
| ('rep2', 'TOMM20') | 832 | 834 | 980 | 970 | 900 | 1281 | 965 |
| ('rep1', 'Tubulin') | 626 | 717 | 750 | 992 | 906 | 891 | 625 |
| ('rep2', 'Tubulin') | 441 | 552 | 690 | 743 | 832 | 1053 | 821 |
| ('rep1', 'mitotracker') | 874 | 900 | 956 | 1103 | 829 | 1222 | 1149 |
| ('rep2', 'mitotracker') | 834 | 850 | 996 | 1071 | 959 | 1347 | 990 |

Total number of whole cells

Color scale: 200, 400, 600, 800, 1000, 1200, 1400, 1600

### Batch 4

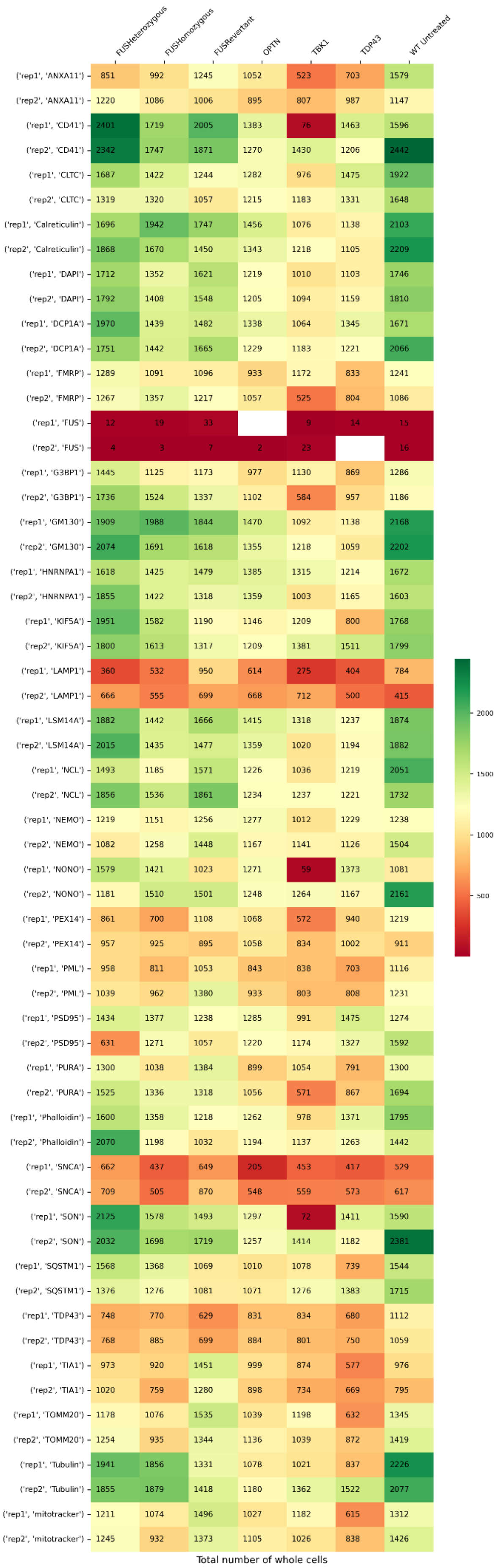

| | FUSHeterozygous | FUSHomozygous | FUSRevertant | OPTN | TBK1 | TDP43 | WT Untreated |
|---|---|---|---|---|---|---|---|
| ('rep1', 'ANXA11') | 851 | 992 | 1245 | 1052 | 523 | 703 | 1579 |
| ('rep2', 'ANXA11') | 1220 | 1086 | 1006 | 895 | 807 | 987 | 1147 |
| ('rep1', 'CD41') | 2401 | 1719 | 2005 | 1383 | 76 | 1463 | 1596 |
| ('rep2', 'CD41') | 2342 | 1747 | 1871 | 1270 | 1430 | 1206 | 2442 |
| ('rep1', 'CLTC') | 1687 | 1422 | 1244 | 1282 | 976 | 1475 | 1922 |
| ('rep2', 'CLTC') | 1319 | 1320 | 1057 | 1215 | 1183 | 1331 | 1648 |
| ('rep1', 'Calreticulin') | 1696 | 1942 | 1747 | 1456 | 1076 | 1138 | 2103 |
| ('rep2', 'Calreticulin') | 1868 | 1670 | 1450 | 1343 | 1218 | 1105 | 2209 |
| ('rep1', 'DAPI') | 1712 | 1352 | 1621 | 1219 | 1010 | 1103 | 1746 |
| ('rep2', 'DAPI') | 1792 | 1408 | 1548 | 1205 | 1094 | 1159 | 1810 |
| ('rep1', 'DCP1A') | 1970 | 1439 | 1482 | 1338 | 1064 | 1345 | 1671 |
| ('rep2', 'DCP1A') | 1751 | 1442 | 1665 | 1229 | 1183 | 1221 | 2066 |
| ('rep1', 'FMRP') | 1289 | 1091 | 1096 | 933 | 1172 | 833 | 1241 |
| ('rep2', 'FMRP') | 1267 | 1357 | 1217 | 1057 | 525 | 804 | 1086 |
| ('rep1', 'FUS') | 12 | 19 | 33 | | 9 | 14 | 15 |
| ('rep2', 'FUS') | 4 | 3 | 7 | 2 | 23 | | 16 |
| ('rep1', 'G3BP1') | 1445 | 1125 | 1173 | 977 | 1130 | 869 | 1286 |
| ('rep2', 'G3BP1') | 1736 | 1524 | 1337 | 1102 | 584 | 957 | 1186 |
| ('rep1', 'GM130') | 1909 | 1988 | 1844 | 1470 | 1092 | 1138 | 2168 |
| ('rep2', 'GM130') | 2074 | 1691 | 1618 | 1355 | 1218 | 1059 | 2202 |
| ('rep1', 'HNRNPA1') | 1618 | 1425 | 1479 | 1385 | 1315 | 1214 | 1672 |
| ('rep2', 'HNRNPA1') | 1855 | 1422 | 1318 | 1359 | 1003 | 1165 | 1603 |
| ('rep1', 'KIF5A') | 1951 | 1582 | 1190 | 1146 | 1209 | 800 | 1768 |
| ('rep2', 'KIF5A') | 1800 | 1613 | 1317 | 1209 | 1381 | 1511 | 1799 |
| ('rep1', 'LAMP1') | 360 | 532 | 950 | 614 | 275 | 404 | 784 |
| ('rep2', 'LAMP1') | 666 | 555 | 699 | 668 | 712 | 500 | 415 |
| ('rep1', 'LSM14A') | 1882 | 1442 | 1666 | 1415 | 1318 | 1237 | 1874 |
| ('rep2', 'LSM14A') | 2015 | 1435 | 1477 | 1359 | 1020 | 1194 | 1882 |
| ('rep1', 'NCL') | 1493 | 1185 | 1571 | 1226 | 1036 | 1219 | 2051 |
| ('rep2', 'NCL') | 1856 | 1536 | 1861 | 1234 | 1237 | 1221 | 1732 |
| ('rep1', 'NEMO') | 1219 | 1151 | 1256 | 1277 | 1012 | 1229 | 1238 |
| ('rep2', 'NEMO') | 1082 | 1258 | 1448 | 1167 | 1141 | 1126 | 1504 |
| ('rep1', 'NONO') | 1579 | 1421 | 1023 | 1271 | 59 | 1373 | 1081 |
| ('rep2', 'NONO') | 1181 | 1510 | 1501 | 1248 | 1264 | 1167 | 2161 |
| ('rep1', 'PEX14') | 861 | 700 | 1108 | 1068 | 572 | 940 | 1219 |
| ('rep2', 'PEX14') | 957 | 925 | 895 | 1058 | 834 | 1002 | 911 |
| ('rep1', 'PML') | 958 | 811 | 1053 | 843 | 838 | 703 | 1116 |
| ('rep2', 'PML') | 1039 | 962 | 1380 | 933 | 803 | 808 | 1231 |
| ('rep1', 'PSD95') | 1434 | 1377 | 1238 | 1285 | 991 | 1475 | 1274 |
| ('rep2', 'PSD95') | 631 | 1271 | 1057 | 1220 | 1174 | 1327 | 1592 |
| ('rep1', 'PURA') | 1300 | 1038 | 1384 | 899 | 1054 | 791 | 1300 |
| ('rep2', 'PURA') | 1525 | 1336 | 1318 | 1056 | 571 | 867 | 1694 |
| ('rep1', 'Phalloidin') | 1600 | 1358 | 1218 | 1262 | 978 | 1371 | 1795 |
| ('rep2', 'Phalloidin') | 2070 | 1198 | 1032 | 1194 | 1137 | 1263 | 1442 |
| ('rep1', 'SNCA') | 662 | 437 | 649 | 205 | 453 | 417 | 579 |
| ('rep2', 'SNCA') | 709 | 505 | 870 | 548 | 559 | 573 | 617 |
| ('rep1', 'SON') | 2125 | 1578 | 1493 | 1297 | 72 | 1411 | 1590 |
| ('rep2', 'SON') | 2032 | 1698 | 1719 | 1257 | 1414 | 1182 | 2381 |
| ('rep1', 'SQSTM1') | 1568 | 1368 | 1069 | 1010 | 1078 | 739 | 1544 |
| ('rep2', 'SQSTM1') | 1376 | 1276 | 1081 | 1071 | 1276 | 1383 | 1715 |
| ('rep1', 'TDP43') | 748 | 770 | 629 | 831 | 834 | 680 | 1112 |
| ('rep2', 'TDP43') | 768 | 885 | 699 | 884 | 801 | 750 | 1059 |
| ('rep1', 'TIA1') | 973 | 920 | 1451 | 999 | 874 | 577 | 976 |
| ('rep2', 'TIA1') | 1020 | 759 | 1280 | 898 | 734 | 669 | 795 |
| ('rep1', 'TOMM20') | 1178 | 1076 | 1535 | 1039 | 1198 | 632 | 1345 |
| ('rep2', 'TOMM20') | 1254 | 935 | 1344 | 1136 | 1039 | 872 | 1419 |
| ('rep1', 'Tubulin') | 1941 | 1856 | 1331 | 1078 | 1021 | 837 | 2226 |
| ('rep2', 'Tubulin') | 1855 | 1879 | 1418 | 1180 | 1362 | 1522 | 2077 |
| ('rep1', 'mitotracker') | 1211 | 1074 | 1496 | 1027 | 1182 | 615 | 1312 |
| ('rep2', 'mitotracker') | 1245 | 932 | 1373 | 1105 | 1026 | 838 | 1426 |

2000
1500
1000
500
Total number of whole cells

### Batch 5

| | FUSHeterozygous | FUSHomozygous | FUSRevertant | OPTN | TBK1 | TDP43 | WT Untreated |
|---|---|---|---|---|---|---|---|
| ('rep1', 'ANXA11') | 1004 | 1139 | 1324 | 1104 | 809 | 998 | 1181 |
| ('rep2', 'ANXA11') | 1079 | 1292 | 1447 | 1158 | 710 | 1114 | 1102 |
| ('rep1', 'CD41') | 1884 | 1521 | 1395 | 1518 | 1169 | 1688 | 1619 |
| ('rep2', 'CD41') | 1463 | 1654 | 1704 | 1189 | 1129 | 1545 | 1683 |
| ('rep1', 'CLTC') | 1705 | 1450 | 1558 | 1319 | 987 | 1441 | 1532 |
| ('rep2', 'CLTC') | 1548 | 1381 | 1260 | 1257 | 1020 | 1243 | 1183 |
| ('rep1', 'Calreticulin') | 1963 | 1919 | 1495 | 1783 | 1254 | 1471 | 1847 |
| ('rep2', 'Calreticulin') | 1938 | 1808 | 1920 | 1485 | 1298 | 1891 | 1768 |
| ('rep1', 'DAPI') | 1570 | 1470 | 1560 | 1367 | 997 | 1389 | 1485 |
| ('rep2', 'DAPI') | 1545 | 1502 | 1582 | 1258 | 1019 | 1375 | 1505 |
| ('rep1', 'DCP1A') | 1750 | 1666 | 1530 | 1458 | 1121 | 1680 | 1775 |
| ('rep2', 'DCP1A') | 1867 | 1707 | 1680 | 1478 | 1330 | 1348 | 1749 |
| ('rep1', 'FMRP') | 1085 | 1252 | 810 | 1208 | 733 | 1199 | 1388 |
| ('rep2', 'FMRP') | 1004 | 1243 | 803 | 1046 | 703 | 1252 | 1348 |
| ('rep1', 'FUS') | 1677 | 1707 | 1573 | 1579 | 1200 | 1752 | 1884 |
| ('rep2', 'FUS') | 1973 | 1830 | 1916 | 1367 | 1229 | 1469 | 1779 |
| ('rep1', 'G3BP1') | 1337 | 1330 | 900 | 1242 | 775 | 1351 | 1534 |
| ('rep2', 'G3BP1') | 1179 | 1387 | 912 | 1106 | 765 | 1393 | 1473 |
| ('rep1', 'GM130') | 1963 | 1919 | 1495 | 1778 | 1254 | 1483 | 1847 |
| ('rep2', 'GM130') | 1939 | 1808 | 1920 | 1478 | 1306 | 1899 | 1768 |
| ('rep1', 'HNRNPA1') | 1299 | 1712 | 1522 | 1434 | 1106 | 1638 | 1573 |
| ('rep2', 'HNRNPA1') | 1240 | 1631 | 1231 | 1359 | 1120 | 1560 | 1591 |
| ('rep1', 'KIF5A') | 1415 | 1185 | 1256 | 1118 | 853 | 973 | 1048 |
| ('rep2', 'KIF5A') | 1488 | 1299 | 1502 | 956 | 855 | 1311 | 1413 |
| ('rep1', 'LAMP1') | 573 | 773 | 947 | 564 | 582 | 462 | 694 |
| ('rep2', 'LAMP1') | 643 | 658 | 819 | 786 | 650 | 732 | 819 |
| ('rep1', 'LSM14A') | 1370 | 1821 | 1801 | 1444 | 1106 | 1646 | 1592 |
| ('rep2', 'LSM14A') | 1521 | 1693 | 1659 | 1374 | 1127 | 1585 | 1607 |
| ('rep1', 'NCL') | 1677 | 1697 | 1558 | 1579 | 1200 | 1742 | 1884 |
| ('rep2', 'NCL') | 1942 | 1835 | 1804 | 1367 | 1229 | 1469 | 1779 |
| ('rep1', 'NEMO') | 1745 | 1654 | 1526 | 1446 | 1103 | 1680 | 1775 |
| ('rep2', 'NEMO') | 1867 | 1700 | 1691 | 1436 | 1314 | 1340 | 1749 |
| ('rep1', 'NONO') | 1397 | 1469 | 1227 | 1344 | 1164 | 1526 | 1411 |
| ('rep2', 'NONO') | 1290 | 1620 | 1457 | 1171 | 1115 | 1415 | 1469 |
| ('rep1', 'PEX14') | 1071 | 1193 | 1008 | 1104 | 759 | 1176 | 1105 |
| ('rep2', 'PEX14') | 1108 | 1055 | 1220 | 1013 | 831 | 1123 | 1069 |
| ('rep1', 'PML') | 839 | 1063 | 1514 | 1089 | 621 | 801 | 748 |
| ('rep2', 'PML') | 1171 | 1120 | 1176 | 1182 | 638 | 962 | 1022 |
| ('rep1', 'PSD95') | 1749 | 1478 | 1565 | 1312 | 986 | 1434 | 1412 |
| ('rep2', 'PSD95') | 1537 | 1381 | 1246 | 1255 | 1020 | 1238 | 130 |
| ('rep1', 'PURA') | 1132 | 1083 | 804 | 1132 | 715 | 1229 | 1214 |
| ('rep2', 'PURA') | 1024 | 1113 | 849 | 920 | 624 | 1143 | 1261 |
| ('rep1', 'Phalloidin') | 1568 | 1438 | 1480 | 1299 | 927 | 1307 | 1539 |
| ('rep2', 'Phalloidin') | 1338 | 1268 | 1181 | 1229 | 929 | 1192 | 1240 |
| ('rep1', 'SNCA') | 704 | 484 | 1375 | 561 | 325 | 544 | 385 |
| ('rep2', 'SNCA') | 769 | 555 | 678 | 471 | 593 | 824 | 432 |
| ('rep1', 'SON') | 1769 | 1531 | 1377 | 1447 | 1169 | 1726 | 1612 |
| ('rep2', 'SON') | 1414 | 1654 | 1638 | 1168 | 1120 | 1520 | 1626 |
| ('rep1', 'SQSTM1') | 1165 | 949 | 1001 | 990 | 831 | 795 | 888 |
| ('rep2', 'SQSTM1') | 1209 | 1254 | 1261 | 898 | 852 | 1166 | 1291 |
| ('rep1', 'TDP43') | 800 | 960 | 556 | 1025 | 671 | 800 | 878 |
| ('rep2', 'TDP43') | 1081 | 1097 | 885 | 1119 | 708 | 1055 | 1084 |
| ('rep1', 'TIA1') | 1228 | 951 | 1030 | 1006 | 622 | 740 | 864 |
| ('rep2', 'TIA1') | 926 | 1040 | 1453 | 804 | 532 | 763 | 690 |
| ('rep1', 'TOMM20') | 1087 | 959 | 932 | 1078 | 773 | 897 | 967 |
| ('rep2', 'TOMM20') | 896 | 1143 | 1299 | 921 | 610 | 953 | 1008 |
| ('rep1', 'Tubulin') | 1701 | 1545 | 1607 | 1281 | 1094 | 1229 | 1407 |
| ('rep2', 'Tubulin') | 1600 | 1593 | 1656 | 1178 | 998 | 1336 | 1687 |
| ('rep1', 'mitotracker') | 1544 | 1016 | 1031 | 1087 | 773 | 913 | 909 |
| ('rep2', 'mitotracker') | 1057 | 1232 | 1475 | 904 | 588 | 919 | 1013 |

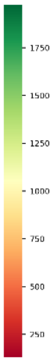


Total number of whole cells

### Batch 6

| | FUSHeterozygous | FUSHomozygous | FUSRevertant | OPTN | TBK1 | TDP43 | WT Untreated |
|---|---|---|---|---|---|---|---|
| ('rep1', 'ANXA11') | 385 | 217 | 545 | 623 | 195 | 887 | 861 |
| ('rep2', 'ANXA11') | 296 | 254 | 380 | 259 | 469 | 510 | 869 |
| ('rep1', 'CD41') | 1991 | 1319 | 1801 | 1825 | 1165 | 1456 | 8 |
| ('rep2', 'CD41') | 1936 | 1398 | 1685 | 1962 | 1179 | 1495 | 2064 |
| ('rep1', 'CLTC') | 1714 | 1316 | 1296 | 1668 | 1125 | 1447 | 1814 |
| ('rep2', 'CLTC') | 1520 | 1364 | 1087 | 1607 | 1220 | 1450 | 1779 |
| ('rep1', 'Calreticulin') | 1689 | 1277 | 1885 | | 1180 | 1546 | 1917 |
| ('rep2', 'Calreticulin') | 1973 | 1389 | 1689 | 1724 | 1187 | 1339 | 1833 |
| ('rep1', 'DAPI') | 1453 | 1064 | 1317 | 1308 | 1007 | 1259 | 1543 |
| ('rep2', 'DAPI') | 1414 | 1119 | 1338 | 1452 | 1092 | 1230 | 1669 |
| ('rep1', 'DCP1A') | 1604 | 1187 | 1362 | 1585 | 1207 | 1253 | 2128 |
| ('rep2', 'DCP1A') | 1683 | 1261 | 1410 | 1880 | 1155 | 1353 | 2054 |
| ('rep1', 'FMRP') | 1320 | 1097 | 994 | 1401 | 986 | 1072 | 1527 |
| ('rep2', 'FMRP') | 1552 | 1318 | 1112 | 1515 | 1134 | 1454 | 1563 |
| ('rep1', 'FUS') | 1870 | 1335 | 1687 | 1570 | 1322 | 1317 | 1875 |
| ('rep2', 'FUS') | 1849 | 1332 | 1846 | 1656 | 1343 | 1490 | 1652 |
| ('rep1', 'G3BP1') | 1369 | 1114 | 1021 | 1356 | 1004 | 1094 | 1619 |
| ('rep2', 'G3BP1') | 1584 | 1338 | 1182 | 1525 | 1174 | 1473 | 1961 |
| ('rep1', 'GM130') | 1710 | 1281 | 1907 | | 1180 | 1546 | 1917 |
| ('rep2', 'GM130') | 1983 | 1389 | 1697 | 1726 | 1187 | 1343 | 1839 |
| ('rep1', 'HNRNPA1') | 1721 | 1392 | 1266 | 1623 | 1052 | 1301 | 1557 |
| ('rep2', 'HNRNPA1') | 1434 | 1305 | 1064 | 1430 | 1188 | 1275 | 1519 |
| ('rep1', 'KIF5A') | 1513 | 1012 | 1537 | 1407 | 769 | 1239 | 1621 |
| ('rep2', 'KIF5A') | 1772 | 1079 | 1516 | 1341 | 994 | 1081 | 1556 |
| ('rep1', 'LAMP1') | 337 | 171 | 656 | 861 | 490 | 1060 | 772 |
| ('rep2', 'LAMP1') | 256 | 225 | 612 | 475 | 549 | 548 | 731 |
| ('rep1', 'LSM14A') | 1894 | 1433 | 1266 | 1655 | 1058 | 1319 | 1619 |
| ('rep2', 'LSM14A') | 1600 | 1326 | 1308 | 1430 | 1193 | 1283 | 1590 |
| ('rep1', 'NCL') | 1870 | 1349 | 1665 | 1570 | 1322 | 1317 | 1875 |
| ('rep2', 'NCL') | 1790 | 1339 | 1767 | 1656 | 1343 | 1479 | 1640 |
| ('rep1', 'NEMO') | 1598 | 1187 | 1360 | 1571 | 1204 | 1260 | 2113 |
| ('rep2', 'NEMO') | 1679 | 1253 | 1400 | 1880 | 1147 | 1353 | 2038 |
| ('rep1', 'NONO') | 1913 | 1323 | 1520 | 1836 | 1160 | 1467 | |
| ('rep2', 'NONO') | 1831 | 1398 | 1487 | 1943 | 1175 | 1502 | 2064 |
| ('rep1', 'PEX14') | 384 | 408 | 586 | 601 | 563 | 690 | 975 |
| ('rep2', 'PEX14') | 500 | 376 | 816 | 871 | 641 | 633 | 1311 |
| ('rep1', 'PML') | 273 | 308 | 166 | 872 | 485 | 1020 | 976 |
| ('rep2', 'PML') | 793 | 701 | 617 | 644 | 643 | 714 | 1086 |
| ('rep1', 'PSD95') | 1712 | 1304 | 1286 | 1668 | 1125 | 1429 | 1811 |
| ('rep2', 'PSD95') | 1484 | 1354 | 1078 | 1543 | 1218 | 1455 | 1775 |
| ('rep1', 'PURA') | 1208 | 897 | 881 | 1193 | 856 | 976 | 1295 |
| ('rep2', 'PURA') | 1185 | 1034 | 1042 | 1429 | 1008 | 1096 | 1769 |
| ('rep1', 'Phalloidin') | 1481 | 1097 | 1234 | 1636 | 1048 | 1343 | 1559 |
| ('rep2', 'Phalloidin') | 1330 | 1221 | 1053 | 1554 | 1042 | 1385 | 1402 |
| ('rep1', 'SNCA') | 413 | 238 | 510 | 528 | 392 | 439 | 586 |
| ('rep2', 'SNCA') | 445 | 181 | 701 | 625 | 411 | 312 | 849 |
| ('rep1', 'SON') | 1972 | 1319 | 1706 | 1827 | 1165 | 1498 | 8 |
| ('rep2', 'SON') | 1886 | 1398 | 1635 | 1944 | 1169 | 1486 | 2053 |
| ('rep1', 'SQSTM1') | 1282 | 834 | 1310 | 959 | 587 | 1022 | 1251 |
| ('rep2', 'SQSTM1') | 1416 | 918 | 1164 | 1066 | 825 | 879 | 1328 |
| ('rep1', 'TDP43') | 260 | 298 | 150 | 737 | 458 | 944 | 860 |
| ('rep2', 'TDP43') | 537 | 683 | 579 | 688 | 622 | 679 | 910 |
| ('rep1', 'TIA1') | 1072 | 496 | 992 | 1067 | 423 | 739 | 939 |
| ('rep2', 'TIA1') | 723 | 728 | 819 | 1201 | 771 | 842 | 730 |
| ('rep1', 'TOMM20') | 1566 | 1023 | 1158 | 1117 | 821 | 980 | 1474 |
| ('rep2', 'TOMM20') | 713 | 1126 | 855 | 1296 | 1060 | 1202 | 1348 |
| ('rep1', 'Tubulin') | 1809 | 1286 | 1671 | 1682 | 967 | 1372 | 2012 |
| ('rep2', 'Tubulin') | 1905 | 1144 | 1620 | 1619 | 1067 | 1036 | 1766 |
| ('rep1', 'mitotracker') | 1765 | 1008 | 1150 | 1144 | 811 | 968 | 1490 |
| ('rep2', 'mitotracker') | 852 | 1112 | 842 | 1226 | 1028 | 1165 | 1342 |

Total number of whole cells
