## Supplementary material for "Organellomics: AI-driven deep organellar phenotyping reveals novel ALS mechanisms in human neurons": Day 8 iPSC-derived cortical neurons (used for model training)- QC Report.pdf

#### Processed Files Validation

Number of sites that survived pre-processing

Batch 1

| | Rep | FUSHomozygous | TDP43 | TBK1 | WT stress | WT Untreated | FUSRevertant | OPTN | FUSHeterozygous | SNCA |
|---|---|---|---|---|---|---|---|---|---|---|
| G3BP1 | rep1 | 80 | 24 | 96 | 56 | 33 | 90 | 78 | 44 | 8 |
| G3BP1 | rep2 | 93 | 100 | 100 | 100 | 100 | 100 | 100 | 62 | 22 |
| NONO | rep1 | 92 | 100 | 95 | 100 | 100 | 23 | 100 | 40 | 12 |
| NONO | rep2 | 96 | 100 | 100 | 100 | 99 | 96 | 100 | 57 | 7 |
| SQSTM1 | rep1 | 77 | 93 | 83 | 57 | 97 | 73 | 87 | 31 | 7 |
| SQSTM1 | rep2 | 85 | 100 | 98 | 83 | 97 | 99 | 98 | 63 | 8 |
| PSD95 | rep1 | 96 | 96 | 100 | 100 | 100 | 100 | 99 | 54 | 10 |
| PSD95 | rep2 | 100 | 100 | 100 | 100 | 100 | 100 | 100 | 86 | 1 |
| NEMO | rep1 | 86 | 98 | 97 | 100 | 100 | 92 | 100 | 73 | 1 |
| NEMO | rep2 | 98 | 100 | 99 | 100 | 99 | 1 | 100 | 37 | 0 |
| GM130 | rep1 | 98 | 100 | 100 | 100 | 100 | 87 | 100 | 46 | 0 |
| GM130 | rep2 | 97 | 100 | 100 | 99 | 100 | 99 | 100 | 52 | 1 |
| NCL | rep1 | 67 | 84 | 100 | 96 | 100 | 90 | 100 | 32 | 3 |
| NCL | rep2 | 82 | 99 | 72 | 100 | 99 | 73 | 100 | 48 | 0 |
| ANXA11 | rep1 | 77 | 99 | 74 | 98 | 100 | 59 | 99 | 60 | 6 |
| ANXA11 | rep2 | 91 | 90 | 98 | 100 | 100 | 100 | 100 | 69 | 10 |
| Calreticulin | rep1 | 71 | 96 | 98 | 85 | 99 | 81 | 100 | 61 | 10 |
| Calreticulin | rep2 | 82 | 93 | 100 | 98 | 97 | 95 | 99 | 68 | 11 |
| mitotracker | rep1 | 100 | 92 | 100 | 100 | 97 | 35 | 99 | 24 | nan |
| mitotracker | rep2 | 100 | 90 | 99 | 100 | 99 | 54 | 100 | 29 | nan |
| KIF5A | rep1 | 77 | 19 | 89 | 43 | 27 | 87 | 67 | 39 | 6 |
| KIF5A | rep2 | 93 | 100 | 99 | 95 | 100 | 98 | 86 | 60 | 26 |
| TDP43 | rep1 | 92 | 100 | 99 | 100 | 100 | 81 | 100 | 47 | 12 |
| TDP43 | rep2 | 96 | 100 | 100 | 100 | 100 | 96 | 100 | 56 | 7 |
| FMRP | rep1 | 72 | 93 | 78 | 48 | 96 | 35 | 81 | 20 | 0 |
| FMRP | rep2 | 88 | 96 | 91 | 81 | 99 | 99 | 97 | 62 | 4 |
| CLTC | rep1 | 96 | 96 | 100 | 100 | 100 | 100 | 99 | 55 | 17 |
| CLTC | rep2 | 100 | 100 | 100 | 100 | 100 | 100 | 100 | 87 | 24 |
| DCP1A | rep1 | 26 | 6 | 56 | 92 | 79 | 9 | 42 | 0 | nan |
| DCP1A | rep2 | 85 | 84 | 81 | 98 | 95 | 99 | 83 | 7 | nan |
| TOMM20 | rep1 | 98 | 100 | 100 | 100 | 100 | 94 | 99 | 60 | 6 |
| TOMM20 | rep2 | 97 | 100 | 100 | 100 | 99 | 100 | 100 | 78 | 5 |
| FUS | rep1 | 67 | 83 | 100 | 88 | 100 | 90 | 100 | 37 | 6 |
| FUS | rep2 | 82 | 99 | 73 | 100 | 99 | 71 | 100 | 50 | 0 |
| SCNA | rep1 | 9 | 13 | 6 | 13 | 19 | 9 | 26 | 12 | 2 |
| SCNA | rep2 | 15 | 6 | 8 | 22 | 28 | 2 | 6 | 15 | 3 |
| LAMP1 | rep1 | 66 | 95 | 98 | 85 | 99 | 82 | 99 | 60 | 13 |
| LAMP1 | rep2 | 79 | 92 | 100 | 94 | 95 | 93 | 97 | 65 | 13 |
| TIA1 | rep1 | nan | nan | nan | nan | nan | nan | nan | nan | nan |
| TIA1 | rep2 | nan | nan | nan | nan | nan | nan | nan | nan | nan |
| PML | rep1 | 88 | 67 | 99 | 100 | 74 | 98 | 95 | 56 | 25 |
| PML | rep2 | 87 | 87 | 95 | 100 | 91 | 95 | 99 | 19 | 9 |
| PURA | rep1 | 78 | 21 | 88 | 54 | 29 | 86 | 71 | 32 | 7 |
| PURA | rep2 | 92 | 100 | 94 | 100 | 99 | 96 | 94 | 61 | 19 |
| CD41 | rep1 | 91 | 100 | 22 | 100 | 100 | 2 | 91 | 26 | 11 |
| CD41 | rep2 | 96 | 100 | 100 | 100 | 100 | 96 | 100 | 56 | 2 |
| Phalloidin | rep1 | 75 | 97 | 80 | 56 | 100 | 85 | 88 | 33 | 6 |
| Phalloidin | rep2 | 90 | 100 | 100 | 83 | 100 | 99 | 100 | 64 | 5 |
| PEX14 | rep1 | 100 | 99 | 100 | 100 | 100 | 99 | 100 | 73 | 27 |
| PEX14 | rep2 | 100 | 100 | 100 | 100 | 100 | 99 | 100 | 65 | 24 |
| DAPI | rep1 | 953 | 1004 | 1052 | 1021 | 1093 | 996 | 1085 | 650 | 236 |
| DAPI | rep2 | 1033 | 1083 | 1073 | 1068 | 1091 | 1063 | 1099 | 777 | 287 |

### Batch 2.1

| | Rep | FUSHomozygous | TDP43 | TBK1 | WT stress | WT Untreated | FUSRevertant | OPTN | FUSHeterozygous | SNCA |
|---|---|---|---|---|---|---|---|---|---|---|
| G3BP1 | rep1 | 100 | 99 | 100 | 100 | 100 | 100 | 100 | 99 | 100 |
| G3BP1 | rep2 | 100 | 100 | 100 | 100 | 100 | 100 | 100 | 100 | 100 |
| NONO | rep1 | 100 | 100 | 100 | 100 | 100 | 100 | 100 | 100 | 98 |
| NONO | rep2 | 100 | 100 | 100 | 100 | 100 | 100 | 100 | 100 | 100 |
| SQSTM1 | rep1 | 99 | 96 | 98 | 100 | 99 | 100 | 97 | 95 | 98 |
| SQSTM1 | rep2 | 98 | 94 | 99 | 100 | 99 | 98 | 97 | 97 | 99 |
| PSD95 | rep1 | 100 | 100 | 99 | 100 | 85 | 100 | 100 | 99 | 98 |
| PSD95 | rep2 | 100 | 100 | 100 | 100 | 99 | 100 | 100 | 100 | 77 |
| NEMO | rep1 | 100 | 99 | 100 | 100 | 100 | 100 | 100 | 100 | 96 |
| NEMO | rep2 | 87 | 99 | 69 | 100 | 84 | 68 | 100 | 98 | 46 |
| GM130 | rep1 | 100 | 100 | 100 | 100 | 100 | 99 | 98 | 99 | 0 |
| GM130 | rep2 | 100 | 100 | 100 | 100 | 97 | 99 | 98 | 100 | 85 |
| NCL | rep1 | 99 | 98 | 99 | 78 | 100 | 99 | 100 | 88 | 90 |
| NCL | rep2 | 100 | 100 | 100 | 98 | 100 | 100 | 100 | 93 | 98 |
| ANXA11 | rep1 | 100 | 100 | 100 | 100 | 99 | 100 | 96 | 98 | 100 |
| ANXA11 | rep2 | 97 | 100 | 99 | 100 | 100 | 100 | 100 | 90 | 100 |
| Calreticulin | rep1 | 100 | 94 | 100 | 100 | 100 | 100 | 98 | 100 | 99 |
| Calreticulin | rep2 | 97 | 45 | 99 | 94 | 95 | 99 | 95 | 97 | 98 |
| mitotracker | rep1 | 100 | 100 | 100 | 100 | 100 | 100 | 100 | 100 | 99 |
| mitotracker | rep2 | 100 | 96 | 100 | 100 | 100 | 100 | 100 | 100 | 100 |
| KIF5A | rep1 | 99 | 98 | 97 | 99 | 100 | 99 | 98 | 99 | 98 |
| KIF5A | rep2 | 98 | 100 | 98 | 99 | 100 | 98 | 98 | 100 | 98 |
| TDP43 | rep1 | 100 | 100 | 100 | 100 | 100 | 100 | 100 | 100 | 98 |
| TDP43 | rep2 | 100 | 100 | 100 | 100 | 100 | 100 | 100 | 100 | 100 |
| FMRP | rep1 | 100 | 99 | 100 | 99 | 100 | 100 | 100 | 100 | 100 |
| FMRP | rep2 | 100 | 100 | 100 | 100 | 99 | 100 | 100 | 99 | 98 |
| CLTC | rep1 | 100 | 100 | 100 | 100 | 99 | 100 | 100 | 99 | 99 |
| CLTC | rep2 | 100 | 100 | 100 | 100 | 100 | 100 | 100 | 100 | 93 |
| DCP1A | rep1 | 97 | 85 | 78 | 95 | 78 | 76 | 39 | 44 | 18 |
| DCP1A | rep2 | 86 | 55 | 64 | 88 | 24 | 29 | 11 | 11 | 5 |
| TOMM20 | rep1 | 100 | 100 | 100 | 100 | 100 | 99 | 100 | 99 | 0 |
| TOMM20 | rep2 | 100 | 100 | 100 | 100 | 100 | 100 | 99 | 98 | 100 |
| FUS | rep1 | 100 | 100 | 100 | 87 | 100 | 100 | 100 | 100 | 99 |
| FUS | rep2 | 100 | 100 | 100 | 100 | 100 | 100 | 100 | 100 | 99 |
| SCNA | rep1 | 98 | 88 | 97 | 89 | 89 | 92 | 85 | 93 | 92 |
| SCNA | rep2 | 70 | 96 | 49 | 91 | 90 | 95 | 92 | 80 | 95 |
| LAMP1 | rep1 | 95 | 88 | 100 | 96 | 99 | 98 | 91 | 96 | 95 |
| LAMP1 | rep2 | 74 | 33 | 86 | 72 | 77 | 83 | 93 | 86 | 91 |
| TIA1 | rep1 | nan | nan | nan | nan | nan | nan | nan | nan | nan |
| TIA1 | rep2 | nan | nan | nan | nan | nan | nan | nan | nan | nan |
| PML | rep1 | 97 | 100 | 55 | 100 | 99 | 96 | 99 | 100 | 97 |
| PML | rep2 | 100 | 100 | 93 | 100 | 100 | 96 | 99 | 100 | 97 |
| PURA | rep1 | 100 | 95 | 99 | 100 | 100 | 99 | 100 | 99 | 100 |
| PURA | rep2 | 99 | 100 | 100 | 100 | 98 | 100 | 99 | 100 | 100 |
| CD41 | rep1 | 100 | 100 | 100 | 100 | 98 | 100 | 90 | 99 | 98 |
| CD41 | rep2 | 100 | 100 | 100 | 100 | 100 | 100 | 95 | 100 | 100 |
| Phalloidin | rep1 | 100 | 100 | 100 | 100 | 100 | 100 | 100 | 100 | 100 |
| Phalloidin | rep2 | 100 | 100 | 100 | 100 | 100 | 100 | 100 | 99 | 100 |
| PEX14 | rep1 | 100 | 100 | 100 | 100 | 100 | 100 | 100 | 100 | 99 |
| PEX14 | rep2 | 100 | 100 | 100 | 100 | 100 | 100 | 100 | 100 | 100 |
| DAPI | rep1 | 1100 | 1093 | 1100 | 1089 | 1099 | 1099 | 1091 | 1096 | 990 |
| DAPI | rep2 | 1098 | 1099 | 1099 | 1100 | 1099 | 1098 | 1099 | 1088 | 1092 |

### Batch 2.2

| | Rep | FUSHomozygous | TDP43 | TBK1 | WT stress | WT Untreated | FUSRevertant | OPTN | FUSHeterozygous | SNCA |
|---|---|---|---|---|---|---|---|---|---|---|
| G3BP1 | rep1 | 100 | 93 | 99 | 100 | 99 | 100 | 99 | 100 | 94 |
| G3BP1 | rep2 | 100 | 100 | 100 | 100 | 100 | 100 | 100 | 100 | 100 |
| NONO | rep1 | 100 | 100 | 100 | 96 | 100 | 100 | 99 | 100 | 98 |
| NONO | rep2 | 100 | 100 | 100 | 99 | 100 | 100 | 100 | 100 | 100 |
| SQSTM1 | rep1 | 99 | 96 | 99 | 99 | 100 | 100 | 97 | 99 | 99 |
| SQSTM1 | rep2 | 99 | 95 | 99 | 95 | 100 | 99 | 99 | 94 | 99 |
| PSD95 | rep1 | 100 | 99 | 99 | 99 | 100 | 99 | 97 | 98 | 99 |
| PSD95 | rep2 | 100 | 100 | 99 | 100 | 99 | 99 | 100 | 97 | 98 |
| NEMO | rep1 | 100 | 100 | 98 | 100 | 100 | 100 | 100 | 100 | 99 |
| NEMO | rep2 | 100 | 100 | 100 | 89 | 100 | 100 | 100 | 100 | 100 |
| GM130 | rep1 | 100 | 100 | 100 | 100 | 99 | 100 | 100 | 100 | 96 |
| GM130 | rep2 | 100 | 99 | 100 | 100 | 100 | 99 | 100 | 100 | 95 |
| NCL | rep1 | 100 | 100 | 100 | 99 | 100 | 100 | 96 | 98 | 75 |
| NCL | rep2 | 100 | 99 | 100 | 100 | 100 | 98 | 100 | 98 | 89 |
| ANXA11 | rep1 | 100 | 100 | 100 | 100 | 100 | 100 | 100 | 98 | 95 |
| ANXA11 | rep2 | 100 | 100 | 95 | 100 | 100 | 100 | 100 | 95 | 100 |
| Calreticulin | rep1 | 100 | 98 | 100 | 100 | 100 | 100 | 98 | 99 | 100 |
| Calreticulin | rep2 | 100 | 100 | 100 | 100 | 100 | 100 | 98 | 100 | 100 |
| mitotracker | rep1 | 100 | 93 | 100 | 100 | 98 | 99 | 98 | 99 | 98 |
| mitotracker | rep2 | 100 | 99 | 100 | 100 | 100 | 100 | 100 | 100 | 99 |
| KIF5A | rep1 | 97 | 93 | 94 | 100 | 96 | 99 | 97 | 98 | 91 |
| KIF5A | rep2 | 99 | 98 | 98 | 96 | 99 | 99 | 97 | 100 | 100 |
| TDP43 | rep1 | 100 | 100 | 100 | 98 | 100 | 100 | 99 | 100 | 98 |
| TDP43 | rep2 | 100 | 100 | 100 | 99 | 100 | 100 | 100 | 100 | 100 |
| FMRP | rep1 | 100 | 100 | 100 | 99 | 100 | 100 | 98 | 99 | 99 |
| FMRP | rep2 | 100 | 99 | 100 | 97 | 100 | 100 | 97 | 99 | 100 |
| CLTC | rep1 | 100 | 99 | 99 | 99 | 100 | 100 | 97 | 98 | 99 |
| CLTC | rep2 | 100 | 100 | 100 | 100 | 99 | 100 | 100 | 97 | 99 |
| DCP1A | rep1 | 98 | 94 | 94 | 96 | 88 | 81 | 62 | 64 | 55 |
| DCP1A | rep2 | 94 | 92 | 94 | 77 | 87 | 81 | 65 | 55 | 22 |
| TOMM20 | rep1 | 100 | 100 | 100 | 100 | 100 | 100 | 100 | 100 | 99 |
| TOMM20 | rep2 | 100 | 99 | 100 | 100 | 100 | 100 | 100 | 100 | 99 |
| FUS | rep1 | 100 | 100 | 100 | 100 | 100 | 100 | 96 | 100 | 87 |
| FUS | rep2 | 100 | 99 | 100 | 100 | 100 | 100 | 100 | 99 | 98 |
| SCNA | rep1 | 91 | 83 | 87 | 85 | 89 | 73 | 93 | 86 | 90 |
| SCNA | rep2 | 88 | 88 | 79 | 89 | 86 | 83 | 96 | 76 | 92 |
| LAMP1 | rep1 | 96 | 94 | 89 | 95 | 99 | 90 | 97 | 89 | 92 |
| LAMP1 | rep2 | 96 | 91 | 90 | 95 | 100 | 90 | 94 | 93 | 92 |
| TIA1 | rep1 | nan | nan | nan | nan | nan | nan | nan | nan | nan |
| TIA1 | rep2 | nan | nan | nan | nan | nan | nan | nan | nan | nan |
| PML | rep1 | 76 | 98 | 48 | 100 | 99 | 74 | 92 | 93 | 66 |
| PML | rep2 | 91 | 99 | 53 | 100 | 100 | 71 | 100 | 85 | 79 |
| PURA | rep1 | 100 | 92 | 97 | 100 | 97 | 98 | 96 | 99 | 89 |
| PURA | rep2 | 99 | 99 | 99 | 100 | 100 | 100 | 100 | 100 | 100 |
| CD41 | rep1 | 100 | 100 | 100 | 98 | 100 | 100 | 99 | 100 | 98 |
| CD41 | rep2 | 100 | 100 | 100 | 95 | 100 | 100 | 100 | 100 | 100 |
| Phalloidin | rep1 | 100 | 100 | 100 | 99 | 100 | 100 | 98 | 98 | 100 |
| Phalloidin | rep2 | 100 | 99 | 100 | 98 | 100 | 100 | 100 | 100 | 100 |
| PEX14 | rep1 | 100 | 98 | 100 | 100 | 99 | 99 | 98 | 99 | 98 |
| PEX14 | rep2 | 100 | 99 | 100 | 100 | 100 | 100 | 100 | 100 | 98 |
| DAPI | rep1 | 1100 | 1082 | 1098 | 1098 | 1098 | 1098 | 1085 | 1091 | 1071 |
| DAPI | rep2 | 1099 | 1095 | 1094 | 1087 | 1099 | 1089 | 1098 | 1091 | 1095 |

### Batch 2.3

| | Rep | FUSHomozygous | TDP43 | TBK1 | WT stress | WT Untreated | FUSRevertant | OPTN | FUSHeterozygous | SNCA |
|---|---|---|---|---|---|---|---|---|---|---|
| G3BP1 | rep1 | 100 | 100 | 99 | 97 | 99 | 100 | 100 | 99 | 96 |
| G3BP1 | rep2 | 100 | 100 | 100 | 100 | 100 | 100 | 99 | 100 | 100 |
| NONO | rep1 | 100 | 100 | 100 | 100 | 100 | 99 | 100 | 100 | 100 |
| NONO | rep2 | 100 | 100 | 100 | 72 | 100 | 100 | 100 | 100 | 100 |
| SQSTM1 | rep1 | 98 | 99 | 98 | 99 | 100 | 96 | 99 | 95 | 93 |
| SQSTM1 | rep2 | 98 | 98 | 100 | 97 | 100 | 93 | 99 | 100 | 97 |
| PSD95 | rep1 | 100 | 97 | 99 | 98 | 100 | 100 | 96 | 98 | 96 |
| PSD95 | rep2 | 100 | 98 | 100 | 100 | 99 | 55 | 98 | 98 | 100 |
| NEMO | rep1 | 100 | 100 | 99 | 100 | 100 | 98 | 100 | 99 | 74 |
| NEMO | rep2 | 100 | 100 | 100 | 100 | 100 | 78 | 100 | 100 | 96 |
| GM130 | rep1 | 100 | 100 | 100 | 99 | 95 | 100 | 91 | 100 | 99 |
| GM130 | rep2 | 100 | 98 | 100 | 97 | 100 | 100 | 87 | 97 | 97 |
| NCL | rep1 | 100 | 98 | 100 | 49 | 88 | 100 | 85 | 96 | 93 |
| NCL | rep2 | 100 | 99 | 97 | 55 | 95 | 100 | 92 | 96 | 91 |
| ANXA11 | rep1 | 100 | 100 | 100 | 100 | 99 | 100 | 100 | 100 | 100 |
| ANXA11 | rep2 | 100 | 100 | 100 | 99 | 100 | 100 | 100 | 100 | 100 |
| Calreticulin | rep1 | 100 | 100 | 100 | 100 | 97 | 100 | 99 | 100 | 98 |
| Calreticulin | rep2 | 100 | 100 | 100 | 100 | 100 | 100 | 98 | 100 | 100 |
| mitotracker | rep1 | 100 | 98 | 100 | 100 | 97 | 100 | 100 | 100 | 100 |
| mitotracker | rep2 | 100 | 99 | 100 | 88 | 99 | 100 | 100 | 100 | 100 |
| KIF5A | rep1 | 98 | 96 | 95 | 96 | 99 | 100 | 95 | 99 | 92 |
| KIF5A | rep2 | 100 | 98 | 98 | 99 | 100 | 97 | 96 | 100 | 96 |
| TDP43 | rep1 | 100 | 100 | 100 | 100 | 100 | 99 | 100 | 100 | 100 |
| TDP43 | rep2 | 100 | 100 | 100 | 73 | 100 | 100 | 100 | 100 | 100 |
| FMRP | rep1 | 99 | 99 | 99 | 100 | 97 | 99 | 99 | 95 | 100 |
| FMRP | rep2 | 100 | 100 | 100 | 97 | 99 | 100 | 100 | 98 | 98 |
| CLTC | rep1 | 100 | 97 | 99 | 99 | 100 | 100 | 96 | 98 | 97 |
| CLTC | rep2 | 100 | 98 | 100 | 100 | 100 | 98 | 98 | 98 | 100 |
| DCP1A | rep1 | 90 | 93 | 74 | 92 | 63 | 52 | 21 | 17 | 0 |
| DCP1A | rep2 | 87 | 68 | 81 | 86 | 35 | 19 | 11 | 29 | 6 |
| TOMM20 | rep1 | 100 | 100 | 100 | 100 | 100 | 100 | 99 | 100 | 100 |
| TOMM20 | rep2 | 100 | 99 | 100 | 99 | 100 | 100 | 93 | 98 | 97 |
| FUS | rep1 | 100 | 98 | 100 | 100 | 100 | 100 | 93 | 100 | 96 |
| FUS | rep2 | 98 | 100 | 100 | 99 | 99 | 100 | 98 | 100 | 98 |
| SCNA | rep1 | 95 | 95 | 96 | 90 | 87 | 89 | 91 | 97 | 95 |
| SCNA | rep2 | 95 | 97 | 88 | 90 | 91 | 95 | 97 | 94 | 91 |
| LAMP1 | rep1 | 99 | 96 | 98 | 93 | 93 | 96 | 90 | 96 | 88 |
| LAMP1 | rep2 | 97 | 85 | 95 | 83 | 96 | 97 | 88 | 94 | 86 |
| TIA1 | rep1 | nan | nan | nan | nan | nan | nan | nan | nan | nan |
| TIA1 | rep2 | nan | nan | nan | nan | nan | nan | nan | nan | nan |
| PML | rep1 | 98 | 98 | 96 | 99 | 99 | 96 | 100 | 97 | 93 |
| PML | rep2 | 95 | 100 | 95 | 95 | 96 | 91 | 97 | 98 | 92 |
| PURA | rep1 | 100 | 99 | 96 | 97 | 99 | 100 | 99 | 99 | 96 |
| PURA | rep2 | 99 | 99 | 97 | 100 | 100 | 99 | 98 | 100 | 99 |
| CD41 | rep1 | 99 | 100 | 98 | 100 | 100 | 98 | 100 | 100 | 100 |
| CD41 | rep2 | 100 | 100 | 100 | 44 | 100 | 100 | 100 | 100 | 100 |
| Phalloidin | rep1 | 100 | 100 | 100 | 99 | 100 | 100 | 100 | 98 | 100 |
| Phalloidin | rep2 | 100 | 100 | 100 | 98 | 100 | 100 | 100 | 100 | 100 |
| PEX14 | rep1 | 100 | 99 | 100 | 100 | 100 | 100 | 100 | 100 | 100 |
| PEX14 | rep2 | 100 | 100 | 100 | 92 | 100 | 100 | 100 | 100 | 100 |
| DAPI | rep1 | 1099 | 1082 | 1097 | 1096 | 1095 | 1097 | 1063 | 1095 | 1078 |
| DAPI | rep2 | 1100 | 1095 | 1098 | 1079 | 1099 | 1097 | 1086 | 1096 | 1095 |

### Batch 2.4

| | Rep | FUSHomozygous | TDP43 | TBK1 | WT stress | WT Untreated | FUSRevertant | OPTN | FUSHeterozygous | SNCA |
|---|---|---|---|---|---|---|---|---|---|---|
| G3BP1 | rep1 | 85 | 100 | 100 | 99 | 100 | 100 | 99 | 97 | 100 |
| G3BP1 | rep2 | 88 | 100 | 100 | 100 | 99 | 100 | 91 | 100 | 97 |
| NONO | rep1 | 100 | 100 | 100 | 100 | 100 | 100 | 100 | 100 | 100 |
| NONO | rep2 | 100 | 100 | 100 | 100 | 100 | 100 | 100 | 100 | 100 |
| SQSTM1 | rep1 | 99 | 100 | 99 | 93 | 100 | 96 | 100 | 91 | 96 |
| SQSTM1 | rep2 | 99 | 97 | 99 | 95 | 99 | 95 | 99 | 100 | 95 |
| PSD95 | rep1 | 100 | 99 | 100 | 100 | 100 | 100 | 99 | 99 | 98 |
| PSD95 | rep2 | 100 | 99 | 100 | 100 | 100 | 100 | 99 | 100 | 100 |
| NEMO | rep1 | 100 | 100 | 100 | 97 | 100 | 100 | 99 | 100 | 99 |
| NEMO | rep2 | 100 | 100 | 100 | 100 | 100 | 100 | 100 | 100 | 100 |
| GM130 | rep1 | 100 | 100 | 100 | 100 | 100 | 0 | 93 | 97 | 93 |
| GM130 | rep2 | 100 | 77 | 100 | 35 | 99 | 100 | 97 | 99 | 87 |
| NCL | rep1 | 96 | 75 | 92 | 42 | 88 | 87 | 89 | 52 | 89 |
| NCL | rep2 | 96 | 95 | 96 | 73 | 94 | 93 | 71 | 65 | 66 |
| ANXA11 | rep1 | 100 | 100 | 100 | 100 | 100 | 100 | 94 | 99 | 98 |
| ANXA11 | rep2 | 100 | 100 | 100 | 100 | 100 | 100 | 100 | 100 | 99 |
| Calreticulin | rep1 | 100 | 100 | 100 | 100 | 100 | 100 | 100 | 100 | 97 |
| Calreticulin | rep2 | 100 | 98 | 100 | 100 | 100 | 100 | 99 | 99 | 100 |
| mitotracker | rep1 | 99 | 100 | 99 | 87 | 100 | 100 | 100 | 99 | 100 |
| mitotracker | rep2 | 100 | 99 | 100 | 100 | 100 | 100 | 100 | 100 | 100 |
| KIF5A | rep1 | 85 | 99 | 99 | 99 | 99 | 97 | 95 | 97 | 99 |
| KIF5A | rep2 | 86 | 96 | 97 | 99 | 97 | 100 | 88 | 100 | 97 |
| TDP43 | rep1 | 100 | 100 | 100 | 100 | 100 | 100 | 100 | 100 | 100 |
| TDP43 | rep2 | 100 | 100 | 100 | 100 | 100 | 100 | 100 | 100 | 100 |
| FMRP | rep1 | 98 | 98 | 99 | 91 | 99 | 100 | 98 | 99 | 99 |
| FMRP | rep2 | 100 | 98 | 100 | 96 | 99 | 99 | 99 | 100 | 100 |
| CLTC | rep1 | 100 | 100 | 100 | 100 | 100 | 100 | 100 | 99 | 98 |
| CLTC | rep2 | 100 | 100 | 100 | 100 | 100 | 100 | 100 | 100 | 100 |
| DCP1A | rep1 | 99 | 91 | 100 | 99 | 91 | 99 | 90 | 93 | 70 |
| DCP1A | rep2 | 100 | 100 | 99 | 100 | 96 | 99 | 37 | 93 | 63 |
| TOMM20 | rep1 | 100 | 99 | 100 | 100 | 100 | 0 | 100 | 97 | 98 |
| TOMM20 | rep2 | 100 | 84 | 100 | 69 | 99 | 100 | 97 | 100 | 98 |
| FUS | rep1 | 75 | 96 | 100 | 96 | 99 | 100 | 95 | 88 | 91 |
| FUS | rep2 | 92 | 100 | 100 | 100 | 100 | 100 | 100 | 29 | 99 |
| SCNA | rep1 | 95 | 83 | 93 | 83 | 88 | 91 | 93 | 93 | 96 |
| SCNA | rep2 | 94 | 95 | 91 | 90 | 88 | 92 | 95 | 92 | 96 |
| LAMP1 | rep1 | 95 | 97 | 100 | 99 | 100 | 97 | 98 | 96 | 96 |
| LAMP1 | rep2 | 96 | 89 | 97 | 92 | 96 | 96 | 96 | 87 | 93 |
| TIA1 | rep1 | nan | nan | nan | nan | nan | nan | nan | nan | nan |
| TIA1 | rep2 | nan | nan | nan | nan | nan | nan | nan | nan | nan |
| PML | rep1 | 100 | 100 | 99 | 91 | 100 | 100 | 100 | 99 | 83 |
| PML | rep2 | 99 | 100 | 99 | 100 | 100 | 99 | 100 | 99 | 96 |
| PURA | rep1 | 85 | 97 | 100 | 99 | 100 | 99 | 98 | 97 | 100 |
| PURA | rep2 | 88 | 100 | 100 | 100 | 98 | 98 | 89 | 100 | 97 |
| CD41 | rep1 | 100 | 100 | 99 | 99 | 100 | 100 | 99 | 100 | 100 |
| CD41 | rep2 | 100 | 100 | 100 | 100 | 100 | 100 | 100 | 100 | 100 |
| Phalloidin | rep1 | 100 | 100 | 100 | 96 | 100 | 100 | 100 | 98 | 100 |
| Phalloidin | rep2 | 100 | 100 | 100 | 99 | 99 | 100 | 100 | 100 | 100 |
| PEX14 | rep1 | 100 | 100 | 99 | 91 | 100 | 100 | 100 | 99 | 100 |
| PEX14 | rep2 | 100 | 98 | 100 | 100 | 100 | 100 | 100 | 100 | 100 |
| DAPI | rep1 | 1087 | 1096 | 1099 | 1086 | 1099 | 1000 | 1086 | 1084 | 1081 |
| DAPI | rep2 | 1088 | 1083 | 1100 | 1045 | 1099 | 1100 | 1087 | 1099 | 1093 |

#### **Total Tile Count**

Batch1

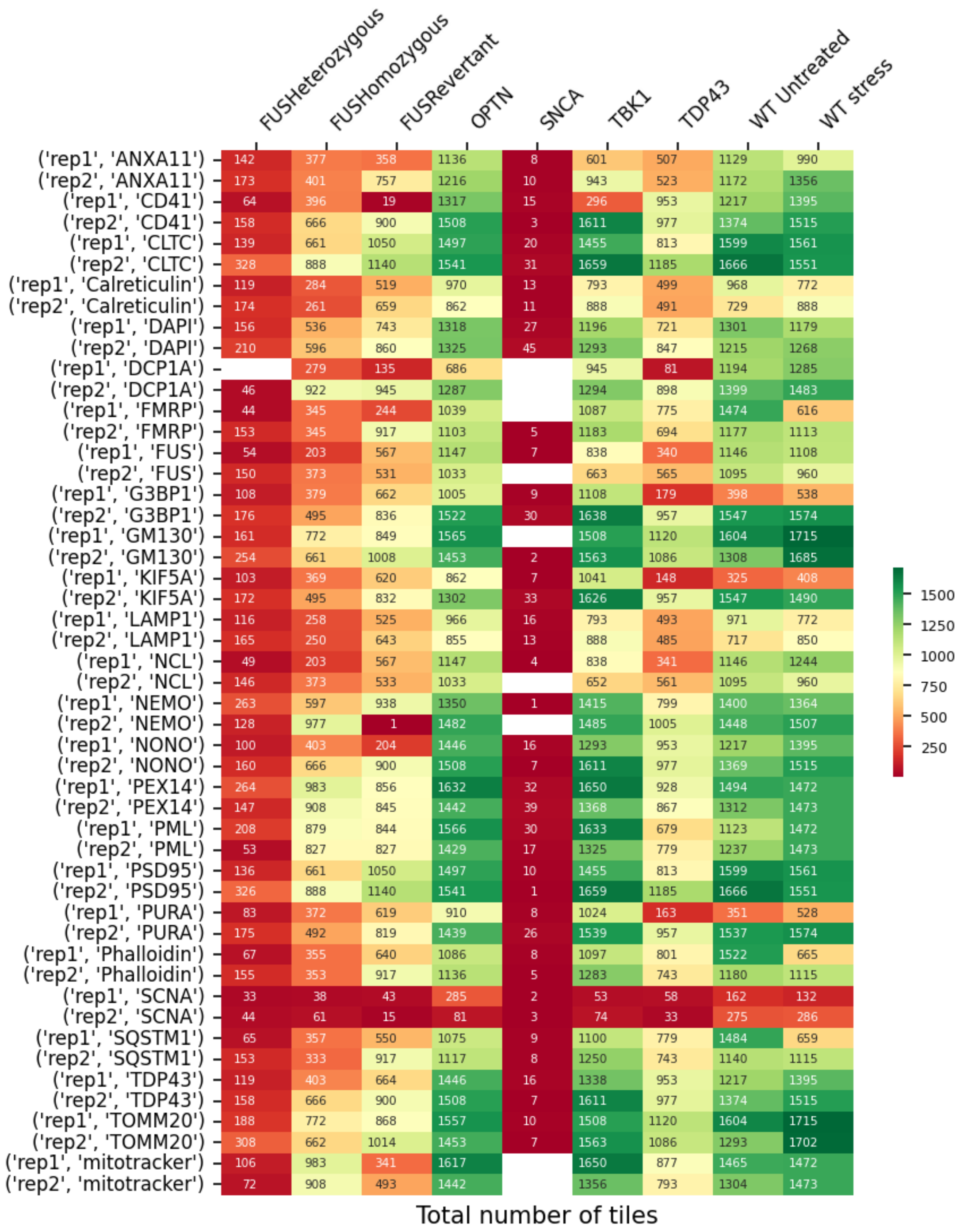

#### Batch 2.1

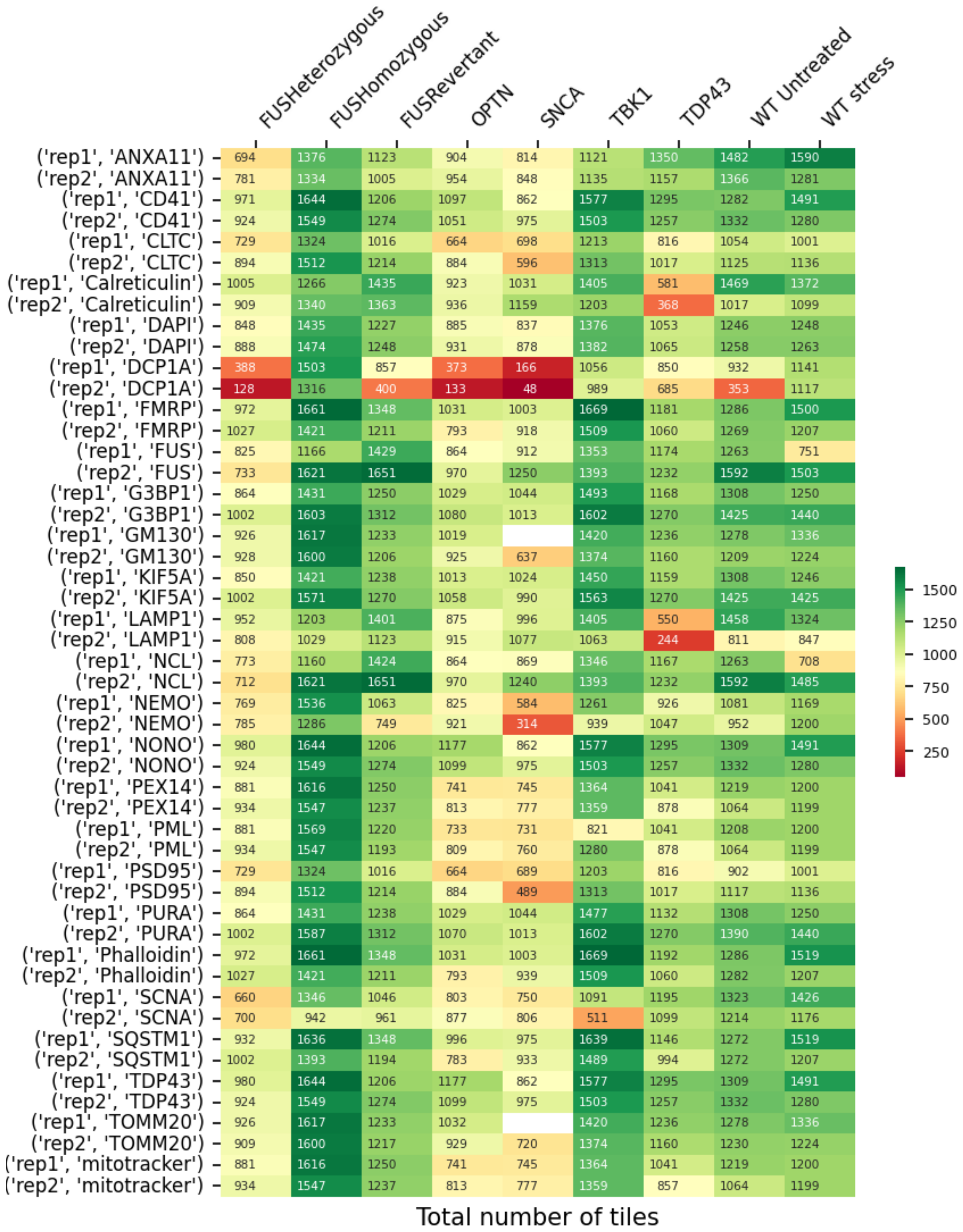
FUSHeterozygous
FUSHomozygous
FUSRevertant
OPTN
SNCA
TBK1
TDP43
WT Untreated
WT stress
('rep1', 'ANXA11') 694 1376 1123 904 814 1121 1350 1482 1590
('rep2', 'ANXA11') 781 1334 1005 954 848 1135 1157 1366 1281
('rep1', 'CD41') 971 1644 1206 1097 862 1577 1295 1282 1491
('rep2', 'CD41') 924 1549 1274 1051 975 1503 1257 1332 1280
('rep1', 'CLTC') 729 1324 1016 664 698 1213 816 1054 1001
('rep2', 'CLTC') 894 1512 1214 884 596 1313 1017 1125 1136
('rep1', 'Calreticulin') 1005 1266 1435 923 1031 1405 581 1469 1372
('rep2', 'Calreticulin') 909 1340 1363 936 1159 1203 368 1017 1099
('rep1', 'DAPI') 848 1435 1227 885 837 1376 1053 1246 1248
('rep2', 'DAPI') 888 1474 1248 931 878 1382 1065 1258 1263
('rep1', 'DCP1A') 388 1503 857 373 166 1056 850 932 1141
('rep2', 'DCP1A') 128 1316 400 133 48 989 685 353 1117
('rep1', 'FMRP') 972 1661 1348 1031 1003 1669 1181 1286 1500
('rep2', 'FMRP') 1027 1421 1211 793 918 1509 1060 1269 1207
('rep1', 'FUS') 825 1166 1429 864 912 1353 1174 1263 751
('rep2', 'FUS') 733 1621 1651 970 1250 1393 1232 1592 1503
('rep1', 'G3BP1') 864 1431 1250 1029 1044 1493 1168 1308 1250
('rep2', 'G3BP1') 1002 1603 1312 1080 1013 1602 1270 1425 1440
('rep1', 'GM130') 926 1617 1233 1019 1420 1236 1278 1336
('rep2', 'GM130') 928 1600 1206 925 637 1374 1160 1209 1224
('rep1', 'KIF5A') 850 1421 1238 1013 1024 1450 1159 1308 1246
('rep2', 'KIF5A') 1002 1571 1270 1058 990 1563 1270 1425 1425
('rep1', 'LAMP1') 952 1203 1401 875 996 1405 550 1458 1324
('rep2', 'LAMP1') 808 1029 1123 915 1077 1063 244 811 847
('rep1', 'NCL') 773 1160 1424 864 869 1346 1167 1263 708
('rep2', 'NCL') 712 1621 1651 970 1240 1393 1232 1592 1485
('rep1', 'NEMO') 769 1536 1063 825 584 1261 926 1081 1169
('rep2', 'NEMO') 785 1286 749 921 314 939 1047 952 1200
('rep1', 'NONO') 980 1644 1206 1177 862 1577 1295 1309 1491
('rep2', 'NONO') 924 1549 1274 1099 975 1503 1257 1332 1280
('rep1', 'PEX14') 881 1616 1250 741 745 1364 1041 1219 1200
('rep2', 'PEX14') 934 1547 1237 813 777 1359 878 1064 1199
('rep1', 'PML') 881 1569 1220 733 731 821 1041 1208 1200
('rep2', 'PML') 934 1547 1193 809 760 1280 878 1064 1199
('rep1', 'PSD95') 729 1324 1016 664 689 1203 816 902 1001
('rep2', 'PSD95') 894 1512 1214 884 489 1313 1017 1117 1136
('rep1', 'PURA') 864 1431 1238 1029 1044 1477 1132 1308 1250
('rep2', 'PURA') 1002 1587 1312 1070 1013 1602 1270 1390 1440
('rep1', 'Phalloidin') 972 1661 1348 1031 1003 1669 1192 1286 1519
('rep2', 'Phalloidin') 1027 1421 1211 793 939 1509 1060 1282 1207
('rep1', 'SCNA') 660 1346 1046 803 750 1091 1195 1323 1426
('rep2', 'SCNA') 700 942 961 877 806 511 1099 1214 1176
('rep1', 'SQSTM1') 932 1636 1348 996 975 1639 1146 1272 1519
('rep2', 'SQSTM1') 1002 1393 1194 783 933 1489 994 1272 1207
('rep1', 'TDP43') 980 1644 1206 1177 862 1577 1295 1309 1491
('rep2', 'TDP43') 924 1549 1274 1099 975 1503 1257 1332 1280
('rep1', 'TOMM20') 926 1617 1233 1032 1420 1236 1278 1336
('rep2', 'TOMM20') 909 1600 1217 929 720 1374 1160 1230 1224
('rep1', 'mitotracker') 881 1616 1250 741 745 1364 1041 1219 1200
('rep2', 'mitotracker') 934 1547 1237 813 777 1359 857 1064 1199
1500
1250
1000
750
500
250
Total number of tiles

Batch 2.2

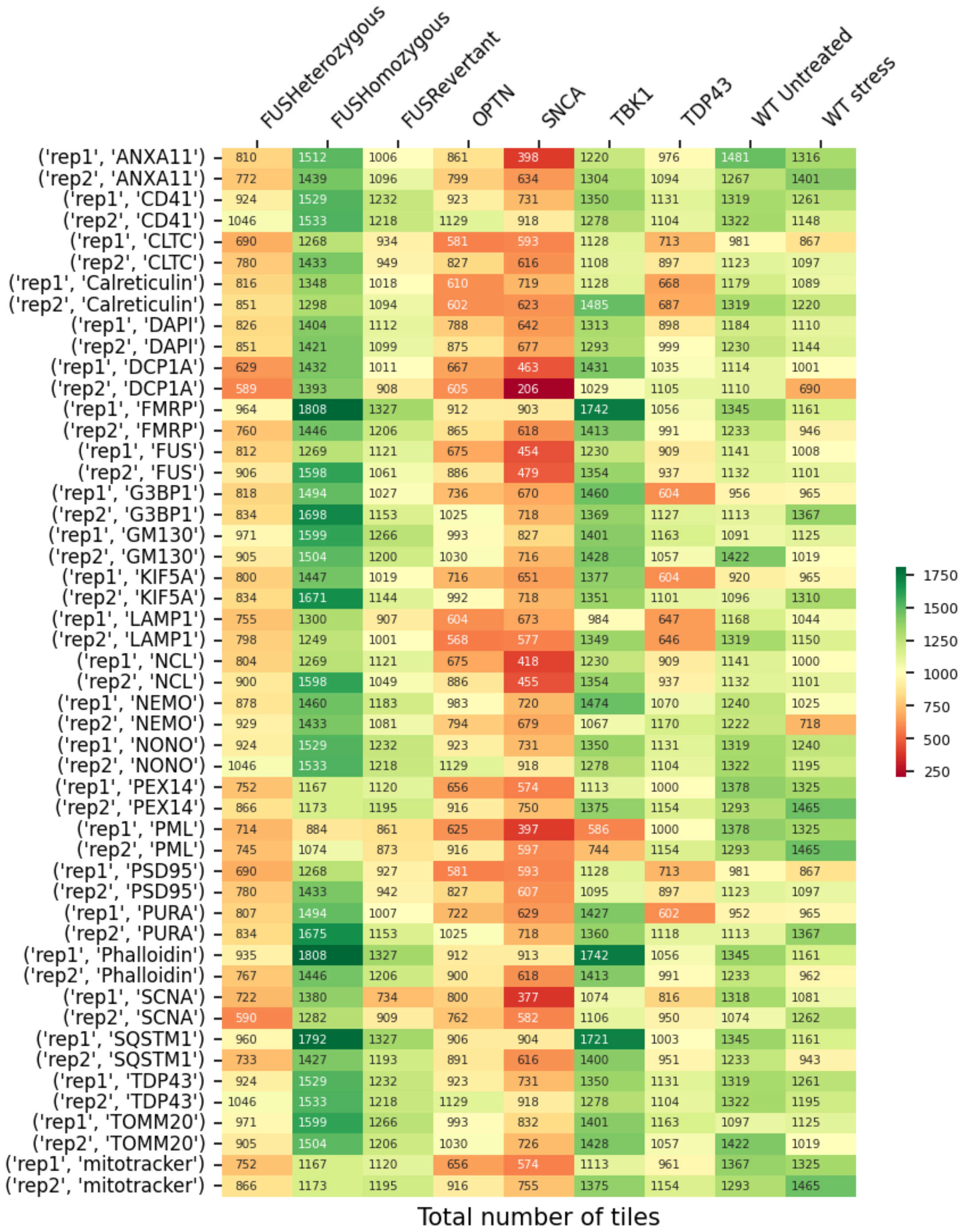


| | FUSHeterozygous | FUSHomozygous | FUSRevertant | OPTN | SNCA | TBK1 | TDP43 | WT Untreated | WT stress |
|---|---|---|---|---|---|---|---|---|---|
| ('rep1', 'ANXA11') | 810 | 1512 | 1006 | 861 | 398 | 1220 | 976 | 1481 | 1316 |
| ('rep2', 'ANXA11') | 772 | 1439 | 1096 | 799 | 634 | 1304 | 1094 | 1267 | 1401 |
| ('rep1', 'CD41') | 924 | 1529 | 1232 | 923 | 731 | 1350 | 1131 | 1319 | 1261 |
| ('rep2', 'CD41') | 1046 | 1533 | 1218 | 1129 | 918 | 1278 | 1104 | 1322 | 1148 |
| ('rep1', 'CLTC') | 690 | 1268 | 934 | 581 | 593 | 1128 | 713 | 981 | 867 |
| ('rep2', 'CLTC') | 780 | 1433 | 949 | 827 | 616 | 1108 | 897 | 1123 | 1097 |
| ('rep1', 'Calreticulin') | 816 | 1348 | 1018 | 610 | 719 | 1128 | 668 | 1179 | 1089 |
| ('rep2', 'Calreticulin') | 851 | 1298 | 1094 | 602 | 623 | 1485 | 687 | 1319 | 1220 |
| ('rep1', 'DAPI') | 826 | 1404 | 1112 | 788 | 642 | 1313 | 898 | 1184 | 1110 |
| ('rep2', 'DAPI') | 851 | 1421 | 1099 | 875 | 677 | 1293 | 999 | 1230 | 1144 |
| ('rep1', 'DCP1A') | 629 | 1432 | 1011 | 667 | 463 | 1431 | 1035 | 1114 | 1001 |
| ('rep2', 'DCP1A') | 589 | 1393 | 908 | 605 | 206 | 1029 | 1105 | 1110 | 690 |
| ('rep1', 'FMRP') | 964 | 1808 | 1327 | 912 | 903 | 1742 | 1056 | 1345 | 1161 |
| ('rep2', 'FMRP') | 760 | 1446 | 1206 | 865 | 618 | 1413 | 991 | 1233 | 946 |
| ('rep1', 'FUS') | 812 | 1269 | 1121 | 675 | 454 | 1230 | 909 | 1141 | 1008 |
| ('rep2', 'FUS') | 906 | 1598 | 1061 | 886 | 479 | 1354 | 937 | 1132 | 1101 |
| ('rep1', 'G3BP1') | 818 | 1494 | 1027 | 736 | 670 | 1460 | 604 | 956 | 965 |
| ('rep2', 'G3BP1') | 834 | 1698 | 1153 | 1025 | 718 | 1369 | 1127 | 1113 | 1367 |
| ('rep1', 'GM130') | 971 | 1599 | 1266 | 993 | 827 | 1401 | 1163 | 1091 | 1125 |
| ('rep2', 'GM130') | 905 | 1504 | 1200 | 1030 | 716 | 1428 | 1057 | 1422 | 1019 |
| ('rep1', 'KIF5A') | 800 | 1447 | 1019 | 716 | 651 | 1377 | 604 | 920 | 965 |
| ('rep2', 'KIF5A') | 834 | 1671 | 1144 | 992 | 718 | 1351 | 1101 | 1096 | 1310 |
| ('rep1', 'LAMP1') | 755 | 1300 | 907 | 604 | 673 | 984 | 647 | 1168 | 1044 |
| ('rep2', 'LAMP1') | 798 | 1249 | 1001 | 568 | 577 | 1349 | 646 | 1319 | 1150 |
| ('rep1', 'NCL') | 804 | 1269 | 1121 | 675 | 418 | 1230 | 909 | 1141 | 1000 |
| ('rep2', 'NCL') | 900 | 1598 | 1049 | 886 | 455 | 1354 | 937 | 1132 | 1101 |
| ('rep1', 'NEMO') | 878 | 1460 | 1183 | 983 | 720 | 1474 | 1070 | 1240 | 1025 |
| ('rep2', 'NEMO') | 929 | 1433 | 1081 | 794 | 679 | 1067 | 1170 | 1222 | 718 |
| ('rep1', 'NONO') | 924 | 1529 | 1232 | 923 | 731 | 1350 | 1131 | 1319 | 1240 |
| ('rep2', 'NONO') | 1046 | 1533 | 1218 | 1129 | 918 | 1278 | 1104 | 1322 | 1195 |
| ('rep1', 'PEX14') | 752 | 1167 | 1120 | 656 | 574 | 1113 | 1000 | 1378 | 1325 |
| ('rep2', 'PEX14') | 866 | 1173 | 1195 | 916 | 750 | 1375 | 1154 | 1293 | 1465 |
| ('rep1', 'PML') | 714 | 884 | 861 | 625 | 397 | 586 | 1000 | 1378 | 1325 |
| ('rep2', 'PML') | 745 | 1074 | 873 | 916 | 597 | 744 | 1154 | 1293 | 1465 |
| ('rep1', 'PSD95') | 690 | 1268 | 927 | 581 | 593 | 1128 | 713 | 981 | 867 |
| ('rep2', 'PSD95') | 780 | 1433 | 942 | 827 | 607 | 1095 | 897 | 1123 | 1097 |
| ('rep1', 'PURA') | 807 | 1494 | 1007 | 722 | 629 | 1427 | 602 | 952 | 965 |
| ('rep2', 'PURA') | 834 | 1675 | 1153 | 1025 | 718 | 1360 | 1118 | 1113 | 1367 |
| ('rep1', 'Phalloidin') | 935 | 1808 | 1327 | 912 | 913 | 1742 | 1056 | 1345 | 1161 |
| ('rep2', 'Phalloidin') | 767 | 1446 | 1206 | 900 | 618 | 1413 | 991 | 1233 | 962 |
| ('rep1', 'SCNA') | 722 | 1380 | 734 | 800 | 377 | 1074 | 816 | 1318 | 1081 |
| ('rep2', 'SCNA') | 590 | 1282 | 909 | 762 | 582 | 1106 | 950 | 1074 | 1262 |
| ('rep1', 'SQSTM1') | 960 | 1792 | 1327 | 906 | 904 | 1721 | 1003 | 1345 | 1161 |
| ('rep2', 'SQSTM1') | 733 | 1427 | 1193 | 891 | 616 | 1400 | 951 | 1233 | 943 |
| ('rep1', 'TDP43') | 924 | 1529 | 1232 | 923 | 731 | 1350 | 1131 | 1319 | 1261 |
| ('rep2', 'TDP43') | 1046 | 1533 | 1218 | 1129 | 918 | 1278 | 1104 | 1322 | 1195 |
| ('rep1', 'TOMM20') | 971 | 1599 | 1266 | 993 | 832 | 1401 | 1163 | 1097 | 1125 |
| ('rep2', 'TOMM20') | 905 | 1504 | 1206 | 1030 | 726 | 1428 | 1057 | 1422 | 1019 |
| ('rep1', 'mitotracker') | 752 | 1167 | 1120 | 656 | 574 | 1113 | 961 | 1367 | 1325 |
| ('rep2', 'mitotracker') | 866 | 1173 | 1195 | 916 | 755 | 1375 | 1154 | 1293 | 1465 |

Batch 2.3

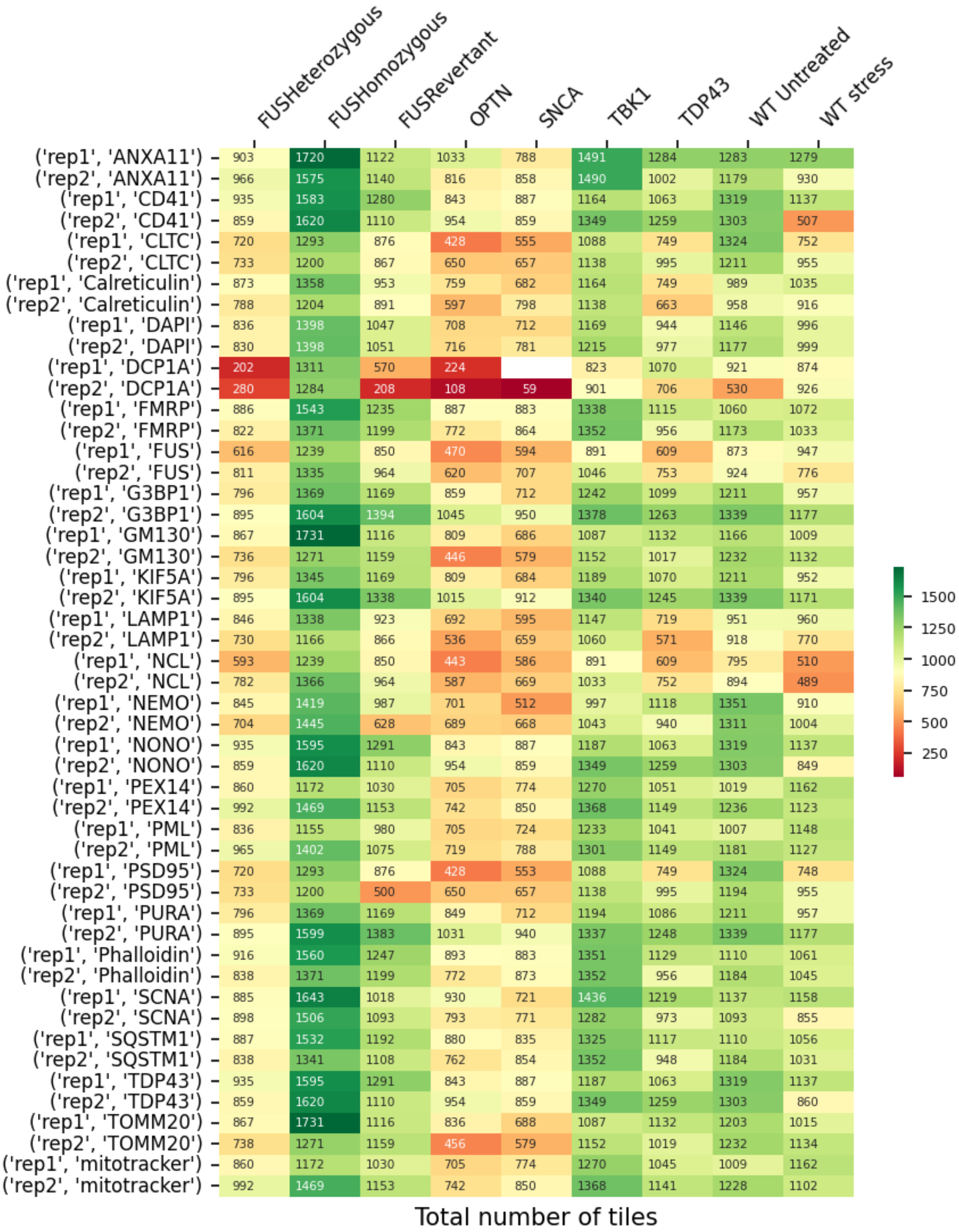
FUSHeterozygous FUSHomozygous FUSRevertant OPTN SNCA TBK1 TDP43 WT Untreated WT stress
('rep1', 'ANXA11') 903 1720 1122 1033 788 1491 1284 1283 1279
('rep2', 'ANXA11') 966 1575 1140 816 858 1490 1002 1179 930
('rep1', 'CD41') 935 1583 1280 843 887 1164 1063 1319 1137
('rep2', 'CD41') 859 1620 1110 954 859 1349 1259 1303 507
('rep1', 'CLTC') 720 1293 876 428 555 1088 749 1324 752
('rep2', 'CLTC') 733 1200 867 650 657 1138 995 1211 955
('rep1', 'Calreticulin') 873 1358 953 759 682 1164 749 989 1035
('rep2', 'Calreticulin') 788 1204 891 597 798 1138 663 958 916
('rep1', 'DAPI') 836 1398 1047 708 712 1169 944 1146 996
('rep2', 'DAPI') 830 1398 1051 716 781 1215 977 1177 999
('rep1', 'DCP1A') 202 1311 570 224 823 1070 921 874
('rep2', 'DCP1A') 280 1284 208 108 59 901 706 530 926
('rep1', 'FMRP') 886 1543 1235 887 883 1338 1115 1060 1072
('rep2', 'FMRP') 822 1371 1199 772 864 1352 956 1173 1033
('rep1', 'FUS') 616 1239 850 470 594 891 609 873 947
('rep2', 'FUS') 811 1335 964 620 707 1046 753 924 776
('rep1', 'G3BP1') 796 1369 1169 859 712 1242 1099 1211 957
('rep2', 'G3BP1') 895 1604 1394 1045 950 1378 1263 1339 1177
('rep1', 'GM130') 867 1731 1116 809 686 1087 1132 1166 1009
('rep2', 'GM130') 736 1271 1159 446 579 1152 1017 1232 1132
('rep1', 'KIF5A') 796 1345 1169 809 684 1189 1070 1211 952
('rep2', 'KIF5A') 895 1604 1338 1015 912 1340 1245 1339 1171
('rep1', 'LAMP1') 846 1338 923 692 595 1147 719 951 960
('rep2', 'LAMP1') 730 1166 866 536 659 1060 571 918 770
('rep1', 'NCL') 593 1239 850 443 586 891 609 795 510
('rep2', 'NCL') 782 1366 964 587 669 1033 752 894 489
('rep1', 'NEMO') 845 1419 987 701 512 997 1118 1351 910
('rep2', 'NEMO') 704 1445 628 689 668 1043 940 1311 1004
('rep1', 'NONO') 935 1595 1291 843 887 1187 1063 1319 1137
('rep2', 'NONO') 859 1620 1110 954 859 1349 1259 1303 849
('rep1', 'PEX14') 860 1172 1030 705 774 1270 1051 1019 1162
('rep2', 'PEX14') 992 1469 1153 742 850 1368 1149 1236 1123
('rep1', 'PML') 836 1155 980 705 724 1233 1041 1007 1148
('rep2', 'PML') 965 1402 1075 719 788 1301 1149 1181 1127
('rep1', 'PSD95') 720 1293 876 428 553 1088 749 1324 748
('rep2', 'PSD95') 733 1200 500 650 657 1138 995 1194 955
('rep1', 'PURA') 796 1369 1169 849 712 1194 1086 1211 957
('rep2', 'PURA') 895 1599 1383 1031 940 1337 1248 1339 1177
('rep1', 'Phalloidin') 916 1560 1247 893 883 1351 1129 1110 1061
('rep2', 'Phalloidin') 838 1371 1199 772 873 1352 956 1184 1045
('rep1', 'SCNA') 885 1643 1018 930 721 1436 1219 1137 1158
('rep2', 'SCNA') 898 1506 1093 793 771 1282 973 1093 855
('rep1', 'SQSTM1') 887 1532 1192 880 835 1325 1117 1110 1056
('rep2', 'SQSTM1') 838 1341 1108 762 854 1352 948 1184 1031
('rep1', 'TDP43') 935 1595 1291 843 887 1187 1063 1319 1137
('rep2', 'TDP43') 859 1620 1110 954 859 1349 1259 1303 860
('rep1', 'TOMM20') 867 1731 1116 836 688 1087 1132 1203 1015
('rep2', 'TOMM20') 738 1271 1159 456 579 1152 1019 1232 1134
('rep1', 'mitotracker') 860 1172 1030 705 774 1270 1045 1009 1162
('rep2', 'mitotracker') 992 1469 1153 742 850 1368 1141 1228 1102
1500
1250
1000
750
500
250
Total number of tiles

Batch 2.4

| | FUSHeterozygous | FUSHomozygous | FUSRevertant | OPTN | SNCA | TBK1 | TDP43 | WT Untreated | WT stress |
|---|---|---|---|---|---|---|---|---|---|
| ('rep1', 'ANXA11') | 858 | 1463 | 1435 | 619 | 570 | 1359 | 1311 | 1540 | 1328 |
| ('rep2', 'ANXA11') | 1080 | 1813 | 1375 | 1134 | 680 | 1573 | 1309 | 1634 | 1360 |
| ('rep1', 'CD41') | 1057 | 1660 | 1513 | 936 | 1035 | 1426 | 1105 | 1246 | 1278 |
| ('rep2', 'CD41') | 1082 | 1622 | 1498 | 1060 | 1043 | 1415 | 1295 | 1269 | 1319 |
| ('rep1', 'CLTC') | 768 | 1256 | 1172 | 628 | 731 | 1496 | 923 | 1054 | 962 |
| ('rep2', 'CLTC') | 1021 | 1597 | 1435 | 924 | 925 | 1581 | 1233 | 1229 | 1172 |
| ('rep1', 'Calreticulin') | 1010 | 1621 | 1325 | 934 | 757 | 1449 | 1125 | 1617 | 1402 |
| ('rep2', 'Calreticulin') | 813 | 1394 | 1261 | 751 | 808 | 1133 | 743 | 1387 | 1309 |
| ('rep1', 'DAPI') | 870 | 1472 | 1312 | 800 | 747 | 1447 | 1070 | 1266 | 1102 |
| ('rep2', 'DAPI') | 939 | 1582 | 1369 | 850 | 831 | 1420 | 1071 | 1262 | 1072 |
| ('rep1', 'DCP1A') | 1038 | 1522 | 1495 | 838 | 637 | 1583 | 1223 | 1187 | 1260 |
| ('rep2', 'DCP1A') | 885 | 1812 | 1562 | 410 | 557 | 1566 | 1375 | 1366 | 1299 |
| ('rep1', 'FMRP') | 883 | 1634 | 1269 | 1016 | 767 | 1339 | 1144 | 1208 | 984 |
| ('rep2', 'FMRP') | 865 | 1474 | 1225 | 844 | 921 | 1385 | 924 | 1193 | 1143 |
| ('rep1', 'FUS') | 431 | 876 | 1022 | 577 | 331 | 1274 | 718 | 1059 | 865 |
| ('rep2', 'FUS') | 231 | 1518 | 1219 | 753 | 718 | 1467 | 977 | 1319 | 1096 |
| ('rep1', 'G3BP1') | 702 | 1177 | 1183 | 748 | 819 | 1525 | 1039 | 1190 | 917 |
| ('rep2', 'G3BP1') | 997 | 1356 | 1333 | 730 | 1031 | 1524 | 1203 | 1246 | 1296 |
| ('rep1', 'GM130') | 1021 | 1539 | | 849 | 652 | 1477 | 1234 | 1379 | 1191 |
| ('rep2', 'GM130') | 783 | 1429 | 1252 | 656 | 637 | 1406 | 329 | 869 | 154 |
| ('rep1', 'KIF5A') | 702 | 1177 | 1154 | 723 | 808 | 1511 | 1022 | 1189 | 917 |
| ('rep2', 'KIF5A') | 997 | 1340 | 1333 | 700 | 1031 | 1475 | 1175 | 1222 | 1280 |
| ('rep1', 'LAMP1') | 973 | 1547 | 1285 | 907 | 755 | 1449 | 1086 | 1617 | 1387 |
| ('rep2', 'LAMP1') | 706 | 1329 | 1210 | 736 | 752 | 1105 | 683 | 1329 | 1205 |
| ('rep1', 'NCL') | 322 | 1215 | 932 | 553 | 324 | 1210 | 598 | 953 | 421 |
| ('rep2', 'NCL') | 638 | 1623 | 1159 | 576 | 535 | 1432 | 948 | 1264 | 847 |
| ('rep1', 'NEMO') | 1083 | 1529 | 1507 | 873 | 791 | 1583 | 1297 | 1268 | 1216 |
| ('rep2', 'NEMO') | 912 | 1812 | 1569 | 918 | 744 | 1567 | 1375 | 1400 | 1299 |
| ('rep1', 'NONO') | 1057 | 1660 | 1513 | 943 | 1035 | 1439 | 1105 | 1246 | 1285 |
| ('rep2', 'NONO') | 1082 | 1622 | 1498 | 1060 | 1043 | 1415 | 1295 | 1269 | 1319 |
| ('rep1', 'PEX14') | 973 | 1543 | 1467 | 918 | 1040 | 1462 | 1095 | 1174 | 743 |
| ('rep2', 'PEX14') | 1190 | 1530 | 1573 | 792 | 828 | 1395 | 1228 | 1353 | 1125 |
| ('rep1', 'PML') | 973 | 1543 | 1467 | 918 | 891 | 1462 | 1095 | 1174 | 743 |
| ('rep2', 'PML') | 1180 | 1514 | 1558 | 792 | 804 | 1388 | 1247 | 1353 | 1125 |
| ('rep1', 'PSD95') | 768 | 1256 | 1172 | 616 | 731 | 1496 | 915 | 1054 | 962 |
| ('rep2', 'PSD95') | 1021 | 1597 | 1435 | 914 | 925 | 1581 | 1218 | 1229 | 1172 |
| ('rep1', 'PURA') | 702 | 1177 | 1164 | 745 | 819 | 1525 | 1010 | 1190 | 917 |
| ('rep2', 'PURA') | 997 | 1356 | 1309 | 722 | 1031 | 1524 | 1203 | 1223 | 1296 |
| ('rep1', 'Phalloidin') | 878 | 1670 | 1269 | 1030 | 773 | 1359 | 1173 | 1220 | 1012 |
| ('rep2', 'Phalloidin') | 865 | 1474 | 1233 | 854 | 921 | 1385 | 949 | 1193 | 1160 |
| ('rep1', 'SCNA') | 808 | 1393 | 1333 | 604 | 554 | 1258 | 1090 | 1346 | 1111 |
| ('rep2', 'SCNA') | 987 | 1710 | 1252 | 1075 | 667 | 1430 | 1251 | 1434 | 1214 |
| ('rep1', 'SQSTM1') | 819 | 1656 | 1211 | 1030 | 754 | 1340 | 1173 | 1220 | 988 |
| ('rep2', 'SQSTM1') | 865 | 1459 | 1160 | 844 | 863 | 1371 | 929 | 1193 | 1125 |
| ('rep1', 'TDP43') | 1057 | 1660 | 1513 | 943 | 1035 | 1439 | 1105 | 1246 | 1285 |
| ('rep2', 'TDP43') | 1082 | 1622 | 1498 | 1060 | 1043 | 1415 | 1295 | 1269 | 1319 |
| ('rep1', 'TOMM20') | 1021 | 1539 | | 882 | 659 | 1477 | 1222 | 1379 | 1191 |
| ('rep2', 'TOMM20') | 785 | 1429 | 1252 | 656 | 686 | 1406 | 344 | 861 | 229 |
| ('rep1', 'mitotracker') | 973 | 1532 | 1467 | 918 | 1040 | 1462 | 1095 | 1174 | 733 |
| ('rep2', 'mitotracker') | 1190 | 1530 | 1573 | 792 | 828 | 1395 | 1242 | 1353 | 1125 |

Total number of tiles

#### **Whole Cell Count**

Batch 1

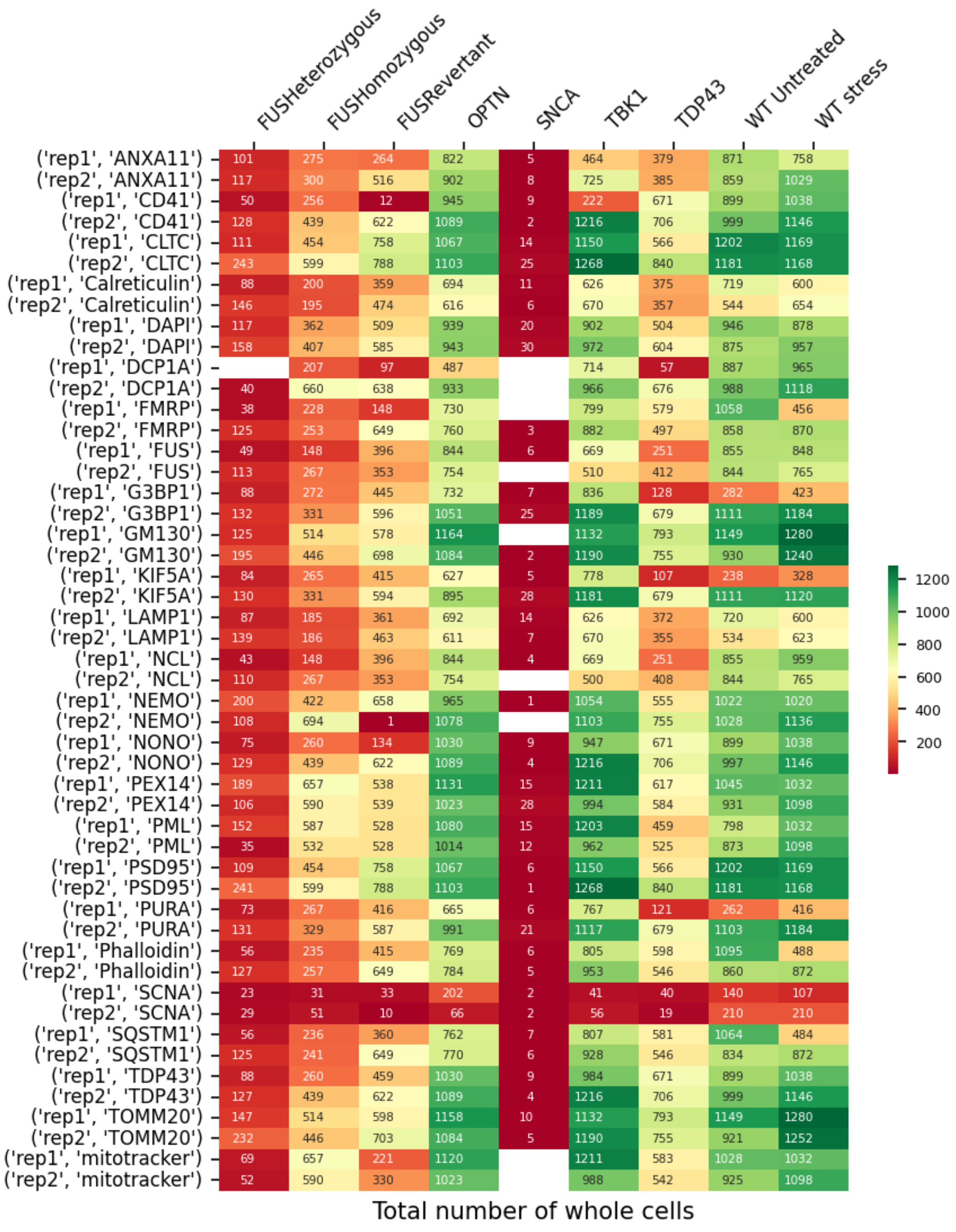

Batch 2.1

| | FUSHeterozygous | FUSHomozygous | FUSRevertant | OPTN | SNCA | TBK1 | TDP43 | WT Untreated | WT stress |
|---|---|---|---|---|---|---|---|---|---|
| ('rep1', 'ANXA11') | 434 | 933 | 719 | 619 | 565 | 779 | 863 | 907 | 1079 |
| ('rep2', 'ANXA11') | 493 | 902 | 655 | 669 | 567 | 767 | 762 | 864 | 851 |
| ('rep1', 'CD41') | 644 | 1083 | 718 | 710 | 575 | 1021 | 814 | 816 | 984 |
| ('rep2', 'CD41') | 563 | 1000 | 819 | 693 | 643 | 987 | 755 | 863 | 827 |
| ('rep1', 'CLTC') | 440 | 912 | 642 | 434 | 460 | 789 | 512 | 670 | 648 |
| ('rep2', 'CLTC') | 581 | 1014 | 836 | 615 | 415 | 874 | 616 | 714 | 749 |
| ('rep1', 'Calreticulin') | 610 | 862 | 962 | 628 | 728 | 959 | 351 | 971 | 876 |
| ('rep2', 'Calreticulin') | 601 | 893 | 921 | 637 | 862 | 811 | 236 | 694 | 807 |
| ('rep1', 'DAPI') | 540 | 974 | 783 | 583 | 564 | 938 | 660 | 796 | 812 |
| ('rep2', 'DAPI') | 562 | 970 | 808 | 632 | 599 | 941 | 680 | 810 | 846 |
| ('rep1', 'DCP1A') | 254 | 1050 | 555 | 254 | 104 | 734 | 549 | 599 | 686 |
| ('rep2', 'DCP1A') | 74 | 845 | 251 | 84 | 36 | 689 | 456 | 207 | 748 |
| ('rep1', 'FMRP') | 603 | 1086 | 856 | 667 | 689 | 1166 | 707 | 844 | 1020 |
| ('rep2', 'FMRP') | 661 | 887 | 736 | 497 | 612 | 993 | 625 | 798 | 769 |
| ('rep1', 'FUS') | 516 | 759 | 902 | 568 | 591 | 903 | 763 | 805 | 460 |
| ('rep2', 'FUS') | 403 | 1052 | 1099 | 651 | 857 | 932 | 760 | 1021 | 1040 |
| ('rep1', 'G3BP1') | 542 | 946 | 771 | 639 | 670 | 960 | 721 | 767 | 781 |
| ('rep2', 'G3BP1') | 588 | 1001 | 795 | 724 | 686 | 1036 | 810 | 871 | 913 |
| ('rep1', 'GM130') | 579 | 1059 | 765 | 673 | | 981 | 754 | 811 | 856 |
| ('rep2', 'GM130') | 588 | 1063 | 758 | 635 | 432 | 986 | 770 | 791 | 803 |
| ('rep1', 'KIF5A') | 538 | 937 | 763 | 629 | 659 | 932 | 715 | 767 | 777 |
| ('rep2', 'KIF5A') | 588 | 981 | 767 | 706 | 672 | 1011 | 810 | 871 | 906 |
| ('rep1', 'LAMP1') | 582 | 819 | 939 | 600 | 705 | 959 | 334 | 964 | 856 |
| ('rep2', 'LAMP1') | 535 | 683 | 749 | 624 | 799 | 714 | 157 | 572 | 609 |
| ('rep1', 'NCL') | 485 | 753 | 897 | 568 | 554 | 898 | 759 | 805 | 436 |
| ('rep2', 'NCL') | 389 | 1052 | 1099 | 651 | 850 | 932 | 760 | 1021 | 1028 |
| ('rep1', 'NEMO') | 489 | 1073 | 683 | 528 | 395 | 858 | 590 | 691 | 706 |
| ('rep2', 'NEMO') | 488 | 827 | 472 | 603 | 211 | 641 | 691 | 569 | 803 |
| ('rep1', 'NONO') | 650 | 1083 | 718 | 758 | 575 | 1021 | 814 | 831 | 984 |
| ('rep2', 'NONO') | 563 | 1000 | 819 | 724 | 643 | 987 | 755 | 863 | 827 |
| ('rep1', 'PEX14') | 612 | 1189 | 847 | 517 | 517 | 1028 | 689 | 846 | 825 |
| ('rep2', 'PEX14') | 635 | 1074 | 808 | 592 | 517 | 960 | 606 | 736 | 839 |
| ('rep1', 'PML') | 612 | 1154 | 831 | 511 | 506 | 618 | 689 | 840 | 825 |
| ('rep2', 'PML') | 635 | 1074 | 780 | 590 | 507 | 898 | 606 | 736 | 839 |
| ('rep1', 'PSD95') | 440 | 912 | 642 | 434 | 455 | 783 | 512 | 565 | 648 |
| ('rep2', 'PSD95') | 581 | 1014 | 836 | 615 | 343 | 874 | 616 | 709 | 749 |
| ('rep1', 'PURA') | 542 | 946 | 765 | 639 | 670 | 952 | 696 | 767 | 781 |
| ('rep2', 'PURA') | 588 | 990 | 795 | 720 | 686 | 1036 | 810 | 849 | 913 |
| ('rep1', 'Phalloidin') | 603 | 1086 | 856 | 667 | 689 | 1166 | 714 | 844 | 1032 |
| ('rep2', 'Phalloidin') | 661 | 887 | 736 | 497 | 624 | 993 | 625 | 803 | 769 |
| ('rep1', 'SCNA') | 411 | 912 | 674 | 551 | 519 | 757 | 755 | 810 | 966 |
| ('rep2', 'SCNA') | 448 | 621 | 621 | 620 | 540 | 342 | 729 | 764 | 782 |
| ('rep1', 'SQSTM1') | 576 | 1068 | 856 | 644 | 668 | 1141 | 687 | 832 | 1032 |
| ('rep2', 'SQSTM1') | 647 | 866 | 724 | 490 | 619 | 980 | 588 | 794 | 769 |
| ('rep1', 'TDP43') | 650 | 1083 | 718 | 758 | 575 | 1021 | 814 | 831 | 984 |
| ('rep2', 'TDP43') | 563 | 1000 | 819 | 724 | 643 | 987 | 755 | 863 | 827 |
| ('rep1', 'TOMM20') | 579 | 1059 | 765 | 678 | | 981 | 754 | 811 | 856 |
| ('rep2', 'TOMM20') | 581 | 1063 | 765 | 638 | 492 | 986 | 770 | 804 | 803 |
| ('rep1', 'mitotracker') | 612 | 1189 | 847 | 517 | 517 | 1028 | 689 | 846 | 825 |
| ('rep2', 'mitotracker') | 635 | 1074 | 808 | 592 | 517 | 960 | 595 | 736 | 839 |

Total number of whole cells

Batch 2.2

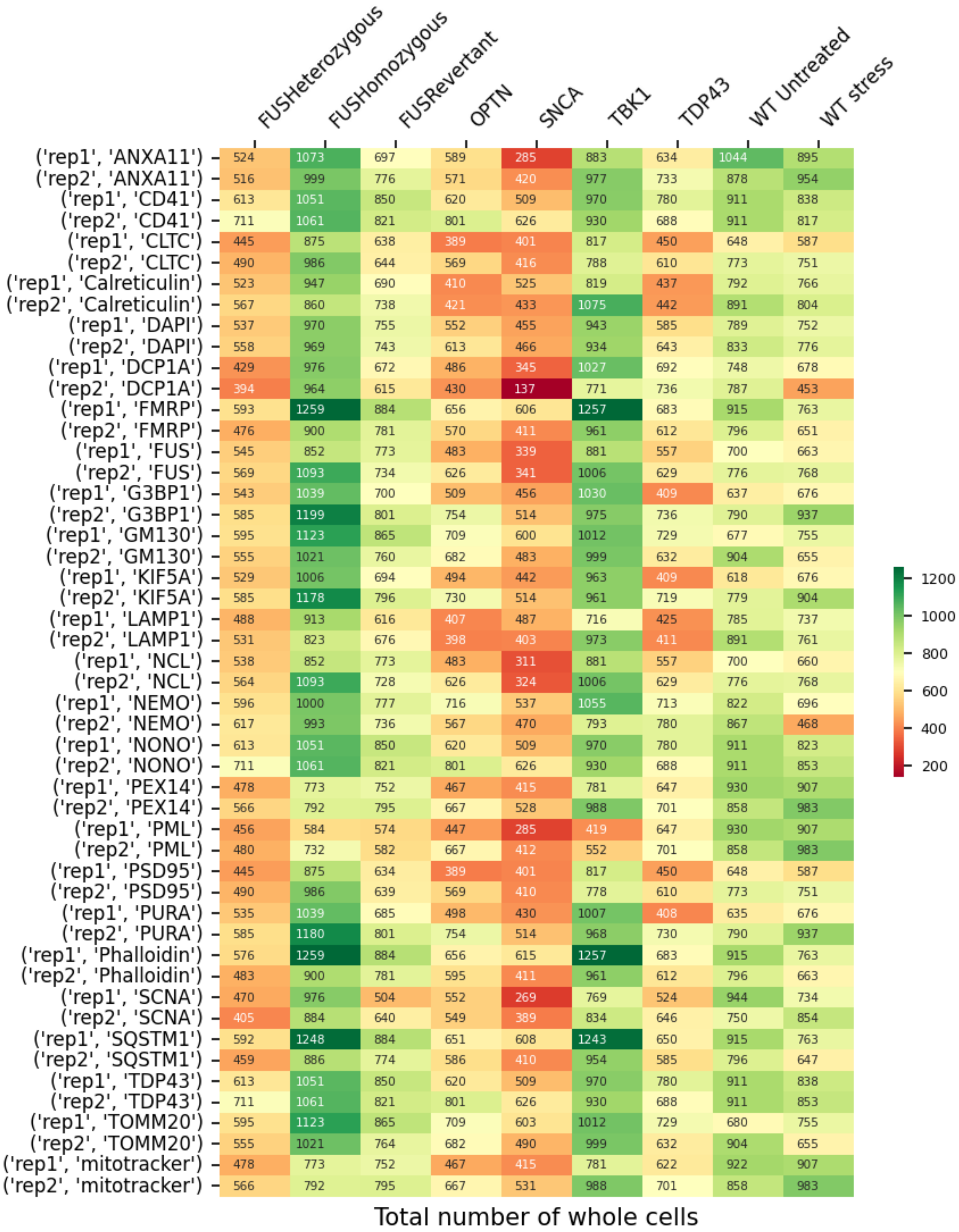
FUSHeterozygous
FUSHomozygous
FUSRevertant
OPTN
SNCA
TBK1
TDP43
WT Untreated
WT stress
('rep1', 'ANXA11') 524 1073 697 589 285 883 634 1044 895
('rep2', 'ANXA11') 516 999 776 571 420 977 733 878 954
('rep1', 'CD41') 613 1051 850 620 509 970 780 911 838
('rep2', 'CD41') 711 1061 821 801 626 930 688 911 817
('rep1', 'CLTC') 445 875 638 389 401 817 450 648 587
('rep2', 'CLTC') 490 986 644 569 416 788 610 773 751
('rep1', 'Calreticulin') 523 947 690 410 525 819 437 792 766
('rep2', 'Calreticulin') 567 860 738 421 433 1075 442 891 804
('rep1', 'DAPI') 537 970 755 552 455 943 585 789 752
('rep2', 'DAPI') 558 969 743 613 466 934 643 833 776
('rep1', 'DCP1A') 429 976 672 486 345 1027 692 748 678
('rep2', 'DCP1A') 394 964 615 430 137 771 736 787 453
('rep1', 'FMRP') 593 1259 884 656 606 1257 683 915 763
('rep2', 'FMRP') 476 900 781 570 411 961 612 796 651
('rep1', 'FUS') 545 852 773 483 339 881 557 700 663
('rep2', 'FUS') 569 1093 734 626 341 1006 629 776 768
('rep1', 'G3BP1') 543 1039 700 509 456 1030 409 637 676
('rep2', 'G3BP1') 585 1199 801 754 514 975 736 790 937
('rep1', 'GM130') 595 1123 865 709 600 1012 729 677 755
('rep2', 'GM130') 555 1021 760 682 483 999 632 904 655
('rep1', 'KIF5A') 529 1006 694 494 442 963 409 618 676
('rep2', 'KIF5A') 585 1178 796 730 514 961 719 779 904
('rep1', 'LAMP1') 488 913 616 407 487 716 425 785 737
('rep2', 'LAMP1') 531 823 676 398 403 973 411 891 761
('rep1', 'NCL') 538 852 773 483 311 881 557 700 660
('rep2', 'NCL') 564 1093 728 626 324 1006 629 776 768
('rep1', 'NEMO') 596 1000 777 716 537 1055 713 822 696
('rep2', 'NEMO') 617 993 736 567 470 793 780 867 468
('rep1', 'NONO') 613 1051 850 620 509 970 780 911 823
('rep2', 'NONO') 711 1061 821 801 626 930 688 911 853
('rep1', 'PEX14') 478 773 752 467 415 781 647 930 907
('rep2', 'PEX14') 566 792 795 667 528 988 701 858 983
('rep1', 'PML') 456 584 574 447 285 419 647 930 907
('rep2', 'PML') 480 732 582 667 412 552 701 858 983
('rep1', 'PSD95') 445 875 634 389 401 817 450 648 587
('rep2', 'PSD95') 490 986 639 569 410 778 610 773 751
('rep1', 'PURA') 535 1039 685 498 430 1007 408 635 676
('rep2', 'PURA') 585 1180 801 754 514 968 730 790 937
('rep1', 'Phalloidin') 576 1259 884 656 615 1257 683 915 763
('rep2', 'Phalloidin') 483 900 781 595 411 961 612 796 663
('rep1', 'SCNA') 470 976 504 552 269 769 524 944 734
('rep2', 'SCNA') 405 884 640 549 389 834 646 750 854
('rep1', 'SQSTM1') 592 1248 884 651 608 1243 650 915 763
('rep2', 'SQSTM1') 459 886 774 586 410 954 585 796 647
('rep1', 'TDP43') 613 1051 850 620 509 970 780 911 838
('rep2', 'TDP43') 711 1061 821 801 626 930 688 911 853
('rep1', 'TOMM20') 595 1123 865 709 603 1012 729 680 755
('rep2', 'TOMM20') 555 1021 764 682 490 999 632 904 655
('rep1', 'mitotracker') 478 773 752 467 415 781 622 922 907
('rep2', 'mitotracker') 566 792 795 667 531 988 701 858 983
1200
1000
800
600
400
200
Total number of whole cells

Batch 2.3

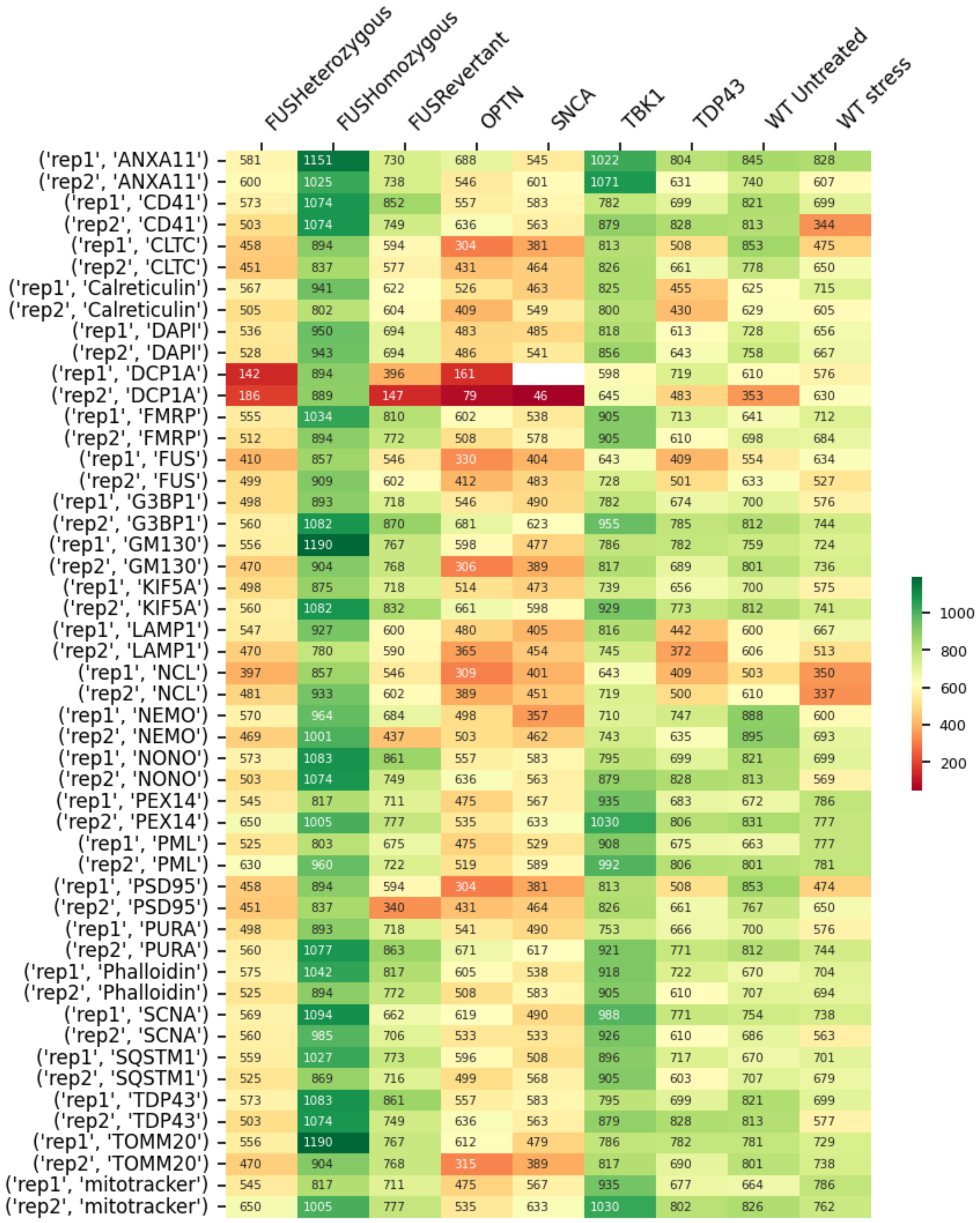


Total number of whole cells

#### Batch 2.4

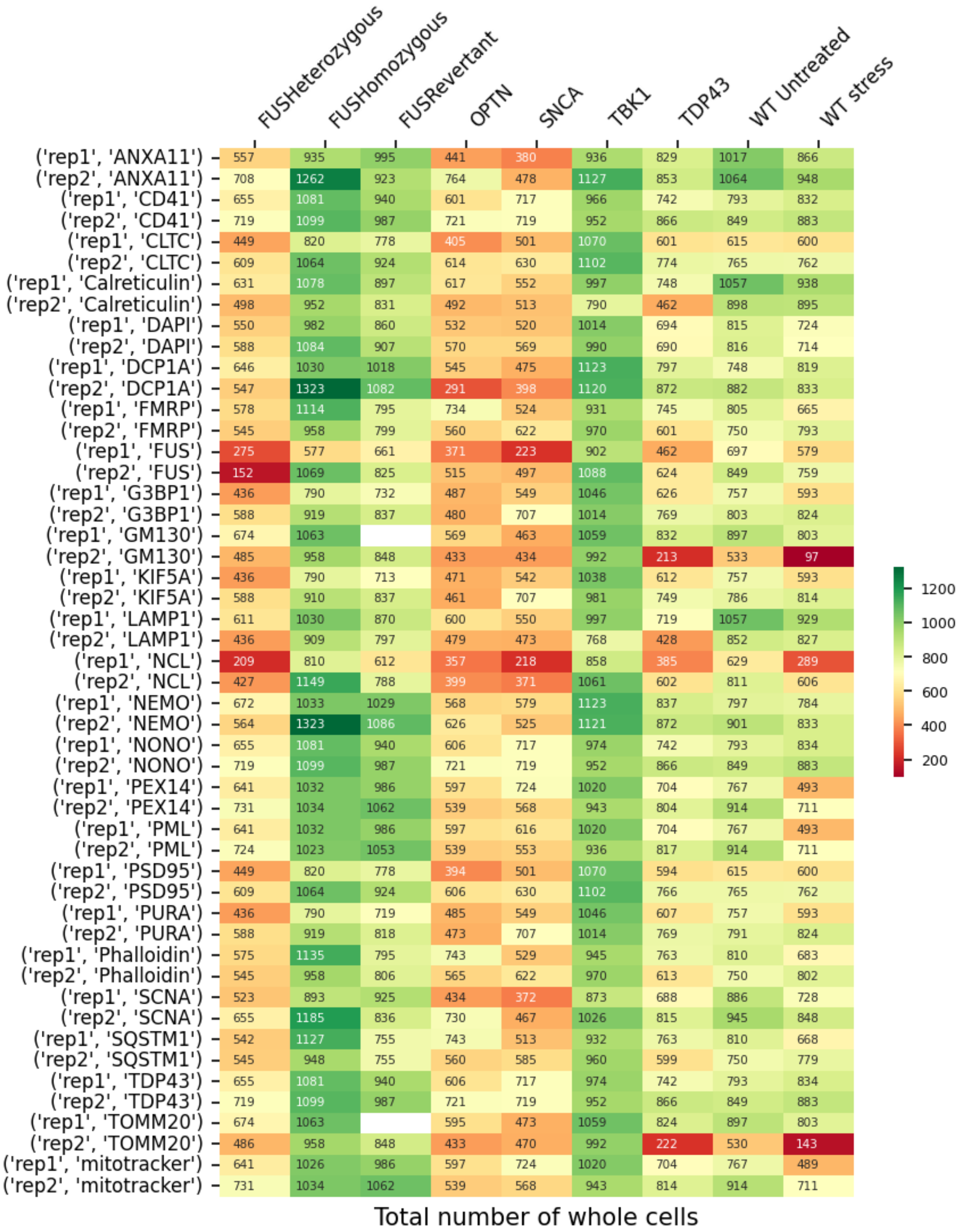
FUSHeterozygous FUSHomozygous FUSRevertant OPTN SNCA TBK1 TDP43 WT Untreated WT stress
('rep1', 'ANXA11') 557 935 995 441 380 936 829 1017 866
('rep2', 'ANXA11') 708 1262 923 764 478 1127 853 1064 948
('rep1', 'CD41') 655 1081 940 601 717 966 742 793 832
('rep2', 'CD41') 719 1099 987 721 719 952 866 849 883
('rep1', 'CLTC') 449 820 778 405 501 1070 601 615 600
('rep2', 'CLTC') 609 1064 924 614 630 1102 774 765 762
('rep1', 'Calreticulin') 631 1078 897 617 552 997 748 1057 938
('rep2', 'Calreticulin') 498 952 831 492 513 790 462 898 895
('rep1', 'DAPI') 550 982 860 532 520 1014 694 815 724
('rep2', 'DAPI') 588 1084 907 570 569 990 690 816 714
('rep1', 'DCP1A') 646 1030 1018 545 475 1123 797 748 819
('rep2', 'DCP1A') 547 1323 1082 291 398 1120 872 882 833
('rep1', 'FMRP') 578 1114 795 734 524 931 745 805 665
('rep2', 'FMRP') 545 958 799 560 622 970 601 750 793
('rep1', 'FUS') 275 577 661 371 223 902 462 697 579
('rep2', 'FUS') 152 1069 825 515 497 1088 624 849 759
('rep1', 'G3BP1') 436 790 732 487 549 1046 626 757 593
('rep2', 'G3BP1') 588 919 837 480 707 1014 769 803 824
('rep1', 'GM130') 674 1063 569 463 1059 832 897 803
('rep2', 'GM130') 485 958 848 433 434 992 213 533 97
('rep1', 'KIF5A') 436 790 713 471 542 1038 612 757 593
('rep2', 'KIF5A') 588 910 837 461 707 981 749 786 814
('rep1', 'LAMP1') 611 1030 870 600 550 997 719 1057 929
('rep2', 'LAMP1') 436 909 797 479 473 768 428 852 827
('rep1', 'NCL') 209 810 612 357 218 858 385 629 289
('rep2', 'NCL') 427 1149 788 399 371 1061 602 811 606
('rep1', 'NEMO') 672 1033 1029 568 579 1123 837 797 784
('rep2', 'NEMO') 564 1323 1086 626 525 1121 872 901 833
('rep1', 'NONO') 655 1081 940 606 717 974 742 793 834
('rep2', 'NONO') 719 1099 987 721 719 952 866 849 883
('rep1', 'PEX14') 641 1032 986 597 724 1020 704 767 493
('rep2', 'PEX14') 731 1034 1062 539 568 943 804 914 711
('rep1', 'PML') 641 1032 986 597 616 1020 704 767 493
('rep2', 'PML') 724 1023 1053 539 553 936 817 914 711
('rep1', 'PSD95') 449 820 778 394 501 1070 594 615 600
('rep2', 'PSD95') 609 1064 924 606 630 1102 766 765 762
('rep1', 'PURA') 436 790 719 485 549 1046 607 757 593
('rep2', 'PURA') 588 919 818 473 707 1014 769 791 824
('rep1', 'Phalloidin') 575 1135 795 743 529 945 763 810 683
('rep2', 'Phalloidin') 545 958 806 565 622 970 613 750 802
('rep1', 'SCNA') 523 893 925 434 372 873 688 886 728
('rep2', 'SCNA') 655 1185 836 730 467 1026 815 945 848
('rep1', 'SQSTM1') 542 1127 755 743 513 932 763 810 668
('rep2', 'SQSTM1') 545 948 755 560 585 960 599 750 779
('rep1', 'TDP43') 655 1081 940 606 717 974 742 793 834
('rep2', 'TDP43') 719 1099 987 721 719 952 866 849 883
('rep1', 'TOMM20') 674 1063 595 473 1059 824 897 803
('rep2', 'TOMM20') 486 958 848 433 470 992 222 530 143
('rep1', 'mitotracker') 641 1026 986 597 724 1020 704 767 489
('rep2', 'mitotracker') 731 1034 1062 539 568 943 814 914 711
1200
1000
800
600
400
200
Total number of whole cells
