## Supplementary material for "Organellomics: AI-driven deep organellar phenotyping reveals novel ALS mechanisms in human neurons": sALS patient-derived motor neurons (day 60) (4 organelles) - QC Report.pdf

### Coyne Lab QC Report

#### Processed Files Validation

Number of sites that survived pre-processing

Batch 1

| | Rep | Controls | sALS Negative cytoTDP43 | sALS Positive cytoTDP43 | c9orf72 ALS patients |
|---|---|---|---|---|---|
| DCP1A | rep1 | 10 | 10 | 10 | 10 |
| DCP1A | rep2 | 10 | 10 | 10 | 10 |
| DCP1A | rep3 | 10 | 0 | 10 | 10 |
| DCP1A | rep4 | 10 | 0 | 10 | 0 |
| DCP1A | rep5 | 10 | 0 | 10 | 0 |
| DCP1A | rep6 | 10 | 0 | 10 | 0 |
| DCP1A | rep7 | 0 | 0 | 10 | 0 |
| DCP1A | rep8 | 0 | 0 | 10 | 0 |
| DCP1A | rep9 | 0 | 0 | 10 | 0 |
| DCP1A | rep10 | 0 | 0 | 10 | 0 |
| Map2 | rep1 | 10 | 10 | 10 | 10 |
| Map2 | rep2 | 10 | 10 | 10 | 10 |
| Map2 | rep3 | 10 | 0 | 10 | 10 |
| Map2 | rep4 | 10 | 0 | 10 | 0 |
| Map2 | rep5 | 10 | 0 | 10 | 0 |
| Map2 | rep6 | 10 | 0 | 10 | 0 |
| Map2 | rep7 | 0 | 0 | 10 | 0 |
| Map2 | rep8 | 0 | 0 | 10 | 0 |
| Map2 | rep9 | 0 | 0 | 10 | 0 |
| Map2 | rep10 | 0 | 0 | 10 | 0 |
| TDP43 | rep1 | 10 | 10 | 10 | 10 |
| TDP43 | rep2 | 10 | 10 | 10 | 10 |
| TDP43 | rep3 | 10 | 0 | 10 | 10 |
| TDP43 | rep4 | 10 | 0 | 10 | 0 |
| TDP43 | rep5 | 10 | 0 | 10 | 0 |
| TDP43 | rep6 | 10 | 0 | 10 | 0 |
| TDP43 | rep7 | 0 | 0 | 10 | 0 |
| TDP43 | rep8 | 0 | 0 | 10 | 0 |
| TDP43 | rep9 | 0 | 0 | 10 | 0 |
| TDP43 | rep10 | 0 | 0 | 10 | 0 |
| DAPI | rep1 | 10 | 10 | 10 | 10 |
| DAPI | rep2 | 10 | 10 | 10 | 10 |
| DAPI | rep3 | 10 | 0 | 10 | 10 |
| DAPI | rep4 | 10 | 0 | 10 | 0 |
| DAPI | rep5 | 10 | 0 | 10 | 0 |
| DAPI | rep6 | 10 | 0 | 10 | 0 |
| DAPI | rep7 | 0 | 0 | 10 | 0 |
| DAPI | rep8 | 0 | 0 | 10 | 0 |
| DAPI | rep9 | 0 | 0 | 10 | 0 |
| DAPI | rep10 | 0 | 0 | 10 | 0 |

**Total Tile Count**

Batch1

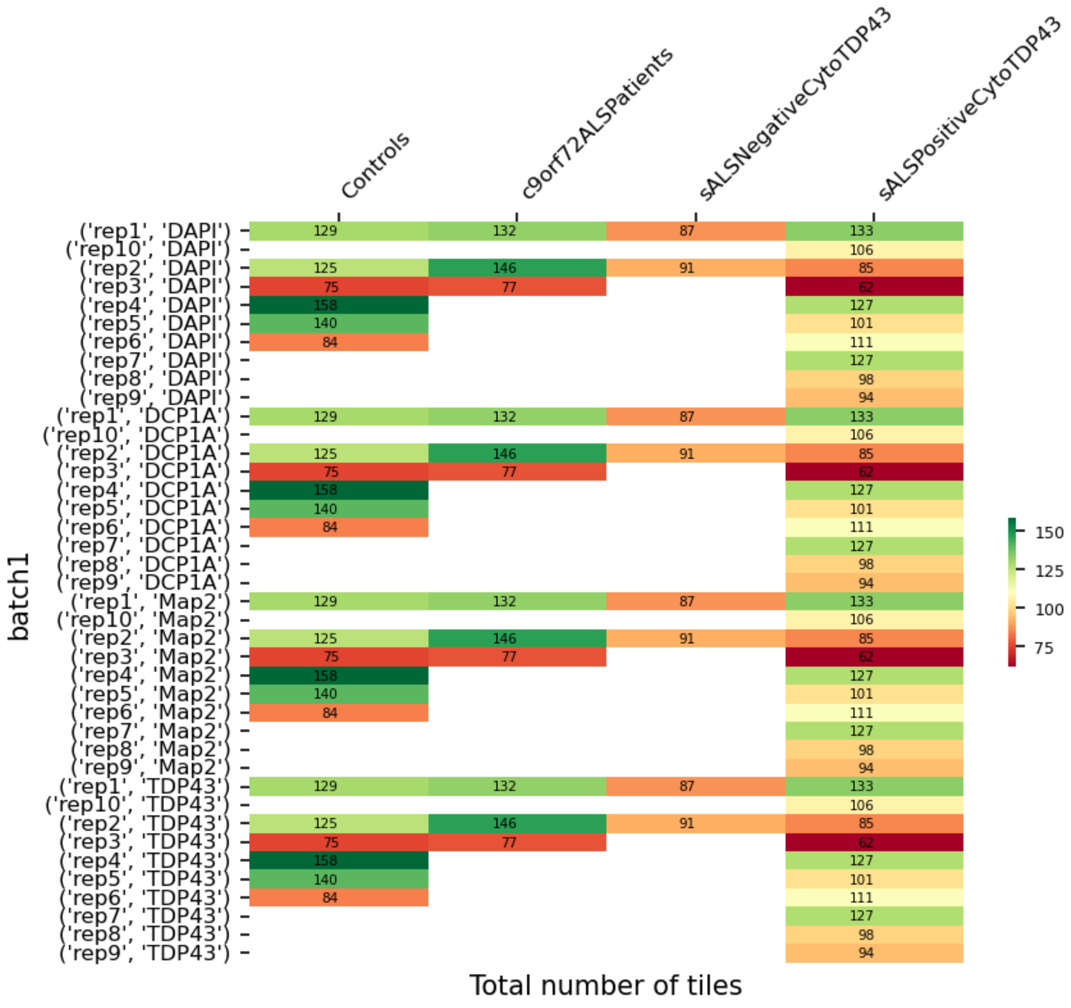

#### **Whole Cell Count**

Batch 1

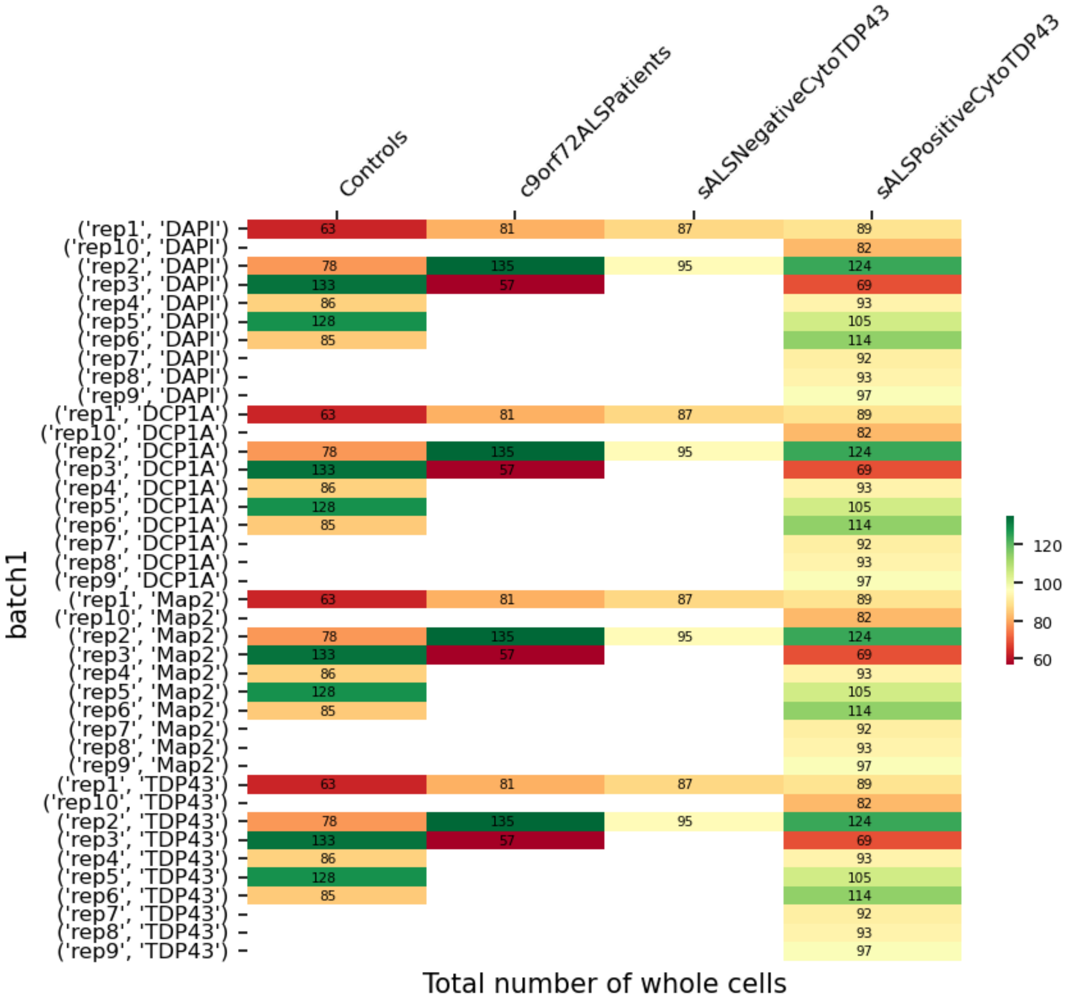
