## Supplementary material for "Organellomics: AI-driven deep organellar phenotyping reveals novel ALS mechanisms in human neurons": TDP-43deltaNLS iPSC-derived neurons - QC Report.pdf

### TDP-43 $^{deltaNLS}$ iPSC-derived neurons - QC Report

#### Processed Files Validation

Number of sites that survived pre-processing

Batch 1

| | Rep | dNLS DOX | dNLS Untreated |
|---|---|---|---|
| G3BP1 | rep1 | 162 | 104 |
| G3BP1 | rep2 | 8 | 74 |
| G3BP1 | rep3 | 77 | 0 |
| NONO | rep1 | 211 | 209 |
| NONO | rep2 | 229 | 121 |
| NONO | rep3 | 212 | 115 |
| SQSTM1 | rep1 | 191 | 172 |
| SQSTM1 | rep2 | 246 | 114 |
| SQSTM1 | rep3 | 227 | 181 |
| PSD95 | rep1 | 61 | 99 |
| PSD95 | rep2 | 147 | 143 |
| PSD95 | rep3 | 45 | 84 |
| NEMO | rep1 | 37 | 6 |
| NEMO | rep2 | 23 | 9 |
| NEMO | rep3 | 46 | 5 |
| GM130 | rep1 | 214 | 65 |
| GM130 | rep2 | 102 | 66 |
| GM130 | rep3 | 167 | 28 |
| NCL | rep1 | 182 | 148 |
| NCL | rep2 | 231 | 176 |
| NCL | rep3 | 219 | 115 |
| LSM14A | rep1 | 94 | 22 |
| LSM14A | rep2 | 91 | 40 |
| LSM14A | rep3 | 86 | 22 |
| TDP43 | rep1 | 28 | 7 |
| TDP43 | rep2 | 21 | 138 |
| TDP43 | rep3 | 150 | 0 |
| ANXA11 | rep1 | 246 | 196 |
| ANXA11 | rep2 | 156 | 246 |
| ANXA11 | rep3 | 225 | 210 |
| PEX14 | rep1 | 124 | 157 |
| PEX14 | rep2 | 129 | 163 |
| PEX14 | rep3 | 110 | 145 |
| mitotracker | rep1 | 91 | 113 |
| mitotracker | rep2 | 104 | 50 |
| mitotracker | rep3 | 45 | 8 |
| FMRP | rep1 | 158 | 104 |
| FMRP | rep2 | 8 | 69 |
| FMRP | rep3 | 74 | 0 |
| SON | rep1 | 211 | 209 |
| SON | rep2 | 229 | 121 |
| SON | rep3 | 212 | 115 |
| KIF5A | rep1 | 193 | 173 |
| KIF5A | rep2 | 245 | 117 |
| KIF5A | rep3 | 228 | 185 |
| CLTC | rep1 | 61 | 99 |
| CLTC | rep2 | 147 | 143 |
| CLTC | rep3 | 45 | 84 |
| DCP1A | rep1 | 37 | 6 |
| DCP1A | rep2 | 23 | 9 |
| DCP1A | rep3 | 46 | 5 |
| Calreticulin | rep1 | 214 | 64 |
| Calreticulin | rep2 | 102 | 65 |
| Calreticulin | rep3 | 169 | 28 |
| FUS | rep1 | 183 | 148 |
| FUS | rep2 | 231 | 176 |
| FUS | rep3 | 219 | 115 |
| HNRNPA1 | rep1 | 94 | 22 |
| HNRNPA1 | rep2 | 91 | 40 |
| HNRNPA1 | rep3 | 86 | 22 |
| PML | rep1 | 28 | 7 |
| PML | rep2 | 21 | 138 |
| PML | rep3 | 150 | 0 |
| LAMP1 | rep1 | 183 | 178 |
| LAMP1 | rep2 | 128 | 210 |
| LAMP1 | rep3 | 203 | 171 |
| SNCA | rep1 | 112 | 143 |
| SNCA | rep2 | 99 | 144 |
| SNCA | rep3 | 99 | 116 |
| TIA1 | rep1 | 91 | 113 |
| TIA1 | rep2 | 105 | 51 |
| TIA1 | rep3 | 46 | 8 |
| PURA | rep1 | 160 | 104 |
| PURA | rep2 | 8 | 74 |
| PURA | rep3 | 77 | 0 |
| Tubulin | rep1 | 192 | 175 |
| Tubulin | rep2 | 246 | 118 |
| Tubulin | rep3 | 229 | 184 |
| Phalloidin | rep1 | 61 | 99 |
| Phalloidin | rep2 | 147 | 143 |
| Phalloidin | rep3 | 45 | 84 |
| TOMM20 | rep1 | 91 | 113 |
| TOMM20 | rep2 | 104 | 50 |
| TOMM20 | rep3 | 45 | 8 |
| DAPI | rep1 | 1571 | 1277 |
| DAPI | rep2 | 1442 | 1292 |
| DAPI | rep3 | 1584 | 861 |

### <u>Batch 2</u>

| | Rep | dNLS DOX | dNLS Untreated |
|---|---|---|---|
| G3BP1 | rep1 | 126 | 91 |
| G3BP1 | rep2 | 3 | 186 |
| G3BP1 | rep3 | 136 | 9 |
| NONO | rep1 | 249 | 181 |
| NONO | rep2 | 205 | 174 |
| NONO | rep3 | 188 | 207 |
| SQSTM1 | rep1 | 191 | 183 |
| SQSTM1 | rep2 | 239 | 127 |
| SQSTM1 | rep3 | 239 | 131 |
| PSD95 | rep1 | 114 | 75 |
| PSD95 | rep2 | 133 | 130 |
| PSD95 | rep3 | 234 | 19 |
| NEMO | rep1 | 47 | 55 |
| NEMO | rep2 | 93 | 131 |
| NEMO | rep3 | 54 | 17 |
| GM130 | rep1 | 187 | 85 |
| GM130 | rep2 | 99 | 134 |
| GM130 | rep3 | 175 | 22 |
| NCL | rep1 | 229 | 116 |
| NCL | rep2 | 164 | 140 |
| NCL | rep3 | 186 | 123 |
| LSM14A | rep1 | 70 | 92 |
| LSM14A | rep2 | 60 | 111 |
| LSM14A | rep3 | 125 | 37 |
| TDP43 | rep1 | 140 | 9 |
| TDP43 | rep2 | 34 | 27 |
| TDP43 | rep3 | 154 | 2 |
| ANXA11 | rep1 | 95 | 46 |
| ANXA11 | rep2 | 124 | 187 |
| ANXA11 | rep3 | 112 | 27 |
| PEX14 | rep1 | 171 | 128 |
| PEX14 | rep2 | 72 | 172 |
| PEX14 | rep3 | 42 | 75 |
| mitotracker | rep1 | 13 | 177 |
| mitotracker | rep2 | 123 | 4 |
| mitotracker | rep3 | 85 | 25 |
| FMRP | rep1 | 126 | 90 |
| FMRP | rep2 | 3 | 184 |
| FMRP | rep3 | 135 | 9 |
| SON | rep1 | 249 | 182 |
| SON | rep2 | 205 | 174 |
| SON | rep3 | 188 | 207 |
| KIF5A | rep1 | 190 | 185 |
| KIF5A | rep2 | 241 | 128 |
| KIF5A | rep3 | 245 | 133 |
| CLTC | rep1 | 114 | 75 |
| CLTC | rep2 | 133 | 130 |
| CLTC | rep3 | 234 | 19 |
| DCP1A | rep1 | 47 | 55 |
| DCP1A | rep2 | 93 | 131 |
| DCP1A | rep3 | 55 | 17 |
| Calreticulin | rep1 | 188 | 86 |
| Calreticulin | rep2 | 101 | 136 |
| Calreticulin | rep3 | 176 | 22 |
| FUS | rep1 | 229 | 116 |
| FUS | rep2 | 164 | 140 |
| FUS | rep3 | 186 | 123 |
| HNRNPA1 | rep1 | 70 | 93 |
| HNRNPA1 | rep2 | 60 | 112 |
| HNRNPA1 | rep3 | 125 | 37 |
| PML | rep1 | 140 | 9 |
| PML | rep2 | 36 | 27 |
| PML | rep3 | 154 | 2 |
| LAMP1 | rep1 | 90 | 45 |
| LAMP1 | rep2 | 120 | 179 |
| LAMP1 | rep3 | 95 | 26 |
| SNCA | rep1 | 133 | 106 |
| SNCA | rep2 | 62 | 138 |
| SNCA | rep3 | 40 | 65 |
| TIA1 | rep1 | 14 | 179 |
| TIA1 | rep2 | 126 | 4 |
| TIA1 | rep3 | 86 | 25 |
| PURA | rep1 | 126 | 91 |
| PURA | rep2 | 3 | 186 |
| PURA | rep3 | 134 | 9 |
| Tubulin | rep1 | 193 | 185 |
| Tubulin | rep2 | 240 | 130 |
| Tubulin | rep3 | 245 | 132 |
| Phalloidin | rep1 | 114 | 75 |
| Phalloidin | rep2 | 133 | 130 |
| Phalloidin | rep3 | 234 | 19 |
| TOMM20 | rep1 | 13 | 177 |
| TOMM20 | rep2 | 123 | 4 |
| TOMM20 | rep3 | 85 | 25 |
| DAPI | rep1 | 1618 | 1229 |
| DAPI | rep2 | 1342 | 1500 |
| DAPI | rep3 | 1720 | 690 |

### Batch 3

| | Rep | dNLS DOX | dNLS Untreated |
|---|---|---|---|
| G3BP1 | rep1 | 221 | 90 |
| G3BP1 | rep2 | 205 | 171 |
| G3BP1 | rep3 | 222 | 147 |
| NONO | rep1 | 246 | 181 |
| NONO | rep2 | 246 | 196 |
| NONO | rep3 | 242 | 217 |
| SQSTM1 | rep1 | 238 | 184 |
| SQSTM1 | rep2 | 235 | 101 |
| SQSTM1 | rep3 | 240 | 229 |
| PSD95 | rep1 | 136 | 170 |
| PSD95 | rep2 | 239 | 218 |
| PSD95 | rep3 | 240 | 190 |
| NEMO | rep1 | 105 | 183 |
| NEMO | rep2 | 217 | 204 |
| NEMO | rep3 | 120 | 55 |
| GM130 | rep1 | 247 | 238 |
| GM130 | rep2 | 237 | 214 |
| GM130 | rep3 | 237 | 215 |
| NCL | rep1 | 249 | 222 |
| NCL | rep2 | 243 | 233 |
| NCL | rep3 | 249 | 214 |
| LSM14A | rep1 | 166 | 143 |
| LSM14A | rep2 | 106 | 103 |
| LSM14A | rep3 | 158 | 107 |
| TDP43 | rep1 | 250 | 134 |
| TDP43 | rep2 | 212 | 241 |
| TDP43 | rep3 | 246 | 30 |
| ANXA11 | rep1 | 247 | 236 |
| ANXA11 | rep2 | 227 | 242 |
| ANXA11 | rep3 | 238 | 205 |
| PEX14 | rep1 | 244 | 248 |
| PEX14 | rep2 | 242 | 239 |
| PEX14 | rep3 | 246 | 232 |
| mitotracker | rep1 | 218 | 221 |
| mitotracker | rep2 | 184 | 198 |
| mitotracker | rep3 | 83 | 235 |
| FMRP | rep1 | 219 | 90 |
| FMRP | rep2 | 194 | 166 |
| FMRP | rep3 | 215 | 144 |
| SON | rep1 | 249 | 181 |
| SON | rep2 | 250 | 196 |
| SON | rep3 | 246 | 217 |
| KIF5A | rep1 | 241 | 186 |
| KIF5A | rep2 | 241 | 101 |
| KIF5A | rep3 | 247 | 232 |
| CLTC | rep1 | 136 | 170 |
| CLTC | rep2 | 239 | 218 |
| CLTC | rep3 | 240 | 190 |
| DCP1A | rep1 | 105 | 183 |
| DCP1A | rep2 | 219 | 204 |
| DCP1A | rep3 | 121 | 55 |
| Calreticulin | rep1 | 247 | 238 |
| Calreticulin | rep2 | 237 | 215 |
| Calreticulin | rep3 | 237 | 215 |
| FUS | rep1 | 249 | 222 |
| FUS | rep2 | 243 | 233 |
| FUS | rep3 | 249 | 214 |
| HNRNPA1 | rep1 | 166 | 143 |
| HNRNPA1 | rep2 | 106 | 103 |
| HNRNPA1 | rep3 | 158 | 107 |
| PML | rep1 | 250 | 134 |
| PML | rep2 | 215 | 241 |
| PML | rep3 | 246 | 30 |
| LAMP1 | rep1 | 216 | 189 |
| LAMP1 | rep2 | 187 | 211 |
| LAMP1 | rep3 | 217 | 187 |
| SNCA | rep1 | 212 | 218 |
| SNCA | rep2 | 189 | 214 |
| SNCA | rep3 | 186 | 185 |
| TIA1 | rep1 | 221 | 229 |
| TIA1 | rep2 | 188 | 205 |
| TIA1 | rep3 | 85 | 241 |
| PURA | rep1 | 221 | 90 |
| PURA | rep2 | 205 | 171 |
| PURA | rep3 | 220 | 147 |
| Tubulin | rep1 | 239 | 185 |
| Tubulin | rep2 | 239 | 101 |
| Tubulin | rep3 | 244 | 231 |
| Phalloidin | rep1 | 136 | 171 |
| Phalloidin | rep2 | 239 | 218 |
| Phalloidin | rep3 | 240 | 190 |
| TOMM20 | rep1 | 218 | 221 |
| TOMM20 | rep2 | 184 | 198 |
| TOMM20 | rep3 | 83 | 235 |
| DAPI | rep1 | 2530 | 2193 |
| DAPI | rep2 | 2537 | 2320 |
| DAPI | rep3 | 2483 | 2040 |

### Batch 4

| | Rep | dNLS DOX | dNLS Untreated |
|---|---|---|---|
| G3BP1 | rep1 | 228 | 228 |
| G3BP1 | rep2 | 220 | 222 |
| G3BP1 | rep3 | 195 | 158 |
| NONO | rep1 | 250 | 247 |
| NONO | rep2 | 249 | 249 |
| NONO | rep3 | 242 | 247 |
| SQSTM1 | rep1 | 237 | 235 |
| SQSTM1 | rep2 | 236 | 243 |
| SQSTM1 | rep3 | 238 | 237 |
| PSD95 | rep1 | 190 | 248 |
| PSD95 | rep2 | 250 | 250 |
| PSD95 | rep3 | 250 | 182 |
| NEMO | rep1 | 112 | 250 |
| NEMO | rep2 | 248 | 238 |
| NEMO | rep3 | 112 | 240 |
| GM130 | rep1 | 250 | 247 |
| GM130 | rep2 | 249 | 250 |
| GM130 | rep3 | 250 | 250 |
| NCL | rep1 | 250 | 250 |
| NCL | rep2 | 250 | 247 |
| NCL | rep3 | 240 | 249 |
| LSM14A | rep1 | 226 | 250 |
| LSM14A | rep2 | 126 | 244 |
| LSM14A | rep3 | 221 | 169 |
| TDP43 | rep1 | 226 | 250 |
| TDP43 | rep2 | 243 | 230 |
| TDP43 | rep3 | 248 | 249 |
| ANXA11 | rep1 | 250 | 250 |
| ANXA11 | rep2 | 249 | 250 |
| ANXA11 | rep3 | 250 | 245 |
| PEX14 | rep1 | 244 | 241 |
| PEX14 | rep2 | 246 | 244 |
| PEX14 | rep3 | 246 | 244 |
| mitotracker | rep1 | 241 | 177 |
| mitotracker | rep2 | 167 | 243 |
| mitotracker | rep3 | 184 | 228 |
| FMRP | rep1 | 226 | 225 |
| FMRP | rep2 | 220 | 221 |
| FMRP | rep3 | 193 | 153 |
| SON | rep1 | 250 | 249 |
| SON | rep2 | 250 | 250 |
| SON | rep3 | 246 | 248 |
| KIF5A | rep1 | 239 | 241 |
| KIF5A | rep2 | 242 | 244 |
| KIF5A | rep3 | 244 | 244 |
| CLTC | rep1 | 190 | 248 |
| CLTC | rep2 | 250 | 250 |
| CLTC | rep3 | 250 | 182 |
| DCP1A | rep1 | 112 | 250 |
| DCP1A | rep2 | 249 | 240 |
| DCP1A | rep3 | 112 | 240 |
| Calreticulin | rep1 | 250 | 250 |
| Calreticulin | rep2 | 249 | 250 |
| Calreticulin | rep3 | 250 | 250 |
| FUS | rep1 | 250 | 250 |
| FUS | rep2 | 250 | 247 |
| FUS | rep3 | 240 | 249 |
| HNRNPA1 | rep1 | 226 | 250 |
| HNRNPA1 | rep2 | 126 | 244 |
| HNRNPA1 | rep3 | 221 | 169 |
| PML | rep1 | 227 | 250 |
| PML | rep2 | 243 | 230 |
| PML | rep3 | 248 | 249 |
| LAMP1 | rep1 | 223 | 210 |
| LAMP1 | rep2 | 221 | 236 |
| LAMP1 | rep3 | 223 | 227 |
| SNCA | rep1 | 214 | 187 |
| SNCA | rep2 | 206 | 203 |
| SNCA | rep3 | 205 | 207 |
| TIA1 | rep1 | 250 | 179 |
| TIA1 | rep2 | 169 | 248 |
| TIA1 | rep3 | 187 | 232 |
| PURA | rep1 | 227 | 228 |
| PURA | rep2 | 219 | 222 |
| PURA | rep3 | 195 | 158 |
| Tubulin | rep1 | 239 | 239 |
| Tubulin | rep2 | 238 | 243 |
| Tubulin | rep3 | 239 | 242 |
| Phalloidin | rep1 | 190 | 248 |
| Phalloidin | rep2 | 250 | 250 |
| Phalloidin | rep3 | 250 | 182 |
| TOMM20 | rep1 | 241 | 177 |
| TOMM20 | rep2 | 167 | 243 |
| TOMM20 | rep3 | 184 | 228 |
| DAPI | rep1 | 2678 | 2816 |
| DAPI | rep2 | 2702 | 2883 |
| DAPI | rep3 | 2645 | 2674 |

### Batch 5

| | Rep | dNLS DOX | dNLS Untreated |
|---|---|---|---|
| G3BP1 | rep1 | 112 | 40 |
| G3BP1 | rep2 | 13 | 124 |
| G3BP1 | rep3 | 112 | 12 |
| NONO | rep1 | 183 | 111 |
| NONO | rep2 | 240 | 140 |
| NONO | rep3 | 204 | 217 |
| SQSTM1 | rep1 | 247 | 150 |
| SQSTM1 | rep2 | 227 | 201 |
| SQSTM1 | rep3 | 245 | 208 |
| PSD95 | rep1 | 14 | 179 |
| PSD95 | rep2 | 142 | 20 |
| PSD95 | rep3 | 59 | 73 |
| NEMO | rep1 | 47 | 0 |
| NEMO | rep2 | 9 | 40 |
| NEMO | rep3 | 35 | 0 |
| GM130 | rep1 | 138 | 72 |
| GM130 | rep2 | 83 | 13 |
| GM130 | rep3 | 27 | 39 |
| NCL | rep1 | 231 | 125 |
| NCL | rep2 | 190 | 172 |
| NCL | rep3 | 194 | 30 |
| LSM14A | rep1 | 33 | 61 |
| LSM14A | rep2 | 55 | 12 |
| LSM14A | rep3 | 4 | 31 |
| TDP43 | rep1 | 59 | 3 |
| TDP43 | rep2 | 64 | 29 |
| TDP43 | rep3 | 109 | 1 |
| ANXA11 | rep1 | 115 | 202 |
| ANXA11 | rep2 | 157 | 19 |
| ANXA11 | rep3 | 125 | 12 |
| PEX14 | rep1 | 28 | 6 |
| PEX14 | rep2 | 27 | 16 |
| PEX14 | rep3 | 17 | 8 |
| mitotracker | rep1 | 13 | 103 |
| mitotracker | rep2 | 97 | 3 |
| mitotracker | rep3 | 57 | 8 |
| FMRP | rep1 | 112 | 39 |
| FMRP | rep2 | 13 | 123 |
| FMRP | rep3 | 108 | 12 |
| SON | rep1 | 185 | 112 |
| SON | rep2 | 246 | 140 |
| SON | rep3 | 205 | 220 |
| KIF5A | rep1 | 248 | 150 |
| KIF5A | rep2 | 229 | 201 |
| KIF5A | rep3 | 247 | 210 |
| CLTC | rep1 | 14 | 178 |
| CLTC | rep2 | 142 | 20 |
| CLTC | rep3 | 59 | 73 |
| DCP1A | rep1 | 47 | 0 |
| DCP1A | rep2 | 9 | 40 |
| DCP1A | rep3 | 35 | 0 |
| Calreticulin | rep1 | 138 | 72 |
| Calreticulin | rep2 | 84 | 13 |
| Calreticulin | rep3 | 27 | 39 |
| FUS | rep1 | 231 | 125 |
| FUS | rep2 | 190 | 172 |
| FUS | rep3 | 194 | 30 |
| HNRNPA1 | rep1 | 33 | 61 |
| HNRNPA1 | rep2 | 55 | 12 |
| HNRNPA1 | rep3 | 4 | 31 |
| PML | rep1 | 62 | 3 |
| PML | rep2 | 65 | 29 |
| PML | rep3 | 110 | 1 |
| LAMP1 | rep1 | 106 | 174 |
| LAMP1 | rep2 | 147 | 18 |
| LAMP1 | rep3 | 122 | 11 |
| SNCA | rep1 | 27 | 6 |
| SNCA | rep2 | 27 | 16 |
| SNCA | rep3 | 17 | 8 |
| TIA1 | rep1 | 13 | 106 |
| TIA1 | rep2 | 99 | 3 |
| TIA1 | rep3 | 57 | 8 |
| PURA | rep1 | 111 | 39 |
| PURA | rep2 | 13 | 123 |
| PURA | rep3 | 112 | 12 |
| Tubulin | rep1 | 247 | 150 |
| Tubulin | rep2 | 231 | 201 |
| Tubulin | rep3 | 248 | 208 |
| Phalloidin | rep1 | 14 | 179 |
| Phalloidin | rep2 | 142 | 20 |
| Phalloidin | rep3 | 59 | 73 |
| TOMM20 | rep1 | 13 | 103 |
| TOMM20 | rep2 | 97 | 3 |
| TOMM20 | rep3 | 57 | 8 |
| DAPI | rep1 | 1212 | 1025 |
| DAPI | rep2 | 1306 | 787 |
| DAPI | rep3 | 1190 | 641 |

### Total Tile Count

#### Batch1

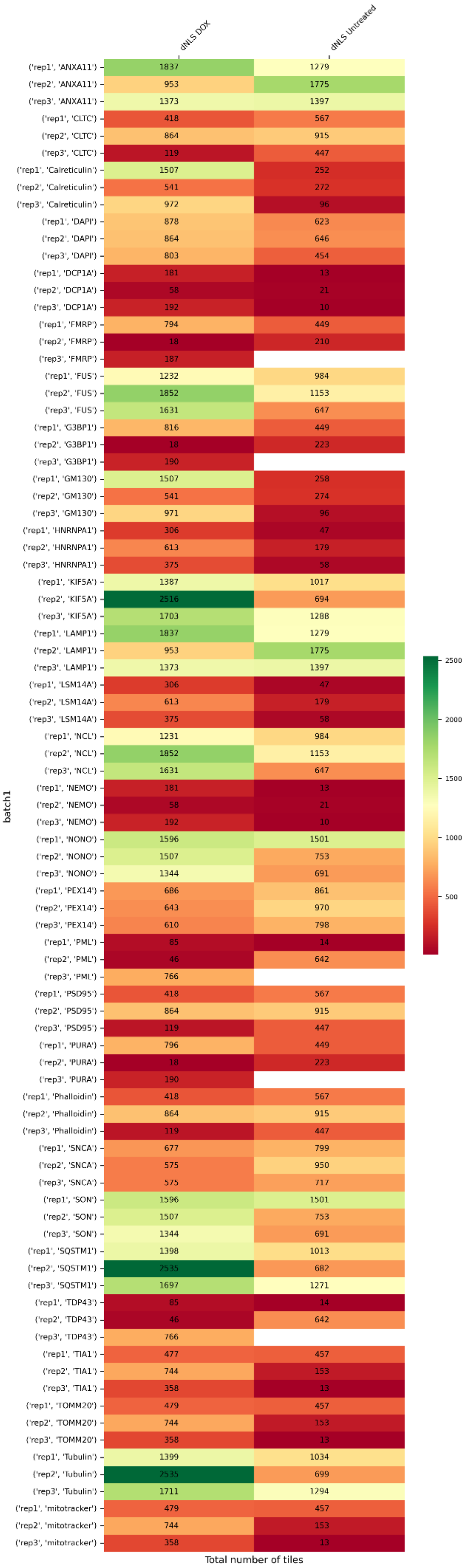

### Batch 2

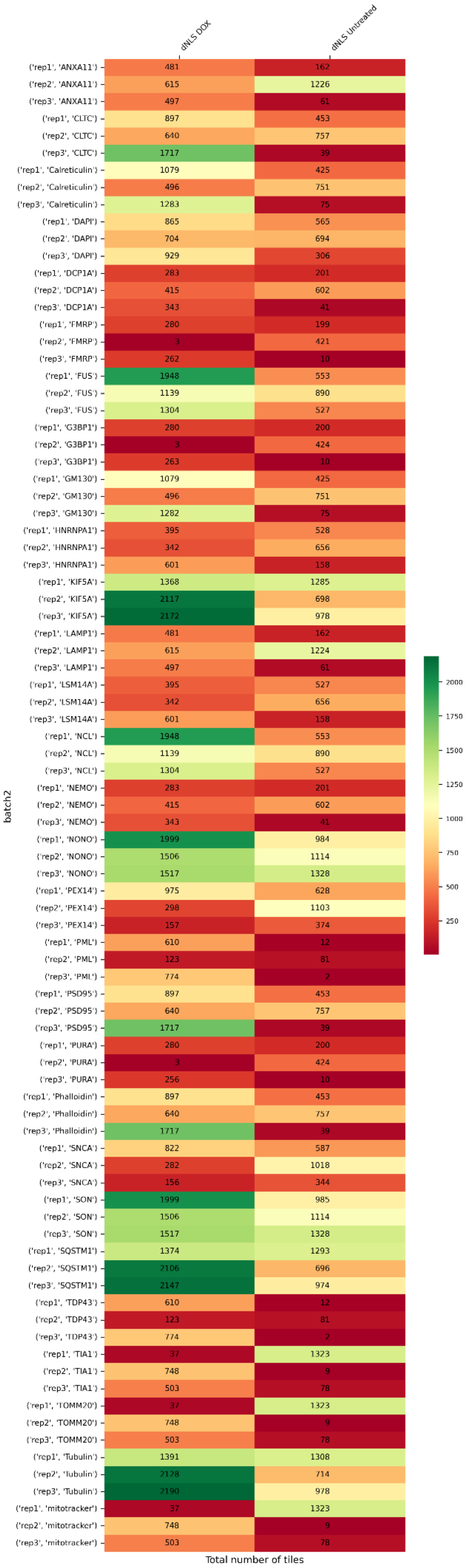

dNLS DOX
dNLS Untreated
('rep1', 'ANXA11') 481 162
('rep2', 'ANXA11') 615 1226
('rep3', 'ANXA11') 497 61
('rep1', 'CLTC') 897 453
('rep2', 'CLTC') 640 757
('rep3', 'CLTC') 1717 39
('rep1', 'Calreticulin') 1079 425
('rep2', 'Calreticulin') 496 751
('rep3', 'Calreticulin') 1283 75
('rep1', 'DAPI') 865 565
('rep2', 'DAPI') 704 694
('rep3', 'DAPI') 929 306
('rep1', 'DCP1A') 283 201
('rep2', 'DCP1A') 415 602
('rep3', 'DCP1A') 343 41
('rep1', 'FMRP') 280 199
('rep2', 'FMRP') 3 421
('rep3', 'FMRP') 262 10
('rep1', 'FUS') 1948 553
('rep2', 'FUS') 1139 890
('rep3', 'FUS') 1304 527
('rep1', 'G3BP1') 280 200
('rep2', 'G3BP1') 3 424
('rep3', 'G3BP1') 263 10
('rep1', 'GM130') 1079 425
('rep2', 'GM130') 496 751
('rep3', 'GM130') 1282 75
('rep1', 'HNRNPA1') 395 528
('rep2', 'HNRNPA1') 342 656
('rep3', 'HNRNPA1') 601 158
('rep1', 'KIF5A') 1368 1285
('rep2', 'KIF5A') 2117 698
('rep3', 'KIF5A') 2172 978
('rep1', 'LAMP1') 481 162
('rep2', 'LAMP1') 615 1224
('rep3', 'LAMP1') 497 61
('rep1', 'LSM14A') 395 527
('rep2', 'LSM14A') 342 656
('rep3', 'LSM14A') 601 158
('rep1', 'NCL') 1948 553
('rep2', 'NCL') 1139 890
('rep3', 'NCL') 1304 527
('rep1', 'NEMO') 283 201
('rep2', 'NEMO') 415 602
('rep3', 'NEMO') 343 41
('rep1', 'NONO') 1999 984
('rep2', 'NONO') 1506 1114
('rep3', 'NONO') 1517 1328
('rep1', 'PEX14') 975 628
('rep2', 'PEX14') 298 1103
('rep3', 'PEX14') 157 374
('rep1', 'PML') 610 12
('rep2', 'PML') 123 81
('rep3', 'PML') 774 7
('rep1', 'PSD95') 897 453
('rep2', 'PSD95') 640 757
('rep3', 'PSD95') 1717 39
('rep1', 'PURA') 280 200
('rep2', 'PURA') 3 424
('rep3', 'PURA') 256 10
('rep1', 'Phalloidin') 897 453
('rep2', 'Phalloidin') 640 757
('rep3', 'Phalloidin') 1717 39
('rep1', 'SNCA') 822 587
('rep2', 'SNCA') 282 1018
('rep3', 'SNCA') 156 344
('rep1', 'SON') 1999 985
('rep2', 'SON') 1506 1114
('rep3', 'SON') 1517 1328
('rep1', 'SQSTM1') 1374 1293
('rep2', 'SQSTM1') 2106 696
('rep3', 'SQSTM1') 2147 974
('rep1', 'TDP43') 610 12
('rep2', 'TDP43') 123 81
('rep3', 'TDP43') 774 7
('rep1', 'TIA1') 37 1323
('rep2', 'TIA1') 748 9
('rep3', 'TIA1') 503 78
('rep1', 'TOMM20') 37 1323
('rep2', 'TOMM20') 748 9
('rep3', 'TOMM20') 503 78
('rep1', 'Tubulin') 1391 1308
('rep2', 'Tubulin') 2128 714
('rep3', 'Tubulin') 2190 978
('rep1', 'mitotracker') 37 1323
('rep2', 'mitotracker') 748 9
('rep3', 'mitotracker') 503 78
batch2
Total number of tiles
2000
1750
1500
1250
1000
750
500
250

<u>Batch 3</u>

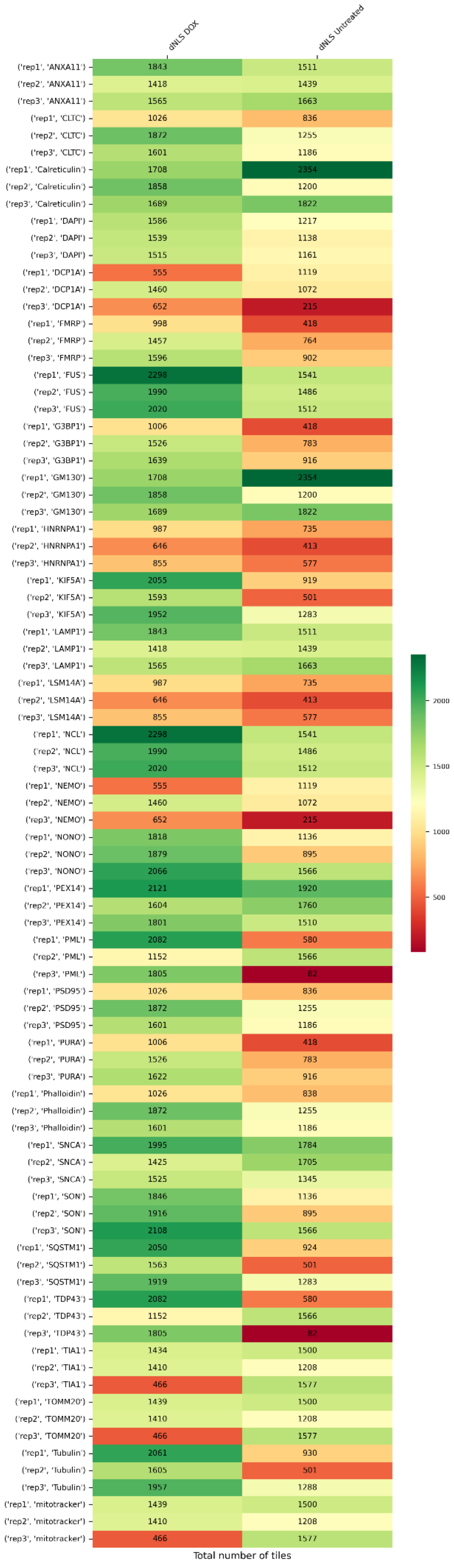
dNLS DOX
dNLS Untreated
('rep1', 'ANXA11') 1843 1511
('rep2', 'ANXA11') 1418 1439
('rep3', 'ANXA11') 1565 1663
('rep1', 'CLTC') 1026 836
('rep2', 'CLTC') 1872 1255
('rep3', 'CLTC') 1601 1186
('rep1', 'Calreticulin') 1708 2354
('rep2', 'Calreticulin') 1858 1200
('rep3', 'Calreticulin') 1689 1822
('rep1', 'DAPI') 1586 1217
('rep2', 'DAPI') 1539 1138
('rep3', 'DAPI') 1515 1161
('rep1', 'DCP1A') 555 1119
('rep2', 'DCP1A') 1460 1072
('rep3', 'DCP1A') 652 215
('rep1', 'FMRP') 998 418
('rep2', 'FMRP') 1457 764
('rep3', 'FMRP') 1596 902
('rep1', 'FUS') 2298 1541
('rep2', 'FUS') 1990 1486
('rep3', 'FUS') 2020 1512
('rep1', 'G3BP1') 1006 418
('rep2', 'G3BP1') 1526 783
('rep3', 'G3BP1') 1639 916
('rep1', 'GM130') 1708 2354
('rep2', 'GM130') 1858 1200
('rep3', 'GM130') 1689 1822
('rep1', 'HNRNPA1') 987 735
('rep2', 'HNRNPA1') 646 413
('rep3', 'HNRNPA1') 855 577
('rep1', 'KIF5A') 2055 919
('rep2', 'KIF5A') 1593 501
('rep3', 'KIF5A') 1952 1283
('rep1', 'LAMP1') 1843 1511
('rep2', 'LAMP1') 1418 1439
('rep3', 'LAMP1') 1565 1663
('rep1', 'LSM14A') 987 735
('rep2', 'LSM14A') 646 413
('rep3', 'LSM14A') 855 577
('rep1', 'NCL') 2298 1541
('rep2', 'NCL') 1990 1486
('rep3', 'NCL') 2020 1512
('rep1', 'NEMO') 555 1119
('rep2', 'NEMO') 1460 1072
('rep3', 'NEMO') 652 215
('rep1', 'NONO') 1818 1136
('rep2', 'NONO') 1879 895
('rep3', 'NONO') 2066 1566
('rep1', 'PEX14') 2121 1920
('rep2', 'PEX14') 1604 1760
('rep3', 'PEX14') 1801 1510
('rep1', 'PML') 2082 580
('rep2', 'PML') 1152 1566
('rep3', 'PML') 1805 82
('rep1', 'PSD95') 1026 836
('rep2', 'PSD95') 1872 1255
('rep3', 'PSD95') 1601 1186
('rep1', 'PURA') 1006 418
('rep2', 'PURA') 1526 783
('rep3', 'PURA') 1622 916
('rep1', 'Phalloidin') 1026 838
('rep2', 'Phalloidin') 1872 1255
('rep3', 'Phalloidin') 1601 1186
('rep1', 'SNCA') 1995 1784
('rep2', 'SNCA') 1425 1705
('rep3', 'SNCA') 1525 1345
('rep1', 'SON') 1846 1136
('rep2', 'SON') 1916 895
('rep3', 'SON') 2108 1566
('rep1', 'SQSTM1') 2050 924
('rep2', 'SQSTM1') 1563 501
('rep3', 'SQSTM1') 1919 1283
('rep1', 'TDP43') 2082 580
('rep2', 'TDP43') 1152 1566
('rep3', 'TDP43') 1805 82
('rep1', 'TIA1') 1434 1500
('rep2', 'TIA1') 1410 1208
('rep3', 'TIA1') 466 1577
('rep1', 'TOMM20') 1439 1500
('rep2', 'TOMM20') 1410 1208
('rep3', 'TOMM20') 466 1577
('rep1', 'Tubulin') 2061 930
('rep2', 'Tubulin') 1605 501
('rep3', 'Tubulin') 1957 1288
('rep1', 'mitotracker') 1439 1500
('rep2', 'mitotracker') 1410 1208
('rep3', 'mitotracker') 466 1577
2000
1500
1000
500
Total number of tiles

### Batch 4

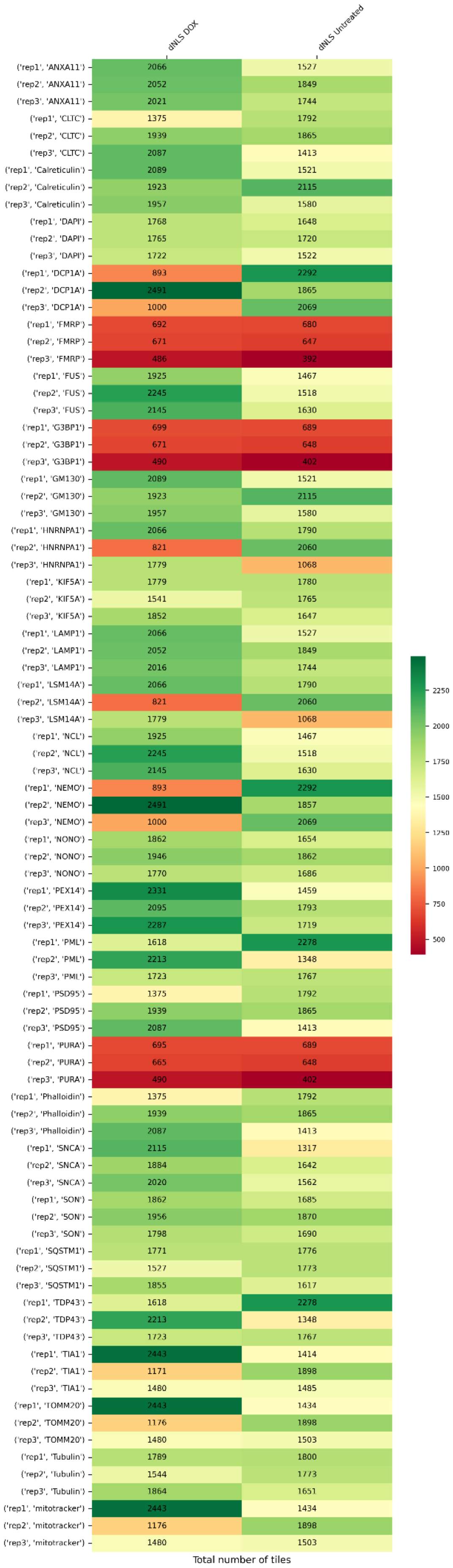
dNLS DOX
dNLS Untreated
('rep1', 'ANXA11') 2066 1527
('rep2', 'ANXA11') 2052 1849
('rep3', 'ANXA11') 2021 1744
('rep1', 'CLTC') 1375 1792
('rep2', 'CLTC') 1939 1865
('rep3', 'CLTC') 2087 1413
('rep1', 'Calreticulin') 2089 1521
('rep2', 'Calreticulin') 1923 2115
('rep3', 'Calreticulin') 1957 1580
('rep1', 'DAPI') 1768 1648
('rep2', 'DAPI') 1765 1720
('rep3', 'DAPI') 1722 1522
('rep1', 'DCP1A') 893 2292
('rep2', 'DCP1A') 2491 1865
('rep3', 'DCP1A') 1000 2069
('rep1', 'FMRP') 692 680
('rep2', 'FMRP') 671 647
('rep3', 'FMRP') 486 392
('rep1', 'FUS') 1925 1467
('rep2', 'FUS') 2245 1518
('rep3', 'FUS') 2145 1630
('rep1', 'G3BP1') 699 689
('rep2', 'G3BP1') 671 648
('rep3', 'G3BP1') 490 402
('rep1', 'GM130') 2089 1521
('rep2', 'GM130') 1923 2115
('rep3', 'GM130') 1957 1580
('rep1', 'HNRNPA1') 2066 1790
('rep2', 'HNRNPA1') 821 2060
('rep3', 'HNRNPA1') 1779 1068
('rep1', 'KIF5A') 1779 1780
('rep2', 'KIF5A') 1541 1765
('rep3', 'KIF5A') 1852 1647
('rep1', 'LAMP1') 2066 1527
('rep2', 'LAMP1') 2052 1849
('rep3', 'LAMP1') 2016 1744
('rep1', 'LSM14A') 2066 1790
('rep2', 'LSM14A') 821 2060
('rep3', 'LSM14A') 1779 1068
('rep1', 'NCL') 1925 1467
('rep2', 'NCL') 2245 1518
('rep3', 'NCL') 2145 1630
('rep1', 'NEMO') 893 2292
('rep2', 'NEMO') 2491 1857
('rep3', 'NEMO') 1000 2069
('rep1', 'NONO') 1862 1654
('rep2', 'NONO') 1946 1862
('rep3', 'NONO') 1770 1686
('rep1', 'PEX14') 2331 1459
('rep2', 'PEX14') 2095 1793
('rep3', 'PEX14') 2287 1719
('rep1', 'PML') 1618 2278
('rep2', 'PML') 2213 1348
('rep3', 'PML') 1723 1767
('rep1', 'PSD95') 1375 1792
('rep2', 'PSD95') 1939 1865
('rep3', 'PSD95') 2087 1413
('rep1', 'PURA') 695 689
('rep2', 'PURA') 665 648
('rep3', 'PURA') 490 402
('rep1', 'Phalloidin') 1375 1792
('rep2', 'Phalloidin') 1939 1865
('rep3', 'Phalloidin') 2087 1413
('rep1', 'SNCA') 2115 1317
('rep2', 'SNCA') 1884 1642
('rep3', 'SNCA') 2020 1562
('rep1', 'SON') 1862 1685
('rep2', 'SON') 1956 1870
('rep3', 'SON') 1798 1690
('rep1', 'SQSTM1') 1771 1776
('rep2', 'SQSTM1') 1527 1773
('rep3', 'SQSTM1') 1855 1617
('rep1', 'TDP43') 1618 2278
('rep2', 'TDP43') 2213 1348
('rep3', 'TDP43') 1723 1767
('rep1', 'TIA1') 2443 1414
('rep2', 'TIA1') 1171 1898
('rep3', 'TIA1') 1480 1485
('rep1', 'TOMM20') 2443 1434
('rep2', 'TOMM20') 1176 1898
('rep3', 'TOMM20') 1480 1503
('rep1', 'Tubulin') 1789 1800
('rep2', 'Tubulin') 1544 1773
('rep3', 'Tubulin') 1864 1651
('rep1', 'mitotracker') 2443 1434
('rep2', 'mitotracker') 1176 1898
('rep3', 'mitotracker') 1480 1503
Total number of tiles
2250
2000
1750
1500
1250
1000
750
500

### Batch 5

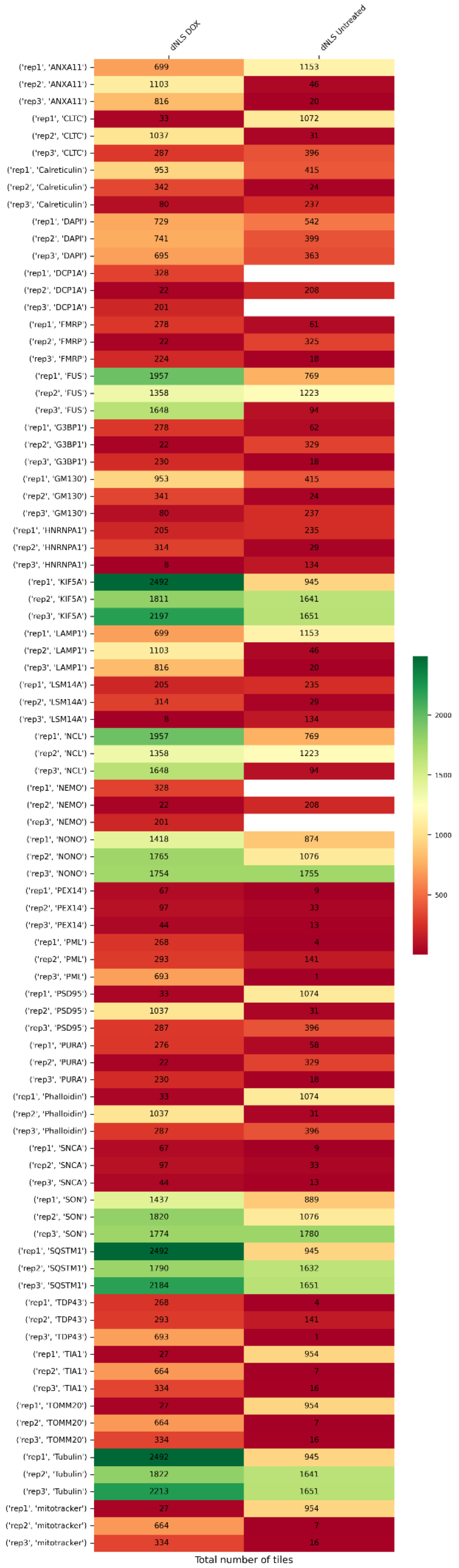
dNLS DOX
dNLS Untreated
('rep1', 'ANXA11') 699 1153
('rep2', 'ANXA11') 1103 46
('rep3', 'ANXA11') 816 20
('rep1', 'CLTC') 33 1072
('rep2', 'CLTC') 1037 31
('rep3', 'CLTC') 287 396
('rep1', 'Calreticulin') 953 415
('rep2', 'Calreticulin') 342 24
('rep3', 'Calreticulin') 80 237
('rep1', 'DAPI') 729 542
('rep2', 'DAPI') 741 399
('rep3', 'DAPI') 695 363
('rep1', 'DCP1A') 328
('rep2', 'DCP1A') 22 208
('rep3', 'DCP1A') 201
('rep1', 'FMRP') 278 61
('rep2', 'FMRP') 22 325
('rep3', 'FMRP') 224 18
('rep1', 'FUS') 1957 769
('rep2', 'FUS') 1358 1223
('rep3', 'FUS') 1648 94
('rep1', 'G3BP1') 278 62
('rep2', 'G3BP1') 22 329
('rep3', 'G3BP1') 230 18
('rep1', 'GM130') 953 415
('rep2', 'GM130') 341 24
('rep3', 'GM130') 80 237
('rep1', 'HNRNPA1') 205 235
('rep2', 'HNRNPA1') 314 29
('rep3', 'HNRNPA1') 8 134
('rep1', 'KIF5A') 2492 945
('rep2', 'KIF5A') 1811 1641
('rep3', 'KIF5A') 2197 1651
('rep1', 'LAMP1') 699 1153
('rep2', 'LAMP1') 1103 46
('rep3', 'LAMP1') 816 20
('rep1', 'LSM14A') 205 235
('rep2', 'LSM14A') 314 29
('rep3', 'LSM14A') 8 134
('rep1', 'NCL') 1957 769
('rep2', 'NCL') 1358 1223
('rep3', 'NCL') 1648 94
('rep1', 'NEMO') 328
('rep2', 'NEMO') 22 208
('rep3', 'NEMO') 201
('rep1', 'NONO') 1418 874
('rep2', 'NONO') 1765 1076
('rep3', 'NONO') 1754 1755
('rep1', 'PEX14') 67 9
('rep2', 'PEX14') 97 33
('rep3', 'PEX14') 44 13
('rep1', 'PML') 268 4
('rep2', 'PML') 293 141
('rep3', 'PML') 693 1
('rep1', 'PSD95') 33 1074
('rep2', 'PSD95') 1037 31
('rep3', 'PSD95') 287 396
('rep1', 'PURA') 276 58
('rep2', 'PURA') 22 329
('rep3', 'PURA') 230 18
('rep1', 'Phalloidin') 33 1074
('rep2', 'Phalloidin') 1037 31
('rep3', 'Phalloidin') 287 396
('rep1', 'SNCA') 67 9
('rep2', 'SNCA') 97 33
('rep3', 'SNCA') 44 13
('rep1', 'SON') 1437 889
('rep2', 'SON') 1820 1076
('rep3', 'SON') 1774 1780
('rep1', 'SQSTM1') 2492 945
('rep2', 'SQSTM1') 1790 1632
('rep3', 'SQSTM1') 2184 1651
('rep1', 'TDP43') 268 4
('rep2', 'TDP43') 293 141
('rep3', 'TDP43') 693 1
('rep1', 'TIA1') 27 954
('rep2', 'TIA1') 664 7
('rep3', 'TIA1') 334 16
('rep1', 'TOMM20') 27 954
('rep2', 'TOMM20') 664 7
('rep3', 'TOMM20') 334 16
('rep1', 'Tubulin') 2492 945
('rep2', 'Tubulin') 1822 1641
('rep3', 'Tubulin') 2213 1651
('rep1', 'mitotracker') 27 954
('rep2', 'mitotracker') 664 7
('rep3', 'mitotracker') 334 16
2000
1500
1000
500
Total number of tiles

### Whole Cell Count

#### Batch 1

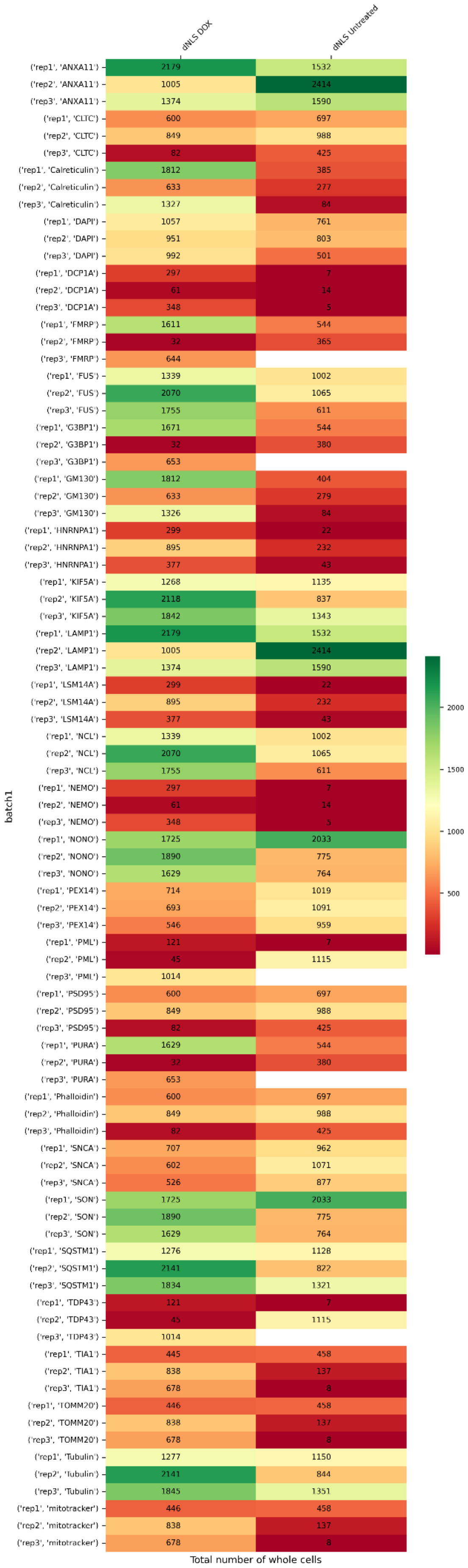

<u>Batch 2</u>

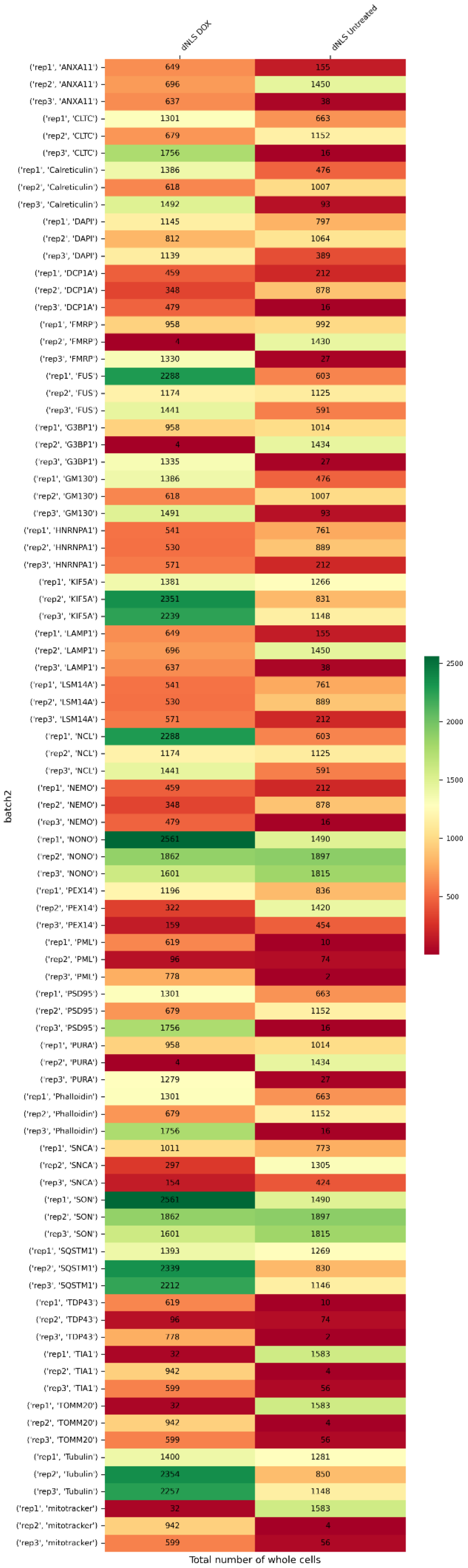
dNLS DOX
dNLS Untreated
batch2
('rep1', 'ANXA11') 649 155
('rep2', 'ANXA11') 696 1450
('rep3', 'ANXA11') 637 38
('rep1', 'CLTC') 1301 663
('rep2', 'CLTC') 679 1152
('rep3', 'CLTC') 1756 16
('rep1', 'Calreticulin') 1386 476
('rep2', 'Calreticulin') 618 1007
('rep3', 'Calreticulin') 1492 93
('rep1', 'DAPI') 1145 797
('rep2', 'DAPI') 812 1064
('rep3', 'DAPI') 1139 389
('rep1', 'DCP1A') 459 212
('rep2', 'DCP1A') 348 878
('rep3', 'DCP1A') 479 16
('rep1', 'FMRP') 958 992
('rep2', 'FMRP') 4 1430
('rep3', 'FMRP') 1330 27
('rep1', 'FUS') 2288 603
('rep2', 'FUS') 1174 1125
('rep3', 'FUS') 1441 591
('rep1', 'G3BP1') 958 1014
('rep2', 'G3BP1') 4 1434
('rep3', 'G3BP1') 1335 27
('rep1', 'GM130') 1386 476
('rep2', 'GM130') 618 1007
('rep3', 'GM130') 1491 93
('rep1', 'HNRNPA1') 541 761
('rep2', 'HNRNPA1') 530 889
('rep3', 'HNRNPA1') 571 212
('rep1', 'KIF5A') 1381 1266
('rep2', 'KIF5A') 2351 831
('rep3', 'KIF5A') 2239 1148
('rep1', 'LAMP1') 649 155
('rep2', 'LAMP1') 696 1450
('rep3', 'LAMP1') 637 38
('rep1', 'LSM14A') 541 761
('rep2', 'LSM14A') 530 889
('rep3', 'LSM14A') 571 212
('rep1', 'NCL') 2288 603
('rep2', 'NCL') 1174 1125
('rep3', 'NCL') 1441 591
('rep1', 'NEMO') 459 212
('rep2', 'NEMO') 348 878
('rep3', 'NEMO') 479 16
('rep1', 'NONO') 2561 1490
('rep2', 'NONO') 1862 1897
('rep3', 'NONO') 1601 1815
('rep1', 'PEX14') 1196 836
('rep2', 'PEX14') 322 1420
('rep3', 'PEX14') 159 454
('rep1', 'PML') 619 10
('rep2', 'PML') 96 74
('rep3', 'PML') 778 2
('rep1', 'PSD95') 1301 663
('rep2', 'PSD95') 679 1152
('rep3', 'PSD95') 1756 16
('rep1', 'PURA') 958 1014
('rep2', 'PURA') 4 1434
('rep3', 'PURA') 1279 27
('rep1', 'Phalloidin') 1301 663
('rep2', 'Phalloidin') 679 1152
('rep3', 'Phalloidin') 1756 16
('rep1', 'SNCA') 1011 773
('rep2', 'SNCA') 297 1305
('rep3', 'SNCA') 154 424
('rep1', 'SON') 2561 1490
('rep2', 'SON') 1862 1897
('rep3', 'SON') 1601 1815
('rep1', 'SQSTM1') 1393 1269
('rep2', 'SQSTM1') 2339 830
('rep3', 'SQSTM1') 2212 1146
('rep1', 'TDP43') 619 10
('rep2', 'TDP43') 96 74
('rep3', 'TDP43') 778 2
('rep1', 'TIA1') 32 1583
('rep2', 'TIA1') 942 4
('rep3', 'TIA1') 599 56
('rep1', 'TOMM20') 32 1583
('rep2', 'TOMM20') 942 4
('rep3', 'TOMM20') 599 56
('rep1', 'Tubulin') 1400 1281
('rep2', 'Tubulin') 2354 850
('rep3', 'Tubulin') 2257 1148
('rep1', 'mitotracker') 32 1583
('rep2', 'mitotracker') 942 4
('rep3', 'mitotracker') 599 56
Total number of whole cells
2500
2000
1500
1000
500

### Batch 3

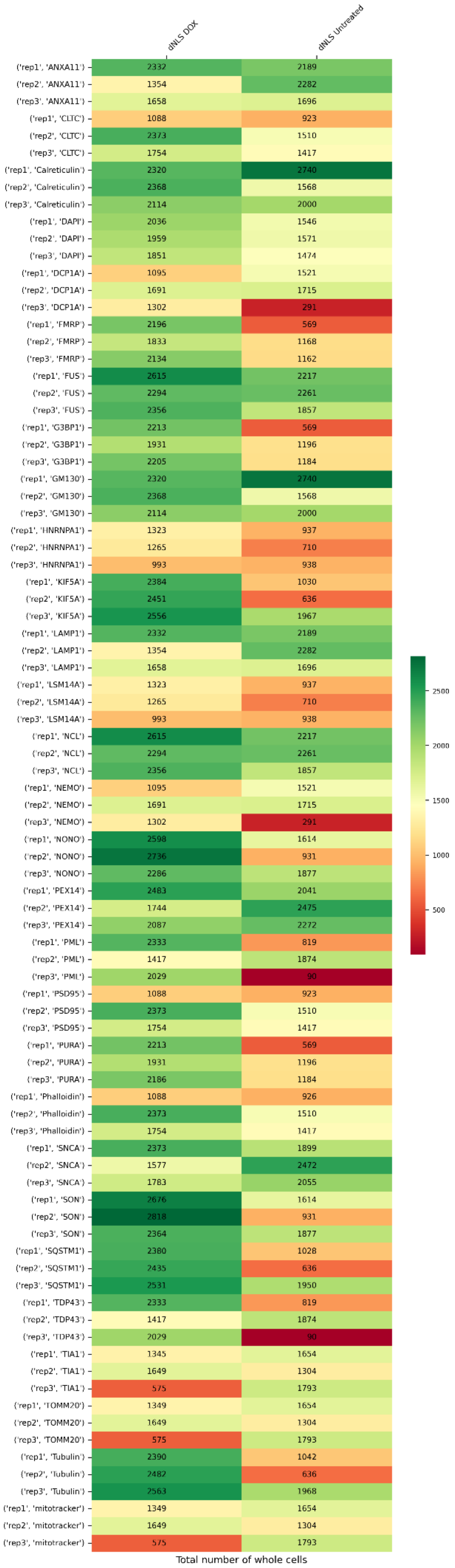
dNLS DOX
dNLS Untreated
('rep1', 'ANXA11') 2332 2189
('rep2', 'ANXA11') 1354 2282
('rep3', 'ANXA11') 1658 1696
('rep1', 'CLTC') 1088 923
('rep2', 'CLTC') 2373 1510
('rep3', 'CLTC') 1754 1417
('rep1', 'Calreticulin') 2320 2740
('rep2', 'Calreticulin') 2368 1568
('rep3', 'Calreticulin') 2114 2000
('rep1', 'DAPI') 2036 1546
('rep2', 'DAPI') 1959 1571
('rep3', 'DAPI') 1851 1474
('rep1', 'DCP1A') 1095 1521
('rep2', 'DCP1A') 1691 1715
('rep3', 'DCP1A') 1302 291
('rep1', 'FMRP') 2196 569
('rep2', 'FMRP') 1833 1168
('rep3', 'FMRP') 2134 1162
('rep1', 'FUS') 2615 2217
('rep2', 'FUS') 2294 2261
('rep3', 'FUS') 2356 1857
('rep1', 'G3BP1') 2213 569
('rep2', 'G3BP1') 1931 1196
('rep3', 'G3BP1') 2205 1184
('rep1', 'GM130') 2320 2740
('rep2', 'GM130') 2368 1568
('rep3', 'GM130') 2114 2000
('rep1', 'HNRNPA1') 1323 937
('rep2', 'HNRNPA1') 1265 710
('rep3', 'HNRNPA1') 993 938
('rep1', 'KIF5A') 2384 1030
('rep2', 'KIF5A') 2451 636
('rep3', 'KIF5A') 2556 1967
('rep1', 'LAMP1') 2332 2189
('rep2', 'LAMP1') 1354 2282
('rep3', 'LAMP1') 1658 1696
('rep1', 'LSM14A') 1323 937
('rep2', 'LSM14A') 1265 710
('rep3', 'LSM14A') 993 938
('rep1', 'NCL') 2615 2217
('rep2', 'NCL') 2294 2261
('rep3', 'NCL') 2356 1857
('rep1', 'NEMO') 1095 1521
('rep2', 'NEMO') 1691 1715
('rep3', 'NEMO') 1302 291
('rep1', 'NONO') 2598 1614
('rep2', 'NONO') 2736 931
('rep3', 'NONO') 2286 1877
('rep1', 'PEX14') 2483 2041
('rep2', 'PEX14') 1744 2475
('rep3', 'PEX14') 2087 2272
('rep1', 'PML') 2333 819
('rep2', 'PML') 1417 1874
('rep3', 'PML') 2029 90
('rep1', 'PSD95') 1088 923
('rep2', 'PSD95') 2373 1510
('rep3', 'PSD95') 1754 1417
('rep1', 'PURA') 2213 569
('rep2', 'PURA') 1931 1196
('rep3', 'PURA') 2186 1184
('rep1', 'Phalloidin') 1088 926
('rep2', 'Phalloidin') 2373 1510
('rep3', 'Phalloidin') 1754 1417
('rep1', 'SNCA') 2373 1899
('rep2', 'SNCA') 1577 2472
('rep3', 'SNCA') 1783 2055
('rep1', 'SON') 2676 1614
('rep2', 'SON') 2818 931
('rep3', 'SON') 2364 1877
('rep1', 'SQSTM1') 2380 1028
('rep2', 'SQSTM1') 2435 636
('rep3', 'SQSTM1') 2531 1950
('rep1', 'TDP43') 2333 819
('rep2', 'TDP43') 1417 1874
('rep3', 'TDP43') 2029 90
('rep1', 'TIA1') 1345 1654
('rep2', 'TIA1') 1649 1304
('rep3', 'TIA1') 575 1793
('rep1', 'TOMM20') 1349 1654
('rep2', 'TOMM20') 1649 1304
('rep3', 'TOMM20') 575 1793
('rep1', 'Tubulin') 2390 1042
('rep2', 'Tubulin') 2482 636
('rep3', 'Tubulin') 2563 1968
('rep1', 'mitotracker') 1349 1654
('rep2', 'mitotracker') 1649 1304
('rep3', 'mitotracker') 575 1793
2500
2000
1500
1000
500
Total number of whole cells

### Batch 4

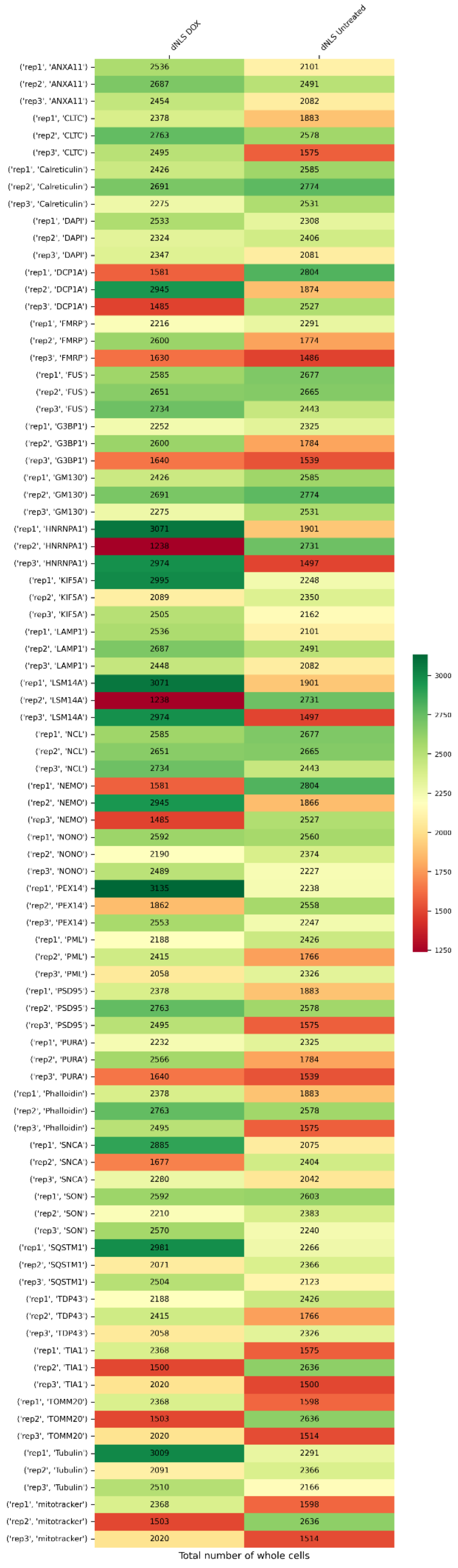

dNLS DOX
dNLS Untreated
('rep1', 'ANXA11') 2536 2101
('rep2', 'ANXA11') 2687 2491
('rep3', 'ANXA11') 2454 2082
('rep1', 'CLTC') 2378 1883
('rep2', 'CLTC') 2763 2578
('rep3', 'CLTC') 2495 1575
('rep1', 'Calreticulin') 2426 2585
('rep2', 'Calreticulin') 2691 2774
('rep3', 'Calreticulin') 2275 2531
('rep1', 'DAPI') 2533 2308
('rep2', 'DAPI') 2324 2406
('rep3', 'DAPI') 2347 2081
('rep1', 'DCP1A') 1581 2804
('rep2', 'DCP1A') 2945 1874
('rep3', 'DCP1A') 1485 2527
('rep1', 'FMRP') 2216 2291
('rep2', 'FMRP') 2600 1774
('rep3', 'FMRP') 1630 1486
('rep1', 'FUS') 2585 2677
('rep2', 'FUS') 2651 2665
('rep3', 'FUS') 2734 2443
('rep1', 'G3BP1') 2252 2325
('rep2', 'G3BP1') 2600 1784
('rep3', 'G3BP1') 1640 1539
('rep1', 'GM130') 2426 2585
('rep2', 'GM130') 2691 2774
('rep3', 'GM130') 2275 2531
('rep1', 'HNRNPA1') 3071 1901
('rep2', 'HNRNPA1') 1238 2731
('rep3', 'HNRNPA1') 2974 1497
('rep1', 'KIF5A') 2995 2248
('rep2', 'KIF5A') 2089 2350
('rep3', 'KIF5A') 2505 2162
('rep1', 'LAMP1') 2536 2101
('rep2', 'LAMP1') 2687 2491
('rep3', 'LAMP1') 2448 2082
('rep1', 'LSM14A') 3071 1901
('rep2', 'LSM14A') 1238 2731
('rep3', 'LSM14A') 2974 1497
('rep1', 'NCL') 2585 2677
('rep2', 'NCL') 2651 2665
('rep3', 'NCL') 2734 2443
('rep1', 'NEMO') 1581 2804
('rep2', 'NEMO') 2945 1866
('rep3', 'NEMO') 1485 2527
('rep1', 'NONO') 2592 2560
('rep2', 'NONO') 2190 2374
('rep3', 'NONO') 2489 2227
('rep1', 'PEX14') 3135 2238
('rep2', 'PEX14') 1862 2558
('rep3', 'PEX14') 2553 2247
('rep1', 'PML') 2188 2426
('rep2', 'PML') 2415 1766
('rep3', 'PML') 2058 2326
('rep1', 'PSD95') 2378 1883
('rep2', 'PSD95') 2763 2578
('rep3', 'PSD95') 2495 1575
('rep1', 'PURA') 2232 2325
('rep2', 'PURA') 2566 1784
('rep3', 'PURA') 1640 1539
('rep1', 'Phalloidin') 2378 1883
('rep2', 'Phalloidin') 2763 2578
('rep3', 'Phalloidin') 2495 1575
('rep1', 'SNCA') 2885 2075
('rep2', 'SNCA') 1677 2404
('rep3', 'SNCA') 2280 2042
('rep1', 'SON') 2592 2603
('rep2', 'SON') 2210 2383
('rep3', 'SON') 2570 2240
('rep1', 'SQSTM1') 2981 2266
('rep2', 'SQSTM1') 2071 2366
('rep3', 'SQSTM1') 2504 2123
('rep1', 'TDP43') 2188 2426
('rep2', 'TDP43') 2415 1766
('rep3', 'TDP43') 2058 2326
('rep1', 'TIA1') 2368 1575
('rep2', 'TIA1') 1500 2636
('rep3', 'TIA1') 2020 1500
('rep1', 'TOMM20') 2368 1598
('rep2', 'TOMM20') 1503 2636
('rep3', 'TOMM20') 2020 1514
('rep1', 'Tubulin') 3009 2291
('rep2', 'Tubulin') 2091 2366
('rep3', 'Tubulin') 2510 2166
('rep1', 'mitotracker') 2368 1598
('rep2', 'mitotracker') 1503 2636
('rep3', 'mitotracker') 2020 1514
3000
2750
2500
2250
2000
1750
1500
1250
Total number of whole cells

#### Batch 5

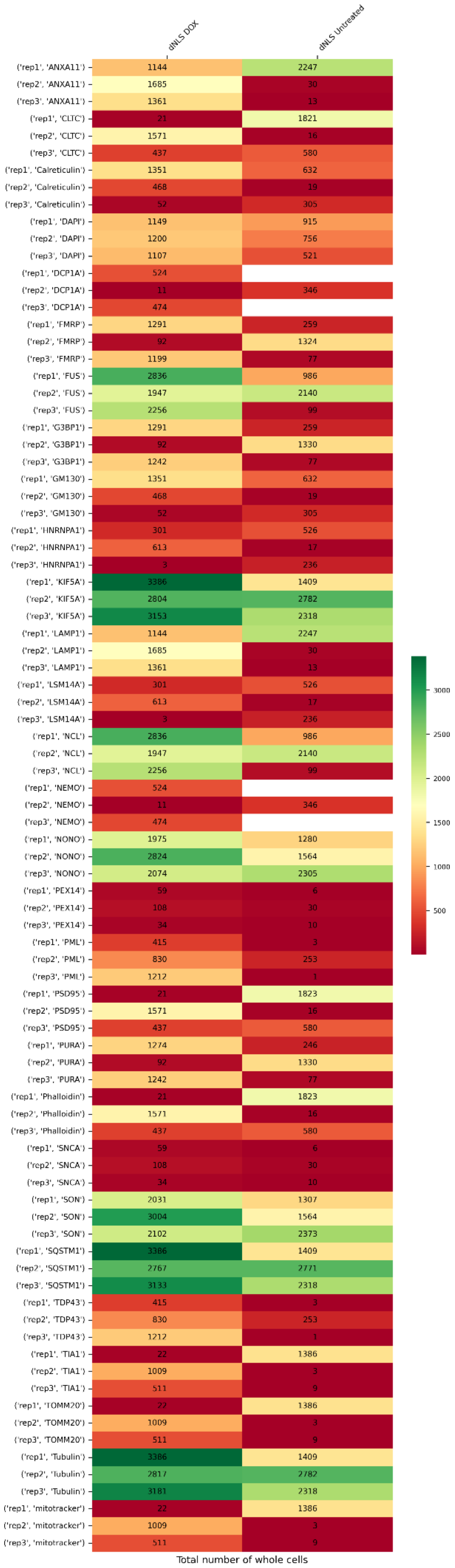
dNLS DOX
dNLS Untreated
('rep1', 'ANXA11') 1144 2247
('rep2', 'ANXA11') 1685 30
('rep3', 'ANXA11') 1361 13
('rep1', 'CLTC') 21 1821
('rep2', 'CLTC') 1571 16
('rep3', 'CLTC') 437 580
('rep1', 'Calreticulin') 1351 632
('rep2', 'Calreticulin') 468 19
('rep3', 'Calreticulin') 52 305
('rep1', 'DAPI') 1149 915
('rep2', 'DAPI') 1200 756
('rep3', 'DAPI') 1107 521
('rep1', 'DCP1A') 524
('rep2', 'DCP1A') 11 346
('rep3', 'DCP1A') 474
('rep1', 'FMRP') 1291 259
('rep2', 'FMRP') 92 1324
('rep3', 'FMRP') 1199 77
('rep1', 'FUS') 2836 986
('rep2', 'FUS') 1947 2140
('rep3', 'FUS') 2256 99
('rep1', 'G3BP1') 1291 259
('rep2', 'G3BP1') 92 1330
('rep3', 'G3BP1') 1242 77
('rep1', 'GM130') 1351 632
('rep2', 'GM130') 468 19
('rep3', 'GM130') 52 305
('rep1', 'HNRNPA1') 301 526
('rep2', 'HNRNPA1') 613 17
('rep3', 'HNRNPA1') 3 236
('rep1', 'KIF5A') 3386 1409
('rep2', 'KIF5A') 2804 2782
('rep3', 'KIF5A') 3153 2318
('rep1', 'LAMP1') 1144 2247
('rep2', 'LAMP1') 1685 30
('rep3', 'LAMP1') 1361 13
('rep1', 'LSM14A') 301 526
('rep2', 'LSM14A') 613 17
('rep3', 'LSM14A') 3 236
('rep1', 'NCL') 2836 986
('rep2', 'NCL') 1947 2140
('rep3', 'NCL') 2256 99
('rep1', 'NEMO') 524
('rep2', 'NEMO') 11 346
('rep3', 'NEMO') 474
('rep1', 'NONO') 1975 1280
('rep2', 'NONO') 2824 1564
('rep3', 'NONO') 2074 2305
('rep1', 'PEX14') 59 6
('rep2', 'PEX14') 108 30
('rep3', 'PEX14') 34 10
('rep1', 'PML') 415 3
('rep2', 'PML') 830 253
('rep3', 'PML') 1212 1
('rep1', 'PSD95') 21 1823
('rep2', 'PSD95') 1571 16
('rep3', 'PSD95') 437 580
('rep1', 'PURA') 1274 246
('rep2', 'PURA') 92 1330
('rep3', 'PURA') 1242 77
('rep1', 'Phalloidin') 21 1823
('rep2', 'Phalloidin') 1571 16
('rep3', 'Phalloidin') 437 580
('rep1', 'SNCA') 59 6
('rep2', 'SNCA') 108 30
('rep3', 'SNCA') 34 10
('rep1', 'SON') 2031 1307
('rep2', 'SON') 3004 1564
('rep3', 'SON') 2102 2373
('rep1', 'SQSTM1') 3386 1409
('rep2', 'SQSTM1') 2767 2771
('rep3', 'SQSTM1') 3133 2318
('rep1', 'TDP43') 415 3
('rep2', 'TDP43') 830 253
('rep3', 'TDP43') 1212 1
('rep1', 'TIA1') 22 1386
('rep2', 'TIA1') 1009 3
('rep3', 'TIA1') 511 9
('rep1', 'TOMM20') 22 1386
('rep2', 'TOMM20') 1009 3
('rep3', 'TOMM20') 511 9
('rep1', 'Tubulin') 3386 1409
('rep2', 'Tubulin') 2817 2782
('rep3', 'Tubulin') 3181 2318
('rep1', 'mitotracker') 22 1386
('rep2', 'mitotracker') 1009 3
('rep3', 'mitotracker') 511 9
3000
2500
2000
1500
1000
500
Total number of whole cells
